# Benchmarking generative AI tools for literature retrieval and summarization in genomic variant interpretation

**DOI:** 10.1101/2025.09.29.679212

**Authors:** Andrea Gazzo, Silvia Berardelli, Matteo Biancospino, Lorenzo Cuollo, Flavia Dei Zotti, Emanuela Ferraro, Antonio Marra, Enrico Tartarotti, Paolo Magni

## Abstract

**Background:** Generative AI is increasingly used to extract structured information across domains, but its reliability in academic and clinical research, where precision and accuracy are essential, remains largely unexplored. This study evaluates the ability of Large Language Models (LLMs)-based algorithms to generate accurate, literature-based summaries of human genomic variants, with a focus on real-world usability.

**Results:** We benchmarked five open-access generative AI platforms—ChatGPT, MistralAI, VarChat, Perplexity, and ScholarAI—across 40 curated variants equally divided between somatic and germline settings. For each variant, summary reports were generated and blindly evaluated by domain experts using five defined metrics. VarChat emerged as the top-ranked tool, showing the highest summarization accuracy, citation relevance, and robustness against hallucinations. Gpt-4o consistently ranked second, showing particularly stable robustness in conditions where the literature was scarce. Perplexity and ScholarAI, despite being literature-focused, ranked lowest across most metrics. Tool performance was strongly influenced by the availability of peer-reviewed literature, confirming that current generative models remain sensitive to data scarcity.

**Conclusions:** Our findings highlight the heterogeneity of current generative AI tools in genomic variant interpretation workflows. While some platforms already provide useful outputs, reliable integration into basic and clinical research requires expert validation and domain-related fine-tuning. This work provides for the first time a curated benchmark for assessing LLM-generated content in variant genomics and underscores the need for caution when using these tools to support variant interpretation.

## Background

Peer-reviewed literature analysis plays a crucial role in guiding scientific research. In genomic research, for instance, the interpretation of genetic variants is particularly relevant, as it can potentially guide not only basic research but ultimately also diagnostic and therapeutic decisions [1]. The integration of knowledge produced by a wide range of analyses, including case studies, functional assays, population studies, and clinical observations, allows researchers and clinicians to better understand the impact of genetic variants and to guide research.

Indeed, the exponential growth of peer-reviewed publications, including those related to human genetic variants, has accumulated a vast amount of knowledge on variants and their association with diseases. At the same time, this drastic accumulation has made it increasingly difficult and time-consuming for clinicians and researchers to efficiently retrieve precise information focused on specific variants [2].

This data overload raises several issues, not only related to the lengthy and time-consuming process of manually extracting customized information from the literature, but also to the risk of overlooking critical evidence that may influence interpretation.

To tackle this problem, several automated or semi-automated tools have been developed to facilitate the retrieval of variant-specific scientific literature. These tools help filter, organize, and summarize the expanding literature corpus [3][4].

LitVar2 retrieves variant-related publications from PubMed and PMC, enriched with disease and drug annotations from PubTator 3.0, which uses tmVar3 for variant recognition and normalization [5–7]. Variomes supports customized searches with historical filters and keyword prioritization [8]. DeepVar applies an end-to-end neural network to identify genomic variants without feature engineering or post-processing [9], while AVADA uses machine learning to extract pathogenic variant evidence from full-text literature [10]. Mastermind, a commercial platform, compiles insights on over 27 million variants [11].

These platforms typically rely on Natural Language Processing (NLP) pipelines [12] and, in some cases, integrate a manual curation step. These tools retrieve the pertinent literature but still require manual synthesis, creating a need for automated, accurate summarization. Generative Artificial Intelligence (AI) tools, particularly Large Language Models (LLMs), have recently emerged as powerful engines for literature summarization across multiple domains, including biomedicine [13]. LLMs are based on Transformer architectures that leverage attention mechanisms to capture relationships across long text sequences [14].

These tools can be broadly categorized into general-purpose models - as ChatGPT or MistralAI -, and literature-based models, such as ScholarAI and VarChat. General-purpose LLMs rely on broad linguistic patterns and reasoning capabilities, offering flexibility but often lacking domain-specific accuracy. In contrast, literature-based models are explicitly designed to retrieve and process scientific articles from databases such as PubMed, PMC, or equivalent databases [15].

Despite their potential, the application of LLMs to genomics remains underexplored. A major limitation is hallucination, where a model fabricates or misrepresents information [16]. In variant interpretation, hallucinations may arise in three different ways: first, incorrect or inconsistent representation of variant nomenclature, gene-disease associations or functional annotations, which can misinform clinical or research decisions. Such inaccuracies can lead to misclassification of pathogenicity, inconsistent evidence presentation, or incorrect inference of variant effects [17]. Secondly, LLMs may also produce pure hallucinations, inventing gene-variant relationships or functional effects unsupported by evidence. Finally, their performance can be biased for rare or underrepresented variants, especially when training data are limited, literature is sparse, or key sources are available only in non-English languages [18]. Moreover, hallucinations may also relate to the literature cited and referenced, which is often inconsistent, inaccurate, or entirely fabricated by the LLMs [19]. To address these concerns, it is essential to evaluate the reliability of generative AI tools in extracting and synthesizing genomic variant-specific information from peer-reviewed literature.

Human experts remain the gold standard for evaluating LLM-generated content [20], but their involvement is time-consuming. As a faster alternative, some studies use independent LLMs for automated benchmark evaluations [21, 22], though these inherit the same limitations as the models they assess and still require human validation [23, 24]. Other work evaluates LLMs on clinical questions with expert review, such as neurologists assessing response quality [25]. Mixed approaches also exist, combining NLP-based assessments of syntax, semantics, and similarity with manual review by clinical staff [26]. In genomics, we believe accurate evaluation ultimately depends on review by selected experts.

Similar studies have been conducted in the biomedical field, where the summarization ability of LLMs has been evaluated. In [27], experts manually evaluate comprehensiveness, readability and faithfulness of five different LLMs on biomedical datasets. In [28], authors investigated the performance of ChatGPT in several biomedical NLP tasks, including entity recognition and paragraph summarization. Furthermore, in [29] authors evaluate the effectiveness of general-purpose and domain-specific LLMs on a large-scale biomedical dataset containing 25,000 records acquired from PubMed in diverse medical domains. The tool LitSumm is another example of application for LLMs summarization of genomic information with focus on RNA, with a manually evaluation made by four expert raters [30]. Finally, [31] presents a systematic comparison between LLM-generated and human-authored clinical reviews.

In this study, we benchmark the performance of five generative AI platforms in producing detailed summaries for 40 curated genomic variants, equally divided between germline and somatic contexts. Each output was blindly evaluated by expert reviewers using five defined metrics, capturing both content quality and usability. The dataset was deliberately constructed to include both germline and somatic variants, reflecting the two main domains where variant interpretation is applied, which often share similarities but also differ substantially. Although the benchmark is limited in size, it spans variants with high, medium, and low literature support, includes well-established and rare pathogenic mutations as well as neutral controls, and thus remains broadly representative of real-world use cases. This evaluation aims to provide a robust assessment of LLM capabilities in supporting variant summarization workflows, for the first time to our knowledge.

## Results

### Evaluation framework and performance on the whole dataset

To evaluate whether generative AI tools can be effectively used to retrieve relevant biological and clinical information on genetic variants from peer-reviewed literature, we benchmarked five platforms across 40 curated variants (see Methods). Each output was scored by expert reviewers using five predefined metrics capturing both content quality and usability. An overview of the workflow and evaluation design is shown in **Figure 1**.

**Figure 1.**
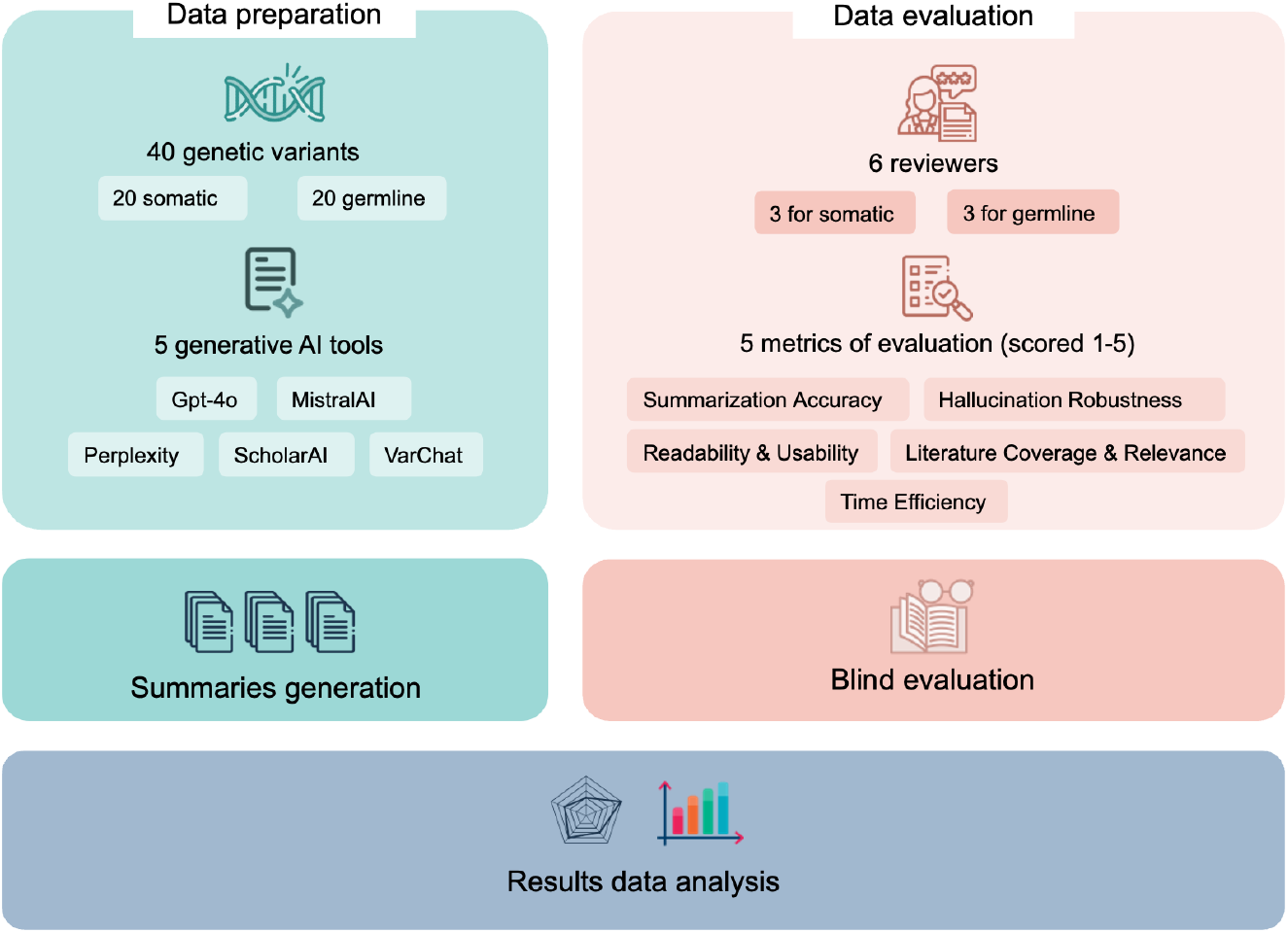
Study workflow and evaluation design. Overview of the benchmarking workflow, including tool input, variant sets (germline and somatic), reviewer distribution, and evaluation metrics.

Aggregated results across all 40 variants and six reviewers (**Figure 2a**) indicate that VarChat achieved the highest Summarization Accuracy score (4.33 ± 0.74, mean ± SD), followed by Gpt-4o (3.74 ± 0.82), MistralAI (3.62 ± 0.85), ScholarAI (3.35 ± 0.94), and Perplexity (3.22 ± 0.96). Hallucination Robustness scores were generally high across all tools (range: 3.92–4.62), with VarChat and Gpt-4o being the least prone to hallucinations (4.62 ± 0.68 and 4.29 ± 0.85, respectively). The lowest-performing tool in this category was Perplexity (3.92 ± 1.10), as reflected in reviewer comments noting occasional inconsistencies in variant descriptions and supporting references. Readability & Usability followed a similar pattern, with VarChat and Gpt-4o again leading in performance (4.33 ± 0.78 and 3.76 ± 0.73, respectively). At the lower end of the ranking, Perplexity, ScholarAI, and MistralAI scored between 3.2 and 3.6, indicating outputs that reviewers often judged less clear and less practical for immediate use. Across all 25 tool–metric pairs, SDs ranged from 0.68 to 1.14 (median ∼0.99), corresponding to 17–29% of the 1– 5 scale range. This reflects moderate inter-reviewer variability, which is expected in subjective scientific extraction tasks of this complexity, and remains within an acceptable range. Literature Coverage & Relevance was the most variable metric (mean SD: 1.07), while Summarization Accuracy and Readability were more consistently scored (both 0.86). VarChat showed the least scoring variability (0.85), while ScholarAI was the most inconsistent (0.99).

**Figure 2.**
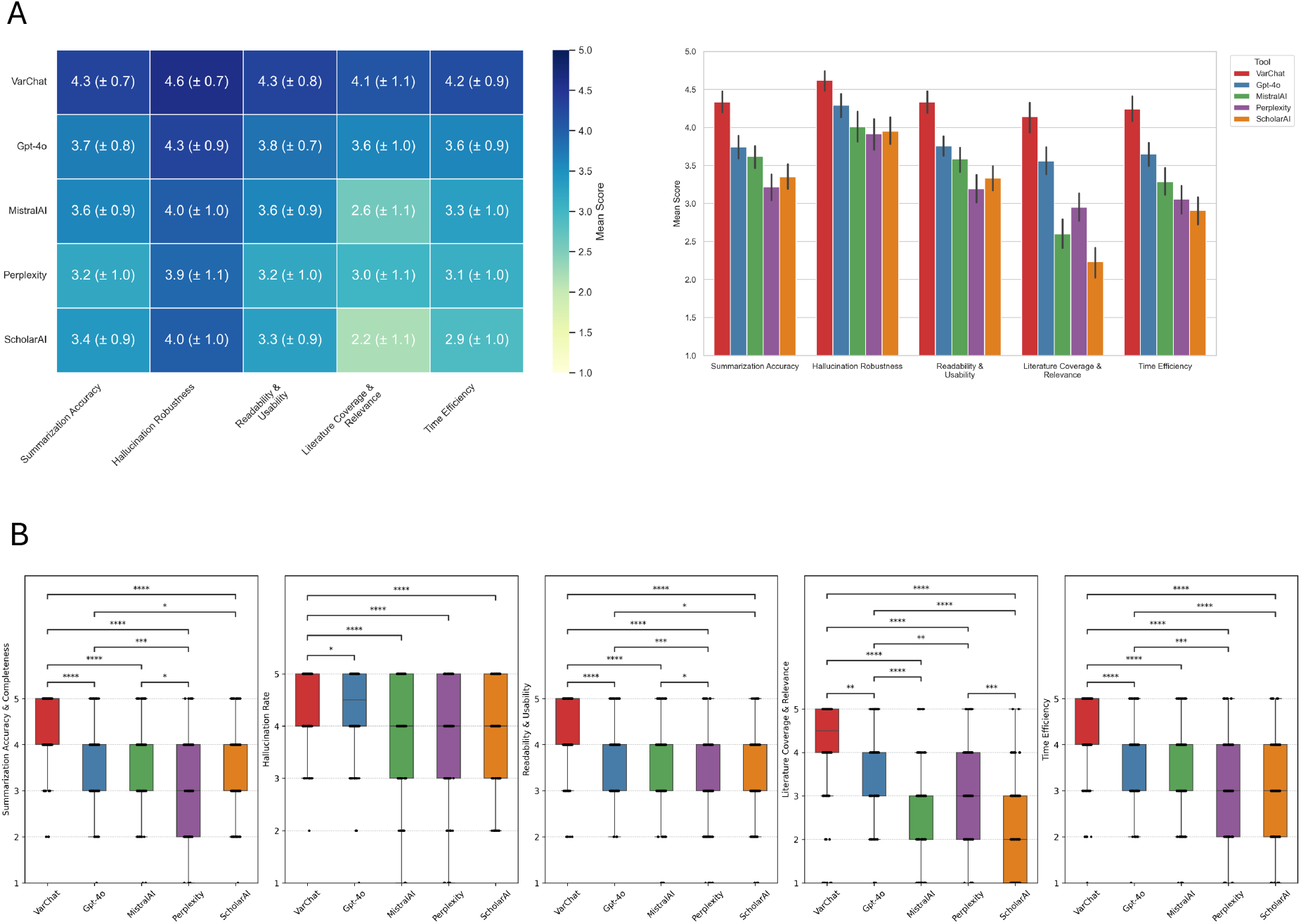
General performance across all variants and metrics. **a**. Left, Heatmap of average scores across five evaluation metrics for all tools (all variants). Right, Bar plots showing mean CI of performance scores per tool and metric. **b**. Boxplots showing distribution of all individual scores per metric and tool; p-values from pairwise comparisons are shown, with significance indicated as p < 0.05 (*), p < 0.01 (**), p < 0.001 (***), and p < 0.0001 (****).

Literature Coverage & Relevance showed an overall drop in performance across all tools relative to the other metrics, although the ranking trend remained similar. Notably, ScholarAI and MistralAI showed a marked decrease in performance in this category. As expected, Time Efficiency scores showed strong correlation with all the other metrics (Spearman ∼ 0.9 in all four pairwise comparisons).

We then compared the tools across all five metrics (**Figure 2b**), observing statistically significant differences (Kruskal p < 0.0001 for all metrics). In Summarization Accuracy, VarChat significantly outperformed all other tools (p = 2.04e-06 vs Gpt-4o, p = 8.09e-09 vs MistralAI, p = 8.95e-19 vs Perplexity, p = 3.79e-16 vs ScholarAI). Gpt-4o also scored significantly higher than ScholarAI (p = 0.0126) and Perplexity (p = 0.00094). The three lowest-performing tools—ScholarAI, Perplexity and MistralAI —did not differ significantly from one another. Significant differences also emerged for Hallucination Robustness (Kruskal p < 0.0001), with VarChat performing significantly better than Perplexity (p = 9.19e-08), MistralAI (p = 4.18e-06), ScholarAI (p = 1.76e-07), and Gpt-4o (p = 0.0255).

No significant differences were observed among the remaining tools (p > 0.05 for all pairwise comparisons not involving VarChat). Readability & Usability and Time Efficiency mirrored the ranking observed for Summarization Accuracy, with VarChat again significantly outperforming all others (p = 2.16e-06 vs Gpt-4o, p = 1.77e-09 vs MistralAI, p = 1.71e-20 vs Perplexity, p = 2.48e-16 vs ScholarAI), and Gpt-4o scoring significantly higher than ScholarAI (p = 0.0103) and Perplexity (p = 0.000145). For Literature Coverage & Relevance, VarChat significantly outperformed all other tools (p = 0.00254 vs Gpt-4o, p = 1.05e-20 vs MistralAI, p = 5.61e-13 vs Perplexity, p = 1.84e-30 vs ScholarAI), followed by Gpt-4o, which also scored significantly higher than ScholarAI (p = 1.14e-14), Perplexity (p = 0.00114), and MistralAI (p = 3.35e-08). Interestingly, ScholarAI—despite being a tool specifically fine-tuned for peer-reviewed scientific content—was the lowest-performing across nearly all metrics, with statistically significant differences compared to all other tools except MistralAI, which ranked as the second-lowest performer.

### Performance on germline variants

In the germline subset (20 variants evaluated by three reviewers), VarChat ranked first across all five metrics, confirming overall strong performance also in this context (**Figure 3a**). It scored highest in Summarization Accuracy (4.27 ± 0.82), followed by MistralAI (3.82 ± 0.72), Gpt-4o (3.53 ± 0.81), ScholarAI (3.48 ± 1.02), and Perplexity (2.97 ± 0.90). VarChat also showed the best Hallucination Robustness (4.73 ± 0.63), with Gpt-4o in second place (4.47 ± 0.85), MistralAI third (4.23 ± 1.01), followed by ScholarAI (4.28 ± 0.99) and Perplexity (4.17 ± 1.01). Readability & Usability ranked VarChat as the highest scored tool (4.37 ± 0.84), followed by MistralAI (3.88 ± 0.85), Gpt-4o (3.77 ± 0.70), ScholarAI (3.65 ± 0.90), and Perplexity (3.15 ± 0.88). For Literature Coverage & Relevance, scores were more variable, with VarChat on top (3.93 ± 1.40), followed by Gpt-4o (3.25 ± 0.93), Perplexity (2.77 ± 0.98), MistralAI (2.65 ± 0.97), and ScholarAI (2.27 ± 1.21). Time Efficiency showed the same trend: VarChat (4.08 ± 1.05) and Gpt-4o (3.47 ± 0.75) as top tools, followed by MistralAI (3.35 ± 0.90), ScholarAI (2.83 ± 1.06), and Perplexity (2.73 ± 0.94).

**Figure 3.**
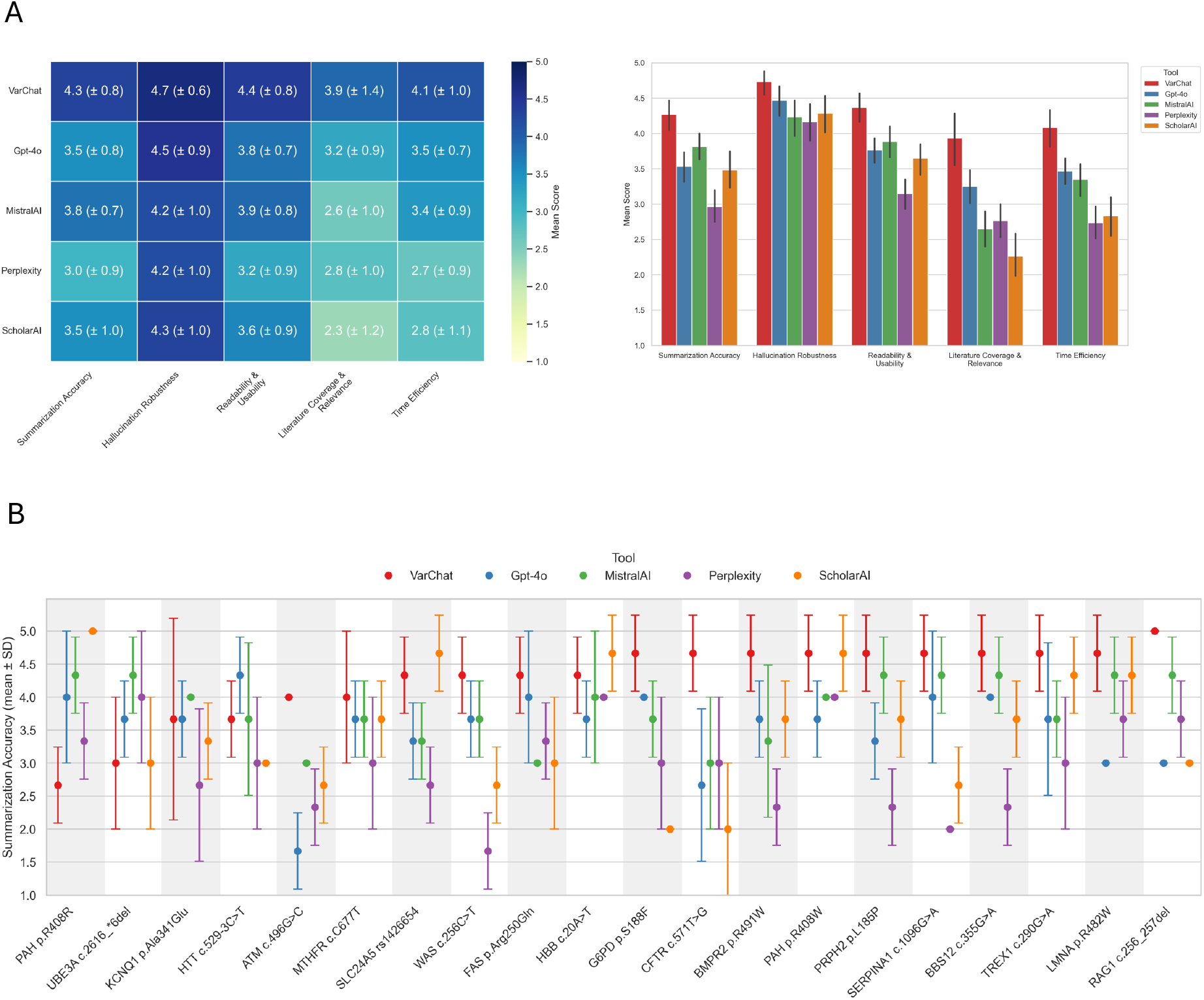
Tool evaluation on germline variants. **a**. Left, Heatmap of average scores across five evaluation metrics for all tools (germline variants). Right, Bar plots showing mean CI of performance scores per tool and metric. **b**. Reviewer score distributions for the 20 germline variants across all tools.

The results in the germline subset are generally very consistent with the overall benchmark. VarChat maintained the top performance across all metrics, with Gpt-4o and MistralAI competing for the second and third positions depending on the metric. The relative underperformance of ScholarAI and Perplexity was also consistent.

VarChat received the highest average score for Summarization Accuracy (4.27), indicating it was most often perceived as the best-performing tool.

In contrast, ScholarAI showed the greatest reviewer consistency (lowest variability), but ranked 4th in mean score (3.48), suggesting that while its outputs were more uniformly interpreted, they were generally rated less favorably. This highlights a potential trade-off between consensus and perceived quality. VarChat was ranked as the best tool for 14 out of 20 germline variants when considering summarization accuracy. The agreement among reviewers was generally acceptable, with a mean STD ranging from 0.525 for ScholarAI to 0.646 for Perplexity. **Figure 3b** shows the reviewer Summarization Accuracy score distributions for all 20 germline variants across tools. The pathogenic variant p.R408W, located in the PAH gene and associated with phenylketonuria, was ranked as the highest on average across all reviewers and tools (mean 4.2). Interestingly, this was also the variant with the strongest inter-reviewer agreement (SD = 0.56). Conversely, c.496G>C in the ATM gene, associated with ataxia-telangiectasia, received the lowest average score (mean 2.73). Despite this, all three reviewers rated VarChat highly for this variant (score 4 or 5), while the lowest-performing tool was Gpt-4o (mean 1.67). Other notable cases included c.571T>G (CFTR) and c.1096G>A (SERPINA1) showed substantial disagreement across both tools and reviewers (SD > 1.18), likely reflecting inconsistencies in evidence accessibility or tool sensitivity to sparse or heterogeneous literature.

### Performance on somatic variants

In the somatic subset (20 variants evaluated by three reviewers), VarChat again ranked first across all five metrics, confirming robust performance also in this context (**Figure 4a**). It scored highest in Summarization Accuracy (4.40 ± 0.64), followed by Gpt-4o (3.95 ± 0.79), Perplexity (3.47 ± 0.96), MistralAI (3.42 ± 0.93), and ScholarAI (3.22 ± 0.85). Hallucination Robustness showed similarly strong results for VarChat (4.50 ± 0.70), with Gpt-4o (4.12 ± 0.83) and MistralAI (3.78 ± 1.03) next in line, while Perplexity (3.67 ± 1.10) and ScholarAI (3.62 ± 0.92) followed closely. Readability & Usability placed VarChat at the top (4.30 ± 0.72), ahead of Gpt-4o (3.75 ± 0.77), MistralAI (3.28 ± 0.88), Perplexity (3.23 ± 1.03), and ScholarAI (3.02 ± 0.79). For Literature Coverage & Relevance, VarChat obtained the highest score (4.35 ± 0.76), followed by Gpt-4o (3.87 ± 1.00), Perplexity (3.13 ± 1.10), MistralAI (2.55 ± 1.19), and ScholarAI (2.20 ± 0.97). Time Efficiency mirrored the general ranking, with VarChat leading (4.40 ± 0.69), followed by Gpt-4o (3.83 ± 0.94), Perplexity (3.38 ± 1.06), MistralAI (3.22 ± 1.08), and ScholarAI (2.98 ± 0.95). Overall, VarChat consistently achieved the highest scores across all metrics, with low variability, confirming its reliability in the somatic setting.

**Figure 4.**
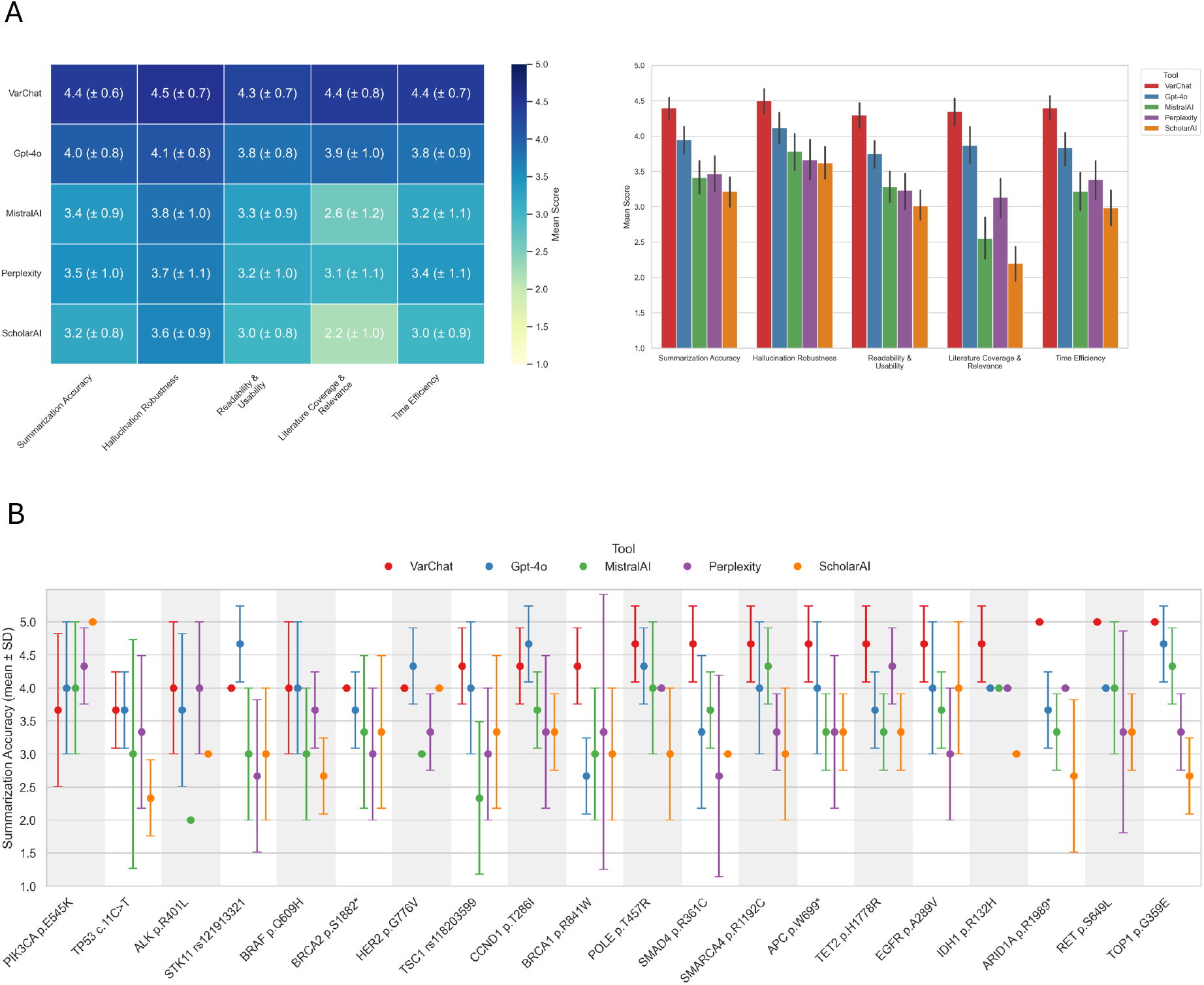
Tool evaluation on somatic variants. **a**. Left, Heatmap of average scores across five evaluation metrics for all tools (somatic variants). Right, Bar plots showing mean CI of performance scores per tool and metric. **b**. Reviewer score distributions for the 20 somatic variants across all tools.

Tool performance showed overall high consistency between the germline and somatic subsets. VarChat ranked first across all five metrics in both contexts. Gpt-4o consistently placed second, while Perplexity and ScholarAI remained the lowest-performing tools. Readability & Usability was the most stable metric across subsets, with nearly identical scores for all tools. Notably, VarChat achieved a higher score for Literature Coverage & Relevance in the somatic setting (4.35 ± 0.76 vs 3.93 ± 1.40).

When examining Summarization Accuracy distributions at the variant level, we observed a wide range of tool behaviors (**Figure 4b**). The somatic variant p.E545K in PIK3CA received the highest average score across all tools and reviewers (mean 4.20), with relatively good agreement among reviewers (SD = 0.86). Conversely, c.11C>T in TP53 was the lowest-ranked variant (mean 3.20), also showing notable inter-reviewer variability (SD = 1.01). Variants such as p.R841W in BRCA1 and rs118203599 in TSC1 exhibited particularly high variability across tools and reviewers (SD > 1.1), suggesting divergent interpretations or tool-specific challenges in retrieving and summarizing the relevant literature. VarChat was ranked as the top-performing tool in 16 out of 20 somatic variants, further supporting its robustness in consistently delivering accurate summaries. Gpt-4o followed with five first-place rankings, while Perplexity and ScholarAI were each ranked first in only one case. These findings reinforce the consistency of the evaluation framework and highlight the variant-specific nuances in tool performance and reviewer agreement.

### Impact of literature availability on tool performance

To assess the performance under varying levels of literature support, we compared tool performance on variants stratified by literature abundance (Low: <10 references, Medium: 10–25, High: >25), as shown in **Figure 5a**. While most tools showed decreased scores across metrics under lower literature availability, the degree of change differed among them. For instance, VarChat average score in Summarization Accuracy decreased from 4.45 to 4.13 (Δ = –0.33, difference in means). Similar declines were observed across other metrics, including Hallucination Robustness (Δ = –0.23), Readability & Usability (Δ = –0.37), and Time Efficiency (Δ = –0.54). Accordingly, Literature Coverage & Relevance show a substantial drop from 4.56 to 3.46 (Δ = –1.10), highlighting the importance of literature availability for VarChat’s performance.

**Figure 5.**
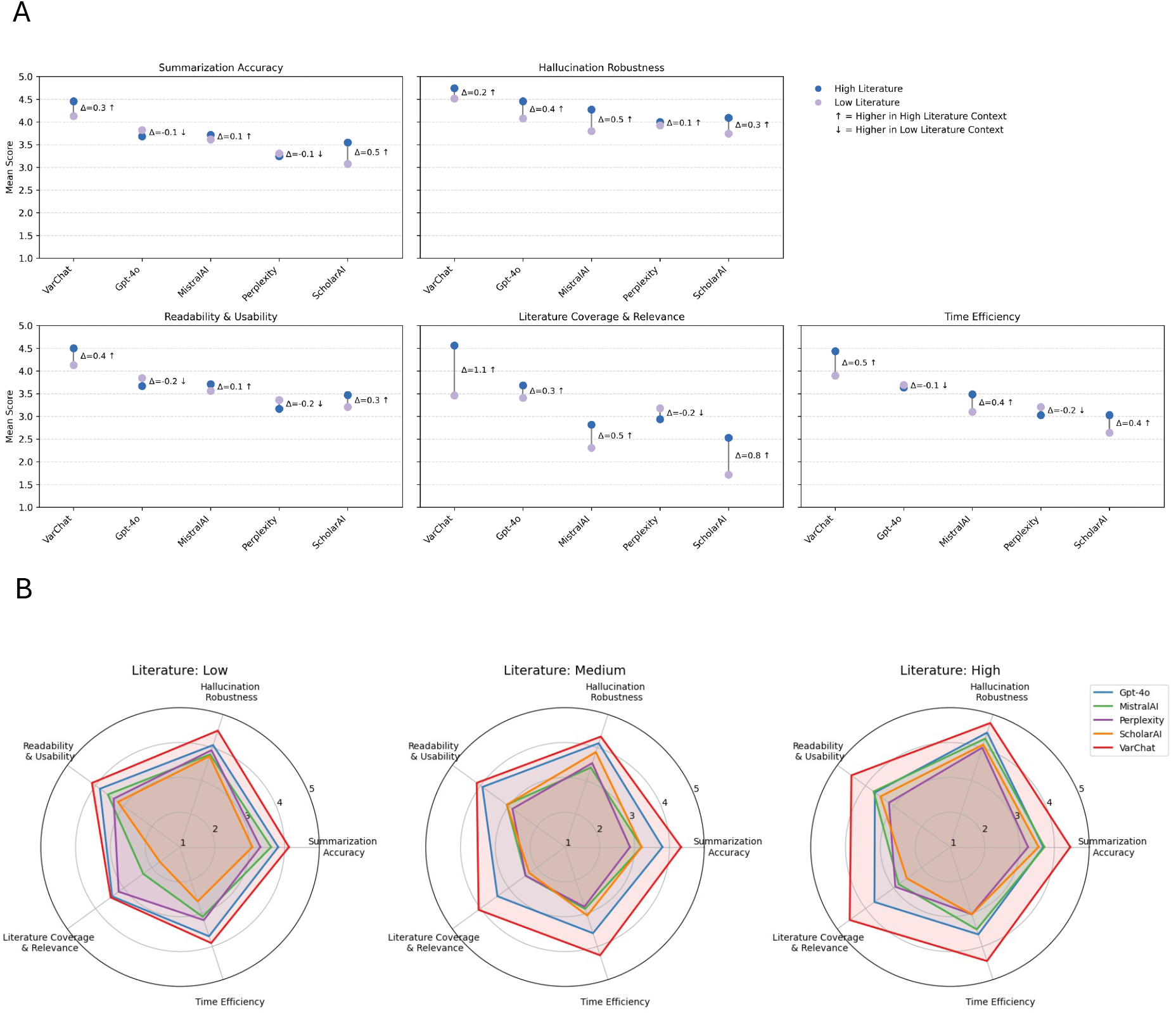
Tool performance stratified by literature support. **a**. Performance comparison between high- and low-literature variants across all tools (each dot-line represents a metric per tool). **b**. Radar plots of tool performance across five metrics for variants with low, medium, and high literature availability.

ScholarAI followed a similar trend but was even more affected overall. It showed the strongest drop in Literature Coverage & Relevance among all tools, falling from 2.53 to 1.72 (Δ = –0.81). Its Summarization Accuracy also declined markedly, from 3.55 to 3.08 (Δ = –0.47), alongside moderate losses in other metrics. MistralAI exhibited a smaller drop in Summarization Accuracy, decreasing only slightly from 3.71 to 3.62 (Δ = –0.10). While this suggests a relatively stable summarization capability, its overall trend remained consistent with the other tools, with lower scores observed under low-literature conditions. The Hallucination Robustness dropped from 4.27 to 3.79 (Δ = –0.48), indicating greater inconsistency or factual errors.

Time Efficiency also decreased (Δ = –0.38), along with a modest reduction in Readability & Usability (Δ = –0.15).

Gpt-4o and Perplexity displayed a distinct pattern compared to the literature-focused tools. Notably, both showed slightly higher scores in Summarization Accuracy under low-literature conditions—Gpt-4o increased from 3.68 to 3.82 (Δ = +0.14), and Perplexity from 3.24 to 3.31 (Δ = +0.07).

Despite this, both tools exhibited declines in Hallucination Robustness—Gpt-4o dropped from 4.45 to 4.08 (Δ = –0.38), and Perplexity from 4.00 to 3.92 (Δ = –0.08)—reflecting reduced factual consistency. This trend was observed across all tools, confirming the idea that less literature correlates with a greater risk of hallucinated or unsupported notions. In Perplexity, additional drops were noted in Time Efficiency (Δ = –0.17) and Readability & Usability (Δ = –0.19), suggesting a general weakening of performance in low-data settings despite the small gain in summarization clarity.

This pattern was further captured by radar plots of the five metrics under the three literature conditions (**Figure 5b**), where the polygon area summarizes each tool’s global performance. VarChat consistently showed the largest area across all tiers, ranging from 38.58 in low to 48.98 in high-literature settings. ScholarAI exhibits the smallest areas, with a minimal value of 19.82 under low-literature conditions. The sharpest area drop was observed for Perplexity (–7.48 units from high to medium), followed by MistralAI (–9.62 from high to medium), while Gpt-4o remained remarkably stable across conditions (range: 33.80–34.72). These values provide an aggregated view of the same trend: lower literature availability generally resulted in reduced global performance, though the magnitude of decline varied by tool.

## Discussion

The emergence of generative AI tools has opened new possibilities for academic literature analysis across scientific fields. In genomics, where variant assessment relies on a rapidly growing number of notions—often fragmented and spread across different peer-reviewed publications—these tools can provide efficient access to relevant and customized information, in a very short time, far less than a manual review. Yet, despite the increasing use of LLMs in biomedical research and the growing interest in their application to clinical and basic research, few studies have rigorously evaluated their performance on specific tasks, such as genomic variant annotation.

To our knowledge, this work represents the first blinded, expert-reviewed benchmark on real clinical variants across five generative platforms. Under controlled prompt conditions, the results provide a snapshot of current tool performance in summarizing biological and clinical knowledge on genomic variants. In detail, we also assessed the ability of models to accurately retrieve and use information from peer-reviewed articles to verify that they provide referenced, relevant, and as up-to-date data as possible. The study reflects the state of these tools as of March 2025, yet the methodology is designed to be re-run periodically, allowing longitudinal tracking of tool evolution and performance over time. To our knowledge, this is one of the first efforts in the field to systematically assess LLM-generated content with expert scoring and defined metrics, using real variants of clinical and biological interest.

Across all metrics and variant types, VarChat consistently ranked as the top-performing tool, likely due to its real-time access to the literature and to a robust prioritization method to identify up to 15 relevant papers. It showed high summarization accuracy, high hallucination robustness, and high usability, both in germline and somatic settings. Gpt-4o was, on average, the second-ranked tool, with robust and stable performance that was less affected by literature-related performance drops—especially considering its general-purpose nature. In clinical genomics, where rare variants may have few publications, a general-purpose model may sometimes yield more usable starting points than a literature-specialized tool. MistralAI showed moderate results, while Perplexity and ScholarAI— despite being positioned as literature-focused tools—ranked lower across nearly all metrics. The time efficiency metric is also consistent with the other metrics, ranking the tools with the same positions. These trends remained consistent across variant types and reviewer subsets, highlighting consistent differences in tool behavior and reliability.

Expectedly, literature availability emerged as a key factor influencing tool performance. Most platforms showed reduced accuracy, consistency, and citation quality when variant-related publications were scarce. In most cases, AI tools operate on open literature. Non-open literature could make a significant contribution to increasing the availability of information to be retrieved. However, LLMs cannot access closed literature unless they operate under special licenses. VarChat maintained the highest scores across all performance metrics even in low-reference settings, although its performance still declined when the literature was limited. ScholarAI proved particularly sensitive to scarcity, with a marked drop in both relevance and summarization accuracy. Interestingly, general-purpose models such as Gpt-4o performed more robustly under these conditions, likely due to broader training data rather than strict reliance on peer-reviewed articles. For example, in the absence of dedicated publications, Gpt-4o retrieves information from databases such as GeneCards, OMIM, or NCBI, but also from poster abstracts hosted on genomic conference websites. These sources, while not peer-reviewed in the strict sense, often provide valid and practically useful information, especially for variants otherwise absent from the literature. Overall, the availability of a reasonable body of peer-reviewed publications remains critical to ensure reliable outputs.

The evaluation design used in this study aimed to reflect real-world use cases, where users input a heterogeneous set of genomic variants, including mutations related to both somatic and germline contexts, different levels of literature support, and a subset of neutral variants used as internal controls, all queried through a fixed prompt specifically optimized for this type of task. This allowed us to test not only how tools perform on well-characterized pathogenic variants, but also how they handle scarcity. All summaries were generated using stable and freely accessible tool versions within a fixed time window, with fresh sessions and controlled prompting to prevent context bias and ensure a fair comparison. Expert reviewers received outputs with no visual formatting and without the tool name they were evaluating, ensuring that scoring was based solely on content quality.

This study presents three main limitations. First, the evaluation is strongly dependent on a specific timepoint—March 2025—using the tool versions available during that period. As both scientific literature and, especially, generative models evolve rapidly, performance assessments may shift quickly—sometimes in months or even weeks. This temporal volatility limits the reproducibility of this study. At the same time, the need to provide a clear assessment of current capabilities remains a crucial step to understand and guide the evolution of this field. Second, an inherent limitation of generative AI is that the reproducibility of results is not guaranteed, and therefore evaluation must also consider the specific values obtained at the time the summaries were generated. For example, a hallucination produced during the execution of a summary may not be repeated or may be repeated in a different form if the request is made in another attempt. Finally, the evaluation was based on 40 variants and six reviewers. This approach cannot be considered a large-scale assessment, but rather a highly curated benchmark: it represents a substantial manual effort, especially given the complexity of the outputs and the need for expert-level evaluation. Unlike purely automated evaluations, this approach required careful human curation—an element still rarely implemented in LLM assessment.

The findings presented here highlight both the potential and current limitations of state-of-the-art open access tools based on generative AI in supporting variant interpretation.

This suggests that LLMs integration into genomic research should be approached with caution, and with a clear understanding of the conditions under which performance is reliable. Future developments in model transparency, citation handling, and evidence grounding will be essential to increase their usability. Moreover, manual expert evaluation remains a critical component of any tool aiming to support research—and even more importantly, clinical and translational decisions.

## Conclusion

Within the rapidly evolving landscape of generative AI, this study offers a structured, expert-reviewed benchmark for evaluating current tools in the task of variant summarization. The summarization task represents just one of many steps contributing to the variant interpretation process. In this context, where timely and accurate information is critical, AI tools have the potential to reduce the manual curation required by clinicians, provided there is a careful understanding of when and how performance remains robust and trustable. Thanks to further improvements, these tools could play a valuable role alongside experts in both genomic research and clinical settings.

## Methods Study Design

The analysis presented in this manuscript aims to determine whether generative AI tools can be effectively used to retrieve relevant biological and clinical information on genetic variants from peer-reviewed scientific literature. The evaluation employed a curated benchmarking dataset consisting of forty selected human genomic variants (i.e. SNPs and small indels): 20 germline variants, mostly pathogenic and associated with inherited disorders (see Methods), and 20 cancer-related variants, primarily somatic and pathogenic. We evaluated five tools, two general-purpose LLMs and three tools specialized in extracting content from peer-reviewed literature.

After implementing fine-tuned prompting methods to extract summaries, a panel of six experts assessed the quality of the generated outputs using five predefined evaluation metrics. Three reviewers evaluated the germline portion of the benchmark, while the other three reviewed the somatic portion. Consequently, each of the 40 variants underwent independent evaluation by three reviewers. In this context, the term ‘reviewers’ refers to the clinicians and researchers who evaluated the tool outputs. They were experienced professionals with diverse backgrounds in oncology, immunology, hematology, and molecular medicine, routinely engaged in genomic data interpretation across research and clinical settings. Their expertise ensured that the evaluation reflected informed domain judgment in variant-related tasks.

The reviewers’ evaluation was conducted in a single-blind fashion. The reviewers assigned scores to each summary, without knowing which tool it was created with and without knowing which ones were included in the benchmark. The sorting within the files given to the reviewers was random, without specific patterns that could infer the association between specific tool and output. The goal was to minimize bias and allow fair evaluation of the tools.

After approximately four weeks, quantitative results and valuable reviewer comments were collected. An analysis of the assigned scores was conducted, and the outcomes are presented through plots in the Results section.

**Figure 1** presents an overview of the study workflow and the evaluation design.

## Variant selection

To assess the reliability of the AI tools in generating accurate and complete variant summaries, we created a curated benchmarking dataset comprising 40 human genomic variants (reported in the Supplementary Materials). The dataset includes 20 germline variants, predominantly pathogenic and associated with inherited disorders, and 20 cancer-related variants, primarily somatic and pathogenic.

Between germline variants, we included both SNPs and small indels, with the aim of covering different types of diseases (including ataxia telangiectasia, Aicardi-Goutières syndrome, cystic fibrosis, severe combined immunodeficiency, phenylketonuria, sickle cell anemia, and many others). We also included a few variants involved in digenic inheritance mechanisms, taken individually, linked respectively to Bardet-Biedl syndrome and Long QT syndrome. We included both pathogenic and benign variants, relying on the information reported in ClinVar.

For somatic variants, we spanned several types of cancer, including breast cancer, gliomas, bladder and endometrial cancer. Across the 20 somatic variants, 16/20 variants are reported as oncogenic, and 4/20 as neutral.

To construct the prompt, shown later in this Methods section, we chose to use the standard nomenclature by which the variant is best known. Specifically, for germline variants we used the coding sequence (standard HGVS coding) for 11/20 variants, the protein sequence (standard HGVS protein) for 8/20 and the dbSNP ID for 1/20 variants. For somatic variants, we used the coding sequence (standard HGVS coding) for 1/20 variants, the protein sequence (standard HGVS protein) for 17/20 and the dbSNP ID for 2/20 variants.

We strategically selected variants to represent varying levels of literature evidence, from extensively studied variants with abundant information available to those minimally documented. We used for literature abundance a qualitative metric (high, medium, low) derived from the quantitative discretization of available peer-reviewed articles: high (≥25 articles), medium (10–24 articles), and low (<10 articles). This metric includes articles identified through searches in PubMed, PMC and Google Scholar, performed for each variant with an appropriate query including alternative nomenclature, to maximize the literature retrieval yield. The benchmark included 13 low, 5 medium, and 22 high-literature variants. The full benchmark dataset used is given in the Supplementary Materials.

## Model selection

We used the described dataset of 40 variants to benchmark a pool of five AI tools in generating concise summaries of genetic variants based on the scientific literature. Since the aim was to mimic the practical need and behavior of a clinician, tools selection was guided by several practical considerations. The tools had to offer freely accessible options at no additional cost; furthermore, it was essential that they feature user-friendly web applications or graphical interfaces to avoid the need for APIs or pre-processing or local deployment steps that could complicate their use. The tools were then selected to represent both generic functionality and access to the specialized scientific literature.

At first, we compared 8 generative AI tools (ChatGPT, MistralAI, DeepSeek, Claude, VarChat, Perplexity, ScholarAI, FAVOR-GPT), evaluating each one based on response time and high-level usability. DeepSeek [32] and Claude [33] were excluded from further analysis: during the initial evaluation, in March 2025, DeepSeek experienced consistent server availability issues while testing it, limiting reliable performance assessment, while Claude lacked the web search capability. The integration of web search capabilities is essential for retrieving up-to-date scientific literature, clinical guidelines, and variant-specific databases, thereby significantly enhancing the clinical relevance of the generated summaries. FAVOR-GPT [34] uses generative AI to synthesize genomic variants, but relies on a retrieval strategy centered on annotations from functional databases, rather than incorporating information extracted directly from the scientific literature. Because of this difference in rationale, we excluded it from the final benchmark.

From this preliminary analysis, five tools were selected for the benchmark evaluation: ChatGPT [35], MistralAI [36], VarChat [37] [38], Perplexity [39], and ScholarAI [40].

Among the tools, ChatGPT, MistralAI and Perplexity are general-purpose AI platforms, with built-in functionalities that allow them to access both external websites and biobank databases. In contrast, VarChat, and ScholarAI are specialized platforms explicitly designed to query and synthesize information exclusively from scientific literature.

However, Perplexity was used with the customized web search based only on scientific papers, with the specific setting “Academic Search”, which allows to consider the tool like VarChat and ScholarAI. For this reason, we grouped together VarChat, ScholarAI and Perplexity, labelling them as “literature based” tools. Detailed specifications, including version information of the tools, are provided in Table 1.

**Table 1.**
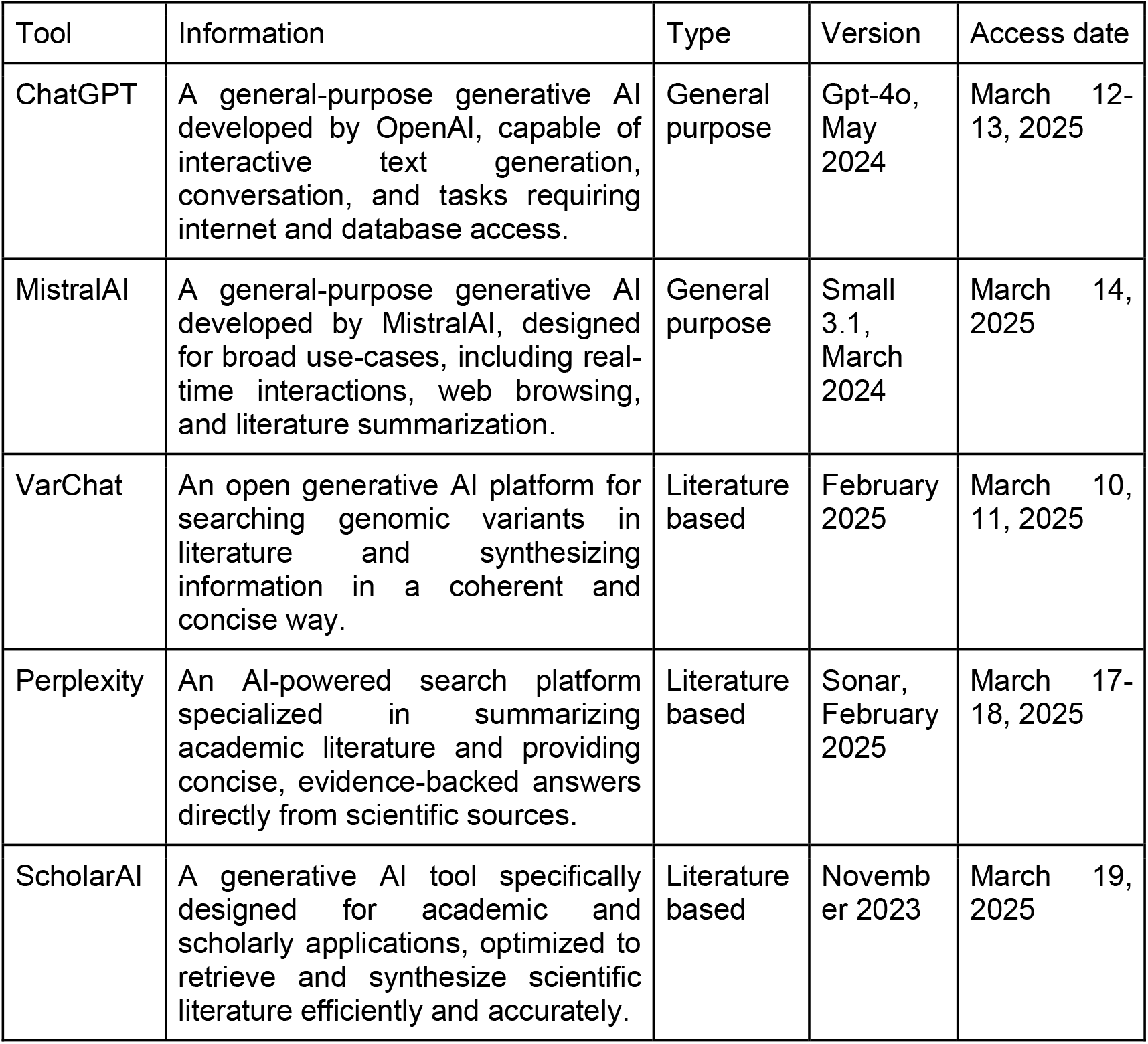
Five AI tools in comparison.

| Tool | Information | Type | Version | Access date |
| --- | --- | --- | --- | --- |
| ChatGPT | A general-purpose generative AI developed by OpenAI, capable of interactive text generation, conversation, and tasks requiring internet and database access. | General purpose | Gpt-4o, May 2024 | March 12-13, 2025 |
| MistralAI | A general-purpose generative AI developed by MistralAI, designed for broad use-cases, including real-time interactions, web browsing, and literature summarization. | General purpose | Small 3.1, March 2024 | March 14, 2025 |
| VarChat | An open generative AI platform for searching genomic variants in literature and synthesizing information in a coherent and concise way. | Literature based | February 2025 | March 10, 11, 2025 |
| Perplexity | An AI-powered search platform specialized in summarizing academic literature and providing concise, evidence-backed answers directly from scientific sources. | Literature based | Sonar, February 2025 | March 17-18, 2025 |
| ScholarAI | A generative AI tool specifically designed for academic and scholarly applications, optimized to retrieve and synthesize scientific literature efficiently and accurately. | Literature based | November 2023 | March 19, 2025 |

## Prompt curation

To ensure that the generated summaries met the practical needs of clinical geneticists and clinicians, we carefully curated a prompt that aims to reflect the clinician perspective and literature-based information requirements, where supported by the respective tool. This prompt specifically instructed the AI platforms to provide a clinically relevant and scientifically rigorous overview of each genetic variant under consideration, thus ensuring both the relevance and practical utility of the generated content.

We refined the prompt with sequential improvements, and the final version includes the following components: (1) explicit task instructions and stylistic guidance, defining the objective of generating a concise summary with a maximum length of 500 words, in plain text format without bullet points or subtitles; (2) nomenclature of the genetic variant and associated gene; (3) scientific details required in the summary, such as the clinical classification of the variant and the biological function of the related gene; (4) preferences for sourcing information, prioritizing peer-reviewed publications when available, along with recommended citation practices; and (5) safety measures aimed at minimizing inaccuracies and hallucinations, maintaining an appropriate professional tone. The complete text of the prompt is reported in Table 2.

**Table 2.**
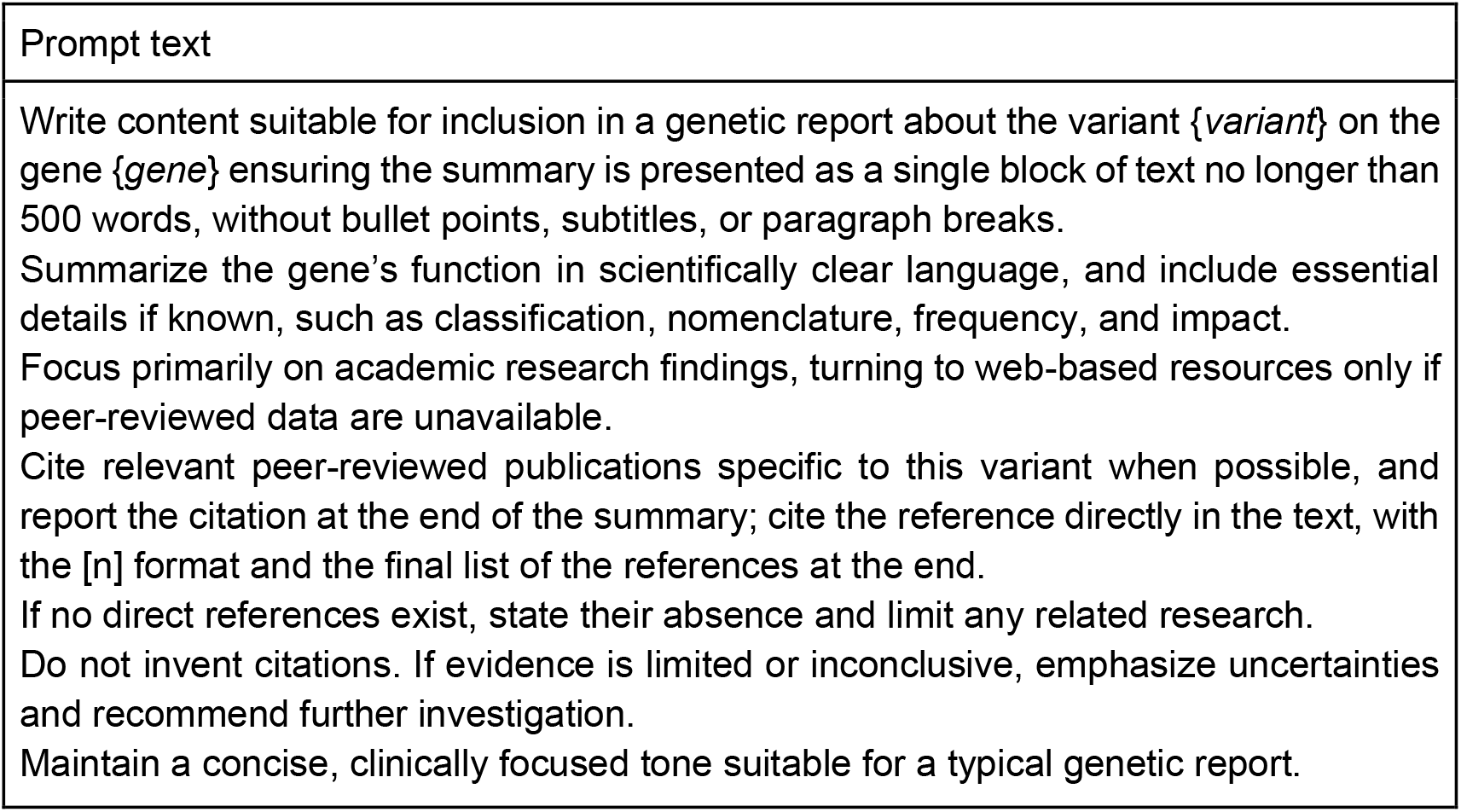
Curated prompt provided to each tool, customizable with information on the specific variant.

Prompt text Write content suitable for inclusion in a genetic report about the variant {*variant*} on the gene {*gene*} ensuring the summary is presented as a single block of text no longer than 500 words, without bullet points, subtitles, or paragraph breaks.

Summarize the gene’s function in scientifically clear language, and include essential details if known, such as classification, nomenclature, frequency, and impact.

Focus primarily on academic research findings, turning to web-based resources only if peer-reviewed data are unavailable.

Cite relevant peer-reviewed publications specific to this variant when possible, and report the citation at the end of the summary; cite the reference directly in the text, with the [n] format and the final list of the references at the end.

If no direct references exist, state their absence and limit any related research.

Do not invent citations. If evidence is limited or inconclusive, emphasize uncertainties and recommend further investigation.

Maintain a concise, clinically focused tone suitable for a typical genetic report.

For each of the 40 variants included in the study, we generated five independent summaries - one with every available tool - using the described prompt wherever the interface allowed. Every tool allows the customized prompt, except for VarChat: it does not permit a fully customized prompt but only allows to query the variant nomenclature. Therefore, we used HGVS notation together with the gene symbol or the dbSNP ID. The full data was generated in two weeks, March 10 to 21, 2025. To prevent conversations from affecting subsequent answers, we submitted every prompt in a fresh chat session, thus guaranteeing complete isolation and removing any memory effects that might influence the LLMs performance. All runs for the same variant were executed with the same, stable release of each tool, ensuring consistency across the full experimental set.

We presented the output text of the models to reviewers. We maintain the references generated by the tools exactly as produced, keeping the original citation format to facilitate manual revision. Only visual platform styling was omitted: the reviewers received the raw text without formatting or graphical embellishments, so that all comparisons would rest solely on the substantive content of the summaries.

A few examples of output summaries are reported in Supplementary Materials.

The choice of reviewers was a key step in the presented method. As mentioned above, manual review by experts lends quality to the evaluation, whereas review by independent AI tools can lead to inaccuracies and hallucinations. To find a compromise between the number of reviewers and the time for assigning the work, evaluating and processing the results, we chose to involve six expert reviewers. Reviewers occupy a variety of roles, including experts in oncology, gynecology, molecular medicine, immunology, and hematology. Three experts evaluated summaries related to germline variants, while the other three focused on somatic variants. As a result, each of the 40 variants was independently evaluated by three reviewers.

## Review metrics

To enable objective and quantitative evaluation, we defined and curated five evaluation metrics: Summarization Accuracy, Hallucination Robustness, Readability & Usability, Literature Coverage & Relevance, and Time Efficiency. Each metric aims to answer qualitative questions with respect to the summary, and the rating includes values from 1 to 5, customized with respect to the metric. The full explanation of each metric is reported in Table 3.

**Table 3.**
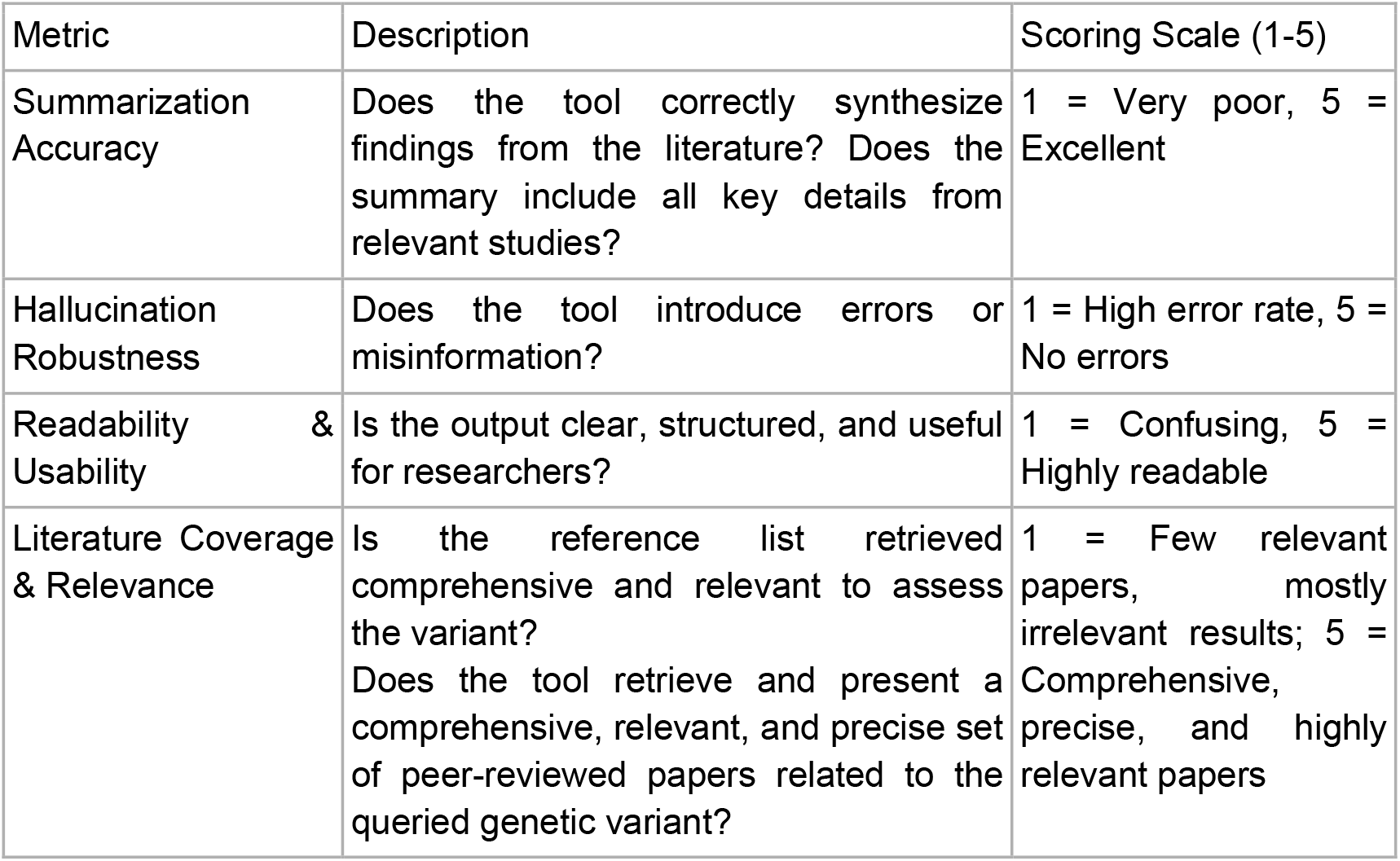
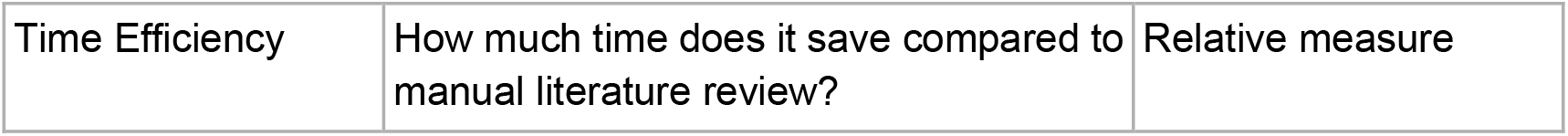
Evaluation metrics.

| Metric | Description | Scoring Scale (1-5) |
| --- | --- | --- |
| Summarization Accuracy | Does the tool correctly synthesize findings from the literature? Does the summary include all key details from relevant studies? | 1 = Very poor, 5 = Excellent |
| Hallucination Robustness | Does the tool introduce errors or misinformation? | 1 = High error rate, 5 = No errors |
| Readability & Usability | Is the output clear, structured, and useful for researchers? | 1 = Confusing, 5 = Highly readable |
| Literature Coverage & Relevance | Is the reference list retrieved comprehensive and relevant to assess the variant?<br>Does the tool retrieve and present a comprehensive, relevant, and precise set of peer-reviewed papers related to the queried genetic variant? | 1 = Few relevant papers, mostly irrelevant results; 5 = Comprehensive, precise, and highly relevant papers |
| Time Efficiency | How much time does it save compared to manual literature review? | Relative measure |

## Abbreviations

AI: Artificial Intelligence
NLP: Natural Language Processing
LLMs: Large Language Models

## Declarations

Andrea Gazzo reports a consultancy relationship with enGenome. Silvia Berardelli was involved in the development of the VarChat tool.

## References

1. Lee, K., Wei, C.-H., Lu, Z.: Recent advances of automated methods for searching and extracting genomic variant information from biomedical literature. Brief. Bioinform. 22, bbaa142 (2021). 10.1093/bib/bbaa142.

2. Zhao, S., Su, C., Lu, Z., Wang, F.: Recent advances in biomedical literature mining. Brief. Bioinform. 22, (2021). 10.1093/bib/bbaa057.

3. Lin, Y.-H., Lu, Y.-C., Chen, T.-F., Hsu, J.S., Lee, K.-H., Cheng, Y.-W., et al.: variant2literature: full text literature search for genetic variants, http://biorxiv.org/lookup/doi/10.1101/583450, (2019). 10.1101/583450.

4. Cejuela, J.M., Bojchevski, A., Uhlig, C., Bekmukhametov, R., Kumar Karn, S., Mahmuti, S., et al.: nala : text mining natural language mutation mentions. Bioinformatics. 33, 1852–1858 (2017). 10.1093/bioinformatics/btx083.

5. Allot, A., Wei, C.-H., Phan, L., Hefferon, T., Landrum, M., Rehm, H.L., et al.: Tracking genetic variants in the biomedical literature using LitVar 2.0. Nat. Genet. 55, 901–903 (2023). 10.1038/s41588-023-01414-x.

6. Wei, C.-H., Allot, A., Lai, P.-T., Leaman, R., Tian, S., Luo, L., et al.: PubTator 3.0: an AI-powered literature resource for unlocking biomedical knowledge. Nucleic Acids Res. 52, W540–W546 (2024). 10.1093/nar/gkae235.

7. Wei, C.-H., Allot, A., Riehle, K., Milosavljevic, A., Lu, Z.: tmVar 3.0: an improved variant concept recognition and normalization tool. Bioinformatics. 38, 4449–4451 (2022). 10.1093/bioinformatics/btac537.

8. Pasche, E., Mottaz, A., Caucheteur, D., Gobeill, J., Michel, P.-A., Ruch, P.: Variomes: a high recall search engine to support the curation of genomic variants. Bioinformatics. 38, 2595–2601 (2022). 10.1093/bioinformatics/btac146.

9. Cheng, C., Tan, F., Wei, Z.: DeepVar: An End-to-End Deep Learning Approach for Genomic Variant Recognition in Biomedical Literature. Proc. AAAI Conf. Artif. Intell. 34, 598–605 (2020). 10.1609/aaai.v34i01.5399.

10. Birgmeier, J., Deisseroth, C.A., Hayward, L.E., Galhardo, L.M.T., Tierno, A.P., Jagadeesh, K.A., et al.: AVADA: toward automated pathogenic variant evidence retrieval directly from the full-text literature. Genet. Med. 22, 362–370 (2020). 10.1038/s41436-019-0643-6.

11. Chunn, L.M., Nefcy, D.C., Scouten, R.W., Tarpey, R.P., Chauhan, G., Lim, M.S., et al.: Mastermind: A Comprehensive Genomic Association Search Engine for Empirical Evidence Curation and Genetic Variant Interpretation. Front. Genet. 11, 577152 (2020). 10.3389/fgene.2020.577152.

12. Chowdhary, K.R.: Natural Language Processing. In: Fundamentals of Artificial Intelligence. pp. 603–649. Springer India, New Delhi (2020). 10.1007/978-81-322-3972-7_19.

13. Tang, L., Sun, Z., Idnay, B., Nestor, J.G., Soroush, A., Elias, P.A., et al.: Evaluating large language models on medical evidence summarization. Npj Digit. Med. 6, (2023). 10.1038/s41746-023-00896-7.

14. Vaswani, A., Shazeer, N., Parmar, N., Uszkoreit, J., Jones, L., Gomez, A.N., et al.: Attention is All you Need. In: Guyon, I., Luxburg, U.V., Bengio, S., Wallach, H., Fergus, R., Vishwanathan, S., et al. (eds.) Advances in Neural Information Processing Systems. Curran Associates, Inc. (2017).

15. Ghali, M.-K., Farrag, A., Won, D., Jin, Y.: Enhancing knowledge retrieval with in-context learning and semantic search through generative AI. Knowl.-Based Syst. 311, 113047 (2025). 10.1016/j.knosys.2025.113047.

16. Zhang, Z., Wang, C., Wang, Y., Shi, E., Ma, Y., Zhong, W., et al.: LLM Hallucinations in Practical Code Generation: Phenomena, Mechanism, and Mitigation. Proc. ACM Softw. Eng. 2, 481–503 (2025). 10.1145/3728894.

17. Du, X., Nagy, A., Oates, M.F., Wang, Y., Wang, X., Plasek, J.M., et al.: Precision Grounding: Augmenting Large Language Models with Evidence-Based Databases for Trustworthy Genetic Variant Summarization, http://medrxiv.org/lookup/doi/10.1101/2025.06.09.25329279, (2025). 10.1101/2025.06.09.25329279.

18. Kumar, C.V., Urlana, A., Kanumolu, G., Garlapati, B.M., Mishra, P.: No LLM is Free From Bias: A Comprehensive Study of Bias Evaluation in Large Language Models, https://arxiv.org/abs/2503.11985, (2025). 10.48550/ARXIV.2503.11985.

19. Walters, W.H., Wilder, E.I.: Fabrication and errors in the bibliographic citations generated by ChatGPT. Sci. Rep. 13, 14045 (2023). 10.1038/s41598-023-41032-5.

20. Tam, T.Y.C., Sivarajkumar, S., Kapoor, S., Stolyar, A.V., Polanska, K., McCarthy, K.R., et al.: A framework for human evaluation of large language models in healthcare derived from literature review. Npj Digit. Med. 7, 258 (2024). 10.1038/s41746-024-01258-7.

21. Croxford, E., Gao, Y., Pellegrino, N., Wong, K., Wills, G., First, E., et al.: Current and future state of evaluation of large language models for medical summarization tasks. Npj Health Syst. 2, 6 (2025). 10.1038/s44401-024-00011-2.

22. Croxford, E., Gao, Y., First, E., Pellegrino, N., Schnier, M., Caskey, J., et al.: Automating Evaluation of AI Text Generation in Healthcare with a Large Language Model (LLM)-as-a-Judge, http://medrxiv.org/lookup/doi/10.1101/2025.04.22.25326219, (2025). 10.1101/2025.04.22.25326219.

23. Shankar, S., Zamfirescu-Pereira, J.D., Hartmann, B., Parameswaran, A., Arawjo, I.: Who Validates the Validators? Aligning LLM-Assisted Evaluation of LLM Outputs with Human Preferences. In: Proceedings of the 37th Annual ACM Symposium on User Interface Software and Technology. pp. 1–14. ACM, Pittsburgh PA USA (2024). 10.1145/3654777.3676450.

24. Chiang, C.-H., Lee, H.: Can Large Language Models Be an Alternative to Human Evaluations? In: Proceedings of the 61st Annual Meeting of the Association for Computational Linguistics (Volume 1: Long Papers). pp. 15607–15631. Association for Computational Linguistics, Toronto, Canada (2023). 10.18653/v1/2023.acl-long.870.

25. Masanneck, L., Meuth, S.G., Pawlitzki, M.: Evaluating base and retrieval augmented LLMs with document or online support for evidence based neurology. Npj Digit. Med. 8, 137 (2025). 10.1038/s41746-025-01536-y.

26. Van Veen, D., Van Uden, C., Blankemeier, L., Delbrouck, J.-B., Aali, A., Bluethgen, C., et al.: Adapted large language models can outperform medical experts in clinical text summarization. Nat. Med. 30, 1134–1142 (2024). 10.1038/s41591-024-02855-5.

27. Salvi, R.C., Panigrahi, S., Jain, D., Yadav, S., Akhtar, Md.S.: Towards Understanding LLM-Generated Biomedical Lay Summaries. In: Proceedings of the Second Workshop on Patient-Oriented Language Processing (CL4Health). pp. 260–268. Association for Computational Linguistics, Albuquerque, New Mexico (2025). 10.18653/v1/2025.cl4health-1.22.

28. Chen, Q., Sun, H., Liu, H., Jiang, Y., Ran, T., Jin, X., et al.: An extensive benchmark study on biomedical text generation and mining with ChatGPT. Bioinformatics. 39, btad557 (2023). 10.1093/bioinformatics/btad557.

29. Toprak, A.G., Onan, A.: Benchmarking Intelligent Large Language Models for Biomedical Text Summarization: A Performance Evaluation. In: Kahraman, C., Cebi, S., Oztaysi, B., Cevik Onar, S., Tolga, C., Ucal Sari, I., et al. (eds.) Intelligent and Fuzzy Systems. pp. 492–500. Springer Nature Switzerland, Cham (2025). 10.1007/978-3-031-97992-7_55.

30. Green, A., Ribas, C.E., Ontiveros-Palacios, N., Griffiths-Jones, S., Petrov, A.I., Bateman, A., et al.: LitSumm: large language models for literature summarization of noncoding RNAs. Database. 2025, baaf006 (2025). 10.1093/database/baaf006.

31. Luo, Z., Qiao, Y., Xu, X., Li, X., Xiao, M., Kang, A., et al.: Cross sectional pilot study on clinical review generation using large language models. Npj Digit. Med. 8, 170 (2025). 10.1038/s41746-025-01535-z.

32. DeepSeek-AI, Liu, A., Feng, B., Xue, B., Wang, B., Wu, B., et al.: DeepSeek-V3 Technical Report, https://arxiv.org/abs/2412.19437, (2024). 10.48550/ARXIV.2412.19437.

33. Claude Opus 4, https://www.anthropic.com/claude/opus, last accessed 2025/07/25.

34. Li, T.C., Zhou, H., Verma, V., Tang, X., Shao, Y., Van Buren, E., et al.: FAVOR-GPT: a generative natural language interface to whole genome variant functional annotations. Bioinforma. Adv. 4, vbae143 (2024). 10.1093/bioadv/vbae143.

35. OpenAI, Hurst, A., Lerer, A., Goucher, A.P., Perelman, A., Ramesh, A., et al.: PT-4o System Card, https://arxiv.org/abs/2410.21276, (2024). 10.48550/ARXIV.2410.21276.

36. Mistral Small 3.1 | MistralAI, https://mistral.ai/news/mistral-small-3-1, last accessed 2025/07/25.

37. De Paoli, F., Berardelli, S., Limongelli, I., Rizzo, E., Zucca, S.: VarChat: the generative AI assistant for the interpretation of human genomic variations. Bioinformatics. 40, btae183 (2024). 10.1093/bioinformatics/btae183.

38. De Paoli, F., Berardelli, S., Tudisco, A., Blindu, A., Parimbelli, E., Zucca, S.: Generative AI Meets Genomics: VarChat, a RAG-Based Approach for Literature-Driven Variant Summarization. In: Bellazzi, R., Juarez Herrero, J.M., Sacchi, L., and Zupan, B. (eds.) Artificial Intelligence in Medicine. pp. 127–131. Springer Nature Switzerland, Cham (2025). 10.1007/978-3-031-95841-0_24.

39. Sonar by Perplexity, https://sonar.perplexity.ai/, last accessed 2025/07/29.

40. Scholar AI | AI Research Assistant for Students, Developers & Clinical Teams, https://scholarai.io, last accessed 2025/08/04.

