## Supplementary material for "Benchmarking generative AI tools for literature retrieval and summarization in genomic variant interpretation": ATM c.496G_C.docx

| **Tool** | **Summary** | **References** |
| --- | --- | --- |
| **1** | The genomic variant c.496G>C p.Glu166Gln rs1565357473 is located in the ATM gene, which encodes the ataxia-telangiectasia mutated protein, a key player in the cellular response to DNA damage. The ATM protein is  involved in the activation of cell cycle checkpoints, DNA repair, and apoptosis, thereby maintaining genomic stability.  The c.496G>C variant results in the substitution of glutamic acid with glutamine at position 166 of the ATM protein (p.Glu166Gln). This missense mutation has been identified in a consanguineous family with two cases of late-onset cerebellar ataxia with neuropathy [1]. The affected individuals presented with cerebellar ataxia, hand tremor, neck dystonia, distal muscle wasting, and diminished tendon reflexes, but without the classic ataxia-telangiectasia symptoms of conjunctival telangiectasia or immunodeficiency. Additionally, slightly elevated alpha-fetoprotein levels and marked cerebellar atrophy with mild brainstem atrophy were observed on brain MRI [1].  Functional analysis of this variant revealed that it leads to aberrant splicing, specifically the skipping of an exon, as demonstrated by RT-PCR analysis [1]. This aberrant splicing event is likely to disrupt the normal function of the ATM protein, contributing to the observed clinical phenotype.  In the context of variant interpretation, rs1565357473 has been submitted to ClinVar with a total of 3 pathogenic submissions and 2 submissions with uncertain significance. The variant is classified as both a missense and splice region variant, which underscores its potential dual impact on protein structure and splicing. The pathogenic submissions in ClinVar support the findings in the family described, suggesting a deleterious effect of the c.496G>C p.Glu166Gln variant on protein function and its association with the disease phenotype [1]. | [[1]](https://pubmed.ncbi.nlm.nih.gov/32259893/) Shimazaki H et al. (2020). "Late-onset autosomal recessive cerebellar ataxia and neuropathy with a novel splicing mutation in the ATM gene." Journal of integrative neuroscience, 19(1) |
|  | Evaluation  **Summarization Accuracy & Completeness:**  **Hallucination Rate:**  **Readability & Usability:**  **Literature Coverage & Relevance:**  **Time Efficiency:** | |
| **2** | The ATM gene encodes a serine/threonine protein kinase crucial for maintaining genomic stability. It orchestrates cellular responses to DNA double-strand breaks by phosphorylating key proteins involved in DNA repair, cell cycle arrest, or apoptosis, including p53, CHK2, BRCA1, and H2AX. Mutations in ATM are linked to ataxia-telangiectasia, a disorder characterized by cerebellar degeneration, immunodeficiency, radiosensitivity, and increased cancer risk. The specific variant ATM c.496G>C results in a glycine-to-alanine substitution at codon 166 (p.Gly166Ala). This variant is not widely reported in the literature, and its clinical significance remains uncertain. The absence of peer-reviewed studies directly addressing this variant necessitates cautious interpretation. Functional assays and comprehensive clinical correlation are recommended to elucidate its potential pathogenicity. Given the limited data, further research is essential to determine the impact of the ATM c.496G>C variant on protein function and its association with disease. | [1] [researchgate.net (PDF) Specifications of the ACMG/AMP variant curation guidelines ...](https://www.researchgate.net/publication/380996014_Specifications_of_the_ACMGAMP_variant_curation_guidelines_for_the_analysis_of_germline_ATM_sequence_variants)  [16 dicembre 2024 — This VCEP made specifications for ACMG/AMP guidelines for the ataxia telangiectasia mutated (ATM) gene according to the Food and Drug Administration (FDA)- ...](https://www.researchgate.net/publication/380996014_Specifications_of_the_ACMGAMP_variant_curation_guidelines_for_the_analysis_of_germline_ATM_sequence_variants)  [2] [medlineplus.gov ATM gene: MedlinePlus Genetics 19 settembre 2022 — The ATM gene provides instructions for making a protein that helps control the rate at which cells grow and divide.](https://medlineplus.gov/genetics/gene/atm/)  [3] [pathsocjournals.onlinelibrary.wiley.com Minigene‐based splicing analysis and ACMG/AMP‐based tentative … 18 giugno 2022 — We decided to classify all 56 ATM variants according to ACMG/AMP variant classification guidelines](https://pathsocjournals.onlinelibrary.wiley.com/doi/full/10.1002/path.5979)  [4] [ncbi.nlm.nih.gov c.496+2T>G AND Ataxia-telangiectasia syndrome - ClinVar - NCBI](https://www.ncbi.nlm.nih.gov/clinvar/RCV001223507.2/)  [23 aprile 2022 — ClinVar archives and aggregates information about relationships](https://www.ncbi.nlm.nih.gov/clinvar/RCV001223507.2/)  [5] [pmc.ncbi.nlm.nih.gov ATM: Functions of ATM Kinase and Its Relevance to Hereditary … 4 gennaio 2022 — Ataxia–telangiectasia mutated (ATM) functions as a key initiator](https://pmc.ncbi.nlm.nih.gov/articles/PMC8745051/)  [6] [Nature Cellular functions of the protein kinase ATM and their relevance to … 24 agosto 2021 — The protein kinase ataxia telangiectasia mutated (ATM) is a ...](https://www.nature.com/articles/s41580-021-00394-2)  [7] [pmc.ncbi.nlm.nih.gov ATM: Main Features, Signaling Pathways, and Its Diverse Roles in … 30 maggio 2021 — Thus, thanks to its central role in coordinating DNA damage repair, ...](https://pmc.ncbi.nlm.nih.gov/articles/PMC8228802/)  [8] [Nature The ATM protein kinase: regulating the cellular response to ... - Nature](https://www.nature.com/articles/nrm3546)  [13 marzo 2013 — The most documented function of ATM is mobilizing and...](https://www.nature.com/articles/nrm3546)  [9] [ncbi.nlm.nih.gov ClinVar Genomic variation as it relates to human health - NCBI](https://www.ncbi.nlm.nih.gov/clinvar/variation/3047/)  [Variant summary: ATM c.496+5G>A alters a conserved nucleotide located ...](https://www.ncbi.nlm.nih.gov/clinvar/variation/3047/)  [10] [pmc.ncbi.nlm.nih.gov](https://pmc.ncbi.nlm.nih.gov/articles/PMC8152746/)  [11] [medlineplus.gov](https://medlineplus.gov/download/genetics/gene/atm.pdf)  [12] [ClinVar Miner Submissions for variant NM_000051.4(ATM):c.496G>C (p ...](https://clinvarminer.genetics.utah.edu/submissions-by-variant/NM_000051.4%28ATM%29%3Ac.496G%3EC%20%28p.Glu166Gln%29)  [Submissions for variant NM_000051.4(ATM):c.496G>C (p.Glu166Gln). dbSNP: rs1565357473. ...](https://clinvarminer.genetics.utah.edu/submissions-by-variant/NM_000051.4%28ATM%29%3Ac.496G%3EC%20%28p.Glu166Gln%29)  [13] [ClinVar Miner List of variants in gene ATM reported as likely pathogenic for ...](https://clinvarminer.genetics.utah.edu/variants-by-gene/ATM/condition/Hereditary%20cancer-predisposing%20syndrome/likely%20pathogenic)  [List of variants in gene ATM reported as likely pathogenic for Hereditary cancer-predisposing syndrome. Minimum submission review status: criteria provided.](https://clinvarminer.genetics.utah.edu/variants-by-gene/ATM/condition/Hereditary%20cancer-predisposing%20syndrome/likely%20pathogenic)  [14][Oxford Academic](https://academic.oup.com/hmg/article/7/10/1555/634692)  [15] [en.wikipedia.org ATM serine/threonine kinase - Wikipedia It phosphorylates several key proteins that initiate activation of the DNA damage checkpoint, leading to cell cycle arrest, ...](https://en.wikipedia.org/wiki/ATM_serine/threonine_kinase)  [16] [repository.cam.ac.uk [PDF] Minigene-based splicing analysis and ACMG/AMP-based tentative … For instance, the ATM variant c.332-1G>A (targeting exon 5 acceptor site) ... 496G>C (p.Glu166Gln) is reported in ClinVar as pathogenic. (late onset A-T ...](https://www.repository.cam.ac.uk/bitstreams/8944bf52-a8b2-4f23-991f-03097498eff3/download)  [17] [imrpress.com Late-onset autosomal recessive cerebellar ataxia and neuropathy ...](https://www.imrpress.com/journal/JIN/19/1/10.31083/j.jin.2020.01.1239/htm)  [Sanger sequencing confirmed the homozygous missense variant (c.496G > C) of ...](https://www.imrpress.com/journal/JIN/19/1/10.31083/j.jin.2020.01.1239/htm)  [18][markerdb.ca Showing biomarker card for c.496G>C (p.Glu166Gln) in ... - MarkerDB](https://markerdb.ca/sequence_variants/40643)  [Showing biomarker card for c.496G>C (p.Glu166Gln) in Serine-protein kinase atm ;](https://markerdb.ca/sequence_variants/40643) |
|  | Evaluation  **Summarization Accuracy & Completeness:**  **Hallucination Rate:**  **Readability & Usability:**  **Literature Coverage & Relevance:**  **Time Efficiency:** | |
| **3** | The ATM gene, located on chromosome 11q22-23, encodes a protein kinase crucial for activating cellular responses to DNA double-strand breaks. This kinase phosphorylates key players in the DNA damage-response pathway, ensuring genomic stability. The ATM c.496G>C variant, also denoted as IVS5+5G>A, is a single nucleotide substitution at the +5 position of intron 5 of the ATM gene. This variant is predicted to disrupt the natural donor splice site, potentially leading to abnormal gene splicing and reduced ATM protein expression. The guanine nucleotide altered in this variant is conserved across species, and its substitution is not observed in large population cohorts, suggesting it is not a common benign variant. The ATM c.496G>C variant has been observed in individuals with Ataxia-Telangiectasia (A-T), an autosomal recessive disorder characterized by progressive neurodegeneration, immunodeficiency, and increased cancer risk. In these cases, the variant was present in a compound heterozygous state with other pathogenic ATM variants. Functional studies have shown that cells harboring this variant exhibit chromosomal instability and reduced ATM protein levels, further supporting its pathogenic nature. However, the exact impact of this variant on cancer risk, particularly breast cancer, remains uncertain. While some studies suggest an association with increased breast cancer risk, others are inconclusive. The variant has also been observed in individuals with a personal history of endometrial cancer and a family history of breast, ovarian, colon, and endometrial cancers, but its direct role in these cancers is unclear. Given the limited evidence and the potential for abnormal splicing, the ATM c.496G>C variant is classified as likely pathogenic. Further research is needed to fully understand its clinical implications and to establish clear genotype-phenotype correlations. | [1] Dork T, Verhagen LM, et al. (2004).  [2] Verhagen LM, et al. (2009).  [3] Cadoo KA, et al. (2019). |
|  | Evaluation  **Summarization Accuracy & Completeness:**  **Hallucination Rate:**  **Readability & Usability:**  **Literature Coverage & Relevance:**  **Time Efficiency:** | |
| **4** | The ATM gene encodes a serine/threonine kinase involved in DNA damage response, cell cycle control, and genomic stability. Variants in ATM are associated with ataxia-telangiectasia, a neurodegenerative disorder, and an increased risk for malignancies such as breast cancer. The ATM c.496G>C variant is a single nucleotide substitution in this gene, but currently, no direct peer-reviewed studies have specifically characterized its pathogenicity, frequency, or clinical impact. In broader analyses of ATM variants, loss-of-function mutations have been linked to a twofold increased risk of breast cancer, and various ATM variants have been classified using ACMG/AMP guidelines to determine their clinical significance [1]. However, some ATM variants remain of uncertain significance due to a lack of functional validation or established genotype-phenotype correlations [2]. The absence of c.496G>C in large-scale sequencing studies suggests it may be a rare or novel variant, requiring further investigation for definitive classification. Given the established role of ATM in cancer predisposition and its function in DNA repair, further research including functional assays, segregation studies, and population frequency analysis is necessary to determine the clinical significance of this specific variant. In the absence of conclusive data, genetic counseling and consideration of broader ATM-related findings are recommended when assessing individual risk. | [1] Bueno-Martínez, E., Sanoguera-Miralles, L. (2022). Minigene-based splicing analysis and ACMG/AMP-based tentative classification of 56 ATM variants. [Pathology](https://app.scholarai.io/paper?paper_id=DOI:10.1002/path.5979&original_url=https%3A%2F%2Fpathsocjournals.onlinelibrary.wiley.com%2Fdoi%2Fabs%2F10.1002%2Fpath.5979).  [2] de Oliveira, K.C., Lindsey, S.C., Olivati, C. (2023). Germline genetic variants in pheochromocytoma/paraganglioma: single-center experience. [Endocrine Oncology](https://eo.bioscientifica.com/view/journals/eo/3/1/EO-22-0091.xml). |
|  | Evaluation  **Summarization Accuracy & Completeness:**  **Hallucination Rate:**  **Readability & Usability:**  **Literature Coverage & Relevance:**  **Time Efficiency:** | |
| **5** | The ATM (ataxia-telangiectasia mutated) gene encodes a protein kinase crucial for DNA damage response, cell cycle control, and genome stability. The ATM protein is activated by DNA double-strand breaks, leading to phosphorylation of downstream targets involved in DNA repair, cell cycle checkpoints, and apoptosis[6](https://pubmed.ncbi.nlm.nih.gov/30888062/)[4](https://www.ncbi.nlm.nih.gov/pmc/articles/PMC5056966/). Mutations in ATM are associated with ataxia-telangiectasia (A-T), a rare autosomal recessive disorder characterized by progressive cerebellar ataxia, immunodeficiency, telangiectasia, radiosensitivity, and cancer predisposition[6](https://pubmed.ncbi.nlm.nih.gov/30888062/). The variant ATM c.496G>C has not been specifically described in the literature. However, many ATM mutations can lead to loss of function of the ATM protein[6](https://pubmed.ncbi.nlm.nih.gov/30888062/). Approximately 14% of A-T-causing mutations are single-nucleotide changes that result in premature termination codons (PTCs)[1](https://pubmed.ncbi.nlm.nih.gov/15498871/). Aminoglycosides can induce PTC read-through, restoring levels of full-length ATM protein and correcting the phenotype of A-T cells[1](https://pubmed.ncbi.nlm.nih.gov/15498871/). In cases where ATM function is reduced but not absent (e.g. 5-20% of normal levels), slower neurological progression has been observed[1](https://pubmed.ncbi.nlm.nih.gov/15498871/). Considering the absence of direct functional data for ATM c.496G>C, its impact remains uncertain, and further investigation is warranted. | [[1](https://pubmed.ncbi.nlm.nih.gov/15498871/)] Correction of ATM gene function by aminoglycoside-induced read-through of premature termination codons. *Hum Mol Genet.* 2004 Nov 2;13(21):2679-88.  [[6](https://pubmed.ncbi.nlm.nih.gov/30888062/)] Two novel variants in the ATM gene causing ataxia‐telangiectasia, including a duplication of 90 kb: Utility of targeted next‐generation sequencing in detection of copy number variation. *Clin Case Rep*. 2019 Jul;7(7):1313-1318  [[4](https://www.ncbi.nlm.nih.gov/pmc/articles/PMC5056966/)] ATM function and its relationship with ATM gene mutations in chronic lymphocytic leukemia with the recurrent deletion (11q22.3-23.2). *Mol Cell Biochem*. 2017 Jan;426(1-2):133-144.  Citations:   1. <https://pubmed.ncbi.nlm.nih.gov/15498871/> 2. <https://pubmed.ncbi.nlm.nih.gov/11536012/> 3. <https://pubmed.ncbi.nlm.nih.gov/21901628/> 4. <https://www.ncbi.nlm.nih.gov/pmc/articles/PMC5056966/> 5. <https://www.ncbi.nlm.nih.gov/pmc/articles/PMC11196715/> 6. <https://pubmed.ncbi.nlm.nih.gov/30888062/> 7. <https://pubmed.ncbi.nlm.nih.gov/28477109/> 8. <https://pubmed.ncbi.nlm.nih.gov/23614766/> |
|  | Evaluation  **Summarization Accuracy & Completeness:**  **Hallucination Rate:**  **Readability & Usability:**  **Literature Coverage & Relevance:**  **Time Efficiency:** | |
