## Supplementary material for "Benchmarking generative AI tools for literature retrieval and summarization in genomic variant interpretation": BBS12 c.355G_A.docx

| **Tool** | **Summary** | **References** |
| --- | --- | --- |
| **1** | The BBS12 gene encodes a protein involved in the normal structure and function of primary cilia, which are essential for cellular signaling and development[1](https://www.ncbi.nlm.nih.gov/pmc/articles/PMC10352915/). BBS12 is classified as a chaperonin-like protein, which is required for the initial assembly of the BBSome, a multiprotein complex that regulates ciliary membrane protein composition[2](https://www.ncbi.nlm.nih.gov/pmc/articles/PMC9325507/). Bardet-Biedl syndrome (BBS) is a rare, pleiotropic disorder caused by mutations in genes affecting the cilia; it is estimated to occur in 1/140,000 to 1/160,000 individuals in Europe[3](https://pubmed.ncbi.nlm.nih.gov/37293956/). The BBS12 c.355G>A variant has been identified in individuals with Bardet-Biedl syndrome (BBS)[1](https://www.ncbi.nlm.nih.gov/pmc/articles/PMC10352915/)[3](https://pubmed.ncbi.nlm.nih.gov/37293956/). One study in a Romanian cohort found BBS12 variants in 35% of the families with BBS, with the homozygous BBS12 p.Arg355* variant being present in seven pedigrees[3](https://pubmed.ncbi.nlm.nih.gov/37293956/). Patients with truncating mutations in BBS genes, including BBS12, tend to exhibit more severe phenotypes, and mutations in chaperonin-like BBS proteins correlate with severe kidney impairment[2](https://www.ncbi.nlm.nih.gov/pmc/articles/PMC9325507/). Genetic analysis is important for the timely diagnosis of oligosymptomatic patients with BBS to prevent long-term complications; diversity in clinical expression can occur even with the same type of mutation[1](https://www.ncbi.nlm.nih.gov/pmc/articles/PMC10352915/). | [1](https://www.ncbi.nlm.nih.gov/pmc/articles/PMC10352915/) Khvostenko I, Rybalko T, and Kulinskyi V. Bardet-Biedl syndrome caused by compound heterozygosity in BBS12 gene: a case report of one family with three affected members. *Pediatr Endocrinol Diabetes Metab*. 2023 Jul 4;29(2):130-134.  [2](https://www.ncbi.nlm.nih.gov/pmc/articles/PMC9325507/) Palfy GG, and Gergely F. Bardet–Biedl syndrome: The pleiotropic role of the chaperonin‐like BBS6, 10, and 12 proteins. *Mol Genet Genomic Med*. 2022 Mar;10(3):e1886.  [3](https://pubmed.ncbi.nlm.nih.gov/37293956/) Albu A, Radu E, Diaconescu A, et al. Exome sequencing in a Romanian Bardet‐Biedl syndrome cohort revealed an overabundance of causal BBS12 variants. *Clin Genet*. 2023 Jun;103(6):567-577.  1 <https://www.ncbi.nlm.nih.gov/pmc/articles/PMC10352915/>  2 <https://www.ncbi.nlm.nih.gov/pmc/articles/PMC9325507/>  3 <https://pubmed.ncbi.nlm.nih.gov/37293956/>  4 <https://pubmed.ncbi.nlm.nih.gov/27485388/>  5 <https://pubmed.ncbi.nlm.nih.gov/30358434/>  6 <https://pubmed.ncbi.nlm.nih.gov/30070157/>  7 <https://pubmed.ncbi.nlm.nih.gov/26945629/>  8 <https://www.ncbi.nlm.nih.gov/pmc/articles/PMC3384976/> |
|  | Evaluation  **Summarization Accuracy & Completeness:**  **Hallucination Rate:**  **Readability & Usability:**  **Literature Coverage & Relevance:**  **Time Efficiency:** | |
| **2** | The BBS12 gene is a protein-coding gene associated with Bardet-Biedl syndrome type 12 (BBS12), a rare genetic disorder characterized by a range of clinical features including retinal degeneration, obesity, polydactyly, cognitive impairment, renal anomalies, and urogenital malformations. BBS12 is part of the chaperonin-containing T-complex (TRiC), a molecular chaperone complex that assists in protein folding upon ATP hydrolysis. This complex plays a crucial role in the assembly of the BBSome, which is involved in ciliogenesis and regulates transport vesicles to the cilia. The BBS12 protein is also involved in adipocyte differentiation, highlighting its multifaceted role in cellular processes. Defects in BBS12 are known to cause Bardet-Biedl syndrome type 12, underscoring the gene's importance in maintaining normal cellular functions.  The variant BBS12 c.355G>A is a missense mutation in the BBS12 gene, where guanine is replaced by adenine at nucleotide position 355. This variant results in an amino acid substitution, which can potentially alter the protein's structure and function. The specific impact of the BBS12 c.355G>A variant on protein function and its association with Bardet-Biedl syndrome or other clinical phenotypes is not well-documented in the available literature. The classification of this variant as pathogenic, likely pathogenic, or of uncertain significance is not explicitly stated in the reviewed sources.The frequency of the BBS12 c.355G>A variant in the general population is not reported in the available literature. However, Bardet-Biedl syndrome is a rare disorder, and pathogenic variants in BBS12 are estimated to account for approximately 8% of all BBS cases. The clinical significance of the BBS12 c.355G>A variant remains uncertain due to the lack of specific research findings. Further investigation, including functional studies and genetic analyses, is necessary to determine the variant's impact on protein function and its potential role in disease pathogenesis.  In summary, the BBS12 gene plays a critical role in protein folding and ciliogenesis, and its defects are associated with Bardet-Biedl syndrome type 12. The BBS12 c.355G>A variant is a missense mutation with an uncertain clinical significance. Additional research is needed to elucidate the variant's impact on protein function and its potential contribution to Bardet-Biedl syndrome or other related disorders. | Focsa, I. O., Budisteanu, M., Burloiu, C. M., Khan, S., Sadeghpour, A., Bohiltea, L. C., ... & Davis, E. E. (2021). A case of Bardet-Biedl syndrome caused by a recurrent variant in BBS12: A case report. Biomedical Reports, 15. |
|  | Evaluation  **Summarization Accuracy & Completeness:**  **Hallucination Rate:**  **Readability & Usability:**  **Literature Coverage & Relevance:**  **Time Efficiency:** | |
| **3** | The BBS12 gene encodes a protein that is a component of the BBSome complex, which plays a crucial role in ciliogenesis and intracellular trafficking. Variants in BBS12 are associated with Bardet-Biedl syndrome (BBS), a ciliopathy characterized by retinal dystrophy, obesity, polydactyly, renal abnormalities, and intellectual disability. The c.355G>A variant in BBS12 leads to an amino acid substitution, the clinical significance of which remains uncertain due to limited direct studies. Available genetic analyses of BBS12 variants have identified both pathogenic and benign mutations, with conflicting interpretations reported in ClinVar and other variant databases [1]. A study involving whole-exome sequencing in Tunisian patients identified a homozygous BBS12 variant, which was classified as benign according to ACMG guidelines, though the classification remains uncertain due to conflicting clinical reports [2]. Broader molecular analyses of BBS12 mutations in Bardet-Biedl syndrome families have contributed to the understanding of disease phenotypes but do not specifically address the c.355G>A variant [3]. Given the variable expressivity and possible oligogenic interactions in BBS, the pathogenicity of BBS12 variants can be influenced by additional genetic factors, requiring comprehensive genetic and functional studies to establish the role of c.355G>A in disease etiology. Further investigation, including functional assays and case-specific segregation studies, is recommended to determine its clinical relevance. | [1] Kheriji et al., 2023, [Whole-exome sequencing reveals novel variants of monogenic diabetes in Tunisia](https://www.frontiersin.org/articles/10.3389/fgene.2023.1224284/full);  [2] Chen et al., 2011, [Molecular analysis of Bardet-Biedl syndrome families](https://iovs.arvojournals.org/article.aspx?articleid=2187597);  [3] Lamont et al., 2016, [Expansion of phenotype and genotypic data in CRB2-related syndrome](https://www.nature.com/articles/ejhg201624). |
|  | Evaluation  **Summarization Accuracy & Completeness:**  **Hallucination Rate:**  **Readability & Usability:**  **Literature Coverage & Relevance:**  **Time Efficiency:** | |
| **4** | The BBS12 gene encodes a protein that functions as a molecular chaperone, assisting in the proper folding of other proteins through ATP hydrolysis. This protein is also implicated in adipocyte differentiation, influencing fat cell development. Mutations in BBS12 are associated with Bardet-Biedl syndrome (BBS), a ciliopathy characterized by features such as pigmentary retinopathy, early-onset obesity, polydactyly, and renal anomalies. The c.355G>A variant in BBS12 results in a glycine-to-serine substitution at position 119 (p.Gly119Ser). This specific variant has been identified in patients exhibiting phenotypes related to BBS; however, its pathogenicity remains uncertain. According to a study by Kheriji et al., the c.355G>A variant was found in a patient alongside a variant in the TTC8 gene, another gene associated with BBS. The c.355G>A variant is rare globally, with a frequency of approximately 0.00016, and is currently classified as benign according to the American College of Medical Genetics and Genomics (ACMG) guidelines. Nonetheless, conflicting interpretations have been reported in ClinVar, where it is listed as a variant of uncertain significance (VUS) (ClinVar ID: 215543). Given the limited and inconclusive evidence regarding the pathogenicity of the BBS12 c.355G>A variant, further functional studies and comprehensive genetic analyses are recommended to elucidate its clinical impact. | [ncbi.nlm.nih.gov BBS12 Bardet-Biedl syndrome 12 [Homo sapiens (human)] - Gene](https://www.ncbi.nlm.nih.gov/gene?Cmd=DetailsSearch&Db=gene&Term=166379)  [4 gennaio 2025 — BBS12 inactivation increases glucd ...](https://www.ncbi.nlm.nih.gov/gene?Cmd=DetailsSearch&Db=gene&Term=166379)  [frontiersin.org [PDF] Whole-exome sequencing reveals novel variants of monogenic ...](https://www.frontiersin.org/journals/genetics/articles/10.3389/fgene.2023.1224284/pdf)  [14 dicembre 2023 — The patient P4 carries two potential pathogenic ...](https://www.frontiersin.org/journals/genetics/articles/10.3389/fgene.2023.1224284/pdf)  [databases.lovd.nl Global Variome shared LOVD](https://databases.lovd.nl/shared/variants/0000261403)  [15 gennaio 2018 — Variant #0000261403 (NC_000004.11:g.123663402G>A, NM_001178007.1:c.355G>A (BBS12)).](https://databases.lovd.nl/shared/variants/0000261403)  [ncbi.nlm.nih.gov VCV000215543.29 - ClinVar - NCBI Identifiers. NM_152618.3(BBS12):c.355G>A (p.Gly119Ser). Variation ID: 215543 Accession:](https://www.ncbi.nlm.nih.gov/clinvar/variation/215543)  [genecards.org BBS12 Gene - Bardet-Biedl Syndrome 12 - GeneCards](https://www.genecards.org/cgi-bin/carddisp.pl?gene=BBS12)  [The encoded protein is a molecular chaperone that aids in protein folding upon ATP](https://www.genecards.org/cgi-bin/carddisp.pl?gene=BBS12)  [genecards.org BBS12 Gene - Bardet-Biedl Syndrome 12 - GeneCards](https://www.genecards.org/cgi-bin/carddisp.pl?gene=BBS12)  [databases.lovd.nl Unique variants in the BBS12 gene - Global Variome shared LOVD](https://databases.lovd.nl/shared/variants/BBS12/unique)  [BBS12(NM_001178007.2):c.355G>A (p.G119S), BBS12(NM_152618.3):c.355G>A (p ... BBS12 c.898C>T, p.Q300X, BBS12, variant 1: c.898C>T/p.Q300*, variant 2: c ...](https://databases.lovd.nl/shared/variants/BBS12/unique)  [en.wikipedia.org](https://en.wikipedia.org/wiki/BBS12)  [BBS12 - Wikipedia BBS12 · photoreceptor cell maintenance · ...](https://en.wikipedia.org/wiki/BBS12)  [myriad.com](https://myriad.com/womens-health/diseases/bardet-biedl-syndrome-bbs12-related/)  [Bardet-Biedl syndrome, BBS12-related - Myriad Genetics](https://myriad.com/womens-health/diseases/bardet-biedl-syndrome-bbs12-related/)  [Bardet-Biedl syndrome (BBS), caused by harmful genetic changes (mutations) in the BBS12 gene, is an inherited disease that causes vision problems.](https://myriad.com/womens-health/diseases/bardet-biedl-syndrome-bbs12-related/)  [olida.ibsquare.be](https://olida.ibsquare.be/detail/SmallVariant/155/)  [Details for BBS12:c.355G>A, p.Gly119Ser - OLIDA](https://olida.ibsquare.be/detail/SmallVariant/155/)  [Details for BBS12:c.355G>A, p.Gly119Ser ; E ...](https://olida.ibsquare.be/detail/SmallVariant/155/)  [Malacards Bardet-Biedl Syndrome 12 (BBS12) - MalaCardsy, ...](https://www.malacards.org/card/bardet_biedl_syndrome_12)  [uniprot.org](https://www.uniprot.org/uniprotkb/Q6ZW61/entry)  [medlineplus.gov](https://medlineplus.gov/genetics/condition/bardet-biedl-syndrome/)  [Bardet-Biedl syndrome - Genetics - MedlinePlus](https://medlineplus.gov/genetics/condition/bardet-biedl-syndrome/)  [The proteins produced from BBS genes are involved in the maintenance and function ...](https://medlineplus.gov/genetics/condition/bardet-biedl-syndrome/)  [q6zw61 · bbs12_human - UniProt](https://www.uniprot.org/uniprotkb/Q6ZW61/entry)  [Component of the chaperonin-containing T-complex (TRiC), a molecular chaperone complex that assists the folding of proteins upon ATP hydrolysis.](https://www.uniprot.org/uniprotkb/Q6ZW61/entry)  [v1.marrvel.org BBS12 GENE - MARRVEL: Search Result](https://v1.marrvel.org/search/gene/BBS12)  [MIM number: 610683. Description: Stoetzel et al. (2007)](https://v1.marrvel.org/search/gene/BBS12)  [mseqdr.org MSeqDR Central Portal BBS12, 1, BBS12_HUMAN, c.355G>A, p.G119S, non-syn, rs77731085, 0.0028, A=3/G=8597;A=51/G=4355;A=54/G=12952, lod=89:449, DAMAGING, D, -, het, 5. 20, 4, 123663402 ...](https://www.mseqdr.org/portal.php?dbsource=genomic&name=BBS12)  [europepmc.org](https://europepmc.org/article/pmc/3176075)  [Molecular analysis of Bardet-Biedl syndrome families - Europe PMC….](https://europepmc.org/article/pmc/3176075) |
|  | Evaluation  **Summarization Accuracy & Completeness:**  **Hallucination Rate:**  **Readability & Usability:**  **Literature Coverage & Relevance:**  **Time Efficiency:** | |
| **5** | The genomic variant c.355G>A p.Gly119Ser rs77731085 is located in the BBS12 gene, which is associated with Bardet-Biedl syndrome (BBS), an autosomal recessive disorder. BBS is characterized by a spectrum of clinical features including retinal degeneration, obesity, polydactyly, cognitive impairment, renal abnormalities, and other symptoms such as diabetes mellitus, neurological issues, and developmental delays [1]. The BBS12 gene plays a crucial role in the structure and function of cilia, which are important for cell signaling and organ development.  The c.355G>A p.Gly119Ser variant results in the substitution of a glycine residue with a serine at the 119th amino acid position of the BBS12 protein. This missense change is located in a region of the protein that is important for its function, potentially impacting the stability and activity of the protein. Despite this, the variant is reported in ClinVar with four benign submissions, suggesting that it may not have a significant pathogenic effect on its own.  In the context of patient P4, the variant was identified in a homozygous state, which is typically necessary for the manifestation of BBS symptoms due to its autosomal recessive inheritance pattern. However, the variant is classified as benign according to the ACMG guidelines, and there are conflicting interpretations reported in ClinVar [2]. This suggests that the pathogenicity of the variant may be influenced by other genetic or environmental factors, and its presence alone may not be sufficient to cause disease.  In summary, the c.355G>A p.Gly119Ser rs77731085 variant in the BBS12 gene has been associated with BBS, but its classification as benign in ClinVar and the conflicting interpretations of its pathogenicity indicate that further research is needed to fully understand its clinical significance. The BBS12 gene's role in ciliary function underscores the importance of this variant in disorders related to ciliary dysfunction. | [[1]](https://pubmed.ncbi.nlm.nih.gov/28912962/) Costa KA et al. (2017). "Gene panel sequencing in Brazilian patients with retinitis pigmentosa." International journal of retina and vitreous, 3()  [[2]](https://pubmed.ncbi.nlm.nih.gov/38162681/) Kheriji N et al. (2023). "Whole-exome sequencing reveals novel variants of monogenic diabetes in Tunisia: impact on diagnosis and healthcare management." Frontiers in genetics, 14()  [[3]](https://pubmed.ncbi.nlm.nih.gov/30309378/) Paracchini L et al. (2018). "Exome sequencing in an Italian family with Alzheimer's disease points to a role for seizure-related gene 6 (SEZ6) rare variant R615H." Alzheimer's research & therapy, 10(1) [SUPP]  [[4]](https://pubmed.ncbi.nlm.nih.gov/35495172/) Aguiar T et al. (2022). "Unraveling the Genetic Architecture of Hepatoblastoma Risk: Birth Defects and Increased Burden of Germline Damaging Variants in Gastrointestinal/Renal Cancer Predisposition and DNA Repair Genes." Frontiers in genetics, 13() [SUPP]  [[5]](https://pubmed.ncbi.nlm.nih.gov/27898983/) Roberts L et al. (2016). "Molecular Diagnosis of Inherited Retinal Diseases in Indigenous African Populations by Whole-Exome Sequencing." Investigative ophthalmology & visual science, 57(14) [SUPP]  [[6]](https://pubmed.ncbi.nlm.nih.gov/26740555/) Richards AL et al. (2016). "Exome arrays capture polygenic rare variant contributions to schizophrenia." Human molecular genetics, 25(5) [SUPP]  [[7]](https://pubmed.ncbi.nlm.nih.gov/26659599/) Holehouse AS et al. (2015). "Reproducible Analysis of Post-Translational Modifications in Proteomes--Application to Human Mutations." PloS one, 10(12) [SUPP]  [[8]](https://pubmed.ncbi.nlm.nih.gov/36192439/) Moreno-Ruiz N et al. (2022). "Assessing the digenic model in rare disorders using population sequencing data." European journal of human genetics : EJHG, 30(12) [SUPP]  [[9]](https://pubmed.ncbi.nlm.nih.gov/28719003/) Leonenko G et al. (2017). "Mutation intolerant genes and targets of FMRP are enriched for nonsynonymous alleles in schizophrenia." American journal of medical genetics. Part B, Neuropsychiatric genetics : the official publication of the International Society of Psychiatric Genetics, 174(7) [SUPP] |
|  | Evaluation  **Summarization Accuracy & Completeness:**  **Hallucination Rate:**  **Readability & Usability:**  **Literature Coverage & Relevance:**  **Time Efficiency:** | |
