## Supplementary material for "Benchmarking generative AI tools for literature retrieval and summarization in genomic variant interpretation": BMPR2 p.R491W.docx

| **Tool** | **Summary** | **References** |
| --- | --- | --- |
| **1** | The BMPR2 gene encodes the bone morphogenetic protein receptor type II, a critical component of the transforming growth factor-beta (TGF-β) superfamily involved in regulating cell growth and differentiation. Mutations in BMPR2 are the most common genetic cause of heritable pulmonary arterial hypertension (PAH), accounting for approximately 75% of familial cases and about 20% of idiopathic cases. The p.R491W variant results from a missense mutation where arginine is replaced by tryptophan at position 491 within the kinase domain of the BMPR-II protein. This substitution disrupts a salt bridge with Glu386 and hydrogen bonds to the backbone carbonyl of Asp485, leading to structural clashes that likely impair receptor function. Functional analyses indicate that such mutations in the kinase domain can lead to a loss of function, contributing to the pathogenesis of PAH. Clinically, individuals carrying BMPR2 mutations, including p.R491W, often present with more severe hemodynamic impairments and at a younger age compared to non-carriers. However, the penetrance of BMPR2 mutations is incomplete, suggesting that additional genetic or environmental factors may influence disease manifestation. While the p.R491W variant has been identified in multiple PAH patients, comprehensive studies specifically detailing its prevalence and distinct clinical impact remain limited. Therefore, further research is warranted to elucidate the precise role of the p.R491W mutation in PAH pathogenesis and to inform potential therapeutic strategies. | <https://pmc.ncbi.nlm.nih.gov/articles/PMC2715351/>  <https://pmc.ncbi.nlm.nih.gov/articles/PMC4737700/>  <https://pmc.ncbi.nlm.nih.gov/articles/PMC6892941/>  <https://respiratory-research.biomedcentral.com/articles/10.1186/1465-9921-12-99>  <https://www.researchgate.net/publication/7777380_BMPR2_mutations_have_short_lifetime_expectancy_in_primary_pulmonary_hypertension>  <https://www.researchgate.net/publication/41121888_Study_of_the_BMPR2_Gene_in_Patients_with_Pulmonary_Arterial_Hypertension>  <https://www.nature.com/articles/s41467-021-25248-5>  <https://www.nature.com/articles/s41598-019-54830-7>  <https://www.biorxiv.org/content/10.1101/786756v1.full.pdf>  <https://onlinelibrary.wiley.com/doi/pdf/10.1002/humu.20285>  <https://pmc.ncbi.nlm.nih.gov/articles/PMC6160702/>  <https://omim.org/entry/600799>  <https://journals.lww.com/cmj/fulltext/2008/03010/a_novel_mutation_in_the_bmpr2_gene_in_familial.4.aspx>  <https://openaccess.sgul.ac.uk/id/eprint/111104/1/Southgate%20et%20al_NRC2019_accepted.pdf>  <https://openaccess.sgul.ac.uk/id/eprint/110211/7/Machado%20et%20al.%202015%20(Supplementary%20Info).pdf>  <https://databases.lovd.nl/shared/variants/BMPR2>  <https://www.researchgate.net/publication/265034009_BMPR2_Mutations_in_Six_Taiwanese_Patients_with_Idiopathic_Pulmonary_Arterial_Hypertension/figures?lo=1>  <https://www.researchgate.net/publication/265034009_BMPR2_Mutations_in_Six_Taiwanese_Patients_with_Idiopathic_Pulmonary_Arterial_Hypertension>  <https://www.mdpi.com/1422-0067/19/9/2499>  <https://d-nb.info/1163977969/34>  <https://publications.ersnet.org/content/erj/34/5/1100> |
|  | Evaluation  **Summarization Accuracy & Completeness:**  **Hallucination Rate:**  **Readability & Usability:**  **Literature Coverage & Relevance:**  **Time Efficiency:** | |
| **2** | The BMPR2 gene encodes the bone morphogenetic protein receptor type 2, a serine/threonine kinase that plays a key role in the transforming growth factor-beta (TGF-β) signaling pathway, which regulates cell growth, apoptosis, and differentiation in vascular endothelial and smooth muscle cells. BMPR2 mutations are the most common genetic cause of heritable pulmonary arterial hypertension (HPAH), with approximately 70% of familial and 15–40% of sporadic idiopathic cases attributed to pathogenic variants in this gene [1]. The missense variant p.R491W (arginine to tryptophan at codon 491) has been identified in individuals with familial pulmonary arterial hypertension and is located within the highly conserved kinase domain of the receptor, which is critical for downstream signaling. Although this specific variant is rare and not widely reported in large population databases, its location and physicochemical change suggest a probable deleterious effect on protein function. Evidence from structural analyses and mutagenesis studies supports that substitutions at this residue may disrupt kinase activity, compromising the receptor's capacity to transduce anti-proliferative signals in the pulmonary vasculature. While the p.R491W variant itself is infrequently reported, broader assessments of BMPR2 mutations underscore their significant phenotypic variability, incomplete penetrance, and age-dependent expressivity, which complicate genotype-phenotype correlations [2]. The variant has not been included in larger variant review cohorts with individual functional confirmation, thus direct experimental evidence specific to p.R491W remains limited and further functional characterization is warranted. Given the established pathogenicity of mutations within this region of BMPR2 and the clinical context in which p.R491W has been observed, this variant is likely pathogenic; however, due to the absence of functional studies directly validating this effect, it is currently best classified as a variant of uncertain significance (VUS) pending further evidence. Continued research and data sharing through variant databases are recommended to refine the interpretation of such rare BMPR2 variants. | [1] [Machado et al. 2006](https://onlinelibrary.wiley.com/doi/abs/10.1002/humu.20285)  [2] [Machado et al. 2015](https://onlinelibrary.wiley.com/doi/abs/10.1002/humu.22904) |
|  | Evaluation  **Summarization Accuracy & Completeness:**  **Hallucination Rate:**  **Readability & Usability:**  **Literature Coverage & Relevance:**  **Time Efficiency:** | |
| **3** | The **BMPR2** gene encodes the bone morphogenetic protein receptor type II, a crucial component in the BMP signaling pathway, which plays a significant role in regulating cellular growth, differentiation, and apoptosis. Mutations in **BMPR2** are strongly associated with pulmonary arterial hypertension (PAH), a severe and often fatal disease characterized by elevated blood pressure in the pulmonary arteries. The variant **p.R491W** is a specific mutation within this gene, but there is limited direct research specifically addressing this variant in the context of PAH or other diseases. Generally, mutations in **BMPR2** can lead to impaired signaling, which is linked to endothelial dysfunction and aberrant vascular remodeling in PAH[1][6]. While **BMPR2** mutations are a leading genetic cause of PAH, not all mutation carriers develop the disease, suggesting that additional factors, such as environmental triggers or other genetic modifiers, may influence disease manifestation[6]. Research on **BMPR2** has also highlighted the role of circular RNAs, like **circ5078**, derived from the **BMPR2** gene, which can influence endothelial proliferation and stress responses in PAH[3][4]. Further investigation is needed to fully understand the impact of specific mutations like **p.R491W** on **BMPR2** function and disease risk. | [1] <https://www.semanticscholar.org/paper/73e7b40ee03de5cdccb9c050905ee7e29e4d21cc>  [2] <https://www.semanticscholar.org/paper/30c810ae4f1dd46a0dbdae2dfc817234882727e0>  [3] <https://www.semanticscholar.org/paper/66349a558336841304bed5a08e4662427e95dfbb>  [4] <https://www.semanticscholar.org/paper/ff1a65bdb85e5f55fbec521493ebb5c1bbf8b91e>  [5] <https://www.semanticscholar.org/paper/c662fb4f45aab39743d7fe66707ebf528f48ae9a>  [6] <https://www.ncbi.nlm.nih.gov/pmc/articles/PMC6811766/>  [7] <https://www.ncbi.nlm.nih.gov/pmc/articles/PMC10543474/>  [8] <https://pubmed.ncbi.nlm.nih.gov/37354366/> |
|  | Evaluation  **Summarization Accuracy & Completeness:**  **Hallucination Rate:**  **Readability & Usability:**  **Literature Coverage & Relevance:**  **Time Efficiency:** | |
| **4** | The BMPR2 gene encodes the bone morphogenetic protein receptor type II, which is a member of the TGF-beta superfamily of receptors. BMPR2 plays a crucial role in the regulation of cellular growth and differentiation, particularly in the vascular system. Mutations in BMPR2 are known to be associated with heritable pulmonary arterial hypertension (PAH), a progressive disorder characterized by the narrowing of the small pulmonary arteries, leading to increased pulmonary vascular resistance, right ventricular failure, and potentially death.  The genomic variant c.1471C>T p.R491W rs137852746 in the BMPR2 gene is a missense mutation that results in the substitution of arginine with tryptophan at position 491 of the protein. This mutation has been identified in multiple individuals with PAH and is considered pathogenic. The R491W mutation is located within the kinase domain of BMPR2, which is essential for its function in signal transduction. The alteration of this highly conserved residue is predicted to disturb the kinase activity of BMPR2, potentially hindering the downstream activation cascade within the BMPR2 pathway [6].  The variant has been reported in several studies and is associated with familial PAH, indicating a high penetrance of the disease when this mutation is present. It has been observed that the R491W mutation segregates within families, defining a familial disease with a clear pattern of inheritance [4]. Furthermore, the mutation has been identified in unrelated patients with idiopathic PAH, suggesting that it also contributes to the sporadic form of the disease [7].  Functional studies have shown that mutations clustering at the C1 dimer interface in the BMPR2 kinase domain, including R491W, disrupt BMPR2 kinase dimerization and downstream SMAD signaling. This disruption leads to a loss of function and a dominant-negative effect on BMPR2 signaling, which is critical for maintaining normal pulmonary vascular structure and function [5].  In summary, the c.1471C>T p.R491W rs137852746 variant in the BMPR2 gene is a pathogenic missense mutation that impairs the receptor's kinase activity and disrupts normal signaling pathways. This mutation is associated with both familial and idiopathic forms of PAH and has a significant impact on the clinical phenotype of affected individuals. ClinVar reports a total of 5 pathogenic submissions for this variant, further supporting its pathogenicity in PAH. | [[1]](https://pubmed.ncbi.nlm.nih.gov/33066286/) van den Heuvel LM et al. (2020). "Genetic Evaluation in a Cohort of 126 Dutch Pulmonary Arterial Hypertension Patients." Genes, 11(10)  [[2]](https://pubmed.ncbi.nlm.nih.gov/21801371/) Pfarr N et al. (2011). "Hemodynamic and clinical onset in patients with hereditary pulmonary arterial hypertension and BMPR2 mutations." Respiratory research, 12(1)  [[3]](https://pubmed.ncbi.nlm.nih.gov/29743074/) Yang H et al. (2018). "Genetic analyses in a cohort of 191 pulmonary arterial hypertension patients." Respiratory research, 19(1)  [[4]](https://pubmed.ncbi.nlm.nih.gov/29843651/) Abou Hassan OK et al. (2018). "Clinical and genetic characteristics of pulmonary arterial hypertension in Lebanon." BMC medical genetics, 19(1)  [[5]](https://pubmed.ncbi.nlm.nih.gov/34400635/) Agnew C et al. (2021). "Structural basis for ALK2/BMPR2 receptor complex signaling through kinase domain oligomerization." Nature communications, 12(1)  [[6]](https://pubmed.ncbi.nlm.nih.gov/14985116/) Zhicheng J et al. (2004). "Bone morphogenetic protein receptor-II mutation Arg491Trp causes malignant phenotype of familial primary pulmonary hypertension." Biochemical and biophysical research communications, 315(4)  [[7]](https://pubmed.ncbi.nlm.nih.gov/28388887/) Higasa K et al. (2017). "A burden of rare variants in BMPR2 and KCNK3 contributes to a risk of familial pulmonary arterial hypertension." BMC pulmonary medicine, 17(1)  [[8]](https://pubmed.ncbi.nlm.nih.gov/32966279/) Jang AY et al. (2020). "Prevalence and clinical features of bone morphogenetic protein receptor type 2 mutation in Korean idiopathic pulmonary arterial hypertension patients: The PILGRIM explorative cohort." PloS one, 15(9)  [[9]](https://pubmed.ncbi.nlm.nih.gov/30084161/) Abbasi Y et al. (2018). "Exome data clouds the pathogenicity of genetic variants in Pulmonary Arterial Hypertension." Molecular genetics & genomic medicine, 6(5)  [[10]](https://pubmed.ncbi.nlm.nih.gov/15059534/) Jing ZC et al. (2004). "[Clinical and genetic characteristics of a Chinese family of primary pulmonary hypertension]." Zhonghua yi xue za zhi, 84(3)  [[11]](https://journals.ajsrp.com/index.php/jaevs/article/view/2018) Mohamed Saeed Elhassan Mohamed, Hind AbdelAziz Eln et al. (2019). "In silico analysis of single nucleotide polymorphism (SNPs) in human [BMPR2] gene: دراسة وتحليل الجين (BMPR2) عن طريق (SNPs) باستخدام برمجيات الحاسب الآلي" Journal of agricultural, environmental and veterinary sciences,  [[12]](https://pubmed.ncbi.nlm.nih.gov/31797984/) Chaikuad A et al. (2019). "Structural consequences of BMPR2 kinase domain mutations causing pulmonary arterial hypertension." Scientific reports, 9(1)  [[13]](https://pubmed.ncbi.nlm.nih.gov/37681932/) Sanchez-Duffhues G et al. (2023). "Human iPSCs as Model Systems for BMP-Related Rare Diseases." Cells, 12(17)  [[14]](https://pubmed.ncbi.nlm.nih.gov/28619995/) Tojais NF et al. (2017). "Codependence of Bone Morphogenetic Protein Receptor 2 and Transforming Growth Factor-beta in Elastic Fiber Assembly and Its Perturbation in Pulmonary Arterial Hypertension." Arteriosclerosis, thrombosis, and vascular biology, 37(8)  [[15]](https://pubmed.ncbi.nlm.nih.gov/26387786/) Machado RD et al. (2015). "Pulmonary Arterial Hypertension: A Current Perspective on Established and Emerging Molecular Genetic Defects." Human mutation, 36(12) |
|  | Evaluation  **Summarization Accuracy & Completeness:**  **Hallucination Rate:**  **Readability & Usability:**  **Literature Coverage & Relevance:**  **Time Efficiency:** | |
| **5** | The BMPR2 gene encodes the bone morphogenetic protein receptor type 2, a serine/threonine receptor kinase involved in the TGF-beta signaling pathway. This receptor is crucial for various cellular functions, including osteogenesis, cell growth, and differentiation. BMPR2 binds bone morphogenetic proteins (BMPs), which are part of the TGF-beta superfamily of ligands, and plays a significant role in paracrine signaling. The BMPR2 protein is expressed on granulosa cells and is essential for follicle development, with its expression regulated by hormones such as estrogen and follicle-stimulating hormone (FSH). Mutations in BMPR2 are strongly associated with pulmonary arterial hypertension (PAH), a condition characterized by abnormally high blood pressure in the pulmonary arteries. Over 350 mutations in BMPR2 have been identified as causative for PAH, with these mutations potentially promoting cell proliferation or preventing cell death, leading to an overgrowth of cells in the smallest arteries throughout the lungs. The p.R491W variant in BMPR2 is classified as pathogenic or likely pathogenic, with evidence suggesting it alters the amino acid electric status, contributing to its pathogenicity. This variant is notably absent in the general reference population but represents a hot-spot site in PAH patients, occurring in approximately 14.6% of cases. The frequency of rare BMPR2 variants, including p.R491W, is significantly higher in PAH patients compared to the general population, highlighting their role in the disease's pathogenesis. The exact mechanisms by which BMPR2 mutations cause PAH remain unclear, but they are thought to disrupt normal signaling pathways, leading to vascular remodeling and increased pulmonary artery pressure. Genetic screening for BMPR2 variants is recommended for PAH patients and their families to identify those at risk and guide appropriate treatment strategies. Further research is needed to fully understand the impact of the p.R491W variant and other BMPR2 mutations on the development and progression of PAH. | [1] Machado RD, Pauciulo MW, Thomson JR, Lane KB, Morgan NV, Wheeler L, Phillips JA 3rd, Newman J, Williams D, Galie N, Manes A, McNeil K, Yacoub M, Mikhail G, Rogers P, Corris P, Humbert M, Donnai D, Martensson G, Tranebjaerg L, Loyd JE, Trembath RC, Nichols WC. BMPR2 haploinsufficiency as the inherited molecular mechanism for primary pulmonary hypertension.  [2] Pfarr N, Fischer C, Ehlken N, Becker-Grünig T, López-González V, Gorenflo M, Hager A, Hinderhofer K, Miera O, Nagel C, Schranz D, Grünig E. Hemodynamic and genetic analysis in children with idiopathic, heritable, and congenital heart disease associated pulmonary arterial hypertension. Respir Res. 2013 Jan 9;14(1):3. doi: 10.1186/1465-9921-14-3.  [3] Oriaku I, LeSieur MN, Nichols WC, Barrios R, Elliott CG, Frost A. A novel BMPR2 mutation with widely disparate heritable pulmonary arterial hypertension clinical phenotype. Pulm Circ. 2020; 10(3): 2045894020931315.  [4] Liu D, Morrell NW. Genetics and the molecular pathogenesis of pulmonary arterial hypertension. Curr Hypertens Rep. 2013; 15(6): 632-637.  [5] Zhang R, Chen S, Han P, et al. Whole exome sequencing identified a homozygous novel variant in CEP290 gene causes Meckel syndrome. |
|  | Evaluation  **Summarization Accuracy & Completeness:**  **Hallucination Rate:**  **Readability & Usability:**  **Literature Coverage & Relevance:**  **Time Efficiency:** | |
