## Supplementary material for "Benchmarking generative AI tools for literature retrieval and summarization in genomic variant interpretation": CFTR c.571T_G.docx

| **Tool** | **Summary** | **References** |
| --- | --- | --- |
| **1** | The CFTR gene encodes the cystic fibrosis transmembrane conductance regulator protein, which functions as a chloride channel crucial for maintaining fluid balance across epithelial surfaces. Variants in this gene can lead to cystic fibrosis (CF), an autosomal recessive disorder characterized by impaired chloride ion transport, resulting in thickened mucus secretions and multiorgan dysfunction, particularly affecting the lungs and pancreas. The CFTR c.571T>G variant, also known as p.Leu191Arg or L191R, is a missense variant that alters the amino acid leucine to arginine at position 191 in the CFTR protein. This variant is not well-characterized in the literature, and its clinical significance remains uncertain.  The CFTR gene, located on chromosome 7q31.2, spans approximately 190 kb and contains 27 exons. It encodes a protein of 1,480 amino acids that functions as a cyclic AMP-regulated chloride channel. The CFTR protein comprises two membrane-spanning domains, two nucleotide-binding domains, and a regulatory domain. Variants in CFTR can disrupt the protein's structure and function, leading to CF or CFTR-related disorders (CFTR-RD). Over 2,000 variants have been identified in CFTR, with varying impacts on protein function and clinical severity. These variants are classified based on their effects on CFTR protein production, processing, and function. Class I variants result in no protein production, Class II variants lead to defective protein processing and trafficking, Class III variants affect channel regulation, Class IV variants reduce channel conductance, Class V variants reduce the number of functional proteins at the cell surface, and Class VI variants decrease protein stability.  The CFTR c.571T>G variant is rare, and its frequency in the general population is not well documented. It is not listed among the 23 pathogenic variants recommended for carrier screening by the American College of Medical Genetics and Genomics (ACMG) and the American College of Obstetricians and Gynecologists (ACOG). The clinical impact of this variant is not clearly established, and it has not been extensively studied in the context of CF or CFTR-RD. In silico prediction tools suggest that this variant may be damaging, but these predictions require validation through functional studies and clinical correlation.  Given the limited information available, the classification of the CFTR c.571T>G variant remains uncertain. It is not definitively categorized as pathogenic, likely pathogenic, or benign. Further research, including functional assays and genotype-phenotype correlation studies, is necessary to determine the clinical significance of this variant. Individuals with this variant should be evaluated in the context of their clinical presentation and family history. Genetic counseling is recommended to discuss the potential implications of this variant and the need for further testing.  In conclusion, the CFTR c.571T>G variant is a rare and poorly characterized missense variant with an uncertain clinical impact. Its classification and significance in CF or CFTR-RD require further investigation. Genetic counseling and additional studies are essential to understand the role of this variant in disease pathogenesis and to guide clinical management. | [1] Watson MS, Cutting GR, Desnick RJ, et al. Cystic fibrosis population carrier screening: 2004 revision of American College of Medical Genetics mutation panel. Genet Med. 2004;6(5):387-391.  [2] Bobadilla JL, Macek M Jr, Fine JP, et al. Cystic fibrosis: a worldwide analysis of CFTR mutations—correlation with incidence data and application to screening. Hum Mutat. 2002;19(6):575-606.  [3] Heim RA, Sugarman EA, Allitto BA. Improved detection of cystic fibrosis mutations in the heterogeneous U.S. population using an expanded, pan-ethnic mutation panel. Genet Med. 2001;3(3):168-176.  [4] Moskowitz SM, Chmiel JF, Sternen DL, et al. Clinical practice and genetic counseling for cystic fibrosis and CFTR-related disorders. Genet Med. 2008;10(12):851-868.  [5] Sugarman EA, Rohlfs EM, Silverman LM, et al. CFTR mutation distribution among U.S. Hispanic and African American individuals: evaluation in cystic fibrosis patient and carrier screening populations. Genet Med. 2004;6(5):392-399. |
|  | Evaluation  **Summarization Accuracy & Completeness:**  **Hallucination Rate:**  **Readability & Usability:**  **Literature Coverage & Relevance:**  **Time Efficiency:** | |
| **2** | There was an issue retrieving peer-reviewed data for the CFTR c.571T>G variant from the literature database. Due to this limitation, I will summarize what is generally known about CFTR and this type of variant while emphasizing uncertainties and recommending further investigation.  The CFTR (Cystic Fibrosis Transmembrane Conductance Regulator) gene encodes a chloride channel essential for ion and water transport across epithelial membranes, with dysfunction leading to cystic fibrosis (CF) and related conditions. The c.571T>G variant, located within the CFTR coding region, represents a single nucleotide substitution that may alter protein function, but its precise pathogenicity remains uncertain due to the lack of conclusive evidence. Variants in CFTR can range from benign polymorphisms to disease-causing mutations, classified according to their impact on protein production, processing, or function. If c.571T>G results in an amino acid change, it may influence CFTR folding, trafficking, or channel activity, potentially contributing to CFTR-related disorders. However, no direct studies establishing its clinical significance or population frequency were identified in available databases, necessitating further investigation through functional assays and patient-based studies. Given the absence of variant-specific research, clinicians should interpret this finding with caution, considering segregation analysis, in silico predictions, and additional CFTR screening when relevant. Further research is required to determine whether this variant contributes to disease phenotypes or remains a rare, functionally neutral polymorphism. |  |
|  | Evaluation  **Summarization Accuracy & Completeness:**  **Hallucination Rate:**  **Readability & Usability:**  **Literature Coverage & Relevance:**  **Time Efficiency:** | |
| **3** | The CFTR gene encodes a transmembrane protein that functions as a chloride channel and regulates the flow of ions and water across cell membranes, crucial for the function of lungs, pancreas, and other organs[2](https://www.ncbi.nlm.nih.gov/pmc/articles/PMC8167067/)[8](https://pubmed.ncbi.nlm.nih.gov/33331662/). Mutations in the CFTR gene can lead to cystic fibrosis (CF), a genetic disease characterized by abnormal salt and water transport, resulting in thick mucus buildup and organ damage[2](https://www.ncbi.nlm.nih.gov/pmc/articles/PMC8167067/)[5](https://www.ncbi.nlm.nih.gov/pmc/articles/PMC8584557/). The c.571T>G variant in the CFTR gene has not been specifically characterized in peer-reviewed publications. Research on other CFTR variants indicates that the functional consequences and clinical implications can vary widely depending on the specific mutation and its impact on CFTR protein trafficking, stability, and channel function[4](https://www.ncbi.nlm.nih.gov/pmc/articles/PMC10063961/). Some variants may disrupt protein folding and trafficking, leading to reduced amounts of CFTR protein at the cell membrane, while others may affect the channel's ability to conduct chloride ions[4](https://www.ncbi.nlm.nih.gov/pmc/articles/PMC10063961/)[8](https://pubmed.ncbi.nlm.nih.gov/33331662/). Studies using patient-derived intestinal organoids and CFBE cell lines have shown that some rare CFTR variants respond to currently available CFTR modulators, like elexacaftor/tezacaftor/ivacaftor (ETI), which can rescue CFTR function[3](https://www.ncbi.nlm.nih.gov/pmc/articles/PMC10572896/)[4](https://www.ncbi.nlm.nih.gov/pmc/articles/PMC10063961/). The efficacy of these modulators often depends on the specific mutation[3](https://www.ncbi.nlm.nih.gov/pmc/articles/PMC10572896/)[5](https://www.ncbi.nlm.nih.gov/pmc/articles/PMC8584557/). In the absence of specific data on c.571T>G, predicting its clinical impact and response to CFTR modulators is challenging. Further investigation is needed to determine the functional consequences of this variant, potentially through in vitro studies using cell lines or patient-derived cells, and to assess its response to CFTR modulator therapies. Such studies would help determine appropriate clinical management and personalized treatment strategies for individuals with this variant[4](https://www.ncbi.nlm.nih.gov/pmc/articles/PMC10063961/)[5](https://www.ncbi.nlm.nih.gov/pmc/articles/PMC8584557/). | 1 <https://www.semanticscholar.org/paper/db646744d07af6b7f306278b7b4d68a0bc1f4c66>  2 <https://www.ncbi.nlm.nih.gov/pmc/articles/PMC8167067/>  3 <https://www.ncbi.nlm.nih.gov/pmc/articles/PMC10572896/>  4 <https://www.ncbi.nlm.nih.gov/pmc/articles/PMC10063961/>  5 <https://www.ncbi.nlm.nih.gov/pmc/articles/PMC8584557/>  6 <https://www.ncbi.nlm.nih.gov/pmc/articles/PMC4400651/>  7 <https://pubmed.ncbi.nlm.nih.gov/31184507/>  8 [https://pubmed.ncbi.nlm.nih.gov/33331662/https://pubmed.ncbi.nlm.nih.gov/33331662/](https://pubmed.ncbi.nlm.nih.gov/33331662/) |
|  | Evaluation  **Summarization Accuracy & Completeness:**  **Hallucination Rate:**  **Readability & Usability:**  **Literature Coverage & Relevance:**  **Time Efficiency:** | |
| **4** | The genomic variant c.571T>G p.Phe191Val rs141482808 on the CFTR gene is a missense mutation that results in the substitution of phenylalanine with valine at position 191 of the CFTR protein. This variant has been reported in patients with cystic fibrosis (CF) and is associated with the disease phenotype. In one study, a patient carrying the p.Phe191Val variant in trans with the common F508del mutation exhibited normal chloride sweat concentration but had a beta-adrenergic sweat secretion rate in the CF range, suggesting minimal CFTR activity associated with this genotype [1]. Another case involved the reclassification of a patient's genotype to CF after the p.Phe191Val mutation was identified as CF-causing, with the patient initially presenting with an intermediate sweat chloride test result and later confirmed through genetic analysis [2].  The CFTR gene encodes the cystic fibrosis transmembrane conductance regulator protein, which functions as a chloride channel in epithelial cells and plays a critical role in the regulation of salt and water transport across cell membranes. Mutations in the CFTR gene can disrupt the function of this channel, leading to the accumulation of thick and sticky mucus in various organs, primarily affecting the respiratory and digestive systems. The p.Phe191Val variant is one of over 2,000 mutations described in the CFTR gene, with the F508del mutation being the most common, particularly in the Caucasian population [4].  The CFTR protein is subject to a classification system based on the primary molecular defect caused by mutations, with classes ranging from defects in synthesis to protein stability. The p.Phe191Val variant has been included in the FDA-approved list for CFTR modulator therapy with ivacaftor, a potentiator that facilitates greater ion passage through the CFTR protein [3]. This inclusion indicates that the variant has been recognized as responsive to targeted therapeutic intervention, which can enhance or restore the function of the defective CFTR channel.  In summary, the c.571T>G p.Phe191Val rs141482808 variant on the CFTR gene is a pathogenic missense mutation associated with cystic fibrosis and has been documented in clinical cases. It affects the CFTR protein's function, leading to disease manifestations, but it is also a target for modulator therapies that aim to correct the defective chloride channel activity. | [[1]](https://pubmed.ncbi.nlm.nih.gov/36751320/) Cornet M et al. (2023). "Proteomic profiling of sweat in patients with cystic fibrosis provides new insights into epidermal homoeostasis." Skin health and disease, 3(1)  [[2]](https://pubmed.ncbi.nlm.nih.gov/36969284/) Gunnett MA et al. (2023). "Outcomes of children with cystic fibrosis screen positive, inconclusive diagnosis/CFTR related metabolic syndrome." Frontiers in pediatrics, 11()  [[3]](https://pubmed.ncbi.nlm.nih.gov/39841779/) Barros LGB et al. (2025). "A new era of cystic fibrosis therapy with CFTR modulators." Jornal brasileiro de pneumologia : publicacao oficial da Sociedade Brasileira de Pneumologia e Tisilogia, 50(6)  [[4]](https://pubmed.ncbi.nlm.nih.gov/35451201/) Izquierdo ME et al. (2022). "DNA sequencing analysis of cystic fibrosis transmembrane conductance regulator gene identifies cystic fibrosis-associated variants in the Severe Asthma Research Program." Pediatric pulmonology, 57(7)  [[5]](https://pubmed.ncbi.nlm.nih.gov/35652053/) Garcia LCE et al. (2022). "Translational Research in Cystic Fibrosis: From Bench to Beside." Frontiers in pediatrics, 10()  [[6]](https://pubmed.ncbi.nlm.nih.gov/32265312/) McCravy MS et al. (2020). "Personalised medicine for non-classic cystic fibrosis resulting from rare CFTR mutations." The European respiratory journal, 56(1)  [[7]](https://pubmed.ncbi.nlm.nih.gov/34541464/) Chen KG et al. (2020). "Genomic stability at the coding regions of the multidrug transporter gene ABCB1: insights into the development of alternative drug resistance mechanisms in human leukemia cells." Cancer drug resistance (Alhambra, Calif.), 3(4) [SUPP]  [[8]](https://pubmed.ncbi.nlm.nih.gov/36670555/) Conti J et al. (2022). "Organoid Technology and Its Role for Theratyping Applications in Cystic Fibrosis." Children (Basel, Switzerland), 10(1) |
|  | Evaluation  **Summarization Accuracy & Completeness:**  **Hallucination Rate:**  **Readability & Usability:**  **Literature Coverage & Relevance:**  **Time Efficiency:** | |
| **5** | The CFTR gene encodes the cystic fibrosis transmembrane conductance regulator, a chloride channel integral to the regulation of salt and water transport across epithelial membranes. The c.571T>G variant, also known as p.Phe191Val, results from a thymine-to-guanine substitution at nucleotide position 571, leading to the replacement of phenylalanine with valine at codon 191. This amino acid change is considered conservative due to the similar properties of phenylalanine and valine. The variant has been identified in individuals diagnosed with non-classic cystic fibrosis (CF) in conjunction with other pathogenic CFTR mutations; however, the phase of these alterations was not provided. The Clinical and Functional Translation of CFTR (CFTR2) project, which evaluates genotype-phenotype correlations, has not classified this variant as CF-causing, suggesting that its pathogenicity remains uncertain. Population frequency data indicate that c.571T>G has a minor allele frequency of approximately 0.00006 in the general population. Given the limited and inconclusive evidence regarding the clinical significance of the c.571T>G variant, it is currently classified as a variant of uncertain significance. Further functional studies and comprehensive genotype-phenotype correlation analyses are necessary to elucidate its impact on CFTR function and its potential role in CF pathogenesis. | [ClinVar Miner List of variants in gene CFTR studied for Cystic fibrosis - ClinVar Miner...](https://clinvarminer.genetics.utah.edu/variants-by-gene/CFTR/condition/Cystic%20fibrosis%3B%20CFTR-related%20disorder)  [ncbi.nlm.nih.gov ClinVar Genomic variation as it relates to human health](https://www.ncbi.nlm.nih.gov/clinvar/variation/495953/)  [avalonhcs.com [PDF] M2017 – Genetic Testing for Cystic Fibrosis Prior Policy Name and N](https://www.avalonhcs.com/wp-content/uploads/sentarahealth/M2017%20Genetic%20Testing%20for%20Cystic%20Fibrosis.pdf)  [1 dicembre 2024 — The Clinical and Functional Translation of CFTR (CFTR2) project continually evaluates … avalonhcs.com [PDF] M2017 – Genetic Testing for Cystic Fibrosis Prior Policy Name and N](https://www.avalonhcs.com/wp-content/uploads/sentarahealth/M2017%20Genetic%20Testing%20for%20Cystic%20Fibrosis.pdf)  [1 dicembre 2024 — The Clinical and Functional Translation of CFTR (CFTR2) project continually evaluates ...](https://www.avalonhcs.com/wp-content/uploads/sentarahealth/M2017%20Genetic%20Testing%20for%20Cystic%20Fibrosis.pdf)  [gimjournal.org Updated recommendations for CFTR carrier screening: A position ...](https://www.gimjournal.org/article/S1098-3600%2823%2900880-8/fulltext)  [13 giugno 2023 — Pathogenic variants in the CFTR gene are causative of cystic fibrosis (CF) ...](https://www.gimjournal.org/article/S1098-3600%2823%2900880-8/fulltext)  [ncbi.nlm.nih.gov NM_000492.3(CFTR):c.571T>G (p.Phe191Val) AND Cystic fibrosis 25 novembre 2018 — 571T>G (p.Phe191Val). ...](https://www.ncbi.nlm.nih.gov/clinvar/35503424/)  [ncbi.nlm.nih.gov ClinVar Genomic variation as it relates to human health - NCBI](https://www.ncbi.nlm.nih.gov/clinvar/variation/495953/)  [ClinVar Miner List of variants in gene CFTR reported as likely pathogenic for Cystic ...](https://clinvarminer.genetics.utah.edu/variants-by-gene/CFTR/condition/Cystic%20fibrosis/likely%20pathogenic)  [List of variants in gene CFTR reported as likely pathogenic for Cystic fibrosis ; NM_000492.4(CFTR):c.2797A>G (p.Arg933Gly), ...](https://clinvarminer.genetics.utah.edu/variants-by-gene/CFTR/condition/Cystic%20fibrosis/likely%20pathogenic)  [ClinVar Miner List of variants in gene CFTR studied for Cystic fibrosis -](https://clinvarminer.genetics.utah.edu/variants-by-gene/CFTR/condition/Cystic%20fibrosis%3B%20CFTR-related%20disorder)  [ClinVar Miner List of variants in gene CFTR reported as uncertain significance for ...](https://clinvarminer.genetics.utah.edu/variants-by-gene/CFTR/condition/Cystic%20fibrosis/uncertain%20significance)  [wadsworth.org [PDF] NYS Cystic Fibrosis Newborn Screening: Custom CFTR Variant ...](https://www.wadsworth.org/sites/default/files/CF%20Archer%202nd%20Tier%20Panel%20Content_v1.1.0_Final%20for%20website.pdf)  [F191V \| p.Phe191Val (c.571T>G)§. D192G \| p.Asp192Gly (c.575A>G)§. Variant ... .](https://www.wadsworth.org/sites/default/files/CF%20Archer%202nd%20Tier%20Panel%20Content_v1.1.0_Final%20for%20website.pdf)  [databases.lovd.nl All variants in the CFTR gene - Whole genome datasets](https://databases.lovd.nl/whole_genome/variants/CFTR) [g.117174411T>G, Copied from the {URL:Exome ...](https://databases.lovd.nl/whole_genome/variants/CFTR)  [journalpulmonology.org Spectrum of CFTR gene sequence variants in a northern Portugal ...](https://www.journalpulmonology.org/en-spectrum-cftr-gene-sequence-variants-articulo-S2531043718300345)  [A total of 30 different CFTR sequence variants, already reported as cystic fibrosis (CF) or CFTR related disorders variants, were detected.](https://www.journalpulmonology.org/en-spectrum-cftr-gene-sequence-variants-articulo-S2531043718300345)  [scientiasalut.gencat.cat [PDF] Standards of care for CFTR variant-specific therapy (including … CFTR gene variants should ...](https://scientiasalut.gencat.cat/bitstream/handle/11351/9241/standards_care_cftr_variant_specific_therapy_including_modulators_people_cystic_fibrosis_2023_material_suplementari.pdf?isAllowed=y&sequence=2)  [nature.com Clinical practice and genetic counseling for cystic fibrosis and CFTR …](https://www.nature.com/articles/gim2008133) |
|  | Evaluation  **Summarization Accuracy & Completeness:**  **Hallucination Rate:**  **Readability & Usability:**  **Literature Coverage & Relevance:**  **Time Efficiency:** | |
