## Supplementary material for "Benchmarking generative AI tools for literature retrieval and summarization in genomic variant interpretation": FAS_p.Arg250Gln.docx

| **Tool** | **Summary** | **References** |
| --- | --- | --- |
| **1** | The genomic variant c.749G>A, resulting in the amino acid substitution p.Arg250Gln in the FAS gene, has been classified as pathogenic with deleterious effects on protein function [2]. This variant is located within the intracellular death domain of the FAS protein, which is crucial for the induction of apoptosis [2]. The FAS gene encodes a cell surface receptor that plays a vital role in the regulation of programmed cell death and is involved in the pathway that leads to apoptosis. The proper functioning of FAS is essential for the maintenance of immune system homeostasis and the prevention of autoimmunity.  The p.Arg250Gln variant has been associated with Autoimmune Lymphoproliferative Syndrome (ALPS), a disorder characterized by abnormal lymphocyte survival, lymphadenopathy, splenomegaly, and an increased risk of lymphoma [2]. Patients with ALPS typically present with defects in lymphocyte apoptosis, leading to the accumulation of autoreactive cells and the clinical manifestations of the disease.  The pathogenicity of the c.749G>A variant is supported by its classification in ClinVar, where it has been submitted as pathogenic five times. This variant has been observed in individuals with clinical features consistent with ALPS, including hepatosplenomegaly, enlarged lymph nodes, and hematological abnormalities such as thrombocytopenia [2]. Furthermore, the variant has been implicated in cases with Hodgkin lymphoma, which is in remission, highlighting the potential impact of FAS dysfunction on lymphoproliferative disorders [2].  In summary, the c.749G>A p.Arg250Gln variant in the FAS gene is a missense mutation that has been associated with pathogenic effects and is implicated in the development of ALPS. The variant's location within the death domain of FAS underscores its significance in the apoptotic function of the protein and its role in maintaining immune system balance. | [[1]](https://pubmed.ncbi.nlm.nih.gov/37392849/) Minafra AR et al. (2023). "Synthetic receptor platform to identify loss-of-function single nucleotide variants and designed mutants in the death receptor Fas/CD95." The Journal of biological chemistry, 299(8)  [[2]](https://pubmed.ncbi.nlm.nih.gov/39060684/) Xu X et al. (2024). "Genetic Testing in Patients with Autoimmune Lymphoproliferative Syndrome: Experience of 802 Patients at Cincinnati Children's Hospital Medical Center." Journal of clinical immunology, 44(7)  [[3]](https://pubmed.ncbi.nlm.nih.gov/30120239/) Braun S et al. (2018). "Decoding a cancer-relevant splicing decision in the RON proto-oncogene using high-throughput mutagenesis." Nature communications, 9(1)  [[4]](https://pubmed.ncbi.nlm.nih.gov/21490157/) Kuehn HS et al. (2011). "FAS haploinsufficiency is a common disease mechanism in the human autoimmune lymphoproliferative syndrome." Journal of immunology (Baltimore, Md. : 1950), 186(10)  [[5]](https://pubmed.ncbi.nlm.nih.gov/19118384/) Scott FL et al. (2009). "The Fas-FADD death domain complex structure unravels signalling by receptor clustering." Nature, 457(7232) [SUPP]  [[6]](https://pubmed.ncbi.nlm.nih.gov/27915290/) Kumar S et al. (2016). "Localized structural frustration for evaluating the impact of sequence variants." Nucleic acids research, 44(21) [SUPP]  [[7]](https://pubmed.ncbi.nlm.nih.gov/26147798/) Ferrarini A et al. (2015). "The Use of Non-Variant Sites to Improve the Clinical Assessment of Whole-Genome Sequence Data." PloS one, 10(7) [SUPP]  [[8]](https://pubmed.ncbi.nlm.nih.gov/26659599/) Holehouse AS et al. (2015). "Reproducible Analysis of Post-Translational Modifications in Proteomes--Application to Human Mutations." PloS one, 10(12) [SUPP] |
|  | Evaluation  **Summarization Accuracy & Completeness:**  **Hallucination Rate:**  **Readability & Usability:**  **Literature Coverage & Relevance:**  **Time Efficiency:** | |
| **2** | The FAS gene encodes a member of the tumor necrosis factor receptor (TNFR) superfamily and plays a critical role in the regulation of programmed cell death (apoptosis). This process is vital for immune system homeostasis and the elimination of autoreactive lymphocytes. Mutations in FAS can disrupt apoptotic signaling, leading to autoimmune lymphoproliferative syndrome (ALPS), characterized by chronic lymphadenopathy, splenomegaly, and autoimmune cytopenias. The p.Arg250Gln (c.749G>A) missense variant results in an amino acid substitution at a highly conserved position within the intracellular death domain of the protein, which is essential for downstream recruitment of FADD and subsequent caspase activation. Although this residue lies in a functionally critical region, there are currently no peer-reviewed studies directly evaluating the pathogenicity or functional consequences of the p.Arg250Gln substitution. It is therefore not possible to classify the variant definitively; it remains a variant of uncertain significance (VUS). Computational predictions may suggest a possible deleterious effect due to its location and conservation, but without empirical evidence, clinical interpretation should remain cautious. Further functional assays, segregation studies, and clinical correlation are needed to determine its impact on FAS-mediated apoptosis and its potential involvement in ALPS or related disorders. | No references available |
|  | Evaluation  **Summarization Accuracy & Completeness:**  **Hallucination Rate:**  **Readability & Usability:**  **Literature Coverage & Relevance:**  **Time Efficiency:** | |
| **3** | The FAS gene encodes a transmembrane receptor crucial for initiating apoptosis through the extrinsic pathway, primarily by binding to its ligand, FasL, leading to the formation of the death-inducing signaling complex (DISC) and subsequent activation of caspases. This mechanism is vital for maintaining immune homeostasis by eliminating autoreactive or excess lymphocytes. The p.Arg250Gln (c.749G>A) variant results in a missense substitution of arginine to glutamine at position 250 within the highly conserved death domain of the FAS protein. This variant has been identified in individuals with autoimmune lymphoproliferative syndrome (ALPS), a disorder characterized by defective lymphocyte apoptosis, leading to lymphadenopathy, hepatosplenomegaly, and autoimmune manifestations . Functional studies have demonstrated that this substitution impairs FAS-mediated apoptotic signaling, contributing to the ALPS phenotype . The variant is absent from large population databases, suggesting it is not a common polymorphism . Given its location in a critical functional domain and its association with disease, the p.Arg250Gln variant is classified as pathogenic. Further studies are warranted to elucidate the full spectrum of clinical manifestations associated with this variant and to inform management strategies for affected individuals. | <https://www.ncbi.nlm.nih.gov/gene/355>  <https://www.ncbi.nlm.nih.gov/clinvar/RCV001382235/>  <https://pmc.ncbi.nlm.nih.gov/articles/PMC11282156/>  <https://www.sciencedirect.com/science/article/pii/S0006497120616172>  <https://www.nature.com/articles/4401179>  <https://pmc.ncbi.nlm.nih.gov/articles/PMC2956119/>  <https://pmc.ncbi.nlm.nih.gov/articles/PMC5037862/>  <https://pubmed.ncbi.nlm.nih.gov/19239902/>  <https://medlineplus.gov/genetics/gene/fas/>  <https://en.wikipedia.org/wiki/Fas_receptor>  <https://www.sciencedirect.com/science/article/pii/S1525157815002615>  <https://www.sciencedirect.com/science/article/pii/S0065277608606720>  <https://www.genecards.org/cgi-bin/carddisp.pl?gene=FAS>  <https://varsome.com/variant/hg38/rs121913080>  <https://rgd.mcw.edu/rgdweb/report/gene/main.html?id=1346266>  <https://v1.marrvel.org/search/gene/FAS>  <https://web.expasy.org/variant_pages/VAR_013427.html> |
|  | Evaluation  **Summarization Accuracy & Completeness:**  **Hallucination Rate:**  **Readability & Usability:**  **Literature Coverage & Relevance:**  **Time Efficiency:** | |
| **4** | The FAS gene, also known as Fas Cell Surface Death Receptor or TNFRSF6, is a critical component of the tumor necrosis factor receptor superfamily. It encodes a protein that plays a central role in programmed cell death, or apoptosis, which is essential for immune system regulation and the prevention of malignancies. The FAS protein contains a death domain that, upon activation, forms a death-inducing signaling complex (DISC) with FADD, caspase 8, and caspase 10. This complex initiates a caspase cascade that leads to apoptosis, a process vital for maintaining homeostasis by eliminating unnecessary or damaged cells. The FAS gene is located on chromosome 10q23.31 and is involved in various pathways, including the dimerization of procaspase-8 and the TNFR1 pathway. Mutations in FAS have been associated with autoimmune lymphoproliferative syndrome (ALPS) and other immune system disorders, as well as an increased risk of developing certain cancers, including lung, breast, and esophageal cancers. The p.Arg250Gln variant of the FAS gene is a missense mutation that results in the substitution of glutamine for arginine at amino acid position 250. This variant has been reported in the context of genetic studies, but its specific impact on FAS function and clinical significance is not well-documented in the available literature. The frequency and classification of this variant are not explicitly stated in peer-reviewed publications, and further investigation is needed to determine its role in disease pathogenesis. The lack of direct references to this variant in academic research suggests that more studies are required to understand its implications fully. Therefore, while the general function of the FAS gene is well-established, the specific effects of the p.Arg250Gln variant remain uncertain and warrant further research. |  |
|  | Evaluation  **Summarization Accuracy & Completeness:**  **Hallucination Rate:**  **Readability & Usability:**  **Literature Coverage & Relevance:**  **Time Efficiency:** | |
| **5** | The FAS gene encodes a transmembrane receptor critical for mediating apoptosis, particularly in immune regulation and lymphocyte homeostasis. It plays a pivotal role in the extrinsic pathway of programmed cell death, which is essential for eliminating autoreactive lymphocytes and maintaining immune tolerance. Variants in FAS can influence susceptibility to autoimmune conditions and immune dysregulation syndromes. The p.Arg250Gln (R250Q) variant in FAS is a missense mutation resulting from a nucleotide change that substitutes arginine with glutamine at amino acid position 250. This variant has been identified in the context of immune dysregulation, with some evidence suggesting it may impair FAS-mediated apoptosis, although direct functional studies specific to R250Q are limited. The classification of this variant remains uncertain, as it has not been definitively linked to a specific disease phenotype in large cohorts, and its frequency in the general population appears low, indicating it may be a rare or private mutation. Literature on this specific variant is sparse, and no peer-reviewed studies directly assess its pathogenicity or functional impact. However, mutations affecting the FAS receptor, especially within the death domain or ligand-binding regions, are known to impair apoptosis and contribute to conditions such as autoimmune lymphoproliferative syndrome (ALPS) and other immune dysregulation disorders. Given the importance of the FAS pathway in immune homeostasis, the R250Q variant warrants further functional analysis to clarify its role in disease. In summary, while the FAS gene is well-characterized as a key regulator of apoptosis with established links to immune disorders, specific data on the p.Arg250Gln variant are limited, and its clinical significance remains to be conclusively determined. Further research, including functional assays and population studies, is necessary to establish its pathogenicity and potential contribution to immune dysregulation syndromes. | [1] <https://www.ncbi.nlm.nih.gov/pmc/articles/PMC9388157/>  [2] <https://www.ncbi.nlm.nih.gov/pmc/articles/PMC5807897/>  [3] <https://www.ncbi.nlm.nih.gov/pmc/articles/PMC8810734/>  [4] <https://pubmed.ncbi.nlm.nih.gov/27391055/>  [5] <https://pubmed.ncbi.nlm.nih.gov/31309545/>  [6] <https://pubmed.ncbi.nlm.nih.gov/23430061/>  [7] <https://www.ncbi.nlm.nih.gov/pmc/articles/PMC2212386/>  [8] <https://pubmed.ncbi.nlm.nih.gov/9108407/> |
|  | Evaluation  **Summarization Accuracy & Completeness:**  **Hallucination Rate:**  **Readability & Usability:**  **Literature Coverage & Relevance:**  **Time Efficiency:** | |
