## Supplementary material for "Benchmarking generative AI tools for literature retrieval and summarization in genomic variant interpretation": G6PD p.S188F.docx

| **Tool** | **Summary** | **References** |
| --- | --- | --- |
| **1** | The glucose-6-phosphate dehydrogenase (G6PD) gene encodes an enzyme crucial for the pentose phosphate pathway, primarily producing NADPH, which is essential for maintaining cellular redox balance and supporting various biosynthetic processes[5][6]. The p.S188F variant, also known as the Mediterranean mutation, is a common cause of G6PD deficiency, leading to reduced enzyme activity[7]. This mutation results in a serine-to-phenylalanine substitution at position 188, significantly impairing the enzyme's function and increasing susceptibility to oxidative stress[7]. G6PD deficiency is an X-linked recessive disorder, affecting more than 400 million people worldwide, with the Mediterranean mutation being particularly prevalent in certain regions[6][7]. In the context of vascular diseases, G6PD activity has been linked to the regulation of smooth muscle cell differentiation and the pathogenesis of metabolic syndrome[1]. Additionally, G6PD plays roles in other cellular processes, such as thermogenesis in brown adipocytes and immune function in cytotoxic T lymphocytes[2][4]. While the p.S188F variant is well-studied in the context of G6PD deficiency, its specific impact on these broader cellular functions requires further investigation. References:[1][2][4][6][7]. | [1] <https://pubmed.ncbi.nlm.nih.gov/33416454/>  [2] <https://pubmed.ncbi.nlm.nih.gov/34521642/>  [3] <https://pubmed.ncbi.nlm.nih.gov/37599085/>  [4] <https://www.ncbi.nlm.nih.gov/pmc/articles/PMC8753452/>  [5] <https://pubmed.ncbi.nlm.nih.gov/10709108/>  [6] <https://pubmed.ncbi.nlm.nih.gov/17516514/>  [7] <https://www.semanticscholar.org/paper/8869d3b531cf3189b36cbafd52349313e06ac9db>  [8] <https://www.semanticscholar.org/paper/911368da789ea9379586323b16684b626cb6cf08> |
|  | Evaluation  **Summarization Accuracy & Completeness:**  **Hallucination Rate:**  **Readability & Usability:**  **Literature Coverage & Relevance:**  **Time Efficiency:** | |
| **2** | The G6PD gene encodes the enzyme glucose-6-phosphate dehydrogenase, which is crucial in the pentose phosphate pathway, particularly for producing NADPH and maintaining the cellular redox state. This enzyme is essential in red blood cells, protecting them from oxidative damage by generating reduced glutathione. The p.S188F variant, also known as G6PD Mediterranean, results from a c.563C>T nucleotide substitution, leading to a serine-to-phenylalanine change at codon 188. This variant is classified as pathogenic, with computational analyses predicting it to be deleterious. The p.S188F variant significantly reduces G6PD enzyme activity, with residual activity ranging from 0-7% compared to the wild type, and it decreases thermostability and catalytic activity. Individuals with this variant are at risk of hemolytic anemia triggered by certain infections, medications, or specific foods. The variant is relatively common in the Mediterranean region and has been associated with a higher risk of stroke in Saudi sickle cell patients. Interestingly, some studies suggest that the p.S188F variant may confer vascular protection, potentially reducing the incidence of vascular diseases. However, the data on this protective effect are not conclusive, and further research is needed. The frequency of the p.S188F variant varies across populations, with a reported frequency of 0.017450 in the South Asian population. Despite the potential protective effects against malaria and vascular diseases, the primary clinical concern remains the risk of hemolytic anemia. Therefore, individuals with this variant should be aware of potential triggers and manage their condition accordingly. Further investigation is recommended to fully understand the variant's impact on vascular health and other potential benefits or risks. | [1] Hellani A, Al-Akoum S, Abu-Amero KK. Genet Test Mol Biomarkers. 2009 Aug;13(4):449-52. doi: 10.1089/gtmb.2009.0011.  [2] Kitagawa A, Kizub I, Jacob C, Michael K, D’Alessandro A, Reisz JA, Grzybowski M, Geurts AM, Rocic P, Gupte R, et al. Hypertension. 2020 Aug;76(2):314-315. doi: 10.1161/HYPERTENSIONAHA.120.14897.  [3] Vulliamy TJ, et al. 1988.  [4] ClinVar archives and aggregates information about relationships among variation and human health. |
|  | Evaluation  **Summarization Accuracy & Completeness:**  **Hallucination Rate:**  **Readability & Usability:**  **Literature Coverage & Relevance:**  **Time Efficiency:** | |
| **3** | The search for peer-reviewed literature specifically referencing the G6PD p.S188F variant encountered a technical error, preventing access to up-to-date data. However, based on current scientific understanding of the G6PD gene, it encodes the enzyme glucose-6-phosphate dehydrogenase, which is essential for the pentose phosphate pathway, particularly in red blood cells where it provides protection against oxidative damage. Deficiency in G6PD can lead to hemolytic anemia triggered by infections, certain drugs, or foods such as fava beans. The p.S188F variant results from a missense mutation substituting serine with phenylalanine at codon 188. While this specific variant does not appear to be extensively reported in the peer-reviewed literature, its location within a highly conserved region of the gene suggests potential functional impact. Mutations affecting similar residues have been associated with reduced enzymatic activity and clinical manifestations of G6PD deficiency. In the absence of direct functional studies or clinical reports concerning p.S188F, the variant should be interpreted with caution, and its pathogenicity remains uncertain. Further biochemical assays or population studies would be needed to determine its significance. No peer-reviewed studies specific to G6PD p.S188F were identified at this time. | No references available |
|  | Evaluation  **Summarization Accuracy & Completeness:**  **Hallucination Rate:**  **Readability & Usability:**  **Literature Coverage & Relevance:**  **Time Efficiency:** | |
| **4** | Glucose-6-phosphate dehydrogenase (G6PD) is a crucial enzyme in the pentose phosphate pathway, responsible for maintaining cellular redox balance by generating nicotinamide adenine dinucleotide phosphate (NADPH). NADPH is essential for protecting red blood cells from oxidative damage. The p.S188F variant, also known as G6PD Mediterranean, results from a c.563C>T substitution in the G6PD gene, leading to the replacement of serine with phenylalanine at position 188. This missense mutation is classified as pathogenic and is associated with G6PD deficiency, a genetic disorder that can cause acute hemolytic anemia in response to oxidative stressors. The p.S188F variant is prevalent among individuals of Mediterranean and Middle Eastern descent, with studies reporting its presence in populations from Greece, southern Italy, Spain, and Saudi Arabia. Functional analyses have demonstrated that this variant leads to significantly reduced enzyme activity, with residual activity ranging from 0% to 7% of normal levels. Additionally, the p.S188F mutation impairs the enzyme's thermostability and catalytic efficiency. In individuals with sickle cell disease, the presence of the G6PD Mediterranean variant has been associated with an increased risk of stroke. Experimental models, such as CRISPR-Cas9-generated rats carrying the S188F mutation, have provided insights into the variant's impact on vascular function and metabolic pathways. These studies suggest that the G6PD S188F variant may confer some protective effects against high-fat diet-induced vascular dysfunction and hypertension. However, the clinical significance of these findings remains to be fully elucidated. Given the established association between the p.S188F variant and G6PD deficiency, individuals carrying this mutation should be advised to avoid known oxidative triggers, such as certain medications and foods, to prevent hemolytic episodes. Further research is warranted to explore the broader clinical implications of this variant and to develop targeted interventions for affected individuals. | Vulliamy TJ, D'Urso M, Battistuzzi G, et al. Diverse point mutations in the human glucose-6-phosphate dehydrogenase gene cause enzyme deficiency and mild or severe hemolytic anemia. *Proc Natl Acad Sci U S A*. 1988;85(14):5171-5175.[NCBI](https://www.ncbi.nlm.nih.gov/clinvar/variation/VCV000100057.5?utm_source=chatgpt.com)  Hellani A, Al-Akoum S, Abu-Amero KK. G6PD Mediterranean S188F codon mutation is common among Saudi sickle cell patients and increases the risk of stroke. *Genet Test Mol Biomarkers*. 2009;13(4):449-452.[AHA Journals+3PubMed+3Liebertpub.com+3](https://pubmed.ncbi.nlm.nih.gov/19594365/?utm_source=chatgpt.com)  Li Y, Pagano PJ. Does the Mediterranean G6PD S188F polymorphism confer vascular protection?: A novel rat model offering CRISPR insight into high fat-induced vascular dysfunction and hypertension. *Hypertension*. 2020;76(2):314-315.[AHA Journals+7PubMed+7AHA Journals+7](https://pubmed.ncbi.nlm.nih.gov/32639882/?utm_source=chatgpt.com)  ClinVar. Variation Report for NM_000402.4(G6PD):c.563C>T (p.Ser188Phe). Available at:<https://www.ncbi.nlm.nih.gov/clinvar/variation/100057/>. Accessed April 8, 2025.[Wikipedia+3NCBI+3NCBI+3](https://www.ncbi.nlm.nih.gov/clinvar/variation/100057/?utm_source=chatgpt.com)  Others:  <https://pubmed.ncbi.nlm.nih.gov/39514761/>  <https://ashpublications.org/bloodadvances/article/9/2/321/525937/Increased-exercise-tolerance-in-humanized-G6PD>  <https://pubmed.ncbi.nlm.nih.gov/38876306/>  <https://pmc.ncbi.nlm.nih.gov/articles/PMC10624141/>  <https://pubmed.ncbi.nlm.nih.gov/36945640/>  <https://pubmed.ncbi.nlm.nih.gov/36372233/>  <https://pubmed.ncbi.nlm.nih.gov/35623546/>  <https://onlinelibrary.wiley.com/doi/10.1155/2021/5529256>  <https://pubmed.ncbi.nlm.nih.gov/33416454/>  <https://ashpublications.org/blood/article/136/11/1225/461549/Glucose-6-phosphate-dehydrogenase-deficiency>  <https://www.ahajournals.org/doi/10.1161/HYPERTENSIONAHA.120.14897>  <https://pubmed.ncbi.nlm.nih.gov/32507041/>  <https://medical.23andme.com/wp-content/uploads/2020/06/zero_variants_male.pdf>  <https://www.nature.com/articles/s41467-018-06447-z>  <https://www.nature.com/articles/jhg200399>  <https://www.nature.com/articles/srep37284>  <https://www.ncbi.nlm.nih.gov/clinvar/variation/VCV000100057.5/>  <https://www.sciencedirect.com/science/article/pii/S0021925824019616>  <https://www.sciencedirect.com/science/article/pii/S1043661821004886>  <https://www.coriell.org/0/sections/Search/Sample_Detail.aspx?Product=CC&Ref=GM01870> |
|  | Evaluation  **Summarization Accuracy & Completeness:**  **Hallucination Rate:**  **Readability & Usability:**  **Literature Coverage & Relevance:**  **Time Efficiency:** | |
| **5** | The genomic variant c.563C>T p.S188F rs5030868 in the G6PD gene is a missense mutation that results in the substitution of serine with phenylalanine at the 188th amino acid position, commonly referred to as the "Mediterranean" variant [1]. This variant has been associated with G6PD deficiency, a condition characterized by reduced activity of the glucose-6-phosphate dehydrogenase enzyme, which plays a critical role in the pentose phosphate pathway, particularly in red blood cells. The enzyme's primary function is to protect the cells from oxidative damage by maintaining levels of the reduced form of nicotinamide adenine dinucleotide phosphate (NADPH), which in turn preserves the integrity of red blood cell membranes by keeping glutathione in its reduced form.  The c.563C>T variant has been reported to be associated with Class II G6PD activity, indicating a severe deficiency with enzyme activity levels typically around 2-10% of normal [2]. This variant is one of the most common disease-causing mutations in the G6PD gene, found on 16 alleles in a study cohort, and is often associated with a residual enzyme activity of less than 10% of normal [2]. The presence of this variant has been linked to an increased risk of moderate hyperbilirubinemia in neonates, which can lead to conditions such as severe neonatal jaundice [4]. Additionally, the residual activity of the enzyme in hemizygote males with the c.563C>T variant has been shown to be significantly lower than in non-c.563C>T G6PD-deficient males and controls, further correlating the mutation with reduced enzyme function and clinical manifestations [5].  The G6PD gene, located on the X chromosome, is vital for the proper functioning of red blood cells, and mutations leading to its deficiency can result in hemolytic anemia under certain stress conditions, such as infection or exposure to certain medications or foods. The c.563C>T variant's pathogenicity is supported by multiple submissions in ClinVar, with a majority indicating a pathogenic status. This evidence underscores the clinical significance of the variant in the context of G6PD deficiency and its related hematological disorders. | [[1]](https://pubmed.ncbi.nlm.nih.gov/24586352/) Jamornthanyawat N et al. (2014). "A population survey of the glucose-6-phosphate dehydrogenase (G6PD) 563C>T (Mediterranean) mutation in Afghanistan." PloS one, 9(2)  [[2]](https://pubmed.ncbi.nlm.nih.gov/33072997/) Ohlsson A et al. (2019). "Incidence of Glucose-6-Phosphate Dehydrogenase Deficiency among Swedish Newborn Infants." International journal of neonatal screening, 5(4)  [[3]](https://pubmed.ncbi.nlm.nih.gov/35306802/) Orman B et al. (2023). "Hemolytic Anemia due to Glucose 6 Phosphate Dehydrogenase Deficiency Triggered by Type 1 Diabetes Mellitus." Journal of clinical research in pediatric endocrinology, 15(4)  [[4]](https://pubmed.ncbi.nlm.nih.gov/22906047/) Moiz B et al. (2012). "Neonatal hyperbilirubinemia in infants with G6PD c.563C > T Variant." BMC pediatrics, 12()  [[5]](https://pubmed.ncbi.nlm.nih.gov/24460025/) Molou E et al. (2014). "Glucose-6-Phosphate Dehydrogenase (G6PD) deficiency in Greek newborns: the Mediterranean C563T mutation screening." Scandinavian journal of clinical and laboratory investigation, 74(3)  [[6]](https://pubmed.ncbi.nlm.nih.gov/38069050/) Signoretti C et al. (2023). "G6PD Orchestrates Genome-Wide DNA Methylation and Gene Expression in the Vascular Wall." International journal of molecular sciences, 24(23)  [[7]](https://pubmed.ncbi.nlm.nih.gov/38876306/) Matsumura S et al. (2024). "Loss-of-function G6PD variant moderated high-fat diet-induced obesity, adipocyte hypertrophy, and fatty liver in male rats." The Journal of biological chemistry, 300(7)  [[8]](https://pubmed.ncbi.nlm.nih.gov/39514761/) Cendali FI et al. (2025). "Increased exercise tolerance in humanized G6PD-deficient mice." Blood advances, 9(2)  [[9]](https://pubmed.ncbi.nlm.nih.gov/39730495/) Errigo A et al. (2024). "Lack of association between common polymorphisms associated with successful aging and longevity in the population of Sardinian Blue Zone." Scientific reports, 14(1)  [[10]](https://pubmed.ncbi.nlm.nih.gov/19594365/) Hellani A et al. (2009). "G6PD Mediterranean S188F codon mutation is common among Saudi sickle cell patients and increases the risk of stroke." Genetic testing and molecular biomarkers, 13(4)  [[11]](https://pubmed.ncbi.nlm.nih.gov/21205543/) Al-Alimi AA et al. (2010). "Mediterranean glucose-6-phosphate dehydrogenase (G6PD(C563T)) mutation among Jordanian females with acute hemolytic crisis." Journal of the College of Physicians and Surgeons--Pakistan : JCPSP, 20(12)  [[12]](https://pubmed.ncbi.nlm.nih.gov/22364808/) Sirdah M et al. (2012). "Hemolysis and Mediterranean G6PD mutation (c.563 C>T) and c.1311 C>T polymorphism among Palestinians at Gaza Strip." Blood cells, molecules & diseases, 48(4)  [[13]](https://pubmed.ncbi.nlm.nih.gov/32507041/) Kitagawa A et al. (2020). "CRISPR-Mediated Single Nucleotide Polymorphism Modeling in Rats Reveals Insight Into Reduced Cardiovascular Risk Associated With Mediterranean G6PD Variant." Hypertension (Dallas, Tex. : 1979), 76(2)  [[14]](https://pubmed.ncbi.nlm.nih.gov/36945640/) Signoretti C et al. (2023). "Studies in CRISPR-generated Mediterranean G6PD variant rats reveal G6PD orchestrates genome-wide DNA methylation and gene expression in vascular wall." bioRxiv : the preprint server for biology, ()  [[15]](https://pubmed.ncbi.nlm.nih.gov/37508669/) Riskin A et al. (2023). "The Genetics of Glucose-6-Phosphate-Dehydrogenase (G6PD) and Uridine Diphosphate Glucuronosyl Transferase 1A1 (UGT1A1) Promoter Gene Polymorphism in Relation to Quantitative Biochemical G6PD Activity Measurement and Neonatal Hyperbilirubinemia." Children (Basel, Switzerland), 10(7) |
|  | Evaluation  **Summarization Accuracy & Completeness:**  **Hallucination Rate:**  **Readability & Usability:**  **Literature Coverage & Relevance:**  **Time Efficiency**: | |
