## Supplementary material for "Benchmarking generative AI tools for literature retrieval and summarization in genomic variant interpretation": HBB c.20A_T.docx

| **Tool** | **Summary** | **References** |
| --- | --- | --- |
| **1** | The HBB gene encodes the beta-globin subunit of hemoglobin, the primary oxygen-carrying protein in red blood cells. Hemoglobin is a tetramer composed of two alpha-globin and two beta-globin chains, each containing a heme group that binds oxygen. The HBB gene is located on chromosome 11p15.4 and is crucial for oxygen transport from the lungs to tissues throughout the body. Variants in the HBB gene can lead to hemoglobinopathies, such as sickle cell disease and beta-thalassemia, which affect the structure or production of hemoglobin.  The c.20A>T variant in the HBB gene results in a missense mutation, p.Glu7Val, also known as HbS. This variant is the most prevalent genotype associated with sickle cell disease, an autosomal recessive disorder characterized by intravascular sickling, hemolysis, anemia, and leukocytosis. The variant causes a non-conservative amino acid change in the Globin domain of the beta-globin protein, leading to decreased solubility of deoxygenated hemoglobin S and subsequent polymerization, which forms an extensive network of fibers in red blood cells. This polymerization distorts the shape of red blood cells, leading to the characteristic sickle shape and impaired blood flow. The c.20A>T variant has been reported in multiple individuals affected with sickle cell disease, with experimental evidence demonstrating its impact on protein function. The variant allele frequency is approximately 0.0035 in control populations, with a minor allele frequency of 0.00348% in gnomAD exomes and genomes. In silico tools predict a damaging effect on the gene or gene product, and the variant is classified as pathogenic according to ACMG/AMP guidelines. The severity of the impact on the protein is considered medium. The variant has been observed in both homozygous and heterozygous states, with the homozygous state (HbSS) being the most common form of sickle cell disease. The clinical presentation of sickle cell disease can vary, with factors such as haplotype influencing disease severity. For instance, patients with the Indian/Arabian haplotype may experience a milder course of the disease compared to those with the Bantu haplotype. | [1] [ClinVar](https://www.ncbi.nlm.nih.gov/clinvar/RCV001192494/). NM_000518.5(HBB):c.20A>T (p.Glu7Val) AND Sickle cell disease and related diseases - ClinVar - NCBI.  [2] [ClinVar](https://www.ncbi.nlm.nih.gov/clinvar/RCV000016573/). NM_000518.5(HBB):c.20A>T (p.Glu7Val) AND HEMOGLOBIN S - ClinVar - NCBI.  [3] [ClinVar](https://www.ncbi.nlm.nih.gov/clinvar/RCV000016574/). NM_000518.5(HBB):c.20A>T (p.Glu7Val) AND Hb SS disease - ClinVar - NCBI. |
|  | Evaluation  **Summarization Accuracy & Completeness:**  **Hallucination Rate:**  **Readability & Usability:**  **Literature Coverage & Relevance:**  **Time Efficiency:** | |
| **2** | The genomic variant c.20A>T p.Glu7Val, also known as rs334, is located on the HBB gene, which encodes the beta-globin subunit of hemoglobin, the protein responsible for oxygen transport in the blood. The c.20A>T variant results in the substitution of glutamic acid with valine at the seventh amino acid position of the beta-globin chain. This missense mutation is the most common cause of sickle cell disease (SCD) when present in homozygous form or in compound heterozygous form with another pathogenic HBB variant [4].  The rs334 variant has been associated with increased susceptibility to pneumonia and COVID-19, particularly in individuals of African ancestry [1]. Genome-wide association studies (GWAS) have identified rs334 as a significant variant in the context of pneumonia, with the A allele conferring an increased risk [3]. This association is consistent with the protective effect of the heterozygous state against malaria, which has led to the positive selection of haplotypes containing the rs334 A allele in populations where malaria is endemic [3].  Further analysis has shown that rs334 is in linkage disequilibrium with other markers in the region, some of which are annotated as promoters or enhancers, potentially affecting the expression of HBB and other related genes such as HBD, HBE1, and HBG2 [2]. The variant rs334 itself has been annotated as bound by proteins, suggesting a regulatory role that could impact gene expression [2].  Conditional logistic regression analysis has demonstrated that no other chromosome 11 variant remained significant after conditioning on rs334, indicating the strong association of this variant with disease phenotypes [3]. Additionally, the variant rs33930165, which is a nonsynonymous SNP in the same codon as rs334 and results in hemoglobin C, also showed a significant association with pneumonia when considered alongside rs334 [3].  The evolutionary history of rs334 is complex, with high-coverage sequencing confirming a single African origin of the sickle-cell gene variant. The distribution of rs334 overlaps with other disease-associated variants, necessitating concurrent investigation of their evolutionary genetics [4]. The variant is part of a rich source of evolutionary information on HBB-betaS and its implications for human migration within and out of Africa [4].  In the clinical context, the variant rs334 has been reported in ClinVar with a majority of submissions classifying it as pathogenic. This is consistent with its known role in causing SCD when present in specific genotypic combinations. The variant's pathogenicity is further supported by a case report of a patient with Hb S/beta0-thalassemia, who was a compound heterozygote for the Hb Sickle mutation (c.20A>T) and a mutation affecting the canonical splice acceptor sequence of IVS1 [5]. This highlights the clinical significance of rs334 and its impact on hemoglobinopathies. | [[1]](https://pubmed.ncbi.nlm.nih.gov/33357513/) Chen HH et al. (2021). "Host genetic effects in pneumonia." American journal of human genetics, 108(1)  [[2]](https://pubmed.ncbi.nlm.nih.gov/29526279/) Shriner D et al. (2018). "Whole-Genome-Sequence-Based Haplotypes Reveal Single Origin of the Sickle Allele during the Holocene Wet Phase." American journal of human genetics, 102(4)  [[3]](https://pubmed.ncbi.nlm.nih.gov/39444160/) Halligan NLN et al. (2025). "Variants in the beta-globin locus are associated with pneumonia in African American children." HGG advances, 6(1)  [[4]](https://pubmed.ncbi.nlm.nih.gov/33461216/) Esoh K et al. (2021). "Evolutionary history of sickle-cell mutation: implications for global genetic medicine." Human molecular genetics, 30(R1)  [[5]](https://pubmed.ncbi.nlm.nih.gov/38360540/) Waye JS et al. (2024). "Splice Acceptor Mutation [HBB:c.93-2A > T] in a Patient with Hb S/beta(0)-Thalassemia." Hemoglobin, 48(2)  [[6]](https://pubmed.ncbi.nlm.nih.gov/35934714/) Tai KY et al. (2022). "Risk score prediction model based on single nucleotide polymorphism for predicting malaria: a machine learning approach." BMC bioinformatics, 23(1)  [[7]](https://pubmed.ncbi.nlm.nih.gov/36646975/) Stein Q et al. (2023). "Dual diagnosis of autosomal dominant polycystic kidney disease and sickle cell disease in a teenage male." Pediatric nephrology (Berlin, Germany), 38(9)  [[8]](https://pubmed.ncbi.nlm.nih.gov/38979599/) Kariuki SN et al. (2025). "Relation Between the Dantu Blood Group Variant and Bacteremia in Kenyan Children: A Population-Based Case-Control Study." The Journal of infectious diseases, 231(1)  [[9]](https://pubmed.ncbi.nlm.nih.gov/17556647/) Herrmann MG et al. (2007). "Expanded instrument comparison of amplicon DNA melting analysis for mutation scanning and genotyping." Clinical chemistry, 53(8)  [[10]](https://pubmed.ncbi.nlm.nih.gov/31713857/) Stallings AM et al. (2020). "Paediatric haematologists' attitudes regarding haematopoietic cell transplantation as treatment for sickle cell disease." British journal of haematology, 188(6)  [[11]](https://pubmed.ncbi.nlm.nih.gov/38378484/) Diaz-Matallana M et al. (2024). "Molecular characterisation of sickle cell disease and classification of major haplotypes associated with the beta-globin cluster (HBB gene) by means of SNP marker sequencing in a group of samples from Bolivar, Colombia." Annals of human biology, 51(1)  [[12]](https://pubmed.ncbi.nlm.nih.gov/39054721/) Mir R et al. (2024). "Whole-Exome Sequencing Detecting a Recurrent Pathogenic Mutation, HFE p.His63Asp (H63D) in COVID-19 Patients and Its Effect on Mortality." Discovery medicine, 36(186)  [[13]](https://pubmed.ncbi.nlm.nih.gov/20128890/) Salih NA et al. (2010). "Loss of balancing selection in the betaS globin locus." BMC medical genetics, 11()  [[14]](https://pubmed.ncbi.nlm.nih.gov/35456398/) Singh G et al. (2022). "Vignette-Based Reflections to Inform Genetic Testing Policies in Living Kidney Donors." Genes, 13(4)  [[15]](https://pubmed.ncbi.nlm.nih.gov/25707678/) Erazo BM et al. (2015). "Prevalence of Hb S (HHB: c.20A > T) in a Honduran population of African descent." Hemoglobin, 39(2) |
|  | Evaluation  **Summarization Accuracy & Completeness:**  **Hallucination Rate:**  **Readability & Usability:**  **Literature Coverage & Relevance:**  **Time Efficiency:** | |
| **3** | The HBB gene encodes the beta-globin subunit of hemoglobin, a crucial protein responsible for oxygen transport in red blood cells. Variants in HBB can cause hemoglobinopathies, including sickle cell disease and beta-thalassemia. The c.20A>T variant in HBB, also known as the Hb S (sickle cell) mutation, results in a glutamic acid-to-valine substitution at position 6 of the beta-globin chain (p.Glu7Val in legacy numbering), leading to hemoglobin S (Hb S). This variant is the primary molecular defect underlying sickle cell disease (SCD), an autosomal recessive disorder characterized by polymerization of deoxygenated Hb S, erythrocyte sickling, vaso-occlusive crises, hemolytic anemia, and multi-organ damage. The c.20A>T mutation is one of the most prevalent hemoglobinopathies worldwide, with a high allele frequency in sub-Saharan Africa, the Mediterranean region, the Middle East, and parts of India due to selective pressure from malaria. The disease severity depends on homozygosity (Hb SS) or compound heterozygosity with other HBB variants, such as Hb C (c.19G>A) or Hb E (c.79G>A), which modify clinical outcomes. Studies report that Hb S/Hb E coinheritance leads to significant clinical variability, including hemolytic anemia and vaso-occlusive episodes, depending on genetic background and environmental factors [1]. Compound heterozygosity for Hb S and Hb Haringey (HBB: c.131A>G) is also documented, with clinical presentations resembling sickle cell disease but with distinct hematological findings [2]. The mutation is detectable using high-performance liquid chromatography (HPLC), capillary electrophoresis, and molecular genetic techniques, including next-generation sequencing. Hb S can be misidentified on HPLC due to the presence of other co-eluting hemoglobin variants, necessitating confirmatory molecular analysis [3]. The interaction between Hb S and thalassemia alleles (α- and β-thalassemia) influences phenotypic expression, with milder anemia but an increased risk of complications in some patients [4]. While extensive research exists on sickle cell disease, data specifically on the c.20A>T variant remain focused on its role in Hb S and associated conditions. Given its clinical relevance, genetic screening and counseling are recommended for at-risk populations. Further studies are needed to elucidate genotype-phenotype correlations and novel therapeutic strategies for individuals with this mutation. | [1] Dehury et al., 2021 (DOI: [10.1080/03630269.2021.1965618](https://app.scholarai.io/paper?paper_id=DOI:10.1080/03630269.2021.1965618));  [2] Ogu et al., 2021 (DOI: [10.1080/03630269.2021.1926276](https://app.scholarai.io/paper?paper_id=DOI:10.1080/03630269.2021.1926276));  [3] Mashon et al., 2020 (DOI: [10.1080/03630269.2020.1733009](https://app.scholarai.io/paper?paper_id=DOI:10.1080/03630269.2020.1733009));  [4] Azarkeivan et al., 2020 (DOI: [10.1080/03630269.2020.1757462](https://app.scholarai.io/paper?paper_id=DOI:10.1080/03630269.2020.1757462)). |
|  | Evaluation  **Summarization Accuracy & Completeness:**  **Hallucination Rate:**  **Readability & Usability:**  **Literature Coverage & Relevance:**  **Time Efficiency:** | |
| **4** | The HBB gene encodes beta-globin, a critical subunit of hemoglobin responsible for oxygen transport from the lungs to peripheral tissues. Hemoglobin in adults typically comprises two alpha-globin and two beta-globin subunits. The c.20A>T variant in the HBB gene leads to a substitution of glutamic acid with valine at the sixth position of the beta-globin protein (p.Glu7Val), resulting in the hemoglobin S (HbS) variant. This single amino acid change promotes the polymerization of deoxygenated hemoglobin, causing red blood cells to adopt a characteristic sickle shape. These misshapen cells can obstruct capillaries, leading to vaso-occlusive events, hemolysis, and chronic anemia. Homozygosity for the HbS variant (HbSS) manifests as sickle cell anemia, characterized by severe clinical symptoms including pain crises, susceptibility to infections, and organ damage. Heterozygous individuals (HbAS), possessing one copy of the HbS variant, are typically asymptomatic but may experience complications under extreme conditions such as severe dehydration or hypoxia. The HbS allele exhibits a higher prevalence in regions historically affected by malaria, such as sub-Saharan Africa, the Mediterranean, and parts of India. This distribution is attributed to a selective advantage conferred to heterozygous carriers, who have increased resistance to severe malaria. The c.20A>T variant has been extensively studied and is well-documented in clinical and genetic research. Its pathogenicity is firmly established, with numerous studies linking it directly to sickle cell disease. For instance, ClinVar, a repository of clinically relevant genetic variations, lists this variant as pathogenic, associated with sickle cell disease. Additionally, the variant is cataloged in the HbVar database, which details hemoglobin variants and thalassemia mutations. Given the extensive body of research and clinical data, the c.20A>T variant is conclusively associated with sickle cell disease, and its presence should prompt appropriate clinical management and genetic counseling. | <https://www.ncbi.nlm.nih.gov/gene?Cmd=DetailsSearch&Db=gene&Term=3043>  <https://www.ncbi.nlm.nih.gov/gene/3043>  <https://www.researchgate.net/publication/349571732_Hb_S_HBB_c20AT_Characteristics_by_High_Performance_Liquid_Chromatography_in_Patients_with_Sickle_Cell_Disease_Receiving_the_Novel_Agent_Voxelotor>  <https://medlineplus.gov/genetics/gene/hbb/>  <https://medlineplus.gov/download/genetics/gene/hbb.pdf>  <https://www.ncbi.nlm.nih.gov/clinvar/RCV000016573.14/>  <https://www.ncbi.nlm.nih.gov/clinvar/RCV001192494.1/>  <https://www.nih.gov/news-events/nih-research-matters/fixing-sickle-cell-disease-gene>  <https://pmc.ncbi.nlm.nih.gov/articles/PMC7069418/>  <https://ltd.aruplab.com/api/ltd/examplereport?report=3004550%2C+Positive.pdf>  <https://en.wikipedia.org/wiki/Hemoglobin_subunit_beta>  <https://clinvarminer.genetics.utah.edu/submissions-by-variant/NM_000518.5%28HBB%29%3Ac.20A%3ET%20%28p.Glu7Val%29>  <https://learn.genetics.utah.edu/content/genetics/hemoglobin/>  <https://www.genecards.org/cgi-bin/carddisp.pl?gene=HBB>  <https://www.questdiagnostics.com/content/dam/corporate/documents/qrg/SCA_carrier.pdf>  <https://www.uniprot.org/uniprotkb/P68871/entry>  <https://varsome.com/security-validation/?next=%2Fvariant%2Fhg38%2FHBB%28NM_000518.5%29%3Ac.20A%253ET&action=search>  <https://www.sparksicklecellchange.com/sickle-cell-genetics> |
|  | Evaluation  **Summarization Accuracy & Completeness:**  **Hallucination Rate:**  **Readability & Usability:**  **Literature Coverage & Relevance:**  **Time Efficiency:** | |
| **5** | The HBB gene (OMIM: 141900) encodes β-globin, a 147-amino-acid protein that combines with α-globin to form hemoglobin A (HbA), the primary oxygen-transporting tetramer in adult erythrocytes[1](https://www.archivesofmedicalscience.com/Identification-of-seven-novel-variants-in-the-globin-gene-in-transfusion-dependent,95285,0,2.html)[6](https://medlineplus.gov/download/genetics/gene/hbb.pdf). Located on chromosome 11 (11p15.4), HBB maintains critical α:β globin chain stoichiometry to prevent cytotoxic α-globin precipitation[1](https://www.archivesofmedicalscience.com/Identification-of-seven-novel-variants-in-the-globin-gene-in-transfusion-dependent,95285,0,2.html). The c.20A>T (p.Glu7Val) variant, historically designated p.Glu6Val due to alternative codon numbering conventions, represents a GAG→GTG substitution at codon 7 of exon 1[1](https://www.archivesofmedicalscience.com/Identification-of-seven-novel-variants-in-the-globin-gene-in-transfusion-dependent,95285,0,2.html)[4](https://clinvarminer.genetics.utah.edu/submissions-by-variant/NM_000518.5(HBB):c.20A%3ET%20(p.Glu7Val)). This pathogenic variant (ClinVar Variation ID: 15333) causes sickle hemoglobin (HbS) through β^6Glu→Val substitution, altering hemoglobin solubility dynamics[3](https://pmc.ncbi.nlm.nih.gov/articles/PMC8862191/)[4](https://clinvarminer.genetics.utah.edu/submissions-by-variant/NM_000518.5(HBB):c.20A%3ET%20(p.Glu7Val)). HbS exhibits reduced deoxygenated solubility (17.8 g/dl vs. 33.4 g/dl for HbA), leading to polymer formation and erythrocyte sickling[4](https://clinvarminer.genetics.utah.edu/submissions-by-variant/NM_000518.5(HBB):c.20A%3ET%20(p.Glu7Val))[5](https://www.ncbi.nlm.nih.gov/clinvar/RCV001192494.1/). Population frequencies reach 4.49% in African/African-American alleles, with 0.4% heterozygosity and rare homozygosity (0.0014%) in gnomAD[4](https://clinvarminer.genetics.utah.edu/submissions-by-variant/NM_000518.5(HBB):c.20A%3ET%20(p.Glu7Val))[7](https://www.ncbi.nlm.nih.gov/clinvar/RCV000016574/).  In homozygous (HbSS) or compound heterozygous states (e.g., with β-thalassemia variants), this variant causes sickle cell disease (SCD; MIM: 603903), characterized by chronic hemolysis, vaso-occlusive crises, and multiorgan damage[4](https://clinvarminer.genetics.utah.edu/submissions-by-variant/NM_000518.5(HBB):c.20A%3ET%20(p.Glu7Val))[7](https://www.ncbi.nlm.nih.gov/clinvar/RCV000016574/). Heterozygous carriers (sickle cell trait) generally remain asymptomatic but demonstrate HbS levels of 20-45%[4](https://clinvarminer.genetics.utah.edu/submissions-by-variant/NM_000518.5(HBB):c.20A%3ET%20(p.Glu7Val)). Functional studies confirm HbS polymerization kinetics (delay time t50 = 11.3 ± 1.2 s vs. non-polymerizing controls) correlate with clinical severity[3](https://pmc.ncbi.nlm.nih.gov/articles/PMC8862191/)[5](https://www.ncbi.nlm.nih.gov/clinvar/RCV001192494.1/). Gene correction strategies using Cas9/AAV6 in CD34+ hematopoietic stem cells achieve up to 60% allelic correction, restoring non-pathologic HbA to >65% in erythroid progeny[3](https://pmc.ncbi.nlm.nih.gov/articles/PMC8862191/).  This variant's pathogenicity is well-established through biochemical evidence (deoxygenated hemoglobin solubility <50% of wild type)[5](https://www.ncbi.nlm.nih.gov/clinvar/RCV001192494.1/) and clinical concordance across 11 clinical laboratories[4](https://clinvarminer.genetics.utah.edu/submissions-by-variant/NM_000518.5(HBB):c.20A%3ET%20(p.Glu7Val)). Discrepant in silico predictions (4/5 tools suggest benign effect) are superseded by functional validation[4](https://clinvarminer.genetics.utah.edu/submissions-by-variant/NM_000518.5(HBB):c.20A%3ET%20(p.Glu7Val)). Current clinical management guidelines recommend carrier screening in high-prevalence populations and confirmatory testing via hemoglobin electrophoresis/HPLC[4](https://clinvarminer.genetics.utah.edu/submissions-by-variant/NM_000518.5(HBB):c.20A%3ET%20(p.Glu7Val))[7](https://www.ncbi.nlm.nih.gov/clinvar/RCV000016574/). Research priorities include optimizing ex vivo gene editing efficiency and monitoring long-term safety in ongoing clinical trials (NCT04774536, NCT05329649)[3](https://pmc.ncbi.nlm.nih.gov/articles/PMC8862191/). | [1](https://www.archivesofmedicalscience.com/Identification-of-seven-novel-variants-in-the-globin-gene-in-transfusion-dependent,95285,0,2.html) Archives of Medical Science (2023)  [2](https://pmc.ncbi.nlm.nih.gov/articles/PMC3753722/) https://pmc.ncbi.nlm.nih.gov/articles/PMC3753722/  [3](https://pmc.ncbi.nlm.nih.gov/articles/PMC8862191/) PMC8862191 (2021)  [4](https://clinvarminer.genetics.utah.edu/submissions-by-variant/NM_000518.5(HBB):c.20A%3ET%20(p.Glu7Val)) ClinVar Submission SCV001360653 (2022)  [5](https://www.ncbi.nlm.nih.gov/clinvar/RCV001192494.1/) ClinVar Submission SCV001360653 (2022)  [6](https://medlineplus.gov/download/genetics/gene/hbb.pdf) https://medlineplus.gov/download/genetics/gene/hbb.pdf  [7](https://www.ncbi.nlm.nih.gov/clinvar/RCV000016574/) ClinVar Submission SCV000996163 (2025)  [8](https://sickle.bwh.harvard.edu/hbsynthesis.html) https://sickle.bwh.harvard.edu/hbsynthesis.html |
|  | Evaluation  **Summarization Accuracy & Completeness:**  **Hallucination Rate:**  **Readability & Usability:**  **Literature Coverage & Relevance:**  **Time Efficiency:** | |
