## Supplementary material for "Benchmarking generative AI tools for literature retrieval and summarization in genomic variant interpretation": HTT c.529-3C_T.docx

| **Tool** | **Summary** | **References** |
| --- | --- | --- |
| **1** | The HTT gene encodes the huntingtin protein, which plays a crucial role in neuronal development and function, particularly in the brain, where it is highly expressed[7]. The gene is well-known for its association with Huntington's disease (HD), a neurodegenerative disorder caused by expansions of CAG repeats beyond 39[3][4]. However, the specific variant c.529-3C>T has not been directly referenced in the available literature. Generally, mutations in the HTT gene can lead to a dominant negative effect on the wild-type protein, contributing to HD phenotypes[4]. The HTT gene's function involves roles in chemical signaling, material transport, protein binding, and protection against apoptosis[7]. While CAG repeats within the non-pathological range have been studied for their influence on cognitive functions and personality traits[3][6], specific information about the c.529-3C>T variant is lacking. Therefore, without direct evidence, it is challenging to determine its classification, frequency, or impact. Further investigation is recommended to clarify the significance of this variant. References:[3][4][6][7]. | [1] <https://www.ncbi.nlm.nih.gov/pmc/articles/PMC10270637/>  [2] <https://pubmed.ncbi.nlm.nih.gov/35829908/>  [3] <https://www.ncbi.nlm.nih.gov/pmc/articles/PMC8965717/>  [4] <https://www.semanticscholar.org/paper/db6b322837cb3385686206a7c16ec37657abc954>  [5] <https://www.ncbi.nlm.nih.gov/pmc/articles/PMC7093057/>  [6] <https://www.ncbi.nlm.nih.gov/pmc/articles/PMC8228729/>  [7] <https://www.semanticscholar.org/paper/794ce7dfe1a458122f95f61dbd4828a40248327a>  [8] <https://www.ncbi.nlm.nih.gov/pmc/articles/PMC5538116/> |
|  | Evaluation  **Summarization Accuracy & Completeness:**  **Hallucination Rate:**  **Readability & Usability:**  **Literature Coverage & Relevance:**  **Time Efficiency:** | |
| **2** | The HTT gene encodes huntingtin, a protein expressed in various tissues, with the highest concentrations in the brain and testes. Huntingtin is involved in numerous cellular processes, including transcription regulation, intracellular signaling, and vesicular transport. Mutations in HTT, particularly expansions of CAG repeats, are known to cause Huntington's disease (HD), a neurodegenerative disorder characterized by motor dysfunction, cognitive decline, and psychiatric symptoms. The specific single nucleotide variant c.529-3C>T is located in intron 7 of the HTT gene, three nucleotides upstream of exon 8. According to ClinVar, this variant is classified as likely benign, with multiple submitters providing criteria and no conflicts reported. The molecular consequence is an intron variant, and it has been assigned the dbSNP ID rs202150164. The Exome Variant Server reports a frequency of 11 occurrences in 12,788 alleles, with all instances found in individuals of European American descent and none in African American individuals. The average frequency in gnomAD v.2.1.1 is not provided. Given its classification as likely benign and its presence in the general population without apparent association with disease, the c.529-3C>T variant is not currently considered to have clinical significance. However, as with all genetic findings, it is essential to interpret this variant in the context of the patient's overall clinical presentation and family history. Further functional studies could provide more definitive insights into any potential effects of this variant on HTT gene function. | [1] ClinVar entry for NM_002111.6(HTT):c.529-3C>T:<https://www.ncbi.nlm.nih.gov/clinvar/variation/VCV001539285/>  [2] Exome Variant Server data for HTT c.529-3C>T:<https://databases.lovd.nl/whole_genome/variants/0000461062>  Other:  <https://databases.lovd.nl/whole_genome/transcripts/00007927>  <https://pmc.ncbi.nlm.nih.gov/articles/PMC10940305/>  [https://genome.ucsc.edu/cgi-bin/hgc?c=chr4&db=hg19&g=clinvarMain&hgsid=2241418966_DvqXzCaZcF48aoLO5i7AqCCyzDjg&i=C%3ET&l=3000710&o=3107080&r=3205710&t=3107081](https://genome.ucsc.edu/cgi-bin/hgc?c=chr4&db=hg19&g=clinvarMain&hgsid=2241418966_DvqXzCaZcF48aoLO5i7AqCCyzDjg&i=C%3ET&l=3000710&o=3107080&r=3205710&t=3107081&utm_source=chatgpt.com)  <https://www.savingforcollege.com/article/roll-over-529-plan-funds-to-a-roth-ira>  <https://www.nature.com/articles/s43856-025-00752-7>  <https://www.irs.gov/taxtopics/tc313>  <https://www.biorxiv.org/content/10.1101/2022.02.02.478920v1.full.pdf>  <https://www.nature.com/articles/s41467-020-20573-7>  <https://elifesciences.org/articles/45056>  <https://www.science.org/doi/10.1126/scitranslmed.aap9927>  <https://insight.jci.org/articles/view/88995>  <https://www.federalregister.gov/documents/2008/01/18/E8-859/guidance-on-qualified-tuition-programs-under-section-529>  <https://www.ncbi.nlm.nih.gov/clinvar/RCV001833094/>  <https://pmc.ncbi.nlm.nih.gov/articles/PMC3178454/>  <https://clinvarminer.genetics.utah.edu/variants-by-gene/HTT/submitter/500031/likely%20benign>  <https://www.law.cornell.edu/uscode/text/26/529>  <https://onlinebanking.huntington.com/rol/Auth/login.aspx> |
|  | Evaluation  **Summarization Accuracy & Completeness:**  **Hallucination Rate:**  **Readability & Usability:**  **Literature Coverage & Relevance:**  **Time Efficiency:** | |
| **3** | Variant c.529-3C>T rs202150164 is a single nucleotide substitution located in the splice region of the HTT gene, specifically affecting the polypyrimidine tract. This variant is classified as a splice region variant, splice polypyrimidine tract variant, and intron variant, indicating its potential impact on the splicing process of the pre-mRNA transcript. The alteration occurs in the consensus sequence of the splice site, which may affect the recognition of the splice site by the splicing machinery and consequently alter the proper splicing of the HTT gene transcript.  The HTT gene encodes the huntingtin protein, which is widely expressed in human tissues, with particularly high expression in the brain. The exact function of huntingtin is not fully understood, but it is known to play a critical role in neuronal function and survival. Huntingtin is involved in various cellular processes, including vesicular transport, postsynaptic signaling, and the regulation of gene transcription. Abnormalities in the HTT gene, particularly expansions of CAG trinucleotide repeats, are known to cause Huntington's disease, a progressive neurodegenerative disorder.  Regarding the specific variant c.529-3C>T rs202150164, it has been reported in ClinVar with one benign submission. This suggests that at least one submitter has provided evidence indicating a lack of pathogenicity for this variant. However, the benign classification does not rule out the possibility of functional consequences, and the variant's impact on splicing should be further investigated through functional assays or additional clinical data to fully elucidate its significance. | No references available |
|  | Evaluation  **Summarization Accuracy & Completeness:**  **Hallucination Rate:**  **Readability & Usability:**  **Literature Coverage & Relevance:**  **Time Efficiency:** | |
| **4** | The HTT gene, located on chromosome 4p16.3, encodes the huntingtin protein, a widely expressed molecule involved in various cellular processes including vesicular transport, transcriptional regulation, and synaptic transmission. Pathogenic expansions of the CAG trinucleotide repeat within exon 1 of HTT are classically associated with Huntington’s disease; however, variants outside the repeat region may also have clinical significance, particularly if they impact splicing or regulatory functions. The c.529-3C>T variant is an intronic change occurring at position -3 upstream of exon 6, situated in the canonical acceptor splice site region. Variants at this location can potentially disrupt normal RNA splicing and lead to aberrant or truncated protein products, although direct functional data for c.529-3C>T is currently lacking in the peer-reviewed literature. No specific studies have been published that directly evaluate the pathogenicity, allele frequency, or clinical implications of the c.529-3C>T substitution in HTT, nor are there curated classifications in ClinVar or locus-specific variant databases as of this report. Given its intronic location near a splice acceptor site, in silico splicing models may predict potential deleterious effects, but such predictions require empirical validation. Without functional studies or observed segregation data, the clinical relevance of this variant remains uncertain. It is recommended that further functional assays or transcript analyses be pursued to clarify any impact on splicing, and that the variant be interpreted with caution, especially in diagnostic settings. At this time, no peer-reviewed publications directly investigate the HTT c.529-3C>T variant [1]. | No references available |
|  | Evaluation  **Summarization Accuracy & Completeness:**  **Hallucination Rate:**  **Readability & Usability:**  **Literature Coverage & Relevance:**  **Time Efficiency:** | |
| **5** | The HTT gene, located on chromosome 4, encodes the protein huntingtin, which plays a crucial role in neuronal function and development.  Although the precise function of huntingtin is not fully understood, it is known to be involved in various cellular processes, including vesicle trafficking, microtubule-mediated transport, and transcriptional regulation. Huntingtin interacts with proteins like HIP1 to mediate endocytosis and is associated with vesicles and microtubules, suggesting roles in cytoskeletal anchoring and mitochondrial transport. It also upregulates brain-derived neurotrophic factor (BDNF) expression, essential for neuronal survival and function. The HTT gene contains a CAG trinucleotide repeat region, which normally repeats 10 to 35 times. Expansion of this repeat beyond 36 repeats is associated with Huntington's disease (HD), a neurodegenerative disorder characterized by progressive motor, cognitive, and psychiatric symptoms. The expanded CAG repeat leads to the production of a mutant huntingtin protein with an elongated polyglutamine tract, resulting in a toxic gain-of-function phenotype. This mutant protein forms aggregates that disrupt cellular homeostasis, impair protein degradation systems, and co-aggregate with other essential proteins, leading to neuronal dysfunction and death, particularly in the striatum and cerebral cortex. The c.529-3C>T variant in the HTT gene is a splice site variant that has not been extensively studied in the literature. Its classification and impact on protein function remain unclear due to limited research. The frequency of this variant in the general population is not well-documented, and its role in disease pathogenesis is uncertain. Further investigation is needed to determine the pathogenicity and clinical significance of the c.529-3C>T variant. Given the lack of direct references, it is recommended to conduct functional studies and gather more data to assess its potential impact on HTT gene function and association with HD or other neurological conditions. | [1] MedlinePlus Genetics. HTT gene. Available at: <https://medlineplus.gov/genetics/gene/htt/> 1.  [2] PMC. The biological function of the Huntingtin protein and its relevance to Huntington’s Disease pathology. Available at: <https://www.ncbi.nlm.nih.gov/pmc/articles/PMC3237673/> 2.  [3] Wikipedia. Huntingtin. Available at: <https://en.wikipedia.org/wiki/Huntingtin> 3.  [4] GeneCards. HTT Gene. Available at: <https://www.genecards.org/cgi-bin/carddisp.pl?gene=HTT> 4. |
|  | Evaluation  **Summarization Accuracy & Completeness:**  **Hallucination Rate:**  **Readability & Usability:**  **Literature Coverage & Relevance:**  **Time Efficiency:** | |
