## Supplementary material for "Benchmarking generative AI tools for literature retrieval and summarization in genomic variant interpretation": KCNQ1 p.Ala341Glu.docx

| **Tool** | **Summary** | **References** |
| --- | --- | --- |
| **1** | The genomic variant c.1022C>A p.Ala341Glu rs12720459 on the KCNQ1 gene is a missense mutation that has been extensively studied due to its association with long QT syndrome type 1 (LQT1), a condition characterized by prolonged cardiac repolarization and a risk of arrhythmias and sudden cardiac death. The KCNQ1 gene encodes the alpha subunit of the voltage-gated potassium channel Kv7.1, which is essential for the cardiac slowly activating delayed-rectifier potassium current (IKs). This current is crucial for the repolarization phase of the cardiac action potential, and its proper function is vital for maintaining normal cardiac rhythm.  Functional studies have demonstrated that the p.Ala341Glu variant results in a channel with altered electrophysiological properties. Specifically, when the mutant KCNQ1 is co-expressed with the wild-type KCNE1 beta-subunit, the resulting current exhibits a smaller current density, slower activation rates, and a depolarized shift in the activation curve compared to the wild-type KCNQ1+KCNE1 channels [1]. This suggests that the p.Ala341Glu variant leads to a functionally defective protein that, despite being trafficked to the cell surface, impairs the IKs current.  Clinical data from a founder effect in a South African population, where many individuals share the mutation identical by descent, have provided insights into the phenotypic heterogeneity and severity associated with this variant. The p.Ala341Glu variant is linked to an unusually severe phenotype, with mutation carriers (MCs) exhibiting a wide range of QTc values and a higher risk of cardiac events, especially when the heart rate is below a certain threshold [2]. This severity is further supported by the observation that MCs with the p.Ala341Glu variant are more symptomatic by age 40 and experience symptoms earlier than those with other LQT1 mutations [6].  The clinical severity of the p.Ala341Glu variant is influenced by its dominant-negative effect on wild-type channels and the impairment of IKs regulation, particularly under conditions of beta-adrenergic stimulation [4]. This is further complicated by the loss of protein kinase A (PKA)-dependent IKs enhancement, which is specific to the p.Ala341Glu variant and correlates with its phenotypic severity [3]. The S6 segment, where p.Ala341Glu is located, is critical for channel activation, and mutations in this region, including those neighboring p.Ala341, are associated with increased arrhythmic risk [3].  The p.Ala341Glu variant also exhibits a dominant-negative suppression of cAMP-dependent Yotiao-mediated IKs upregulation, which is essential for the response to beta-adrenergic stimulation. This suppression is due to reduced phosphorylation of KCNQ1 at S27, even in heterozygous conditions, indicating that phosphorylation defects in some mutant KCNQ1 subunits can completely suppress IKs upregulation [4].  In addition to the functional and clinical data, the p.Ala341Glu variant has been characterized in human-induced pluripotent stem cell-derived cardiomyocytes (hiPSC-CMs), where it showed normal protein trafficking, a mild dominant-negative effect, and impaired PKA-dependent IKs stimulation [5].  Overall, the c.1022C>A p.Ala341Glu rs12720459 variant on the KCNQ1 gene is pathogenic, with multiple studies confirming its association with a severe phenotype in LQT1 patients. The variant affects the channel's biophysical properties and regulatory mechanisms, leading to significant clinical implications for affected individuals. ClinVar submissions further support the pathogenicity of this variant, with a total of 13 pathogenic submissions to date. | [[1]](https://pubmed.ncbi.nlm.nih.gov/21854832/) Mikuni I et al. (2011). "Partial restoration of the long QT syndrome associated KCNQ1 A341V mutant by the KCNE1 beta-subunit." Biochimica et biophysica acta, 1810(12)  [[2]](https://pubmed.ncbi.nlm.nih.gov/16246960/) Brink PA et al. (2005). "Phenotypic variability and unusual clinical severity of congenital long-QT syndrome in a founder population." Circulation, 112(17)  [[3]](https://pubmed.ncbi.nlm.nih.gov/34505893/) Schwartz PJ et al. (2021). "Mutation location and IKs regulation in the arrhythmic risk of long QT syndrome type 1: the importance of the KCNQ1 S6 region." European heart journal, 42(46)  [[4]](https://pubmed.ncbi.nlm.nih.gov/22095730/) Heijman J et al. (2012). "Dominant-negative control of cAMP-dependent IKs upregulation in human long-QT syndrome type 1." Circulation research, 110(2)  [[5]](https://pubmed.ncbi.nlm.nih.gov/37099628/) Giannetti F et al. (2023). "Gene- and variant-specific efficacy of serum/glucocorticoid-regulated kinase 1 inhibition in long QT syndrome types 1 and 2." Europace : European pacing, arrhythmias, and cardiac electrophysiology : journal of the working groups on cardiac pacing, arrhythmias, and cardiac cellular electrophysiology of the European Society of Cardiology, 25(5)  [[6]](https://pubmed.ncbi.nlm.nih.gov/17984373/) Crotti L et al. (2007). "The common long-QT syndrome mutation KCNQ1/A341V causes unusually severe clinical manifestations in patients with different ethnic backgrounds: toward a mutation-specific risk stratification." Circulation, 116(21)  [[7]](https://pubmed.ncbi.nlm.nih.gov/25087618/) de Villiers CP et al. (2014). "AKAP9 is a genetic modifier of congenital long-QT syndrome type 1." Circulation. Cardiovascular genetics, 7(5)  [[8]](https://pubmed.ncbi.nlm.nih.gov/15028050/) Kobori A et al. (2004). "Additional gene variants reduce effectiveness of beta-blockers in the LQT1 form of long QT syndrome." Journal of cardiovascular electrophysiology, 15(2)  [[9]](https://pubmed.ncbi.nlm.nih.gov/16627448/) Anastasakis A et al. (2006). "Phenotype reveals genotype in a Greek long QT syndrome family." Europace : European pacing, arrhythmias, and cardiac electrophysiology : journal of the working groups on cardiac pacing, arrhythmias, and cardiac cellular electrophysiology of the European Society of Cardiology, 8(4)  [[10]](https://pubmed.ncbi.nlm.nih.gov/17010804/) Heradien MJ et al. (2006). "Does pregnancy increase cardiac risk for LQT1 patients with the KCNQ1-A341V mutation?" Journal of the American College of Cardiology, 48(7)  [[11]](https://pubmed.ncbi.nlm.nih.gov/18308161/) Schwartz PJ et al. (2008). "Neural control of heart rate is an arrhythmia risk modifier in long QT syndrome." Journal of the American College of Cardiology, 51(9)  [[12]](https://pubmed.ncbi.nlm.nih.gov/25585005/) Mikuni I et al. (2015). "Enhanced effects of isoflurane on the long QT syndrome 1-associated A341V mutant." Anesthesiology, 122(4)  [[13]](https://pubmed.ncbi.nlm.nih.gov/31398660/) Mura M et al. (2019). "Generation of two human induced pluripotent stem cell (hiPSC) lines from a long QT syndrome South African founder population." Stem cell research, 39()  [[14]](https://pubmed.ncbi.nlm.nih.gov/24705789/) Bari V et al. (2014). "Multiscale complexity analysis of the cardiac control identifies asymptomatic and symptomatic patients in long QT syndrome type 1." PloS one, 9(4)  [[15]](https://pubmed.ncbi.nlm.nih.gov/16155735/) Lai LP et al. (2005). "Denaturing high-performance liquid chromatography screening of the long QT syndrome-related cardiac sodium and potassium channel genes and identification of novel mutations and single nucleotide polymorphisms." Journal of human genetics, 50(9) |
|  | Evaluation  **Summarization Accuracy & Completeness:**  **Hallucination Rate:**  **Readability & Usability:**  **Literature Coverage & Relevance:**  **Time Efficiency:** | |
| **2** | The KCNQ1 gene encodes the Kv7.1 potassium channel α-subunit, which is crucial for cardiac repolarization by providing the slow delayed rectifier current (IKs) essential for maintaining a regular heart rhythm[2][3]. This gene is also expressed in the inner ear, where it helps maintain ion balance necessary for normal hearing, and in other tissues like the kidney, lung, stomach, and intestine[3]. Variants in KCNQ1 are associated with several conditions, including long QT syndrome type 1 (LQT1), which is characterized by loss-of-function mutations leading to QT prolongation and increased risk of cardiac arrhythmias[2][5]. The specific variant p.Ala341Glu in the KCNQ1 gene has not been directly referenced in the available literature, suggesting that its impact and classification may require further investigation. Generally, KCNQ1 mutations can lead to either haploinsufficiency or dominant-negative effects, influencing the risk of cardiac events[5]. While KCNQ1 variants have been linked to other conditions, such as type 2 diabetes and ischemic stroke[1][6], the direct implications of the p.Ala341Glu variant remain unclear due to a lack of specific research findings. Therefore, additional studies are necessary to determine its clinical significance and potential impact on cardiac function or other related conditions. | [1] https://pubmed.ncbi.nlm.nih.gov/26970180/  [2] https://pubmed.ncbi.nlm.nih.gov/33504163/  [3] https://www.semanticscholar.org/paper/5e40c8e0b93442bad6d1e8e86630cab86dd7f52f  [4] https://www.ncbi.nlm.nih.gov/pmc/articles/PMC11120621/  [5] https://pubmed.ncbi.nlm.nih.gov/17470695/  [6] https://pubmed.ncbi.nlm.nih.gov/30816480/  [7] https://www.ncbi.nlm.nih.gov/pmc/articles/PMC7069191/  [8] https://www.ncbi.nlm.nih.gov/pmc/articles/PMC10206659/ |
|  | Evaluation  **Summarization Accuracy & Completeness:**  **Hallucination Rate:**  **Readability & Usability:**  **Literature Coverage & Relevance:**  **Time Efficiency:** | |
| **3** | The KCNQ1 gene encodes the alpha subunit of a voltage-gated potassium channel, Kv7.1, which is crucial for the repolarization phase of the cardiac action potential. This channel, in conjunction with its auxiliary subunit KCNE1, generates the slowly activating delayed rectifier potassium current (IKs), essential for maintaining normal heart rhythm. Pathogenic variants in KCNQ1 are the most common genetic cause of congenital long QT syndrome (LQTS), specifically type 1 (LQTS1), and are also implicated in Jervell and Lange-Nielsen syndrome (JLNS), which combines LQTS with congenital sensorineural hearing loss. The variant p.Ala341Glu in KCNQ1 is a missense mutation where alanine is replaced by glutamic acid at position 341. This variant has been identified in individuals with LQTS, highlighting its potential impact on cardiac function. The precise frequency of this variant in the general population is not well-documented, but it is considered rare. Functional studies have shown that variants in KCNQ1, including p.Ala341Glu, can disrupt the normal function of the potassium channel, leading to prolonged QT intervals and increased risk of life-threatening arrhythmias. The classification of this variant according to the American College of Medical Genetics and Genomics (ACMG) guidelines is likely pathogenic, based on its association with LQTS and the functional data supporting its deleterious effect on the potassium channel. However, the exact mechanisms by which p.Ala341Glu affects channel function and contributes to the clinical phenotype are not fully elucidated. Further research is needed to understand the genotype-phenotype correlation and to develop targeted therapies for individuals carrying this variant. The clinical management of patients with this variant typically involves beta-blocker therapy to reduce the risk of arrhythmias, along with regular cardiac monitoring. Genetic counseling is also recommended for families with a history of LQTS or JLNS to discuss the inheritance pattern and potential risks to other family members. In conclusion, the p.Ala341Glu variant in KCNQ1 is a significant contributor to LQTS, underscoring the importance of genetic testing and clinical surveillance in affected individuals. | [1] Nishimura M, Ueda M, Ebata R, Utsuno E, Ishii T, Matsushita K, Ohara O, Shimojo N, Kobayashi Y, Nomura F. A novel KCNQ1 nonsense variant in the isoform-specific first exon causes both jervell and Lange-Nielsen syndrome 1 and long QT syndrome 1: a case report. BMC Med Genet. 2017.  [2] Vyas B, Puri RD, Namboodiri N, Nair M, Sharma D, Movva S, Saxena R, Bohora S, Aggarwal N, Vora A, Kumar J, Singh T, Verma IC. KCNQ1 mutations associated with Jervell and Lange-Nielsen syndrome and autosomal recessive Romano-Ward syndrome in India-expanding the spectrum of long QT syndrome type 1. Am J Med Genet A.  [3] Zhang S, et al. LQTS in Northern BC: homozygosity for KCNQ1 V205M presents with a more severe cardiac phenotype but with minimal impact on auditory function. BMC Med Genet. 2008.  [4] Zehelein J, Kathoefer S, Khalil M, Alter M, Thomas D, Brockmeier K, et al. Skipping of exon 1 in the KCNQ1 gene causes jervell and Lange-Nielsen syndrome. Nat Genet. 1997. |
|  | Evaluation  **Summarization Accuracy & Completeness:**  **Hallucination Rate:**  **Readability & Usability:**  **Literature Coverage & Relevance:**  **Time Efficiency:** | |
| **4** | The KCNQ1 gene encodes a voltage-gated potassium channel alpha subunit critical to the cardiac action potential, particularly in the repolarization phase, by contributing to the slow delayed rectifier potassium current (IKs). Variants in KCNQ1 are most commonly associated with Long QT Syndrome type 1 (LQT1), a condition that predisposes individuals to potentially fatal arrhythmias such as torsades de pointes and sudden cardiac death. The missense variant p.Ala341Glu (also referred to as A341E, resulting from c.1022C>A) substitutes alanine with glutamic acid at position 341 in the S6 transmembrane segment of the KCNQ1 protein, a region crucial for channel gating and ion conductance. While p.Ala341Glu has been frequently reported in the literature in individuals with LQT1, no specific peer-reviewed studies directly evaluating the functional or clinical consequences of this variant were found in the current academic databases searched. However, the residue A341 has been implicated in other variants (e.g., A341V) associated with pathogenic outcomes, suggesting that substitutions at this position may disrupt channel function. Functional analyses of variants at nearby residues indicate significant reductions in IKs current and dominant-negative effects on channel activity, reinforcing the likelihood that A341E is a deleterious variant, although this has not been confirmed for this exact substitution. In clinical practice, p.Ala341Glu is generally classified as likely pathogenic based on its location within a highly conserved transmembrane domain, its rarity in general population databases, and consistent segregation with disease in affected families. However, due to the absence of variant-specific functional or large-scale epidemiological data, the classification should be treated with caution, and further experimental validation or family studies are recommended. No peer-reviewed publications directly characterizing p.Ala341Glu were identified [1]. | [1] No direct peer-reviewed literature specific to p.Ala341Glu (A341E) in KCNQ1 found; search conducted via [Nature Genetics](https://www.nature.com/articles/s41431-020-00740-6). |
|  | Evaluation  **Summarization Accuracy & Completeness:**  **Hallucination Rate:**  **Readability & Usability:**  **Literature Coverage & Relevance:**  **Time Efficiency:** | |
| **5** | The KCNQ1 gene encodes the alpha subunit of the Kv7.1 potassium channel, which is crucial for the slow delayed rectifier potassium current (I_Ks) responsible for the repolarization phase of the cardiac action potential . Mutations in KCNQ1 are associated with long QT syndrome type 1 (LQT1), a condition characterized by a prolonged QT interval on electrocardiograms, leading to an increased risk of arrhythmias and sudden cardiac events . The specific missense variant p.Ala341Glu (c.1022C>A) results in the substitution of alanine with glutamic acid at position 341 of the Kv7.1 protein. This variant has been classified as pathogenic and is implicated in LQT1 . Functional studies indicate that the p.Ala341Glu mutation leads to a significant reduction in I_Ks currents, approximately by 23%, due to the formation of almost inactive homomeric channels . Clinically, individuals heterozygous for this mutation may exhibit a mild LQTS phenotype, with rare clinical symptoms. However, the phenotypic expression can vary, and some studies suggest that mutations at position 341, such as p.Ala341Val, are associated with unusually severe clinical manifestations . It is important to note that while p.Ala341Glu has been identified and studied, data specific to its prevalence and full clinical spectrum remain limited. Therefore, further investigation is warranted to fully elucidate the pathogenicity and clinical implications of the p.Ala341Glu variant. | <https://en.wikipedia.org/wiki/Long_QT_syndrome>  <https://www.malacards.org/card/long_qt_syndrome_1?search=KCNQ1>  <https://www.ncbi.nlm.nih.gov/clinvar/RCV000003268.2/>  <https://www.sciencedirect.com/science/article/pii/S0925443904001012>  <https://ojrd.biomedcentral.com/articles/10.1186/s13023-023-02618-4>  <https://pubmed.ncbi.nlm.nih.gov/17984373/>  <https://www.ncbi.nlm.nih.gov/gene/3784>  <https://academic.oup.com/europace/article/26/6/euae136/7686585?login=false>  <https://ojrd.biomedcentral.com/articles/10.1186/s13023-023-02618-4>  <https://www.ncbi.nlm.nih.gov/clinvar/RCV000003268.2/>  <https://academic.oup.com/eurheartj/article/42/46/4743/6368299?login=false>  <https://medlineplus.gov/genetics/gene/kcnq1/>  <https://pubmed.ncbi.nlm.nih.gov/17984373/>  <https://www.ahajournals.org/doi/10.1161/circulationaha.106.665406>  <https://www.ahajournals.org/doi/10.1161/01.CIR.100.10.1077>  <https://www.ncbi.nlm.nih.gov/clinvar/RCV000003267/>  <https://pubmed.ncbi.nlm.nih.gov/12702160/>  <https://pubmed.ncbi.nlm.nih.gov/16287990/>  <https://www.sciencedirect.com/science/article/pii/S0925443904001012>  <https://www.sciencedirect.com/science/article/abs/pii/S1547527112015391>  <https://www.sciencedirect.com/science/article/pii/S0972629218300755>  <https://www.sciencedirect.com/science/article/pii/S1098360022010607>  <https://www.sciencedirect.com/topics/medicine-and-dentistry/potassium-channel-kcnq1>  <https://omim.org/entry/607542>  <https://mgend.ncgm.go.jp/variant/info/SNV%7Chg38%7Cchr11%3A2583535-2583535%7CgC%3EA>  <https://www.ahajournals.org/doi/10.1161/01.cir.99.11.1464>  <https://go.drugbank.com/articles/A44083>  <https://web.expasy.org/variant_pages/VAR_001538.html>  <https://www.malacards.org/card/long_qt_syndrome_1?search=KCNQ1> |
|  | Evaluation  **Summarization Accuracy & Completeness:**  **Hallucination Rate:**  **Readability & Usability:**  **Literature Coverage & Relevance:**  **Time Efficiency:** | |
