## Supplementary material for "Benchmarking generative AI tools for literature retrieval and summarization in genomic variant interpretation": LMNA p.R482W.docx

| **Tool** | **Summary** | **References** |
| --- | --- | --- |
| **1** | The LMNA gene encodes lamin A and lamin C, key components of the nuclear lamina that maintain nuclear structure and regulate gene expression by interacting with chromatin and nuclear envelope proteins. The missense variant p.R482W, a substitution of arginine to tryptophan at codon 482 within exon 8, is among the most studied mutations in LMNA due to its strong association with familial partial lipodystrophy type 2 (FPLD2), also known as Dunnigan-type lipodystrophy. This pathogenic variant disrupts the C-terminal immunoglobulin-like fold domain of lamin A/C, impairing adipocyte differentiation and lipid metabolism, and is characterized by abnormal fat redistribution, insulin resistance, diabetes mellitus, and cardiovascular complications. Studies have identified p.R482W as the most common LMNA variant causing FPLD2, with consistent reports of variable expressivity and incomplete penetrance. Clinical heterogeneity is observed even among carriers within the same family, indicating that additional genetic or environmental modifiers may influence phenotypic outcomes [1]. In a systematic review of 115 LMNA-related lipodystrophy cases, p.R482W was the predominant variant and was strongly associated with metabolic complications and altered fat distribution patterns [2]. Research suggests that the R482 residue plays a critical role in DNA binding and chromatin organization; substitution with tryptophan likely disrupts these functions, promoting ectopic gene expression and adipose tissue abnormalities [3]. A separate study comparing R482 with other variants in the same exon confirmed that p.R482W carriers exhibit a more severe phenotype with earlier disease onset and greater metabolic impairment [4]. Although the variant is rare in the general population, it is enriched in familial lipodystrophy cohorts and is classified as pathogenic based on ACMG criteria due to its consistent association with disease, functional impact, and segregation in affected families. No significant population frequency is noted in control databases such as gnomAD. Despite extensive literature on the variant’s clinical manifestations, uncertainties remain regarding long-term outcomes and therapeutic response variability, underlining the need for individualized monitoring and further research into modifier genes or epigenetic factors that may alter disease expression. | [1] [Araujo-Vilar et al., 2021](https://www.mdpi.com/2077-0383/10/6/1259)  [2] [Besci & de Freitas, 2024](https://academic.oup.com/jcem/article/doi/10.1210/clinem/dgad606/7318161)  [3] [Florwick et al., 2017](https://www.frontiersin.org/articles/10.3389/fgene.2017.00079/full)  [4] [Decaudain et al., 2007](https://academic.oup.com/jcem/article-pdf/92/12/4835/9054956/jcem4835.pdf) |
|  | Evaluation  **Summarization Accuracy & Completeness:**  **Hallucination Rate:**  **Readability & Usability:**  **Literature Coverage & Relevance:**  **Time Efficiency:** | |
| **2** | The LMNA gene encodes A-type lamins, primarily lamin A and lamin C, which are intermediate filament proteins forming the nuclear lamina, a fibrous meshwork underlying the inner nuclear membrane. This structure is essential for maintaining nuclear shape, mechanical stability, and chromatin organization, thereby influencing gene expression and cellular mechanotransduction. The p.R482W variant in LMNA is a missense mutation involving the substitution of arginine by tryptophan at position 482, a residue located in the lamin A/C tail domain critical for protein-protein interactions and nuclear architecture. This variant is well-documented in the context of laminopathies, particularly familial partial lipodystrophy type 2 (FPLD2), characterized by abnormal adipose tissue distribution, metabolic disturbances, and increased risk of cardiovascular disease. The p.R482W mutation disrupts the normal function of lamin A/C, leading to altered nuclear mechanics, impaired chromatin organization, and dysregulated gene expression, which contribute to the multisystemic phenotype observed. Although the precise molecular mechanisms remain incompletely understood, evidence suggests that this variant affects the interaction of lamins with chromatin and other nuclear proteins, resulting in defective nuclear integrity and altered cellular responses to mechanical stress. The mutation is rare in the general population but recurrent in affected individuals, consistent with its pathogenic classification. Functional studies have demonstrated that p.R482W leads to nuclear shape abnormalities and altered transcriptional profiles in patient-derived cells. The variant’s impact extends to cardiomyopathy and muscular dystrophy phenotypes in some cases, reflecting the broad spectrum of laminopathies associated with LMNA mutations. No curative treatments currently exist; management is symptomatic and tailored to organ involvement. Given the complexity and variability of phenotypes linked to p.R482W, further research is warranted to elucidate detailed pathogenic pathways and to develop targeted therapies. Peer-reviewed publications specifically addressing p.R482W include studies on its role in lipodystrophy and nuclear envelope dysfunction, but direct references focusing exclusively on this variant’s mechanistic effects remain limited. Therefore, clinical interpretation should consider the variant in the context of established laminopathy phenotypes and recommend genetic counseling and multidisciplinary follow-up for affected individuals. Further functional assays and genotype-phenotype correlation studies are needed to refine prognostic assessments. No direct citations exclusively on p.R482W were found in the recent literature; however, the general pathogenic role of LMNA mutations in nuclear structure and disease phenotypes is well-supported[1][2][3]. | [1] <https://pubmed.ncbi.nlm.nih.gov/37125775/>  [2] <https://www.ncbi.nlm.nih.gov/pmc/articles/PMC10976794/>  [3] <https://www.ncbi.nlm.nih.gov/pmc/articles/PMC11818007/>  [4] <https://www.semanticscholar.org/paper/127cef6487a7d1d52fc54950c7d4557ad13c22d5>  [5] <https://www.ncbi.nlm.nih.gov/pmc/articles/PMC6413092/>  [6] <https://www.semanticscholar.org/paper/10eb2e96f88c44a5213bb876955d13255fb51de5>  [7] <https://www.ncbi.nlm.nih.gov/pmc/articles/PMC9778678/>  [8] <https://www.semanticscholar.org/paper/6eff96c5b33b1fd18b969ab18137992e335a5be4> |
|  | Evaluation  **Summarization Accuracy & Completeness:**  **Hallucination Rate:**  **Readability & Usability:**  **Literature Coverage & Relevance:**  **Time Efficiency:** | |
| **3** | The LMNA gene encodes lamin A and lamin C, structural proteins of the nuclear lamina that provide mechanical support to the nucleus and are involved in chromatin organization and gene regulation. The p.R482W variant, a missense mutation substituting arginine with tryptophan at codon 482, is associated with familial partial lipodystrophy type 2 (FPLD2), also known as Dunnigan-type lipodystrophy. This autosomal dominant condition is characterized by postpubertal loss of subcutaneous fat from the limbs and trunk, with fat accumulation in the face and neck, insulin resistance, dyslipidemia, and increased cardiovascular risk. The p.R482W mutation impairs the interaction between lamin A and transcriptional regulators such as SREBP1, leading to dysregulation of adipogenic gene expression and contributing to the lipodystrophic phenotype . Additionally, this variant disrupts nuclear architecture and chromatin organization, affecting gene networks involved in mesodermal and vascular differentiation . Clinically, individuals with the p.R482W mutation exhibit early-onset atherosclerosis, endothelial dysfunction, and, in some cases, muscular and cardiac abnormalities, including limb-girdle muscular dystrophy and cardiac conduction defects . These findings underscore the multisystem impact of the p.R482W variant and highlight the need for comprehensive clinical evaluation and management of affected individuals. | <https://www.ncbi.nlm.nih.gov/gene/4000>  <https://www.sciencedirect.com/science/article/pii/S2090123225000013> ù  <https://www.frontiersin.org/journals/genetics/articles/10.3389/fgene.2024.1468878/full>  <https://www.science.org/doi/10.1126/scitranslmed.aax9276>  <https://www.frontiersin.org/journals/physiology/articles/10.3389/fphys.2020.00761/full>  <https://medlineplus.gov/download/genetics/gene/lmna.pdf>  <https://pubmed.ncbi.nlm.nih.gov/23846499/>  <https://www.ahajournals.org/doi/full/10.1161/atvbaha.113.301933>  <https://www.ahajournals.org/doi/abs/10.1161/atvbaha.113.301933>  <https://www.ncbi.nlm.nih.gov/books/NBK6151/>  <https://pubmed.ncbi.nlm.nih.gov/15531479/>  <https://academic.oup.com/hmg/article/24/7/2096/598363>  <https://academic.oup.com/hmg/article/27/8/1447/4847145>  <https://www.genecards.org/cgi-bin/carddisp.pl?gene=LMNA>  <https://www.frontiersin.org/journals/pharmacology/articles/10.3389/fphar.2022.820857/full>  <https://omim.org/entry/150330>  <https://en.wikipedia.org/wiki/Prelamin-A/C>  <https://www.sciencedirect.com/topics/biochemistry-genetics-and-molecular-biology/lmna> |
|  | Evaluation  **Summarization Accuracy & Completeness:**  **Hallucination Rate:**  **Readability & Usability:**  **Literature Coverage & Relevance:**  **Time Efficiency:** | |
| **4** | The genomic variant c.1444C>T p.R482W rs57920071 in the LMNA gene is a well-documented mutation associated with familial partial lipodystrophy of Dunnigan type (FPLD2), an autosomal dominant disorder characterized by the atrophy of adipose tissue in the extremities, visceral and cervical fat accumulation, muscle hypertrophy, and severe metabolic complications including insulin resistance, diabetes, and dyslipidemia [2]. This variant has been identified as a hotspot mutation within the LMNA gene and is one of the most frequent mutations underlying lipodystrophic laminopathies [2].  The LMNA gene encodes A-type lamins, which are intermediate filament proteins that form a meshwork underlying the inner nuclear membrane. A-type lamins, including lamins A/C, play a critical role in maintaining nuclear structure, chromatin organization, and the regulation of gene expression [3]. Mutations in LMNA can lead to a variety of diseases collectively known as laminopathies, which include muscle dystrophies, cardiomyopathies, peripheral neuropathies, and premature aging syndromes, in addition to lipodystrophies [3].  The p.R482W variant specifically affects the immunoglobulin fold of lamin A, which is involved in interactions with DNA and nucleosomes. Although the R482 residue is located on the surface of the fold and does not severely disrupt the structure, the mutation impairs the interaction of the immunoglobulin fold with DNA and nucleosomes in vitro, potentially perturbing associations of lamin A with chromatin and spatial genome conformation [2].  In cellular models, the R482W mutation has been shown to lead to deficiencies in adipogenesis and mesodermal and endothelial differentiation. It also results in defects in nuclear morphology, adipogenic transcription factor compartmentalization, and signal transduction [2]. Furthermore, the mutation has been linked to early-onset atherosclerosis and cardiovascular pathologies, which are not solely explained by metabolic risk factors. In endothelial cells, the p.R482W-prelamin-A accumulates abnormally at the nuclear envelope, leading to endothelial dysfunction, oxidative stress, DNA damage, and inflammation [4].  The R482W variant has been associated with a direct proatherogenic effect in endothelial cells, contributing to early atherosclerosis in patients. This effect is suggested to be due to the accumulation of farnesylated p.R482W-prelamin-A at the nuclear envelope, which is a toxic event leading to cellular oxidative stress and endothelial dysfunction [4].  Additionally, the R482W mutation has been implicated in the deregulation of signaling pathways, nucleus and cell mechanosensitivity, and nuclear architecture. It leads to adipogenic differentiation defects and impairs the interaction of LMNA with the adipogenic factor SREBP1 and with DNA in vitro [5]. The mutation also affects the regulation of miR-335, which is involved in adipogenic gene expression, suggesting a complex interplay between the mutant lamin A and epigenetic as well as chromatin state regulation [5].  In summary, the c.1444C>T p.R482W rs57920071 variant in the LMNA gene is a pathogenic mutation that causes FPLD2 through a complex mechanism involving impaired protein-DNA interactions, defects in adipogenic differentiation, and endothelial dysfunction. The mutation has a significant impact on nuclear architecture and gene expression regulation, which contributes to the diverse clinical manifestations observed in patients with this variant. | [[1]](https://pubmed.ncbi.nlm.nih.gov/39550450/) Rajan R et al. (2024). "A series of genetically confirmed congenital lipodystrophy and diabetes in adult southern Indian patients." Scientific reports, 14(1)  [[2]](https://pubmed.ncbi.nlm.nih.gov/30057899/) Briand N et al. (2018). "Lamin A, Chromatin and FPLD2: Not Just a Peripheral Menage-a-Trois." Frontiers in cell and developmental biology, 6()  [[3]](https://pubmed.ncbi.nlm.nih.gov/37190051/) Walker SG et al. (2023). "Drosophila Models Reveal Properties of Mutant Lamins That Give Rise to Distinct Diseases." Cells, 12(8)  [[4]](https://pubmed.ncbi.nlm.nih.gov/23846499/) Bidault G et al. (2013). "Lipodystrophy-linked LMNA p.R482W mutation induces clinical early atherosclerosis and in vitro endothelial dysfunction." Arteriosclerosis, thrombosis, and vascular biology, 33(9)  [[5]](https://pubmed.ncbi.nlm.nih.gov/28751304/) Oldenburg A et al. (2017). "A lipodystrophy-causing lamin A mutant alters conformation and epigenetic regulation of the anti-adipogenic MIR335 locus." The Journal of cell biology, 216(9)  [[6]](https://pubmed.ncbi.nlm.nih.gov/29438482/) Briand N et al. (2018). "The lipodystrophic hotspot lamin A p.R482W mutation deregulates the mesodermal inducer T/Brachyury and early vascular differentiation gene networks." Human molecular genetics, 27(8)  [[7]](https://pubmed.ncbi.nlm.nih.gov/37701573/) Jebane C et al. (2023). "Enhanced cell viscosity: A new phenotype associated with lamin A/C alterations." iScience, 26(10)  [[8]](https://pubmed.ncbi.nlm.nih.gov/28811278/) Elzeneini E et al. (2017). "Lipodystrophic laminopathy: Lamin A mutation relaxes chromatin architecture to impair adipogenesis." The Journal of cell biology, 216(9)  [[9]](https://pubmed.ncbi.nlm.nih.gov/38739524/) Morguetti MJ et al. (2024). "Podocytopathies associated with familial partial lipodystrophy due to LMNA variants: report of two cases." Archives of endocrinology and metabolism, 68()  [[10]](https://pubmed.ncbi.nlm.nih.gov/24108105/) Oldenburg AR et al. (2014). "Deregulation of Fragile X-related protein 1 by the lipodystrophic lamin A p.R482W mutation elicits a myogenic gene expression program in preadipocytes." Human molecular genetics, 23(5)  [[11]](https://pubmed.ncbi.nlm.nih.gov/25524705/) Vadrot N et al. (2015). "The p.R482W substitution in A-type lamins deregulates SREBP1 activity in Dunnigan-type familial partial lipodystrophy." Human molecular genetics, 24(7)  [[12]](https://pubmed.ncbi.nlm.nih.gov/26756202/) Demir T et al. (2016). "Familial partial lipodystrophy linked to a novel peroxisome proliferator activator receptor -gamma (PPARG) mutation, H449L: a comparison of people with this mutation and those with classic codon 482 Lamin A/C (LMNA) mutations." Diabetic medicine : a journal of the British Diabetic Association, 33(10)  [[13]](https://pubmed.ncbi.nlm.nih.gov/27504462/) Panikkath R et al. (2016). "An Uncommon Association of Familial Partial Lipodystrophy, Dilated Cardiomyopathy, and Conduction System Disease." Journal of investigative medicine high impact case reports, 4(3)  [[14]](https://pubmed.ncbi.nlm.nih.gov/11344241/) Schmidt HH et al. (2001). "Dyslipemia in familial partial lipodystrophy caused by an R482W mutation in the LMNA gene." The Journal of clinical endocrinology and metabolism, 86(5)  [[15]](https://pubmed.ncbi.nlm.nih.gov/11792811/) Vigouroux C et al. (2001). "Nuclear envelope disorganization in fibroblasts from lipodystrophic patients with heterozygous R482Q/W mutations in the lamin A/C gene." Journal of cell science, 114(Pt 24) |
|  | Evaluation  **Summarization Accuracy & Completeness:**  **Hallucination Rate:**  **Readability & Usability:**  **Literature Coverage & Relevance:**  **Time Efficiency:** | |
| **5** | The LMNA gene, located on chromosome 1q21, encodes nuclear lamins A and C, which are crucial structural proteins of the nuclear envelope. These proteins play essential roles in maintaining nuclear integrity, chromatin organization, and gene regulation. Mutations in LMNA are associated with a variety of disorders collectively known as laminopathies, which include muscular dystrophies, cardiomyopathies, neuropathies, premature aging syndromes, and lipodystrophies. One notable variant of LMNA is the p.R482W missense mutation, which is strongly linked to familial partial lipodystrophy type 2 (FPLD2), also known as Dunnigan-type lipodystrophy. FPLD2 is an autosomal dominant condition characterized by abnormal fat distribution, with a loss of subcutaneous adipose tissue in the limbs and trunk, and an accumulation of fat in the face, neck, and intra-abdominal regions. This fat redistribution leads to metabolic complications such as insulin resistance, diabetes mellitus, hypertriglyceridemia, and early-onset atherosclerosis. The p.R482W mutation affects the C-terminal globular tail of lamin A, disrupting its interactions with other proteins and altering nuclear envelope structure and function. Studies have shown that this mutation impairs adipocyte differentiation and function, leading to increased lipolysis and altered insulin signaling. Additionally, the p.R482W mutation has been found to deregulate the activity of SREBP1, a transcription factor involved in lipid metabolism and adipocyte differentiation, further contributing to the pathophysiology of FPLD2. The frequency of the p.R482W variant is not well-documented in the general population, but it is considered the most common mutation associated with FPLD2. The classification of this variant is based on its pathogenic effects on adipose tissue and metabolic function, as observed in clinical and molecular studies. While the exact prevalence and impact of the p.R482W mutation may vary, it is clear that it plays a significant role in the development of FPLD2 and its associated metabolic disorders. Further research is needed to fully understand the molecular mechanisms underlying this mutation and to develop targeted therapies for affected individuals. | [1] FPLD2 LMNA mutation R482W dysregulates iPSC-derived adipocyte function and lipid metabolism - ScienceDirect  [2] LMNA p.R482W mutation related to FPLD2 alters SREBP1-A type lamin interactions in human fibroblasts and adipose stem cells \| Orphanet Journal of Rare Diseases \| Full Text  [3] p.R482W substitution in A-type lamins deregulates SREBP1 activity in Dunnigan-type familial partial lipodystrophy \| Human Molecular Genetics \| Oxford Academic  [4] Structure of the lamin A/C R482W mutant responsible for dominant familial partial lipodystrophy (FPLD) - PMC  [5] LMNA gene: MedlinePlus Genetics |
|  | Evaluation  **Summarization Accuracy & Completeness:**  **Hallucination Rate:**  **Readability & Usability:**  **Literature Coverage & Relevance:**  **Time Efficiency:** | |
