## Supplementary material for "Benchmarking generative AI tools for literature retrieval and summarization in genomic variant interpretation": MTHFR c.C677T.docx

| **Tool** | **Summary** | **References** |
| --- | --- | --- |
| **1** | The genomic variant c.C677T in the MTHFR gene results in a substitution of cytosine for thymine at nucleotide position 677. This mutation leads to the production of a thermolabile variant of the methylenetetrahydrofolate reductase enzyme with reduced activity [1]. The MTHFR gene plays a crucial role in folate metabolism, where it catalyzes the conversion of 5,10-methylenetetrahydrofolate to 5-methyltetrahydrofolate, a substrate for the remethylation of homocysteine to methionine [2]. This process is essential for DNA methylation and synthesis, and disruptions can have various clinical implications.  Individuals who are homozygous for the c.C677T variant (approximately 10% of North Americans) have an increased predisposition to mild hyperhomocysteinemia, particularly when folate status is low [1]. This condition has been associated with an increased risk for neural tube defects and vascular disease [1]. Moreover, the presence of the c.C677T variant has been linked to altered distribution of red blood cell folates, with formylated tetrahydrofolate polyglutamates being present in addition to methylated derivatives in homozygous individuals [3]. This altered folate distribution is consistent with an in vivo impairment of the thermolabile MTHFR variant's activity.  The c.C677T variant's impact on health is influenced by folate status, as low folate levels in individuals with the homozygous mutation are associated with diminished DNA methylation [4]. This interaction between folate status and the c.C677T variant is further evidenced by the fact that individuals with the homozygous mutation and low folate levels have a significantly reduced level of genomic DNA methylation compared to those with the wild-type genotype [4].  The c.C677T variant has also been implicated in psychiatric disorders, with meta-analyses demonstrating associations between the homozygous variant and increased risks for unipolar depression, schizophrenia, and bipolar disorder [5]. These findings suggest potential implications for the use of folate in the treatment and prevention of these conditions.  In the context of cardiovascular disease, while the c.C677T variant is a major cause of mild hyperhomocysteinemia, meta-analyses have not found it to increase cardiovascular risk, suggesting that the mild hyperhomocysteinemia often observed in vascular disease patients may not be causally related to the pathogenesis of the disease [6].  The c.C677T variant has also been studied in relation to colorectal neoplasia, with findings indicating that individuals with the homozygous variant may have a moderately reduced risk of colorectal cancer, particularly when folate levels are higher or when alcohol intake is lower [7]. Additionally, the variant has been associated with an increased risk of neural tube defects, reinforcing the importance of folate intake for individuals carrying the mutation [8].  In pediatric leukemias, the c.C677T variant has been associated with a protective effect against leukemias with MLL translocations, suggesting a complex role of folate and one-carbon metabolism in the development of childhood leukemia [9]. Lastly, the variant has been identified as a risk factor for cerebral small vessel disease, particularly ischaemic leukoaraiosis, with the association potentially mediated by endothelial dysfunction [10].  In summary, the c.C677T variant in the MTHFR gene has significant implications for folate metabolism and is associated with a range of health outcomes, including neural tube defects, psychiatric disorders, and certain cancers. Its effects are modulated by folate status, highlighting the importance of nutritional factors in the clinical expression of genetic variants. | [[1]](https://pubmed.ncbi.nlm.nih.gov/9719624/) Ilan S. Weisberg et al. (1998). "A second genetic polymorphism in methylenetetrahydrofolate reductase (MTHFR) associated with decreased enzyme activity." Molecular genetics and metabolism, 64 3  [[2]](https://pubmed.ncbi.nlm.nih.gov/10791559/) L. Botto et al. (2000). "5,10-Methylenetetrahydrofolate reductase gene variants and congenital anomalies: a HuGE review." American journal of epidemiology, 151 9  [[3]](https://pubmed.ncbi.nlm.nih.gov/9789068/) P. Bagley et al. (1998). "A common mutation in the methylenetetrahydrofolate reductase gene is associated with an accumulation of formylated tetrahydrofolates in red blood cells." Proceedings of the National Academy of Sciences of the United States of America, 95 22  [[4]](https://pubmed.ncbi.nlm.nih.gov/11929966/) S. Friso et al. (2002). "A common mutation in the 5,10-methylenetetrahydrofolate reductase gene affects genomic DNA methylation through an interaction with folate status" Proceedings of the National Academy of Sciences of the United States of America, 99  [[5]](https://pubmed.ncbi.nlm.nih.gov/17074966/) S. Gilbody et al. (2006). "Methylenetetrahydrofolate reductase (MTHFR) genetic polymorphisms and psychiatric disorders: a HuGE review." American journal of epidemiology, 165 1  [[6]](https://pubmed.ncbi.nlm.nih.gov/9843457/) L. Brattström et al. (1998). "Common methylenetetrahydrofolate reductase gene mutation leads to hyperhomocysteinemia but not to vascular disease: the result of a meta-analysis." Circulation, 98 23  [[7]](https://pubmed.ncbi.nlm.nih.gov/14977639/) L. Sharp et al. (2004). "Polymorphisms in genes involved in folate metabolism and colorectal neoplasia: a HuGE review." American journal of epidemiology, 159 5  [[8]](https://pubmed.ncbi.nlm.nih.gov/11368417/) N. M. van der Put et al. (2001). "Folate, Homocysteine and Neural Tube Defects: An Overview" Experimental Biology and Medicine, 226  [[9]](https://pubmed.ncbi.nlm.nih.gov/11274424/) J. Wiemels et al. (2001). "Methylenetetrahydrofolate reductase (MTHFR) polymorphisms and risk of molecularly defined subtypes of childhood acute leukemia" Proceedings of the National Academy of Sciences of the United States of America, 98  [[10]](https://pubmed.ncbi.nlm.nih.gov/14607791/) Ahamad Hassan et al. (2004). "Homocysteine is a risk factor for cerebral small vessel disease, acting via endothelial dysfunction." Brain : a journal of neurology, 127 Pt 1  [[11]](https://pubmed.ncbi.nlm.nih.gov/25771069/) Y. Huo et al. (2015). "Efficacy of folic acid therapy in primary prevention of stroke among adults with hypertension in China: the CSPPT randomized clinical trial." JAMA, 313 13  [[12]](https://pubmed.ncbi.nlm.nih.gov/18505952/) Linda M. Dong et al. (2008). "Genetic susceptibility to cancer: the role of polymorphisms in candidate genes." JAMA, 299 20  [[13]](https://pubmed.ncbi.nlm.nih.gov/20385928/) G. Kenet et al. (2010). "Impact of Thrombophilia on Risk of Arterial Ischemic Stroke or Cerebral Sinovenous Thrombosis in Neonates and Children: A Systematic Review and Meta-Analysis of Observational Studies" Circulation, 121  [[14]](https://pubmed.ncbi.nlm.nih.gov/15671130/) A. Coppen et al. (2005). "Treatment of depression: time to consider folic acid and vitamin B12" Journal of Psychopharmacology, 19  [[15]](https://pubmed.ncbi.nlm.nih.gov/10835445/) G. Kenet et al. (2000). "Factor V Leiden and antiphospholipid antibodies are significant risk factors for ischemic stroke in children." Stroke, 31 6 |
|  | Evaluation  **Summarization Accuracy & Completeness:**  **Hallucination Rate:**  **Readability & Usability:**  **Literature Coverage & Relevance:**  **Time Efficiency:** | |
| **2** | The methylenetetrahydrofolate reductase (MTHFR) gene encodes an enzyme critical for folate metabolism and homocysteine regulation, playing a vital role in DNA synthesis, repair, and methylation. The c.C677T variant (rs1801133), a common single nucleotide polymorphism (SNP), results in an alanine-to-valine substitution at position 222, leading to a thermolabile enzyme with reduced activity. This polymorphism is associated with increased plasma homocysteine levels, particularly in individuals with low folate intake, which has been implicated in cardiovascular disease, neural tube defects, and certain cancers. Homozygous individuals (TT genotype) exhibit a 30-40% reduction in enzyme activity compared to the CC genotype, while heterozygous individuals (CT) show an intermediate reduction. Epidemiological studies suggest an association between MTHFR C677T and an increased risk of ischemic stroke and coronary artery disease, particularly in populations with inadequate folate levels [1]. Additionally, meta-analyses indicate that the variant contributes to colorectal cancer susceptibility under a recessive model, with a modest impact on disease risk [2]. Studies also report a potential influence on neurological conditions, including migraine and epilepsy, though findings remain inconclusive [3]. The global frequency of the T allele varies significantly, with higher prevalence in East Asian and Hispanic populations (30-40%) and lower frequencies in African populations (<10%). Despite these associations, the clinical utility of MTHFR genetic testing remains controversial, as many of the proposed disease links are modified by environmental factors, including dietary folate intake and lifestyle [4]. Given these complexities, while the MTHFR C677T variant may contribute to disease risk in specific populations, current evidence does not support routine genetic testing for clinical decision-making. Further research is required to clarify its role in complex diseases and potential gene-environment interactions. | [1] Cortese & Motti (2001),  [2] Hubner & Houlston (2007),  [3] Liew & Gupta (2015),  [4] Long & Goldblatt (2016). |
|  | Evaluation  **Summarization Accuracy & Completeness:**  **Hallucination Rate:**  **Readability & Usability:**  **Literature Coverage & Relevance:**  **Time Efficiency:** | |
| **3** | Methylenetetrahydrofolate reductase (MTHFR) is an enzyme that plays a crucial role in the folate metabolism pathway, specifically catalyzing the conversion of 5,10-methylenetetrahydrofolate to 5-methyltetrahydrofolate, which is essential for the remethylation of homocysteine to methionine. The c.C677T variant in the MTHFR gene results in an alanine-to-valine substitution at codon 222, leading to reduced enzyme activity and increased thermolability. This polymorphism exhibits varying frequencies across populations; for instance, approximately 10% of the North American population are homozygous for the T allele, with higher prevalence observed in Mediterranean and Hispanic populations compared to Caucasians and African-Americans. Individuals homozygous for the 677T allele (TT genotype) exhibit significantly reduced MTHFR activity, approximately 18–22% of normal, compared to heterozygous individuals (CT genotype) with about 56% activity, and those with the CC genotype maintaining 66–67% activity. This diminished enzyme function in TT individuals is associated with mild hyperhomocysteinemia, particularly under conditions of low dietary folate intake, as they have less active MTHFR available to produce 5-methyltetrahydrofolate, which is used to decrease homocysteine. Elevated homocysteine levels have been linked to various health risks, including neural tube defects, cardiovascular diseases, and certain cancers. However, it is important to note that while the MTHFR C677T polymorphism can influence homocysteine concentrations, the clinical significance of this variant as an independent risk factor for these conditions remains a subject of ongoing research and debate. The American College of Medical Genetics and Genomics (ACMG) currently recommends against routine testing for MTHFR variants, citing a lack of sufficient evidence to support a direct causal relationship between this polymorphism and specific health outcomes. Therefore, the presence of the MTHFR C677T variant should be interpreted with caution, and clinical decisions should not be based solely on MTHFR genotyping without considering other established risk factors and clinical context. | <https://en.wikipedia.org/wiki/Methylenetetrahydrofolate_reductase>  <https://bmccancer.biomedcentral.com/articles/10.1186/s12885-025-13546-w>  <https://bmcwomenshealth.biomedcentral.com/articles/10.1186/s12905-024-03244-3>  <https://www.cdc.gov/folic-acid/data-research/mthfr/index.html>  <https://www.mdpi.com/2218-1989/13/2/251>  <https://onlinelibrary.wiley.com/doi/10.1111/jhn.13061>  <https://pmc.ncbi.nlm.nih.gov/articles/PMC9775363/>  <https://www.ahajournals.org/doi/10.1161/circulationaha.114.013311>  <https://www.nature.com/articles/5201914>  <https://bmcmedicine.biomedcentral.com/articles/10.1186/1741-7015-2-3>  <https://pmc.ncbi.nlm.nih.gov/articles/PMC375543/>  <https://pmc.ncbi.nlm.nih.gov/articles/PMC9433753/>  <https://www.frontiersin.org/journals/genetics/articles/10.3389/fgene.2023.1299599/full>  <https://www.frontiersin.org/journals/psychiatry/articles/10.3389/fpsyt.2021.560948/full>  <https://www.aging-us.com/article/202827/text>  <https://www.racgp.org.au/afp/2016/april/mthfr-genetic-testing-controversy-and-clinical-imp>  <https://www.scirp.org/journal/paperinformation?paperid=125731> |
|  | Evaluation  **Summarization Accuracy & Completeness:**  **Hallucination Rate:**  **Readability & Usability:**  **Literature Coverage & Relevance:**  **Time Efficiency:** | |
| **4** | The MTHFR gene encodes methylenetetrahydrofolate reductase, a critical enzyme in folate metabolism that catalyzes the conversion of 5,10-methylenetetrahydrofolate to 5-methyltetrahydrofolate, essential for homocysteine remethylation to methionine. This process is vital for DNA synthesis, repair, and methylation, which regulate gene expression and cellular functions. The MTHFR gene is located on chromosome 1 at position 1p36.3, and its proper functioning is crucial for maintaining normal homocysteine levels and supporting various biological processes, including neurotransmitter production and amino acid metabolism. The c.C677T variant, specifically, involves a substitution of cytosine (C) with thymine (T) at nucleotide position 677, leading to an alanine-to-valine change at amino acid position 222. This variant is classified as a single nucleotide polymorphism (SNP) and is one of the most common genetic variants in the MTHFR gene, with a higher prevalence in certain ethnic groups, such as Hispanics. The c.C677T variant results in a thermolabile enzyme with reduced activity, particularly under conditions of low folate intake, leading to elevated homocysteine levels. This hyperhomocysteinemia is associated with an increased risk of various health conditions, including cardiovascular diseases, neural tube defects, and certain types of cancer. Individuals homozygous for the c.C677T variant (TT genotype) exhibit significantly reduced MTHFR enzyme activity, approximately 30% of normal, compared to 65% in heterozygous individuals (CT genotype). The frequency of the c.C677T variant varies by geography and ethnicity, with higher incidences in North Indian patients with vascular disease and among Hispanic populations. Research has linked the c.C677T variant to increased susceptibility to migraine with aura, vascular diseases, and adverse pregnancy outcomes, including recurrent miscarriages and fetal developmental defects. However, the clinical significance of this variant remains a subject of ongoing research, as some studies have not consistently reproduced these associations. Current guidelines do not recommend routine testing for the c.C677T variant in individuals with recurrent pregnancy loss or for determining the risk of cardiovascular diseases, as the utility of such testing is limited. Management strategies for individuals with the c.C677T variant often include dietary and lifestyle modifications to optimize folate intake and reduce homocysteine levels. Further investigation is needed to fully understand the impact of the c.C677T variant on health outcomes and to develop targeted interventions for affected individuals. | [1] Methylenetetrahydrofolate reductase (MTHFR) C677T polymorphism: epidemiology, metabolism and the associated diseases - PubMed.  [2] MTHFR Gene Variant and Folic Acid Facts \| Folic Acid \| CDC.  [3] The methylenetetrahydrofolate reductase gene variant C677T influences susceptibility to migraine with aura - PMC. |
|  | Evaluation  **Summarization Accuracy & Completeness:**  **Hallucination Rate:**  **Readability & Usability:**  **Literature Coverage & Relevance:**  **Time Efficiency:** | |
| **5** | The Methylenetetrahydrofolate Reductase (MTHFR) gene provides instructions for making an enzyme that is vital in processing folate (vitamin B9)[1](https://www.semanticscholar.org/paper/21c15ee5f719a3b758c1a4b749ba627af921dd09)[5](https://pubmed.ncbi.nlm.nih.gov/19604253/). MTHFR converts 5,10-methyltetrahydrofolate to 5-methyltetrahydrofolate, which is used to convert homocysteine to methionine. High homocysteine levels are associated with various health problems[1](https://www.semanticscholar.org/paper/21c15ee5f719a3b758c1a4b749ba627af921dd09)[5](https://pubmed.ncbi.nlm.nih.gov/19604253/). The MTHFR c.C677T variant (also known as A222V) is a single nucleotide polymorphism (SNP) in the MTHFR gene[1](https://www.semanticscholar.org/paper/21c15ee5f719a3b758c1a4b749ba627af921dd09)[7](https://www.ncbi.nlm.nih.gov/pmc/articles/PMC9168012/). The MTHFR C677T polymorphism is clinically important, and the variant MTHFR (A222V) enzyme has reduced activity, hence increasing the requirement for folic acid[1](https://www.semanticscholar.org/paper/21c15ee5f719a3b758c1a4b749ba627af921dd09). Individuals with the C677T polymorphism may have higher homocysteine levels, especially with low folate intake, which can impact DNA synthesis, repair, and methylation[1](https://www.semanticscholar.org/paper/21c15ee5f719a3b758c1a4b749ba627af921dd09)[5](https://pubmed.ncbi.nlm.nih.gov/19604253/). Studies have explored the association of MTHFR C677T with various conditions. Some research suggests a potential link between the MTHFR C677T variant and male infertility, as well as an increased risk of severe erectile dysfunction[2](https://www.ncbi.nlm.nih.gov/pmc/articles/PMC11426628/)[3](https://www.ncbi.nlm.nih.gov/pmc/articles/PMC10846823/). One study found a significantly higher frequency of the 677TT polymorphism in severe and moderate ED (P = 0.02)[3](https://www.ncbi.nlm.nih.gov/pmc/articles/PMC10846823/). However, other studies did not find a significant association between the C677T polymorphism and male infertility[3](https://www.ncbi.nlm.nih.gov/pmc/articles/PMC10846823/)[4](https://www.semanticscholar.org/paper/685e1ee204643b17427359e6ed83c26c4a56d533). In a Bahraini population, the mutant genotypes (CT and TT) of the MTHFR did not differ between patients and control subjects (OR=0.49; 95% CI 0.26–0.92), indicating that this mutation is unlikely to play a major role in stroke predisposition in the studied sample[8](https://www.semanticscholar.org/paper/82636c759ba962144efae1601201ffaa2f07ccc0). The prevalence of the MTHFR C677T polymorphism varies across different populations[2](https://www.ncbi.nlm.nih.gov/pmc/articles/PMC11426628/). Further research is needed to fully elucidate the clinical significance of this variant. | [1](https://www.semanticscholar.org/paper/21c15ee5f719a3b758c1a4b749ba627af921dd09) Detrimental Effects of Methylenetetrahydrofolate Reductase (MTHFR) Gene Polymorphism on Human Reproductive Health: A Review.  [2](https://www.ncbi.nlm.nih.gov/pmc/articles/PMC11426628/) Correlation of MTHFR C677T Polymorphism with Male Infertility among Indian Population: Case-Control Study.  [3](https://www.ncbi.nlm.nih.gov/pmc/articles/PMC10846823/) Association between MTHFR c.677C>T variant and erectile dysfunction among males attending fertility clinic.  [4](https://www.semanticscholar.org/paper/685e1ee204643b17427359e6ed83c26c4a56d533) Role of MTHFR Gene Polymorphisms in Male Infertility.  [5](https://pubmed.ncbi.nlm.nih.gov/19604253/) Comment on Rubino et al., ‘Association of the C677T Polymorphism in the MTHFR Gene with Migraine: A Meta-Analysis’.  [6](https://www.ncbi.nlm.nih.gov/pmc/articles/PMC7463445/) MTHFR Gene Polymorphisms Prevalence and Cardiovascular Risk Factors Involved in Cardioembolic Stroke Type and Severity.  [7](https://www.ncbi.nlm.nih.gov/pmc/articles/PMC9168012/) Genetic polymorphism in Methylenetetrahydrofolate Reductase chloride transport protein 6 (MTHFR CLCN6) gene is associated with keratinocyte skin cancer in a cohort of renal transplant recipients.  [8](https://www.semanticscholar.org/paper/82636c759ba962144efae1601201ffaa2f07ccc0) Association Between Methylenetetrahy drofolate Reductase (MTHFR) Gene (C677T) Polymorphism and Stroke predisposition in Bahraini Population.   1. <https://www.semanticscholar.org/paper/21c15ee5f719a3b758c1a4b749ba627af921dd09> 2. <https://www.ncbi.nlm.nih.gov/pmc/articles/PMC11426628/> 3. <https://www.ncbi.nlm.nih.gov/pmc/articles/PMC10846823/> 4. <https://www.semanticscholar.org/paper/685e1ee204643b17427359e6ed83c26c4a56d533> 5. <https://pubmed.ncbi.nlm.nih.gov/19604253/> 6. <https://www.ncbi.nlm.nih.gov/pmc/articles/PMC7463445/> 7. <https://www.ncbi.nlm.nih.gov/pmc/articles/PMC9168012/> 8. <https://www.semanticscholar.org/paper/82636c759ba962144efae1601201ffaa2f07ccc0> |
|  | Evaluation  **Summarization Accuracy & Completeness:**  **Hallucination Rate:**  **Readability & Usability:**  **Literature Coverage & Relevance:**  **Time Efficiency:** | |
