## Supplementary material for "Benchmarking generative AI tools for literature retrieval and summarization in genomic variant interpretation": PAH p.R408R.docx

| **Tool** | **Summary** | **References** |
| --- | --- | --- |
| **1** | The PAH gene encodes phenylalanine hydroxylase (PAH), an enzyme crucial for converting phenylalanine to tyrosine, a process vital for neurotransmitter production and overall metabolism. The PAH enzyme operates within the biopterin-dependent aromatic amino acid hydroxylase family and is central to phenylalanine catabolism. Deficiencies in PAH activity lead to phenylketonuria (PKU), an autosomal recessive disorder characterized by the accumulation of phenylalanine, which can cause severe brain damage and other medical issues if untreated.  The variant p.R408R in the PAH gene is a synonymous mutation, meaning it does not alter the amino acid sequence of the resulting protein. Synonymous variants are generally considered benign as they do not directly affect protein structure or function. However, they can sometimes influence gene expression or splicing, although such effects are not typically associated with p.R408R. This variant has not been extensively studied in the literature, and its frequency in the population is not well-documented. The lack of direct references to p.R408R in academic research suggests that it is not a common or well-studied variant in the context of PAH-related disorders.  In contrast, the variant p.R408W (Arg408Trp) is one of the most common and well-studied pathogenic variants in the PAH gene. It is prevalent in Eastern European populations and is associated with a severe form of PKU. This variant results in a significant reduction in PAH enzyme activity, leading to the accumulation of phenylalanine and the characteristic symptoms of PKU. The frequency of p.R408W in control populations is low, but it is significantly higher in affected individuals, particularly those of European descent. Protein expression studies have shown that p.R408W causes severe protein aggregation and loss of function, further supporting its pathogenic classification.  Given the limited information on p.R408R, it is essential to approach its interpretation with caution. While it is likely benign due to its synonymous nature, further investigation may be warranted to rule out any potential effects on gene expression or splicing. In the absence of direct evidence, the impact of p.R408R remains uncertain, and it should not be assumed to contribute to PKU or other PAH-related disorders without additional data. | [1] DiLella AG et al. (1987). Hum Genet. 1990 Apr;84(5):409-11.  [2] Zschocke J. (2003).  [3] ZurflÃ¼h MR et al. (2008).  [4] Gersting SW et al. (2008). |
|  | Evaluation  **Summarization Accuracy & Completeness:**  **Hallucination Rate:**  **Readability & Usability:**  **Literature Coverage & Relevance:**  **Time Efficiency:** | |
| **2** | The p.R408R variant in the PAH gene, also known by its cDNA nomenclature as c.1224C>T, is a synonymous substitution that does not alter the encoded amino acid (arginine) at position 408. The PAH gene encodes phenylalanine hydroxylase, a liver-specific enzyme responsible for the hydroxylation of phenylalanine to tyrosine, a critical step in phenylalanine metabolism. Mutations in this gene can lead to phenylketonuria (PKU), a metabolic disorder characterized by elevated levels of phenylalanine in the blood, which can result in intellectual disability and other neurological complications if untreated. The p.R408R variant is generally considered a benign polymorphism because it does not change the amino acid sequence of the protein and has not been reported to disrupt splicing or regulatory sequences in a consistent or clinically significant manner. It is commonly found in population databases, including gnomAD, where it is observed with relatively high allele frequencies across multiple populations, further supporting its classification as a likely benign or benign variant. As of now, there are no peer-reviewed publications that directly associate the p.R408R variant with altered PAH function or PKU phenotypes, nor is there evidence from functional studies suggesting any pathogenic mechanism. In silico predictive tools uniformly classify this variant as having no functional impact. However, due to the location of this variant in a region that has been implicated in mRNA splicing for other substitutions, the possibility of subtle effects on transcript processing has been suggested in broader discussions of synonymous variants, though such impacts have not been demonstrated specifically for p.R408R. In the absence of direct functional or clinical association studies specific to this variant, its classification remains benign, with no current evidence indicating a need for clinical concern. Continued genomic surveillance and functional analyses are recommended to confirm the absence of splicing effects or regulatory disruptions, especially in cases where it is observed in compound heterozygosity with known pathogenic variants. No peer-reviewed publications currently provide direct evidence on the functional or clinical impact of the PAH p.R408R variant. | No references available |
|  | Evaluation  **Summarization Accuracy & Completeness:**  **Hallucination Rate:**  **Readability & Usability:**  **Literature Coverage & Relevance:**  **Time Efficiency:** | |
| **3** | The phenylalanine hydroxylase (PAH) gene encodes an enzyme crucial for the metabolism of the amino acid phenylalanine (Phe) into tyrosine. This process is essential for preventing the accumulation of phenylalanine, which can lead to phenylketonuria (PKU), a severe metabolic disorder characterized by neurological and neuropsychological symptoms if untreated or late diagnosed. The variant p.R408R in the PAH gene is not commonly discussed in the literature as a pathogenic variant, suggesting it may not be associated with significant clinical implications or may be a synonymous variant. However, the PAH gene is known to harbor numerous pathogenic variants, including missense mutations like p.Arg408Trp (c.1222C>T), which is a well-documented cause of PKU[6]. Deep intronic variants in the PAH gene have also been identified as contributing to PKU in cases where traditional sequencing methods failed to provide a diagnosis[2]. Given the lack of specific information on the p.R408R variant in peer-reviewed literature, its clinical significance remains uncertain, and further investigation may be necessary to determine its impact on PAH enzyme function and disease risk. References:[2][6]. | [1] <https://www.semanticscholar.org/paper/effddea6a73405c5c0ad6b67e141cc1d060c6998>  [2] <https://www.ncbi.nlm.nih.gov/pmc/articles/PMC10214626/>  [3] <https://pubmed.ncbi.nlm.nih.gov/29653233/>  [4] <https://pubmed.ncbi.nlm.nih.gov/38546978/>  [5] <https://www.semanticscholar.org/paper/b7ceedca9dd53cac832dc687cbd38cd82727a12f>  [6] <https://www.ncbi.nlm.nih.gov/pmc/articles/PMC5081532/>  [7] <https://www.ncbi.nlm.nih.gov/pmc/articles/PMC8112021/>  [8] <https://www.ncbi.nlm.nih.gov/pmc/articles/PMC11806585/> |
|  | Evaluation  **Summarization Accuracy & Completeness:**  **Hallucination Rate:**  **Readability & Usability:**  **Literature Coverage & Relevance:**  **Time Efficiency:** | |
| **4** | Variant Description:  The genomic variant c.1224G>A is a single nucleotide substitution occurring in the PAH gene. This variant results in a synonymous change at the codon level, where the codon for arginine at position 408 (CGG) is altered to another codon for arginine (CGA), hence the protein sequence remains unchanged with the arginine (R) still present at position 408 (p.R408R).  Gene Function:  The PAH gene encodes the enzyme phenylalanine hydroxylase, which is responsible for the conversion of the amino acid phenylalanine to tyrosine. This enzymatic activity is crucial for the catabolism of phenylalanine and the biosynthesis of tyrosine, which is a precursor for neurotransmitters and other biologically important molecules. Proper function of the PAH enzyme is essential for maintaining normal levels of phenylalanine in the blood and preventing its toxic accumulation. | No references available |
|  | Evaluation  **Summarization Accuracy & Completeness:**  **Hallucination Rate:**  **Readability & Usability:**  **Literature Coverage & Relevance:**  **Time Efficiency:** | |
| **5** | The PAH gene encodes phenylalanine hydroxylase, a hepatic enzyme that catalyzes the conversion of the amino acid phenylalanine to tyrosine, a crucial step in phenylalanine catabolism. Mutations in PAH can lead to phenylketonuria (PKU), a metabolic disorder characterized by elevated phenylalanine levels, which, if untreated, may result in intellectual disability and other neurological complications. The variant c.1224G>A (p.R408R) in the PAH gene is a synonymous mutation, meaning it does not alter the amino acid sequence of the enzyme, as arginine remains at position 408. Synonymous mutations are typically considered benign; however, they can sometimes affect gene expression or splicing mechanisms. Current literature does not provide specific evidence regarding the clinical significance or impact of the p.R408R variant on PAH function or PKU manifestation. Notably, the c.1222C>T (p.R408W) mutation at the same codon, resulting in an arginine-to-tryptophan substitution, is recognized as a common pathogenic variant associated with PKU. In the absence of direct studies on the p.R408R variant, its role remains uncertain. Further research is necessary to determine any potential effects on PAH enzyme activity, mRNA stability, or splicing. In clinical settings, the detection of the p.R408R variant should be interpreted with caution, and additional functional analyses are recommended to elucidate its significance. | <https://www.genecards.org/cgi-bin/carddisp.pl?gene=PAH>  <https://www.ncbi.nlm.nih.gov/gene/5053>  <https://www.youtube.com/watch?v=XnVgT8KhR4A>  <https://medlineplus.gov/genetics/gene/pah/>  <https://www.mayoclinic.org/diseases-conditions/phenylketonuria/symptoms-causes/syc-20376302>  <https://respiratory-research.biomedcentral.com/articles/10.1186/s12931-018-0789-9>  <https://www.youtube.com/watch?v=GWBPUt8UwwY>  <https://citeseerx.ist.psu.edu/document?doi=6dab2e6b0cafdaa0d2f2dd09fd25023b597bfa21&repid=rep1&type=pdf>  <https://www.ncbi.nlm.nih.gov/clinvar/RCV000000607/>  <https://pubmed.ncbi.nlm.nih.gov/23457044/>  <https://www.sciencedirect.com/science/article/abs/pii/S109671921630004X>  <https://www.sciencedirect.com/science/article/pii/S0002929723003580>  <https://en.wikipedia.org/wiki/Phenylalanine_hydroxylase>  <https://www.jax.org/strain/039069>  <https://myriad.com/womens-health/diseases/phenylalanine-hydroxylase-deficiency/>  <https://www.uniprot.org/uniprotkb/P00439/entry>  <https://omim.org/entry/612349> |
|  | Evaluation  **Summarization Accuracy & Completeness:**  **Hallucination Rate:**  **Readability & Usability:**  **Literature Coverage & Relevance:**  **Time Efficiency:** | |
