## Supplementary material for "Benchmarking generative AI tools for literature retrieval and summarization in genomic variant interpretation": PAH p.R408W.docx

| **Tool** | **Summary** | **References** |
| --- | --- | --- |
| **1** | The c.1222C>T p.R408W variant in the PAH gene is a missense mutation that results in the substitution of arginine with tryptophan at position 408 of the phenylalanine hydroxylase enzyme. This variant is associated with phenylketonuria (PKU), a metabolic disorder characterized by an inability to metabolize the amino acid phenylalanine due to deficient activity of the PAH enzyme. The PAH gene encodes for the enzyme phenylalanine hydroxylase, which is crucial for the hydroxylation of phenylalanine to tyrosine, a precursor for neurotransmitters and other important biological molecules.  The c.1222C>T variant has been reported to be pathogenic with a high allele frequency in certain populations and is associated with a severe phenotype of classical PKU, characterized by a significant reduction in enzyme activity to approximately 2% of normal [2][4]. This variant is also known as "Celtic" and has been identified as the most frequently occurring variant in a study group, suggesting its importance in the genetic landscape of PKU [2].  Computational predictors of pathogenicity have labeled the c.1222C>T variant as damaging, indicating its potential impact on the stability and function of the PAH protein [1]. The amino acid at position 408 is highly conserved across species, underscoring its functional significance within the enzyme [1]. Furthermore, the variant has been associated with a pathogenic classification according to ACMG/AMP criteria, with multiple lines of evidence supporting its deleterious effect on protein function [4]. In vitro studies have demonstrated that the expression of PAH protein is significantly reduced in cells transfected with the c.1222C>T variant compared to wild-type cells, providing functional evidence of its impact on PAH enzyme activity [3]. Additionally, recent advancements in gene editing technology have explored the potential of base editing to correct the c.1222C>T variant in human hepatocytes, which could offer a therapeutic approach for individuals with this mutation [5]. In summary, the c.1222C>T p.R408W variant in the PAH gene is a well-characterized pathogenic mutation with significant clinical relevance for PKU. Its impact on enzyme activity, conservation across species, and the potential for gene editing correction underscore the importance of this variant in the context of genetic diagnosis and therapeutic development for PKU. | [[1]](https://pubmed.ncbi.nlm.nih.gov/39434944/) Rahimzadeh A et al. (2024). "A Rare Combination of Compound Heterozygous Mutations in the PAH Gene in Three Unrelated Consanguineous Iranian Families with Classical Phenylketonuria." Advanced biomedical research, 13()  [[2]](https://pubmed.ncbi.nlm.nih.gov/37189584/) Iuhas A et al. (2023). "PAH Pathogenic Variants and Clinical Correlations in a Group of Hyperphenylalaninemia Patients from North-Western Romania." Diagnostics (Basel, Switzerland), 13(8)  [[3]](https://pubmed.ncbi.nlm.nih.gov/27786189/) Pan Y et al. (2016). "CRISPR RNA-guided FokI nucleases repair a PAH variant in a phenylketonuria model." Scientific reports, 6()  [[4]](https://pubmed.ncbi.nlm.nih.gov/38136067/) Vela-Amieva M et al. (2023). "In Silico Structural Protein Evaluation of the Phenylalanine Hydroxylase p.(Tyr77His) Variant Associated with Benign Hyperphenylalaninemia as Identified through Mexican Newborn Screening." Children (Basel, Switzerland), 10(12)  [[5]](https://pubmed.ncbi.nlm.nih.gov/37922902/) Brooks DL et al. (2024). "A base editing strategy using mRNA-LNPs for in vivo correction of the most frequent phenylketonuria variant." HGG advances, 5(1)  [[6]](https://pubmed.ncbi.nlm.nih.gov/7825588/) Eisensmith RC et al. (1995). "Recurrence of the R408W mutation in the phenylalanine hydroxylase locus in Europeans." American journal of human genetics, 56(1)  [[7]](https://pubmed.ncbi.nlm.nih.gov/7833927/) Byck S et al. (1994). "Evidence for origin, by recurrent mutation, of the phenylalanine hydroxylase R408W mutation on two haplotypes in European and Quebec populations." Human molecular genetics, 3(9)  [[8]](https://pubmed.ncbi.nlm.nih.gov/8335244/) Ivashchenko TE et al. (1993). "[A simple and reliable method for detection of the R408W mutation in exon 12 of the phenylalanine hydroxylase gene in the molecular diagnosis of phenylketonuria]." Genetika, 29(5)  [[9]](https://pubmed.ncbi.nlm.nih.gov/9156326/) Giannattasio S et al. (1997). "Phenylketonuria mutations and linked haplotypes in the Lithuanian population: origin of the most common R408W mutation." Human heredity, 47(3)  [[10]](https://pubmed.ncbi.nlm.nih.gov/12173030/) O'Donnell KA et al. (2002). "The mutation spectrum of hyperphenylalaninaemia in the Republic of Ireland: the population history of the Irish revisited." European journal of human genetics : EJHG, 10(9)  [[11]](https://pubmed.ncbi.nlm.nih.gov/12655548/) Tighe O et al. (2003). "Genetic diversity within the R408W phenylketonuria mutation lineages in Europe." Human mutation, 21(4)  [[12]](https://pubmed.ncbi.nlm.nih.gov/37924808/) Brooks DL et al. (2023). "Efficient in vivo prime editing corrects the most frequent phenylketonuria variant, associated with high unmet medical need." American journal of human genetics, 110(12)  [[13]](https://pubmed.ncbi.nlm.nih.gov/38712058/) Whittaker MN et al. (2024). "Improved specificity and efficacy of base-editing therapies with hybrid guide RNAs." bioRxiv : the preprint server for biology, ()  [[14]](https://pubmed.ncbi.nlm.nih.gov/39600089/) Yin S et al. (2025). "Amelioration of metabolic and behavioral defects through base editing in the Pah(R408W) phenylketonuria mouse model." Molecular therapy : the journal of the American Society of Gene Therapy, 33(1)  [[15]](https://pubmed.ncbi.nlm.nih.gov/36046396/) Fishchuk L et al. (2022). "The analysis of using a panel of the most common variants in the PAH gene for the newborn screening in Ukraine." Molecular genetics and metabolism reports, 32() |
|  | Evaluation  **Summarization Accuracy & Completeness:**  **Hallucination Rate:**  **Readability & Usability:**  **Literature Coverage & Relevance:**  **Time Efficiency:** | |
| **2** | The PAH gene encodes phenylalanine hydroxylase, a hepatic enzyme that catalyzes the hydroxylation of phenylalanine to tyrosine, a critical step in amino acid metabolism. Mutations in PAH can lead to phenylketonuria (PKU), an autosomal recessive disorder characterized by elevated phenylalanine levels, resulting in intellectual disability and other complications if untreated. The c.1222C>T (p.Arg408Trp) variant, commonly known as R408W, is the most frequent pathogenic mutation in PAH associated with PKU. This missense mutation substitutes arginine with tryptophan at codon 408, impairing enzyme function and leading to phenylalanine accumulation. The R408W mutation exhibits a west-to-east gradient in frequency across Europe, with higher prevalence in Eastern European populations. Studies have shown that the PAH activity of the p.[R408W];[A300S] genotype is approximately 18% of wild-type activity; however, 88% of patients with this genotype exhibit mild hyperphenylalaninemia (MHP), indicating a complex genotype-phenotype correlation. Given the variability in clinical outcomes associated with the R408W variant, individualized management and further research into its phenotypic implications are recommended. | [1] Efficient in vivo prime editing corrects the most frequent pathogenic mutation in phenylketonuria. *The American Journal of Human Genetics*, 2023.[ScienceDirect+1Wikipedia+1](https://www.sciencedirect.com/science/article/pii/S0002929723003580?utm_source=chatgpt.com)  [2] Genetic diversity within the R408W phenylketonuria mutation. *Human Mutation*, 2003.[Wikipedia+2Wiley Online Library+2Wikipedia+2](https://onlinelibrary.wiley.com/doi/pdf/10.1002/humu.10195?utm_source=chatgpt.com)  [3] Co-expression of phenylalanine hydroxylase variants and effects of individual mutations on enzyme activity and stability. *Molecular Genetics and Metabolism*, 2016.[Wikipedia+2PubMed+2Wikipedia+2](https://pubmed.ncbi.nlm.nih.gov/26803807/?utm_source=chatgpt.com)  Others:  <https://www.genecards.org/cgi-bin/carddisp.pl?gene=PAH&utm_source=chatgpt.com>  <https://www.sciencedirect.com/science/article/pii/S0002929723003580>  <https://onlinelibrary.wiley.com/doi/pdf/10.1002/humu.10195>  <https://pubmed.ncbi.nlm.nih.gov/26803807/>  <https://www.ncbi.nlm.nih.gov/gene/5053>  <https://medlineplus.gov/genetics/gene/pah/>  <https://www.mayoclinic.org/diseases-conditions/phenylketonuria/symptoms-causes/syc-20376302>  <https://pubs.acs.org/doi/10.1021/acsomega.9b02955>  <https://www.nature.com/articles/jhg2013136>  <https://www.nature.com/articles/jhg200937>  <https://www.ncbi.nlm.nih.gov/clinvar/variation/577/>  <https://pmc.ncbi.nlm.nih.gov/articles/PMC1801324/>  <https://pmc.ncbi.nlm.nih.gov/articles/PMC9421484/>  <https://www.sciencedirect.com/science/article/abs/pii/S1096719212001163>  <https://www.sciencedirect.com/science/article/pii/S0002929723003580>  <https://www.sciencedirect.com/science/article/abs/pii/S0890850884710449>  <https://en.wikipedia.org/wiki/Phenylalanine_hydroxylase>  <https://www.nature.com/articles/5200153.pdf>  <https://academic.oup.com/hmg/article/33/12/1074/7634000?login=false>  <https://www.jax.org/strain/039069>  <https://omim.org/entry/612349>  https://onlinelibrary.wiley.com/doi/pdf/10.1002/humu.10195 |
|  | Evaluation  **Summarization Accuracy & Completeness:**  **Hallucination Rate:**  **Readability & Usability:**  **Literature Coverage & Relevance:**  **Time Efficiency:** | |
| **3** | The PAH gene encodes the enzyme phenylalanine hydroxylase (PAH), which catalyzes the conversion of phenylalanine to tyrosine, a crucial step in phenylalanine metabolism. Mutations in the PAH gene are associated with phenylketonuria (PKU), an autosomal recessive disorder characterized by the accumulation of phenylalanine, leading to severe neurological impairments if untreated. The variant p.R408W (c.1222C>T) in the PAH gene is one of the most common and well-studied mutations associated with PKU. This missense variant results from a C to T substitution at nucleotide position 1222 in exon 12, causing an arginine to tryptophan substitution at amino acid position 408. The p.R408W variant is classified as pathogenic and is linked to classical PKU, which presents with severe phenylalanine hydroxylase deficiency. The frequency of this variant varies across populations but is notably high in European populations, where it accounts for a significant proportion of PKU cases. In Poland, the p.R408W mutation was observed in 68% of PKU patients, while in Estonia, it was present on 84% of mutant alleles. The variant is associated with null to very low PAH activity, leading to the classical PKU phenotype. Genotyping for p.R408W has clinical utility in diagnosing PKU and predicting the potential response to co-factor therapy, such as tetrahydrobiopterin (BH4). The variant's high prevalence in certain populations underscores its importance in genetic screening and counseling for PKU. However, the exact mechanisms by which p.R408W affects PAH enzyme function and the variability in clinical outcomes remain areas of ongoing research. The recurrence of p.R408W on different haplotypes suggests multiple origins of this mutation, further complicating its genetic landscape. Despite extensive research, uncertainties persist regarding the full spectrum of p.R408W's impact on PAH function and PKU phenotypes, warranting further investigation. | [1] Byck S, Morgan K, Tyfield L, Dworniczak B, Scriver CR. Hum Mol Genet. 1994 Sep;3(9):1675-7. doi: 10.1093/hmg/3.9.1675.  [2] Eisensmith RC, Goltsov AA, O'Neill C, Tyfield LA, Schwartz EI, Kuzmin AI, Baranovskaya SS, Tsukerman GL, Treacy E, Scriver CR, et al. Am J Hum Genet. 1995 Jan;56(1):278-86.  [3] DiLella AG, Marvit J, Lichter-Konecki U, Lidsky AS, Guthrie R, Konecki D, Woo SLC. Hum Genet. 1987;77(2):148-52. doi: 10.1007/BF00289264.  [4] Jaruzelska J, Zurawiecka A, Krajewska-Walasek M, Gregersen N, Guttler F. Hum Mutat. 1998;11(5):390-6. doi: 10.1002/(SICI)1098-1004(1998)11:5<390::AID-HUMU4>3.0.CO;2-J.  [5] Lilleväli H, Petrovic I, Käärik R, Reimann E, Ounap K. Eur J Hum Genet. 1996;4(5):296-300. doi: 10.1159/000472217. |
|  | Evaluation  **Summarization Accuracy & Completeness:**  **Hallucination Rate:**  **Readability & Usability:**  **Literature Coverage & Relevance:**  **Time Efficiency:** | |
| **4** | The PAH gene encodes the enzyme phenylalanine hydroxylase, which plays a critical role in phenylalanine metabolism by catalyzing the hydroxylation of phenylalanine to tyrosine. Deficiency in this enzyme results in phenylalanine accumulation, causing phenylketonuria (PKU), an autosomal recessive metabolic disorder. The variant p.R408W (c.1222C>T) is one of the most common and severe missense mutations in the PAH gene, particularly prevalent among individuals of European descent, and is typically associated with the classical, most severe phenotype of PKU. The mutation leads to a substitution of arginine with tryptophan at position 408, which severely disrupts enzyme structure and stability, causing loss of function. Functional studies have demonstrated that the R408W variant results in minimal residual enzyme activity and poor responsiveness to cofactor therapy with tetrahydrobiopterin (BH4), complicating treatment options and necessitating strict dietary management. The high pathogenicity of this mutation is well-documented in diverse populations, including Russian, Italian, and Brazilian cohorts, where it frequently occurs in homozygous or compound heterozygous states and is strongly associated with elevated phenylalanine levels and severe metabolic phenotype [1](https://onlinelibrary.wiley.com/doi/abs/10.1002/humu.20526), [2](https://www.researchgate.net/profile/Rosa-Santacroce/publication/280447809), [3](https://onlinelibrary.wiley.com/doi/abs/10.1002/mgg3.408), [4](https://cyberleninka.ru/article/n/determining-the-frequency-of-pah-mutations-in-moscow-region-residents-with-phenylketonuria-using-a-combination-of-real-time-pcr-and-next). Studies of protein expression in vitro further confirm the R408W variant’s detrimental impact on PAH protein folding and stability, leading to its rapid degradation [5](https://www.sciencedirect.com/science/article/pii/S109671921830283X). Given its well-characterized pathogenicity, p.R408W is consistently classified as a pathogenic variant in clinical and molecular genetics databases and is included in mutation panels for diagnostic screening of PKU. Although the understanding of p.R408W's functional and clinical effects is robust, ongoing studies continue to assess its prevalence in underrepresented populations and explore potential therapeutic avenues. | [1] Scriver CR. *The PAH gene, phenylketonuria, and a paradigm shift*. [DOI:10.1002/humu.20526](https://onlinelibrary.wiley.com/doi/abs/10.1002/humu.20526)  [2] Trunzo R et al. *Phenylalanine hydroxylase deficiency in south Italy: Genotype–phenotype correlations*. [PDF](https://www.researchgate.net/profile/Rosa-Santacroce/publication/280447809)  [3] Vieira Neto E et al. *Mutation analysis of the PAH gene in phenylketonuria patients from Rio de Janeiro*. [DOI:10.1002/mgg3.408](https://onlinelibrary.wiley.com/doi/abs/10.1002/mgg3.408)  [4] Nikiforova AI, Abramov DD. *Determining the frequency of PAH mutations in Moscow region residents with phenylketonuria*. [Full text](https://cyberleninka.ru/article/n/determining-the-frequency-of-pah-mutations-in-moscow-region-residents-with-phenylketonuria-using-a-combination-of-real-time-pcr-and-next)  [5] Himmelreich N et al. *Relationship between genotype, phenylalanine hydroxylase expression and activity*. [PDF](https://www.academia.edu/download/93329098/j.ymgme.2018.06.01120221030-1-xe98t6.pdf) |
|  | Evaluation  **Summarization Accuracy & Completeness:**  **Hallucination Rate:**  **Readability & Usability:**  **Literature Coverage & Relevance:**  **Time Efficiency:** | |
| **5** | The phenylalanine hydroxylase (PAH) gene encodes an enzyme crucial for the metabolism of the amino acid phenylalanine. This enzyme catalyzes the conversion of phenylalanine to tyrosine, a process essential for maintaining normal levels of phenylalanine in the body. Variants in the PAH gene can lead to phenylketonuria (PKU), an autosomal recessive disorder characterized by elevated phenylalanine levels, which can cause severe neurological and neuropsychological symptoms if untreated or late diagnosed.  The variant p.R408W (c.1222C>T) is one of the most common mutations in the PAH gene, with a reported allele frequency of approximately 21.4% in certain populations[6]. This mutation results in the substitution of arginine with tryptophan at position 408 of the phenylalanine hydroxylase protein. The p.R408W variant is classified as pathogenic and significantly impairs the enzyme's activity, leading to PKU.  Research has shown that correcting this variant using genome editing techniques, such as the CRISPR/Cas9 system modified with FokI endonuclease, can restore PAH activity in vitro[6]. This approach suggests potential therapeutic applications for treating PKU by correcting disease-causing variants in the PAH gene.  While the p.R408W variant is well-studied in the context of PKU, further research is needed to fully understand its impact on protein function and its role in disease severity. The classification and frequency of this variant highlight its importance in genetic diagnostics for PKU. | [1] <https://www.semanticscholar.org/paper/effddea6a73405c5c0ad6b67e141cc1d060c6998>  [2] <https://www.ncbi.nlm.nih.gov/pmc/articles/PMC10214626/>  [3] <https://pubmed.ncbi.nlm.nih.gov/29653233/>  [4] <https://pubmed.ncbi.nlm.nih.gov/38546978/>  [5] <https://www.semanticscholar.org/paper/b7ceedca9dd53cac832dc687cbd38cd82727a12f>  [6] <https://www.ncbi.nlm.nih.gov/pmc/articles/PMC5081532/>  [7] <https://www.ncbi.nlm.nih.gov/pmc/articles/PMC8112021/>  [8] <https://www.ncbi.nlm.nih.gov/pmc/articles/PMC11806585/> |
|  | Evaluation  **Summarization Accuracy & Completeness:**  **Hallucination Rate:**  **Readability & Usability:**  **Literature Coverage & Relevance:**  **Time Efficiency:** | |
