## Supplementary material for "Benchmarking generative AI tools for literature retrieval and summarization in genomic variant interpretation": PRPH2 p.L185P.docx

| **Tool** | **Summary** | **References** |
| --- | --- | --- |
| **1** | The PRPH2 gene, also known as RDS, encodes peripherin-2, a protein crucial for normal vision. Peripherin-2 is integral to the structure and function of photoreceptor cells in the retina, specifically in the outer segment discs of rods and cones. It acts as an adhesion molecule, stabilizing and compacting these discs, which are essential for phototransduction. Mutations in PRPH2 are associated with various retinal dystrophies, including retinitis pigmentosa, macular degeneration, and pattern dystrophies, highlighting its critical role in maintaining retinal integrity.  The p.L185P variant in the PRPH2 gene is a missense mutation that has been linked to retinal diseases. This variant was first identified in association with autosomal dominant retinitis pigmentosa and has since been implicated in other retinal disorders. The substitution of leucine with proline at position 185 alters the protein structure, potentially affecting its ability to form stable complexes with other proteins, such as ROM-1, which is essential for the maintenance of photoreceptor outer segment discs. Studies have shown that the p.L185P mutation can cause a digenic form of retinitis pigmentosa when combined with a null mutation in the ROM1 gene, indicating a complex genetic interaction.  The frequency of the p.L185P variant is not well-documented in the general population, but it has been observed in multiple cases of inherited retinal diseases. Its impact on protein function is significant, leading to the degeneration of photoreceptor cells and progressive vision loss. The classification of this variant is typically as pathogenic or likely pathogenic, based on its association with severe retinal phenotypes. However, the exact mechanisms by which p.L185P contributes to disease pathogenesis remain unclear, and further research is needed to elucidate its role in retinal degeneration.  In summary, the p.L185P variant in the PRPH2 gene is a rare but significant mutation that disrupts the normal function of peripherin-2, leading to various retinal dystrophies. Its classification as pathogenic underscores the need for continued investigation into its molecular effects and potential therapeutic targets. | [1] Farrar et al., 1991, Kajiwara et al., 1991  [2] Wang et al., 2013  [3] Kajiwara et al., 1994 |
|  | Evaluation  **Summarization Accuracy & Completeness:**  **Hallucination Rate:**  **Readability & Usability:**  **Literature Coverage & Relevance:**  **Time Efficiency:** | |
| **2** | The PRPH2 gene encodes Peripherin-2, a membrane-associated glycoprotein that is crucial for the function of photoreceptor outer segments. The c.554T>C p.L185P rs121918563 variant in the PRPH2 gene results in the substitution of leucine with proline at position 185 of the protein. This missense variant is located in a critical region of Peripherin-2, specifically in the second intradiscal loop, which is essential for the formation of PRPH2 homotetramers and PRPH2/ROM1 heterotetramers. These complexes are necessary for proper disc morphogenesis, and their impairment can lead to photoreceptor degeneration and visual loss [1].  Functional studies have shown that the PRPH2 L185P variant, when expressed alone, is unable to form homotetramers. However, it can still form heterotetramers with the wild-type ROM1 protein. Despite this, the formation of higher-order complexes that span the entire circumference of the disc is compromised in the presence of the L185P variant. This suggests that individuals carrying the L185P variant may have a reduced ability to form these crucial higher-order complexes, potentially leading to the phenotypic manifestations of pattern dystrophy (PD) observed in affected individuals [1].  The L185P variant has been associated with retinal disorders, including autosomal dominant PD, with clinical features such as adult-onset vitelliform macular dystrophy and butterfly macular dystrophy. These conditions typically progress to retinal pigment epithelial irregularities and central macular atrophy over time. The variant has been identified in individuals with retinopathies characterized by low genetic susceptibility to age-related macular degeneration (AMD) and a younger age of onset. The mean age at diagnosis for individuals with the L185P variant was reported to be 56 years. The variant has been associated with better visual acuity during follow-up compared to another pathogenic variant in the PRPH2 gene [1].  In a mouse model, the L185P mutation in combination with a null mutation in Rom1 has been shown to cause a rare digenic form of retinitis pigmentosa (RP). Mice heterozygous for both mutations exhibited late-onset thinning of the outer nuclear layer and reduced scotopic electroretinograms, providing insights into the pathophysiology of PRPH2-related diseases [2].  The L185P variant is reported in ClinVar with eight pathogenic submissions, underscoring its clinical significance in the context of hereditary retinal dystrophies. | [[1]](https://pubmed.ncbi.nlm.nih.gov/39693084/) Seddon JM et al. (2024). "Clinical and Imaging Characteristics of PRPH2 Retinopathies in a Longitudinal Cohort and Diagnostic Implications." Investigative ophthalmology & visual science, 65(14)  [[2]](https://pubmed.ncbi.nlm.nih.gov/32213850/) Tebbe L et al. (2020). "The Interplay between Peripherin 2 Complex Formation and Degenerative Retinal Diseases." Cells, 9(3)  [[3]](https://pubmed.ncbi.nlm.nih.gov/9331261/) Dryja TP et al. (1997). "Dominant and digenic mutations in the peripherin/RDS and ROM1 genes in retinitis pigmentosa." Investigative ophthalmology & visual science, 38(10)  [[4]](https://pubmed.ncbi.nlm.nih.gov/10800708/) Goldberg AF et al. (2000). "Expression and characterization of peripherin/rds-rom-1 complexes and mutants implicated in retinal degenerative diseases." Methods in enzymology, 316()  [[5]](https://pubmed.ncbi.nlm.nih.gov/11297544/) Loewen CJ et al. (2001). "Molecular characterization of peripherin-2 and rom-1 mutants responsible for digenic retinitis pigmentosa." The Journal of biological chemistry, 276(25)  [[6]](https://pubmed.ncbi.nlm.nih.gov/11427722/) Kedzierski W et al. (2001). "Deficiency of rds/peripherin causes photoreceptor death in mouse models of digenic and dominant retinitis pigmentosa." Proceedings of the National Academy of Sciences of the United States of America, 98(14)  [[7]](https://pubmed.ncbi.nlm.nih.gov/38474159/) Fernandez-Caballero L et al. (2024). "PRPH2-Related Retinal Dystrophies: Mutational Spectrum in 103 Families from a Spanish Cohort." International journal of molecular sciences, 25(5)  [[8]](https://pubmed.ncbi.nlm.nih.gov/38540785/) Hitti-Malin RJ et al. (2024). "Towards Uncovering the Role of Incomplete Penetrance in Maculopathies through Sequencing of 105 Disease-Associated Genes." Biomolecules, 14(3)  [[9]](https://pubmed.ncbi.nlm.nih.gov/23847139/) Wang X et al. (2013). "Comprehensive molecular diagnosis of 179 Leber congenital amaurosis and juvenile retinitis pigmentosa patients by targeted next generation sequencing." Journal of medical genetics, 50(10)  [[10]](https://pubmed.ncbi.nlm.nih.gov/24154662/) Wang F et al. (2014). "Next generation sequencing-based molecular diagnosis of retinitis pigmentosa: identification of a novel genotype-phenotype correlation and clinical refinements." Human genetics, 133(3)  [[11]](https://pubmed.ncbi.nlm.nih.gov/25304133/) Daiger SP et al. (2014). "Genes and Mutations Causing Autosomal Dominant Retinitis Pigmentosa." Cold Spring Harbor perspectives in medicine, 5(10)  [[12]](https://pubmed.ncbi.nlm.nih.gov/38743414/) Heath Jeffery RC et al. (2024). "Retinal Dystrophies Associated With Peripherin-2: Genetic Spectrum and Novel Clinical Observations in 241 Patients." Investigative ophthalmology & visual science, 65(5)  [[13]](https://pubmed.ncbi.nlm.nih.gov/20300562/) Sato H et al. (2010). "A monogenic dominant mutation in Rom1 generated by N-ethyl-N-nitrosourea mutagenesis causes retinal degeneration in mice." Molecular vision, 16()  [[14]](https://pubmed.ncbi.nlm.nih.gov/22131869/) Ferrari S et al. (2011). "Retinitis pigmentosa: genes and disease mechanisms." Current genomics, 12(4)  [[15]](https://pubmed.ncbi.nlm.nih.gov/31126147/) Diakatou M et al. (2019). "Genome Editing as a Treatment for the Most Prevalent Causative Genes of Autosomal Dominant Retinitis Pigmentosa." International journal of molecular sciences, 20(10) |
|  | Evaluation  **Summarization Accuracy & Completeness:**  **Hallucination Rate:**  **Readability & Usability:**  **Literature Coverage & Relevance:**  **Time Efficiency:** | |
| **3** | The PRPH2 gene encodes peripherin-2, a photoreceptor-specific transmembrane glycoprotein critical for the morphogenesis and structural maintenance of photoreceptor outer segment discs in the retina. Mutations in PRPH2 are associated with a range of autosomal dominant retinal dystrophies including retinitis pigmentosa (RP), pattern dystrophy, and adult-onset vitelliform macular dystrophy. The p.L185P (leucine to proline substitution at codon 185) variant is a rare missense mutation located in the second intradiscal loop of the peripherin-2 protein, a region vital for protein–protein interactions and disc rim structure. The L185P variant has been reported in patients with autosomal dominant RP, often in compound or double heterozygous states, suggesting variable expressivity and incomplete penetrance. One study described a Chinese patient harboring the p.L185P mutation in a double heterozygous state who presented with RP symptoms, while others with the same variant showed varying degrees of severity or even asymptomatic phenotypes, indicating a possible role for genetic modifiers or oligogenic inheritance in disease expression [1]. Although pathogenicity is supported by clinical correlation and the known functional importance of the affected protein region, the low frequency of this variant and phenotypic variability limit the strength of genotype-phenotype assertions. To date, no large cohort studies have definitively established the pathogenic classification of p.L185P, and it remains a variant of uncertain significance (VUS) in several clinical contexts. The mutation does not appear in large population databases at appreciable frequencies, consistent with a potentially deleterious effect. Nonetheless, additional functional assays and broader clinical studies are required to confirm its impact. In conclusion, while the p.L185P variant in PRPH2 is likely to contribute to retinal degeneration in certain contexts, the available evidence remains limited and inconclusive, warranting further investigation for clinical interpretation and genetic counseling. | [1] [Xiao et al. (2017](https://app.scholarai.io/paper?paper_id=DOI:10.5301/ejo.5000971&original_url=https%3A%2F%2Fjournals.sagepub.com%2Fdoi%2Fabs%2F10.5301%2Fejo.5000971) |
|  | Evaluation  **Summarization Accuracy & Completeness:**  **Hallucination Rate:**  **Readability & Usability:**  **Literature Coverage & Relevance:**  **Time Efficiency:** | |
| **4** | The PRPH2 gene encodes peripherin-2, a critical glycoprotein integral to the structure and function of photoreceptor cells in the human retina. Peripherin-2 is essential for the morphogenesis and stability of the outer segment disks in both rod and cone cells, playing a pivotal role in phototransduction and overall visual function . The c.554T>C (p.Leu185Pro) variant in PRPH2 has been identified as pathogenic and is associated with autosomal dominant pattern dystrophy (PD), including adult-onset vitelliform macular dystrophy (AVMD) and butterfly-shaped pigment dystrophy. Individuals heterozygous for the p.L185P variant typically present with macular abnormalities in their fourth to fifth decades of life, progressing to retinal pigment epithelial irregularities and central macular atrophy over approximately two decades. Notably, this variant is associated with a relatively better visual prognosis compared to other PRPH2 mutations . The p.L185P variant has also been implicated in digenic retinitis pigmentosa (RP) when co-inherited with specific ROM1 mutations, such as p.G113E, leading to a more severe retinal degeneration phenotype . Given the potential for variable expressivity and the possibility of digenic interactions, comprehensive genetic counseling and longitudinal ophthalmic evaluations are recommended for individuals carrying the p.L185P variant to monitor disease progression and guide management strategies. | Seddon JM, et al. Clinical and Imaging Characteristics of PRPH2 Retinopathies in a Longitudinal Cohort and Diagnostic Implications. *Investigative Ophthalmology & Visual Science*. 2024.[ResearchGate+1PMC+1](https://www.researchgate.net/publication/387179022_Clinical_and_Imaging_Characteristics_of_PRPH2_Retinopathies_in_a_Longitudinal_Cohort_and_Diagnostic_Implications?utm_source=chatgpt.com)  Keen TJ, et al. Rom1 converts Y141C-Prph2-associated pattern dystrophy to retinitis pigmentosa. *Human Molecular Genetics*. 2017.[UniProt+4Academic OUP+4ScienceDirect+4](https://academic.oup.com/hmg/article/26/3/509/2805388?utm_source=chatgpt.com)  Molday RS, et al. Molecular Characterization of Peripherin-2 and Rom-1 Mutants Responsible for Digenic Retinitis Pigmentosa. *Journal of Biological Chemistry*. 1999.[OMIM+6ScienceDirect+6PMC+6](https://www.sciencedirect.com/science/article/pii/S0021925820785182?utm_source=chatgpt.com)  Others:  <https://en.wikipedia.org/wiki/Peripherin_2>  <https://academic.oup.com/hmg/article/26/3/509/2805388>  <https://www.sciencedirect.com/science/article/pii/S0021925820785182>  <https://www.ncbi.nlm.nih.gov/gene/5961>  <https://www.mdpi.com/1422-0067/25/5/2913>  <https://www.uniprot.org/uniprotkb/P23942/entry>  <https://www.fightingblindness.org/diseases/prph2>  <https://ojrd.biomedcentral.com/articles/10.1186/s13023-021-01759-8>  <https://www.ncbi.nlm.nih.gov/clinvar/variation/VCV000013165.12/>  <https://pmc.ncbi.nlm.nih.gov/articles/PMC4208711/>  <https://iovs.arvojournals.org/article.aspx?articleid=2802377>  <https://tvst.arvojournals.org/article.aspx?articleid=2802113>  <https://omim.org/entry/179605>  <https://rgd.mcw.edu/rgdweb/report/gene/main.html?id=RGD%3A3549>  <https://www.genecards.org/cgi-bin/carddisp.pl?gene=PRPH2>  <https://academic.oup.com/hmg/article/26/3/509/2805388?login=false>  <https://medlineplus.gov/download/genetics/gene/prph2.pdf>  <https://www.sciencedirect.com/science/article/pii/S0021925820785182>  <https://www.sciencedirect.com/science/article/abs/pii/S1350946215300045> |
|  | Evaluation  **Summarization Accuracy & Completeness:**  **Hallucination Rate:**  **Readability & Usability:**  **Literature Coverage & Relevance:**  **Time Efficiency:** | |
| **5** | The PRPH2 gene encodes peripherin 2, a photoreceptor-specific glycoprotein crucial for the formation and maintenance of outer segments in rod and cone photoreceptors[1][5]. Mutations in PRPH2 are associated with inherited retinal diseases (IRDs), including retinitis pigmentosa and central areolar choroidal dystrophy[1][2]. The variant p.L185P in the PRPH2 gene has not been specifically documented in the available literature, suggesting a need for further research to understand its pathogenicity and clinical implications. Generally, PRPH2 mutations can lead to significant heterogeneity in clinical presentation, affecting visual acuity and retinal structure[1]. Animal models, such as Prph2 mutant mice, have been used to study the disease pathophysiology and test therapeutic approaches, including gene replacement therapy, which has shown promise in improving retinal function in models with PRPH2-related retinal degeneration[2][5]. Establishing genotype-phenotype correlations and understanding the fundamental biology of PRPH2 are key challenges in developing effective treatments for PRPH2-associated IRDs[1]. Given the lack of specific information on the p.L185P variant, further investigation is recommended to determine its clinical significance and potential impact on retinal health. References:[1][2][5]. | [1] <https://www.ncbi.nlm.nih.gov/pmc/articles/PMC11469193/>  [2] <https://www.ncbi.nlm.nih.gov/pmc/articles/PMC10620171/>  [3] <https://www.semanticscholar.org/paper/a808574e9f9a13070465db510c2d282091416792>  [4] <https://pubmed.ncbi.nlm.nih.gov/10888879/>  [5] <https://pubmed.ncbi.nlm.nih.gov/12950066/>  [6] <https://pubmed.ncbi.nlm.nih.gov/35941228/>  [7] <https://pubmed.ncbi.nlm.nih.gov/11689482/>  [8] <https://pubmed.ncbi.nlm.nih.gov/30926958/> |
|  | Evaluation  **Summarization Accuracy & Completeness:**  **Hallucination Rate:**  **Readability & Usability:**  **Literature Coverage & Relevance:**  **Time Efficiency:** | |
