## Supplementary material for "Benchmarking generative AI tools for literature retrieval and summarization in genomic variant interpretation": RAG1 c.256_257del.docx

| **Tool** | **Summary** | **References** |
| --- | --- | --- |
| **1** | The RAG1 gene encodes a crucial component of the V(D)J recombinase, which mediates the rearrangement and recombination of immunoglobulin and T-cell receptor genes in developing lymphocytes, a process essential for adaptive immunity[6](https://pubmed.ncbi.nlm.nih.gov/35948399/)[8](https://pubmed.ncbi.nlm.nih.gov/25745109/). Specifically, RAG1, along with its partner protein RAG2, initiates DNA cleavage at recombination signal sequences (RSS) flanking V, D, and J gene segments[8](https://pubmed.ncbi.nlm.nih.gov/25745109/). This process generates the diverse repertoire of antigen receptors necessary for recognizing and responding to a wide range of pathogens. Mutations in RAG1 can lead to various immunodeficiencies, ranging from severe combined immunodeficiency (SCID) to Omenn syndrome (OS), depending on the severity of the mutation and its impact on protein function[2](https://pubmed.ncbi.nlm.nih.gov/38324638/). The c.256_257del variant in RAG1 refers to a deletion of two nucleotides at positions 256 and 257 within the coding sequence of the gene. This deletion typically results in a frameshift, leading to a premature stop codon and a truncated, non-functional protein. Such frameshift mutations are generally classified as pathogenic due to their disruptive effect on protein structure and function. The absence of a functional RAG1 protein severely impairs V(D)J recombination, preventing the development of mature B and T cells[2](https://pubmed.ncbi.nlm.nih.gov/38324638/). Consequently, individuals with this mutation may present with SCID, characterized by a profound deficiency of both B and T lymphocytes, or Omenn Syndrome[7](https://www.ncbi.nlm.nih.gov/pmc/articles/PMC2740248/). Clinical presentation can vary based on the zygosity and the presence of other possible mutations. At this time, specific data on the frequency, classification, and nomenclature of the c.256_257del variant is not available in the reviewed literature. However, gene editing strategies have been developed to target a corrective sequence into the human RAG1 gene by homology-directed repair (HDR), which can rescue RAG1 expression and function in human hematopoietic stem and progenitor cells[2](https://pubmed.ncbi.nlm.nih.gov/38324638/). | [2](https://pubmed.ncbi.nlm.nih.gov/38324638/) Exonic knockout and knockin gene editing in hematopoietic stem and progenitor cells rescues RAG1 immunodeficiency.  [6](https://pubmed.ncbi.nlm.nih.gov/35948399/) The RAG1 Ubiquitin Ligase Domain Stimulates Recombination of TCRβ and TCRα Genes and Influences Development of αβ T Cell Lineages.  [7](https://www.ncbi.nlm.nih.gov/pmc/articles/PMC2740248/) Homozygous R396H mutation of the RAG1 gene in a Saudi infant with Omenn’s syndrome: a case report.  [8](https://pubmed.ncbi.nlm.nih.gov/25745109/) Mapping and Quantitation of the Interaction between the Recombination Activating Gene Proteins RAG1 and RAG2*♦.   1. <https://www.semanticscholar.org/paper/f54c580c4569dd6ec450d68b2a22e8969fd9e9e2> 2. <https://pubmed.ncbi.nlm.nih.gov/38324638/> 3. <https://www.semanticscholar.org/paper/a9f287ab883193b59bd861c7a45f810cfade0dcf> 4. <https://www.ncbi.nlm.nih.gov/pmc/articles/PMC10454224/> 5. <https://www.ncbi.nlm.nih.gov/pmc/articles/PMC9859925/> 6. <https://pubmed.ncbi.nlm.nih.gov/35948399/> 7. <https://www.ncbi.nlm.nih.gov/pmc/articles/PMC2740248/> 8. <https://pubmed.ncbi.nlm.nih.gov/25745109/> |
|  | Evaluation  **Summarization Accuracy & Completeness:**  **Hallucination Rate:**  **Readability & Usability:**  **Literature Coverage & Relevance:**  **Time Efficiency:** | |
| **2** | The RAG1 gene encodes the recombination-activating protein 1, a crucial component of the RAG complex responsible for initiating V(D)J recombination in developing B and T lymphocytes. This process generates diverse antigen receptor repertoires essential for adaptive immunity by rearranging variable (V), diversity (D), and joining (J) gene segments. The c.256_257del variant in RAG1 denotes a deletion of two nucleotides at positions 256 and 257 of the coding sequence. This deletion likely results in a frameshift mutation, leading to a premature stop codon and truncated protein product, potentially disrupting normal RAG1 function. Such loss-of-function mutations in RAG1 are associated with severe immunodeficiency disorders, including Omenn syndrome and severe combined immunodeficiency (SCID), characterized by impaired lymphocyte development and function. The specific c.256_257del variant has not been extensively documented in peer-reviewed literature, indicating a need for further investigation to determine its clinical significance. Given the potential impact on immune function, individuals harboring this variant should undergo comprehensive immunological evaluation to assess lymphocyte populations and function. Additionally, genetic counseling is recommended to discuss potential implications and guide clinical management. Further research is essential to elucidate the precise effects of the c.256_257del variant on RAG1 function and its contribution to immunodeficiency phenotypes. | <https://www.ncbi.nlm.nih.gov/gene/5896>  <https://www.ncbi.nlm.nih.gov/clinvar/RCV000282657/>  <https://www.medicaljournals.se/acta/content/html/10.2340/00015555-3042>  <https://medlineplus.gov/genetics/gene/rag1/>  <https://medlineplus.gov/download/genetics/gene/rag1.pdf>  <https://www.uniprot.org/uniprotkb/P15918/entry>  <https://www.ncbi.nlm.nih.gov/clinvar/RCV002222473.1/>  <https://pmc.ncbi.nlm.nih.gov/articles/PMC109053/>  <https://pmc.ncbi.nlm.nih.gov/articles/PMC2676217/>  <https://en.wikipedia.org/wiki/RAG1>  <https://en.wikipedia.org/wiki/Recombination-activating_gene>  <https://erepo.clinicalgenome.org/evrepo/ui/interpretation/e2029994-f2fc-46df-8931-f8afe1e193ef>  <https://www.genecards.org/cgi-bin/carddisp.pl?gene=RAG1>  <https://clinvarminer.genetics.utah.edu/variants-by-mondo-condition/11338/gene/RAG1/pathogenic>  <https://clinvarminer.genetics.utah.edu/variants-by-gene/RAG1/condition/Combined%20immunodeficiency%20due%20to%20partial%20RAG1%20deficiency/pathogenic>  <https://www.geniadb.net/app/gene/info.php?id=26682>  <https://frontiersin.figshare.com/articles/dataset/Table_1_The_Clinical_and_Genetic_Spectrum_of_82_Patients_With_RAG_Deficiency_Including_a_c_256_257delAA_Founder_Variant_in_Slavic_Countries_xlsx/12457754>  <https://www.frontiersin.org/journals/immunology/articles/10.3389/fimmu.2023.1210818/full> |
|  | Evaluation  **Summarization Accuracy & Completeness:**  **Hallucination Rate:**  **Readability & Usability:**  **Literature Coverage & Relevance:**  **Time Efficiency:** | |
| **3** | The genomic variant c.256_257del, also known as p.Lys86ValfsTer33 and designated with the rs number rs772962160, is a frameshift deletion found in the RAG1 gene. This variant results from the deletion of two adjacent adenine nucleotides, leading to a frameshift and the creation of a premature stop codon at position 33 downstream of the lysine at position 86 in the RAG1 protein sequence [1]. The presence of this variant has been associated with a range of immunodeficiency phenotypes, including severe combined immunodeficiency (SCID), Omenn syndrome (OS), and other related conditions [1].  The RAG1 gene encodes one of the two lymphoid-specific components of the V(D)J recombinase complex, which is essential for the somatic recombination process that generates the diverse repertoire of immunoglobulin and T-cell receptor genes during lymphocyte development. Mutations in RAG1 can therefore lead to a spectrum of immunodeficiency disorders, depending on the residual activity of the RAG1 protein [1].  The c.256_257del variant has been identified in both compound heterozygous and homozygous states in patients, with the homozygous state often observed in populations with a higher frequency of consanguinity [1]. This variant has been reported to be particularly prevalent among Slavic populations, with a notable frequency in Polish cohorts, suggesting a possible founder effect in this region [2]. The distribution of this variant has been mapped, showing a concentration in the Vistula river basin area of Poland [4].  Clinical manifestations associated with the c.256_257del variant are variable, ranging from incomplete OS to typical OS and SCID without OS features [1]. The variant has been detected in patients using Sanger sequencing and is included in ClinVar with multiple pathogenic submissions, underscoring its clinical relevance in the context of immunodeficiency disorders.  In summary, the c.256_257del p.Lys86ValfsTer33 rs772962160 variant in the RAG1 gene is a pathogenic frameshift deletion that has significant implications for lymphocyte development and function, leading to a range of immunodeficiency phenotypes. Its prevalence in certain Slavic populations indicates a potential founder effect, and its detection is crucial for the diagnosis and management of affected individuals. | [[1]](https://pubmed.ncbi.nlm.nih.gov/28083621/) Szaflarska A et al. (2016). "Mutation c.256_257delAA in RAG1 Gene in Polish Children with Severe Combined Immunodeficiency: Diversity of Clinical Manifestations." Archivum immunologiae et therapiae experimentalis, 64(Suppl 1)  [[2]](https://pubmed.ncbi.nlm.nih.gov/32655540/) Sharapova SO et al. (2020). "The Clinical and Genetic Spectrum of 82 Patients With RAG Deficiency Including a c.256_257delAA Founder Variant in Slavic Countries." Frontiers in immunology, 11()  [[3]](https://pubmed.ncbi.nlm.nih.gov/25502423/) Moens LN et al. (2014). "Diagnostics of primary immunodeficiency diseases: a sequencing capture approach." PloS one, 9(12)  [[4]](https://pubmed.ncbi.nlm.nih.gov/39026935/) Volodashchik TP et al. (2024). "Infant with diffuse large B-cell lymphoma identified postmortem with homozygous founder Slavic RAG1 variant: a case report and literature review." Frontiers in pediatrics, 12()  [[5]](https://pubmed.ncbi.nlm.nih.gov/31031743/) Cifaldi C et al. (2019). "Targeted NGS Platforms for Genetic Screening and Gene Discovery in Primary Immunodeficiencies." Frontiers in immunology, 10()  [[6]](https://pubmed.ncbi.nlm.nih.gov/23085344/) Sharapova SO et al. (2013). "Late-onset combined immune deficiency associated to skin granuloma due to heterozygous compound mutations in RAG1 gene in a 14 years old male." Human immunology, 74(1)  [[7]](https://pubmed.ncbi.nlm.nih.gov/25516070/) Buchbinder D et al. (2015). "Identification of patients with RAG mutations previously diagnosed with common variable immunodeficiency disorders." Journal of clinical immunology, 35(2)  [[8](https://pubmed.ncbi.nlm.nih.gov/26596586/)] Sharapova SO et al. (2016). "Molecular Characteristics, Clinical and Immunologic Manifestations of 11 Children with Omenn Syndrome in East Slavs (Russia, Belarus, Ukraine)." Journal of clinical immunology, 36(1)  [[9]](https://pubmed.ncbi.nlm.nih.gov/32000930/) Tallar M et al. (2020). "Omenn Syndrome Identified by Newborn Screening." Clinics in perinatology, 47(1)  [[10]](https://pubmed.ncbi.nlm.nih.gov/29856523/) C. Maas et al. (2018). "EBV‐positive B‐cell lymphoma manifestation of the liver in an infant with RAG1 severe combined immunodeficiency disease" Pediatric Blood & Cancer, 65  [[11]](https://pubmed.ncbi.nlm.nih.gov/21185069/) Pessach IM et al. (2011). "Induced pluripotent stem cells: a novel frontier in the study of human primary immunodeficiencies." The Journal of allergy and clinical immunology, 127(6)  [[12]](https://pubmed.ncbi.nlm.nih.gov/39235410/) Bakardjieva M et al. (2024). "Tviblindi algorithm identifies branching developmental trajectories of human B-cell development and describes abnormalities in RAG-1 and WAS patients." European journal of immunology, 54(12)  [[13]](https://pubmed.ncbi.nlm.nih.gov/39720732/) Chen X et al. (2024). "Case report: Identification of a Chinese patient with RAG1 mutations initially presenting as autoimmune hemolytic anemia." Frontiers in immunology, 15()  [[14]](https://pubmed.ncbi.nlm.nih.gov/24290284/) Lee YN et al. (2014). "A systematic analysis of recombination activity and genotype-phenotype correlation in human recombination-activating gene 1 deficiency." The Journal of allergy and clinical immunology, 133(4)  [[15]](https://pubmed.ncbi.nlm.nih.gov/30877075/) Farmer JR et al. (2019). "Outcomes and Treatment Strategies for Autoimmunity and Hyperinflammation in Patients with RAG Deficiency." The journal of allergy and clinical immunology. In practice, 7(6) |
|  | Evaluation  **Summarization Accuracy & Completeness:**  **Hallucination Rate:**  **Readability & Usability:**  **Literature Coverage & Relevance:**  **Time Efficiency:** | |
| **4** | The recombination-activating gene 1 (RAG1) is critical for V(D)J recombination, a process essential for the development of functional B and T lymphocytes. Pathogenic variants in RAG1 can result in severe immunodeficiency disorders such as severe combined immunodeficiency (SCID) and Omenn syndrome. The c.256_257del variant in RAG1 is a frameshift mutation that likely leads to a truncated or non-functional protein, impairing immune system development. Although no direct studies on this specific variant are available, research on similar RAG1 mutations suggests a severe phenotype, often presenting as early-onset immunodeficiency with high susceptibility to infections and possible lymphoproliferative complications [1]. Cases of lymphoma have been reported in individuals with RAG1 mutations, further emphasizing the potential impact of this variant [2]. Current classification systems would likely categorize c.256_257del as a pathogenic variant due to its predicted loss-of-function nature. The frequency of this specific deletion remains unknown, but loss-of-function mutations in RAG1 are generally rare. Functional studies on RAG1-deficient models indicate that even partial loss of function can severely impair T and B cell development, leading to immune system failure [3]. Due to the absence of direct studies on c.256_257del, functional assays and clinical correlation are necessary to confirm its precise pathogenicity. Genetic counseling and early intervention, potentially including hematopoietic stem cell transplantation, are recommended for individuals harboring severe RAG1 mutations. Further research is necessary to determine genotype-phenotype correlations and therapeutic strategies. | [1] Volodashchik TP, Polyakova EA. "Infant with diffuse large B-cell lymphoma identified postmortem with homozygous founder Slavic RAG1 variant: a case report and literature review." Front Pediatr. 2024. <https://www.frontiersin.org/articles/10.3389/fped.2024.1415020/full>  [2] Volodashchik T, Polyakova E, Mikhalevskaya T. "Diffuse large B-cell lymphoma in 1-year male with founder Slavic RAG1 mutation: case report." Sci Direct. 2023. <https://www.sciencedirect.com/science/article/pii/S1521661623002012>  [3] Brauer PM, Pessach IM, Clarke E. "Modeling altered T-cell development with induced pluripotent stem cells from patients with RAG1-dependent immune deficiencies." Blood. 2016. <https://ashpublications.org/blood/article-abstract/128/6/783/35760> |
|  | Evaluation  **Summarization Accuracy & Completeness:**  **Hallucination Rate:**  **Readability & Usability:**  **Literature Coverage & Relevance:**  **Time Efficiency:** | |
| **5** | The RAG1 gene, or Recombination Activating Gene 1, plays a crucial role in the development of the adaptive immune system by facilitating V(D)J recombination, a process essential for generating a diverse repertoire of immunoglobulins and T-cell receptors in developing B and T lymphocytes. This gene encodes a protein that, together with RAG2, forms the RAG complex, which initiates V(D)J recombination by introducing double-strand breaks at specific recombination signal sequences (RSS) adjacent to the coding segments of immunoglobulin and T-cell receptor genes. The RAG1 protein is responsible for DNA binding and cleavage activities within this complex, recognizing and binding to the conserved RSS and catalyzing the necessary DNA cleavage to enable subsequent ligation of the V, D, and J segments. This process is critical for the maturation of functional B and T cells, which are essential for immune response and defense against pathogens. Defects in RAG1 can lead to various immunodeficiencies, including severe combined immunodeficiency (SCID) and Omenn syndrome, characterized by the absence or dysfunction of T and B cells, resulting in severe infections and immune dysregulation. The RAG1 gene is highly conserved across species, reflecting its fundamental role in immune system development and function.  The variant c.256_257del in the RAG1 gene is a dinucleotide deletion that results in a frameshift mutation, leading to a premature stop codon and a truncated protein (p.Lys86ValfsTer33). This variant has been identified in patients with severe combined immunodeficiency (SCID) and is particularly prevalent in Slavic populations, where it is often found in a homozygous state or as a compound heterozygous mutation. The c.256_257del variant significantly impairs the function of the RAG1 protein, affecting its ability to facilitate V(D)J recombination and leading to a profound deficiency in T and B cell development. Patients carrying this variant typically present with clinical features such as recurrent infections, lymphopenia, and reduced immunoglobulin levels, highlighting the severe impact of this mutation on immune system function. The high frequency of this variant in specific populations suggests a founder effect, and its clinical manifestations underscore the importance of genetic screening and early diagnosis in managing immune deficiencies. Further investigation is needed to fully understand the molecular mechanisms underlying the pathogenicity of this variant and to develop targeted therapeutic strategies for affected individuals. | [1] Mutation c.256_257delAA in RAG1 Gene in Polish Children with Severe Combined Immunodeficiency: Diversity of Clinical Manifestations - PubMed [1](https://pubmed.ncbi.nlm.nih.gov/28083621/)  [2] The Clinical and Genetic Spectrum of 82 Patients With RAG Deficiency Including a c.256_257delAA Founder Variant in Slavic Countries - PubMed [2](https://pubmed.ncbi.nlm.nih.gov/32655540/)  [3] A variant of RAG1 gene identified in severe combined immunodeficiency: a case report - PMC [3](https://pmc.ncbi.nlm.nih.gov/articles/PMC9896705/)  [4] Late-onset combined immune deficiency associated to skin granuloma due to heterozygous compound mutations in RAG1 gene in a 14 years old male - PubMed [4](https://pubmed.ncbi.nlm.nih.gov/23085344/)  [5] RAG1 gene: MedlinePlus Genetics [5](https://medlineplus.gov/genetics/gene/rag1/) |
|  | Evaluation  **Summarization Accuracy & Completeness:**  **Hallucination Rate:**  **Readability & Usability:**  **Literature Coverage & Relevance:**  **Time Efficiency:** | |
