## Supplementary material for "Benchmarking generative AI tools for literature retrieval and summarization in genomic variant interpretation": SERPINA1 c.1096G_A.docx

| **Tool** | **Summary** | **References** |
| --- | --- | --- |
| **1** | The genomic variant c.1096G>A p.Glu366Lys, also known as rs28929474, is located in the SERPINA1 gene, which encodes the alpha-1-antitrypsin (AAT) protein. AAT is a serine protease inhibitor primarily produced in the liver, playing a crucial role in protecting lung tissue from enzymes released by inflammatory cells, particularly neutrophil elastase. Deficiency in AAT can lead to uncontrolled enzyme activity, resulting in lung and liver diseases.  The rs28929474 variant is associated with the Z allele of the SERPINA1 gene and is a known pathogenic variant that can lead to alpha-1 antitrypsin deficiency (A1AD). Individuals carrying this variant in a heterozygous state have been found to suffer from common variable immune deficiency (CVID), although they displayed normal AAT serum levels [1]. This variant is also associated with decreased lung function and chronic obstructive pulmonary disease (COPD), particularly in smokers. Homozygosity for the Z allele (PiZZ genotype) is consistently linked with emphysema and reduced lung function, while heterozygotes also show significantly lower pulmonary function [2].  In addition to its pulmonary implications, the rs28929474 variant has been associated with liver disease, as A1AD can lead to liver dysfunction due to the accumulation of the aberrant AAT protein in hepatocytes [4]. The variant has been identified as a missense mutation, resulting in the substitution of glutamic acid with lysine in the AAT protein, which affects its folding and function.  In summary, the c.1096G>A p.Glu366Lys rs28929474 variant in the SERPINA1 gene is a pathogenic variant with significant clinical implications for lung and liver health. It is associated with A1AD, which can manifest as lung diseases such as emphysema and COPD, particularly in the context of environmental factors like smoking, as well as liver disease. ClinVar submissions corroborate the pathogenic nature of this variant, with multiple entries supporting its association with disease phenotypes. | [[1]](https://pubmed.ncbi.nlm.nih.gov/38791420/) Sarrou S et al. (2024). "Heterozygous SERPINA1 Defects and Their Impact on Clinical Manifestations of Patients with Predominantly Antibody Deficiencies." International journal of molecular sciences, 25(10)  [[2]](https://pubmed.ncbi.nlm.nih.gov/30068317/) Li X et al. (2018). "Genome-wide association study of lung function and clinical implication in heavy smokers." BMC medical genetics, 19(1)  [[3]](https://pubmed.ncbi.nlm.nih.gov/28053854/) de Seynes C et al. (2017). "Identification of a novel alpha1-antitrypsin variant." Respiratory medicine case reports, 20()  [[4]](https://pubmed.ncbi.nlm.nih.gov/28887542/) Prins BP et al. (2017). "Genome-wide analysis of health-related biomarkers in the UK Household Longitudinal Study reveals novel associations." Scientific reports, 7(1)  [[5]](https://pubmed.ncbi.nlm.nih.gov/28121484/) Feng L et al. (2017). "Ubiquitin ligase SYVN1/HRD1 facilitates degradation of the SERPINA1 Z variant/alpha-1-antitrypsin Z variant via SQSTM1/p62-dependent selective autophagy." Autophagy, 13(4)  [[6]](https://pubmed.ncbi.nlm.nih.gov/38388492/) Ferrarotti I et al. (2024). "Rare variants in alpha 1 antitrypsin deficiency: a systematic literature review." Orphanet journal of rare diseases, 19(1)  [[7]](https://pubmed.ncbi.nlm.nih.gov/38523412/) Kamuda K et al. (2024). "A novel pathological mutant reveals the role of torsional flexibility in the serpin breach in adoption of an aggregation-prone intermediate." The FEBS journal, 291(13)  [[8]](https://pubmed.ncbi.nlm.nih.gov/21138453/) Mihalache F et al. (2011). "Heterozygosity for the alpha1-antitrypsin Z allele may confer genetic risk of cholangiocarcinoma." Alimentary pharmacology & therapeutics, 33(3)  [[9]](https://pubmed.ncbi.nlm.nih.gov/30351692/) Smirnova EL et al. (2016). "Forecasting role of genetic markers in outlining features of pneumoconiosis course in post-contact period." Meditsina truda i promyshlennaia ekologiia, (11)  [[10]](https://pubmed.ncbi.nlm.nih.gov/38723785/) Ramos-Diaz R et al. (2024). "Real-time PCR detection of PI*S and PI*Z alleles of SERPINA1 gene using SYBR green." Gene, 921()  [[11]](https://pubmed.ncbi.nlm.nih.gov/39881831/) Ivanova AA et al. (2024). "Detection of Major Mutations in CFTR, SERPINA1, HFE Genes in Benign Unconjugated Hyperbilirubinemia Phenotype." Sovremennye tekhnologii v meditsine, 16(4)  [[12]](https://pubmed.ncbi.nlm.nih.gov/23696881/) Wood AR et al. (2013). "Imputation of variants from the 1000 Genomes Project modestly improves known associations and can identify low-frequency variant-phenotype associations undetected by HapMap based imputation." PloS one, 8(5)  [[13]](https://pubmed.ncbi.nlm.nih.gov/34002692/) Tumpara S et al. (2021). "Polymerization of misfolded Z alpha-1 antitrypsin protein lowers CX3CR1 expression in human PBMCs." eLife, 10()  [[14]](https://pubmed.ncbi.nlm.nih.gov/37685316/) Fernandez-Gomez B et al. (2023). "Utility of the Serum Protein Electrophoresis in the Opportunistic Screening for the Deficiency of Alpha-1 Antitrypsin." Diagnostics (Basel, Switzerland), 13(17)  [[15]](https://pubmed.ncbi.nlm.nih.gov/25454901/) Perez-Rubio G et al. (2015). "Prevalence of alpha-1 antitrypsin high-risk variants in Mexican mestizo population and their association with lung function values." Archivos de bronconeumologia, 51(2) |
|  | Evaluation  **Summarization Accuracy & Completeness:**  **Hallucination Rate:**  **Readability & Usability:**  **Literature Coverage & Relevance:**  **Time Efficiency:** | |
| **2** | The SERPINA1 gene encodes alpha-1 antitrypsin (AAT), a serine protease inhibitor primarily synthesized in the liver and secreted into the bloodstream[4](https://www.ncbi.nlm.nih.gov/pmc/articles/PMC6444226/). AAT's main function is to inhibit neutrophil elastase, an enzyme that can break down lung tissue[4](https://www.ncbi.nlm.nih.gov/pmc/articles/PMC6444226/). Deficiency in AAT, often due to genetic variants, can lead to excessive elastase activity, resulting in pulmonary emphysema and, in some cases, liver disease[5](https://pubmed.ncbi.nlm.nih.gov/31661293/). The c.1096G>A variant in the SERPINA1 gene is a single nucleotide variant (SNV) that requires computational tools to predict its effects on AAT protein structure and function[1](https://www.ncbi.nlm.nih.gov/pmc/articles/PMC10970068/). Determining the clinical significance of SERPINA1 variants, like c.1096G>A, is essential for tailoring treatment options for individuals with alpha-1 antitrypsin deficiency (AATD)[1](https://www.ncbi.nlm.nih.gov/pmc/articles/PMC10970068/). At this time, there are no specific peer-reviewed publications detailing the classification, nomenclature, frequency, or precise impact of the c.1096G>A variant. As such, its clinical significance remains uncertain, and further investigation is warranted to determine its effect on AAT function and associated disease risk. Further research, potentially including bioinformatic analyses, is needed to fully elucidate the functional consequences of this variant[1](https://www.ncbi.nlm.nih.gov/pmc/articles/PMC10970068/). | [1](https://www.ncbi.nlm.nih.gov/pmc/articles/PMC10970068/) PMC10970068  [2](https://www.ncbi.nlm.nih.gov/pmc/articles/PMC8048251/) PMC8048251  [3](https://www.semanticscholar.org/paper/9e5b866e8561ccc8c91a7e469f036f2b2381dda4) PMC6444226  [4](https://www.ncbi.nlm.nih.gov/pmc/articles/PMC6444226/) PMC11074302  [5](https://pubmed.ncbi.nlm.nih.gov/31661293/) PMC3691238  [6](https://pmc.ncbi.nlm.nih.gov/articles/PMC11074302/) PMC11074302  [7](https://pmc.ncbi.nlm.nih.gov/articles/PMC6103990/) PMC6103990 |
|  | Evaluation  **Summarization Accuracy & Completeness:**  **Hallucination Rate:**  **Readability & Usability:**  **Literature Coverage & Relevance:**  **Time Efficiency:** | |
| **3** | The SERPINA1 gene encodes alpha-1-antitrypsin (AAT), a serine protease inhibitor that primarily inhibits neutrophil elastase, protecting the lungs from proteolytic damage. The c.1096G>A variant in SERPINA1, commonly known as the Z allele, results in a glutamic acid to lysine substitution at position 366 (p.Glu366Lys) in the AAT protein. This variant is one of the most common pathogenic mutations associated with alpha-1-antitrypsin deficiency (AATD), which can lead to emphysema and liver disease. The Z allele causes AAT to misfold and polymerize within hepatocytes, reducing its secretion into the bloodstream and leading to a significant decrease in AAT levels. Homozygosity for the Z allele (PI*ZZ genotype) is particularly detrimental, resulting in severe AATD with only 10-15% of normal AAT levels. The frequency of the Z allele in the general population is up to 1.8%, highlighting its significance in genetic screening for AATD. The pathogenic nature of this variant is well-documented, supported by multiple lines of computational and clinical evidence. It is classified as pathogenic according to the American College of Medical Genetics and Genomics (ACMG) guidelines. The clinical impact of the Z allele underscores the importance of genetic testing and counseling for individuals at risk for AATD and their families. Further investigation is recommended to understand the full spectrum of clinical outcomes associated with this variant, especially in compound heterozygous states with other SERPINA1 mutations. | [1] OMIM. SERPINA1 gene. Available at: <https://omim.org/entry/107400>  [2] ClinVar. SERPINA1 c.1096G>A. Available at: <https://www.ncbi.nlm.nih.gov/clinvar/RCV000148877/>  [3] MedlinePlus. SERPINA1 gene. Available at: <https://medlineplus.gov/genetics/gene/serpina1/> |
|  | Evaluation  **Summarization Accuracy & Completeness:**  **Hallucination Rate:**  **Readability & Usability:**  **Literature Coverage & Relevance:**  **Time Efficiency:** | |
| **4** | The SERPINA1 gene encodes alpha-1 antitrypsin (AAT), a serine protease inhibitor that primarily regulates neutrophil elastase activity in the lungs, preventing excessive tissue damage. Pathogenic variants in SERPINA1 are associated with alpha-1 antitrypsin deficiency (AATD), a condition that predisposes individuals to chronic obstructive pulmonary disease (COPD) and liver disease due to the misfolding and accumulation of defective AAT protein. The c.1096G>A variant in SERPINA1 results in an amino acid substitution that may impact protein function, but its clinical significance remains incompletely defined. A study identified this variant in a patient with a cystic fibrosis-like phenotype, suggesting a potential role in pulmonary pathology, though further research is needed to establish its pathogenicity [1]. Broader sequencing studies of SERPINA1 variants highlight the genetic heterogeneity of AATD and emphasize the need for comprehensive functional assays to determine the consequences of rare variants [2,3]. In particular, the association of heterozygous SERPINA1 variants with antibody deficiencies suggests that even single copies of pathogenic alleles could contribute to immune dysregulation [4]. While established pathogenic alleles like Z (Glu342Lys) and S (Glu264Val) are well characterized, the impact of rarer mutations such as c.1096G>A remains uncertain, necessitating further biochemical and clinical studies to assess their contribution to AATD-related phenotypes. Given the variability in phenotypic expression and potential modifier effects from other genetic or environmental factors, additional investigations using computational modeling and functional validation studies are recommended to clarify the clinical relevance of c.1096G>A [5]. | 1. Ramos MD, Trujillano D, Olivar R, Sotillo F. *Extensive sequence analysis of CFTR, SCNN1A, SCNN1B, SCNN1G and SERPINA1 suggests an oligogenic basis for cystic fibrosis-like phenotypes.* [DOI:10.1111/cge.12234](https://app.scholarai.io/paper?paper_id=DOI:10.1111/cge.12234&original_url=https%3A%2F%2Fonlinelibrary.wiley.com%2Fdoi%2Fabs%2F10.1111%2Fcge.12234) 2. Gupta N, Gaudreault N, Thériault S. *Granularity of SERPINA1 alleles by DNA sequencing in CanCOLD.* [DOI:10.1183/13993003.00958-2020](https://publications.ersnet.org/content/erj/56/4/2000958.abstract) 3. Ferrarotti I, Wencker M. *Rare variants in alpha 1 antitrypsin deficiency: a systematic literature review.* [DOI:10.1186/s13023-024-03069-1](https://link.springer.com/article/10.1186/s13023-024-03069-1) 4. Sarrou S, Voulgaridi I, Fousika A, Dadouli K. *Heterozygous SERPINA1 defects and their impact on clinical manifestations of patients with predominantly antibody deficiencies.* [DOI:10.3390/ijms25105382](https://app.scholarai.io/paper?paper_id=DOI:10.3390/ijms25105382&original_url=https%3A%2F%2Fwww.mdpi.com%2F1422-0067%2F25%2F10%2F5382) 5. Mróz J, Pelc M, Mitusińska K. *Computational Tools to Assist in Analyzing Effects of the SERPINA1 Gene Variation on Alpha-1 Antitrypsin (AAT).* [DOI:10.3390/genes15030340](https://app.scholarai.io/paper?paper_id=DOI:10.3390/genes15030340&original_url=https%3A%2F%2Fwww.mdpi.com%2F2073-4425%2F15%2F3%2F340) |
|  | Evaluation  **Summarization Accuracy & Completeness:**  **Hallucination Rate:**  **Readability & Usability:**  **Literature Coverage & Relevance:**  **Time Efficiency:** | |
| **5** | The SERPINA1 gene encodes alpha-1 antitrypsin (AAT), a serine protease inhibitor primarily synthesized in the liver. AAT's main function is to protect tissues from enzymes released by inflammatory cells, notably neutrophil elastase, which can degrade elastin in the lungs if unregulated. The c.1096G>A variant in SERPINA1 results in a glutamic acid to lysine substitution at position 366 (p.Glu366Lys), commonly referred to as the Z allele. This mutation leads to misfolding of the AAT protein, causing its accumulation in hepatocytes and reduced secretion into the bloodstream. Consequently, individuals with the Z allele have decreased AAT levels, impairing elastase inhibition and increasing the risk of developing chronic obstructive pulmonary disease (COPD) and liver diseases such as cirrhosis. The Z allele is prevalent in approximately 1-2% of individuals of European descent, with homozygous carriers (ZZ genotype) at a higher risk for these conditions. Heterozygous carriers (MZ genotype) may also have an elevated risk, particularly if exposed to environmental risk factors like smoking. Clinical management includes regular monitoring of lung and liver function, lifestyle modifications to reduce risk factors, and, in severe cases, augmentation therapy with purified AAT. | <https://medlineplus.gov/genetics/gene/serpina1/>  <https://journals.sagepub.com/doi/10.1177/20406223211015954?icid=int.sj-full-text.citing-articles.58>  <https://www.atsjournals.org/doi/10.1164/rccm.201904-0769OC>  <https://translational-medicine.biomedcentral.com/articles/10.1186/s12967-015-0585-y>  <https://www.jmdjournal.org/article/S1525-1578%2815%2900151-8/pdf>  <https://www.ncbi.nlm.nih.gov/clinvar/RCV000148877/>  <https://www.ncbi.nlm.nih.gov/clinvar/RCV000768543/>  <https://databases.lovd.nl/shared/variants/SERPINA1?search_position_c_end=1096&search_position_c_end_intron=&search_position_c_start=1096&search_position_c_start_intron=&search_transcriptid=00000182&search_vot_clean_dna_change=%3D%22c.1096G%3EA%22>  <https://my.clevelandclinic.org/health/diseases/21175-alpha-1-antitrypsin-deficiency>  <https://clinvarminer.genetics.utah.edu/variants-by-gene/SERPINA1/condition/Alpha-1-antitrypsin%20deficiency>  <https://www.genecards.org/cgi-bin/carddisp.pl?gene=SERPINA1>  <https://en.wikipedia.org/wiki/Alpha-1_antitrypsin>  <https://clevelandcliniclabs.com/wp-content/uploads/2020/02/Alpha-1-Antitrypsin-Technical-Brief-Cleveland-Clinic-Laboratories.pdf>  <https://www.atsjournals.org/doi/10.1164/rccm.201911-2209ED>  <https://mgend.ncgm.go.jp/variant/info/SNV%7Chg19%7Cchr14%3A94844947-94844947%7CgC%3ET> |
|  | Evaluation  **Summarization Accuracy & Completeness:**  **Hallucination Rate:**  **Readability & Usability:**  **Literature Coverage & Relevance:**  **Time Efficiency:** | |
