## Supplementary material for "Benchmarking generative AI tools for literature retrieval and summarization in genomic variant interpretation": SLC24A5 rs1426654.docx

| **Tool** | **Summary** | **References** |
| --- | --- | --- |
| **1** | The SLC24A5 gene encodes a protein known as sodium/potassium/calcium exchanger 5 (NCKX5), which is integral to melanin production in melanocytes. A notable single nucleotide polymorphism (SNP) within this gene, rs1426654, results in an amino acid substitution at position 111, changing alanine (Ala) to threonine (Thr). This variant has been identified as a significant determinant of human skin pigmentation differences. The derived threonine allele (Ala111Thr) is prevalent in European populations, with frequencies approaching fixation, while the ancestral alanine form is predominant in Sub-Saharan African, East Asian, and Indigenous American populations. This single nucleotide change accounts for approximately 25–38% of the variation in skin melanin index between individuals of African and European ancestry. Functional studies have demonstrated that NCKX5 is localized to the trans-Golgi network in melanocytes and plays a crucial role in melanin synthesis. Disruption or alteration of NCKX5 activity, such as the Ala111Thr substitution, significantly impacts melanin production, leading to observable differences in pigmentation. The high prevalence of the threonine allele in European populations suggests strong positive selection, likely due to adaptive advantages in regions with lower ultraviolet radiation, facilitating vitamin D synthesis. In contrast, the alanine variant's predominance in darker-skinned populations underscores its evolutionary role in providing protection against intense UV radiation. These findings highlight the pivotal role of the rs1426654 variant in the genetic architecture of human skin color variation. | <https://en.wikipedia.org/wiki/Sodium/potassium/calcium_exchanger_5>  <https://www.researchgate.net/publication/51411413_A_study_of_a_single_variant_allele_rs1426654_of_the_pigmentation-related_gene_SLC24A5_in_Greek_subjects>  <https://pmc.ncbi.nlm.nih.gov/articles/PMC3820762/>  <https://www.researchgate.net/publication/51411413_A_study_of_a_single_variant_allele_rs1426654_of_the_pigmentation-related_gene_SLC24A5_in_Greek_subjects>  <https://www.nature.com/articles/s41598-024-68437-0>  <https://elifesciences.org/articles/77514>  <https://bmcgenomdata.biomedcentral.com/articles/10.1186/s12863-019-0765-5>  <https://academic.oup.com/gbe/article/11/4/1066/5416147>  <https://www.nature.com/articles/s41467-018-08147-0>  <https://pmc.ncbi.nlm.nih.gov/articles/PMC3820762/>  <https://onlinelibrary.wiley.com/doi/10.1111/j.1600-0625.2008.00758.x>  <https://onlinelibrary.wiley.com/doi/full/10.1111/j.1755-148X.2008.00456.x>  <https://pmc.ncbi.nlm.nih.gov/articles/PMC2674805/>  <https://pubmed.ncbi.nlm.nih.gov/18637132/>  <https://www.science.org/doi/10.1126/science.1116238>  <https://www.science.org/doi/10.1126/science.aan8433>  <https://journals.plos.org/plosone/article?id=10.1371%2Fjournal.pone.0083926>  <https://www.infino.me/snp/rs1426654/> |
|  | Evaluation  **Summarization Accuracy & Completeness:**  **Hallucination Rate:**  **Readability & Usability:**  **Literature Coverage & Relevance:**  **Time Efficiency:** | |
| **2** | The SLC24A5 gene, located on chromosome 15q21.1, encodes the NCKX5 protein, a member of the potassium-dependent sodium-calcium exchanger family. This gene plays a crucial role in human pigmentation by regulating melanin production in melanocytes. The variant rs1426654, a non-synonymous single nucleotide polymorphism (SNP) in the third exon of SLC24A5, results in an alanine-to-threonine substitution at amino acid position 111 (A111T). This variant is strongly associated with lighter skin pigmentation, particularly in European populations, where it reaches near fixation. The rs1426654 allele frequency shows significant differences between European and non-European populations, with Europeans exhibiting a much higher frequency of the threonine allele. This variant accounts for 25-38% of the skin color variation between Europeans and West Africans, highlighting its substantial impact on human pigmentation. The lighter skin tone associated with rs1426654 is believed to have arisen from a single origin, with the allele being monophyletic and occurring on a common haplotype background in both South Asian and West Eurasian populations. Functional studies have demonstrated that the NCKX5 protein, encoded by SLC24A5, is essential for melanogenesis, and the rs1426654 variant directly influences this process. The alanine variant, prevalent in West Africans and East Asians, is associated with darker skin pigmentation. The strong selective pressure on SLC24A5 is evident from the reduced heterozygosity and high pairwise FST values surrounding the gene, indicating it is one of the most strongly selected regions of the human genome. While the exact mechanisms by which rs1426654 affects pigmentation are still under investigation, it is clear that this variant plays a pivotal role in determining skin color. Further research is needed to fully elucidate the molecular pathways involved and to understand the evolutionary forces that have shaped the distribution of this allele across different populations. | [1] Lamason et al. (2005)  [2] Stokowski et al. (2007)  [3] Quillen et al. (2008)  [4] Pemberton et al. (2013)  [5] Basu Mallick et al. (2013) |
|  | Evaluation  **Summarization Accuracy & Completeness:**  **Hallucination Rate:**  **Readability & Usability:**  **Literature Coverage & Relevance:**  **Time Efficiency:** | |
| **3** | The SLC24A5 gene encodes a potassium-dependent sodium/calcium exchanger involved in melanosome ion transport and plays a significant role in human pigmentation. The rs1426654 variant is a non-synonymous single nucleotide polymorphism (SNP) resulting in an alanine-to-threonine substitution at codon 111 (A111T), located in exon 3. This SNP exhibits a strong allelic differentiation between populations, with the derived A allele nearly fixed in European populations and the ancestral G allele predominating in African and East Asian populations. Studies have demonstrated that rs1426654 is a major contributor to skin pigmentation variation, explaining approximately 25–38% of melanin index differences among global populations [1]. Genetic analysis suggests that the rs1426654 A allele underwent strong positive selection in non-African populations, potentially linked to adaptation to reduced ultraviolet radiation exposure in higher latitudes [2]. Functional studies indicate that the A allele decreases melanin production, leading to lighter skin pigmentation, which has been confirmed by association studies in multiple ethnic groups, including South Asians, Europeans, and Native Americans [3]. The allele frequency distribution and identity-by-descent analysis suggest that the A allele was introduced into South Asian populations through gene flow from European ancestors rather than an independent mutation [4]. The variant has also been studied in forensic genetics for phenotype prediction models and has been incorporated into genetic ancestry analysis due to its high predictive value for skin color [5]. Moreover, emerging research has explored its interaction with other pigmentation genes such as MC1R and TYRP1, reinforcing its role in polygenic skin pigmentation determination [6]. While the rs1426654 variant is primarily associated with pigmentation, some studies have suggested potential pleiotropic effects, including associations with metabolic traits, though these findings remain inconclusive [7]. Given its well-established role in pigmentation variation and evolutionary selection, rs1426654 remains one of the most studied SNPs in human pigmentation genetics, with implications in anthropology, forensic science, and medical genetics. Further research is needed to fully elucidate any potential functional consequences beyond pigmentation. | [1] Dimisianos et al., 2009 (DOI:10.1111/j.1600-0625.2008.00758.x);  [2] Mallick et al., 2013 (PLoS Genetics);  [3] Rivera-Paredez et al., 2024 (DOI:10.1038/s41598-024-68437-0);  [4] Mukherjee et al., 2013 (Springer);  [5] de Araújo Lima et al., 2015 (ScienceDirect);  [6] Reis et al., 2020 (Springer);  [7] Sarkar & Nandineni, 2018 (DOI:10.1002/ajhb.23068). |
|  | Evaluation  **Summarization Accuracy & Completeness:**  **Hallucination Rate:**  **Readability & Usability:**  **Literature Coverage & Relevance:**  **Time Efficiency:** | |
| **4** | The gene SLC24A5 encodes NCKX5, a potassium-dependent sodium/calcium exchanger 5, which is involved in the maturation of melanosomes, though its precise function remains unclear[3](https://pubmed.ncbi.nlm.nih.gov/34870899/). SLC24A5 is known to play a key role in human pigmentation[2](https://pubmed.ncbi.nlm.nih.gov/18637132/). Certain mutations in SLC24A5 can result in oculocutaneous albinism type 6 (OCA6), a non-syndromic type of albinism characterized by distinct ocular symptoms and variable cutaneous hypopigmentation[3](https://pubmed.ncbi.nlm.nih.gov/34870899/)[5](https://pubmed.ncbi.nlm.nih.gov/33504991/). Genetic analysis of a Japanese patient with OCA6 revealed compound heterozygous variants in SLC24A5, c.590 + 1dupG, and c.598G>A (p.G200R)[3](https://pubmed.ncbi.nlm.nih.gov/34870899/). rs1426654 is a variant allele of the SLC24A5 gene[2](https://pubmed.ncbi.nlm.nih.gov/18637132/). A study of Greek subjects showed a high prevalence of the Thr111 allele, even among those with darker skin pigmentation[2](https://pubmed.ncbi.nlm.nih.gov/18637132/). Research has shown significant genetic variations between groups in women of African descent with melasma[6](https://www.ncbi.nlm.nih.gov/pmc/articles/PMC11818098/). Further investigation is warranted to fully elucidate the functional significance of SLC24A5 variants and their impact on pigmentation and related disorders[3](https://pubmed.ncbi.nlm.nih.gov/34870899/). | [2](https://pubmed.ncbi.nlm.nih.gov/18637132/) A study of a single variant allele (rs1426654) of the pigmentation‐related gene SLC24A5 in Greek subjects. *J Eur Acad Dermatol Venereol*. 2009 Feb;23(2):223-4.  [3](https://pubmed.ncbi.nlm.nih.gov/34870899/) Impact of a SLC24A5 variant on the retinal pigment epithelium of a Japanese patient with oculocutaneous albinism type 6. *Pigment Cell Melanoma Res*. 2022 Jan;35(1):184-188.  [6](https://www.ncbi.nlm.nih.gov/pmc/articles/PMC11818098/) Association of Genetic Polymorphisms in SLC45A2, TYR, HERC2, and SLC24A in African Women with Melasma: A Pilot Study. *J Cosmet Dermatol*. 2024 Jan 29.   1. <https://www.semanticscholar.org/paper/adda9dfb9ad00ec3d701c6ad65331f087118a929> 2. <https://pubmed.ncbi.nlm.nih.gov/18637132/> 3. <https://pubmed.ncbi.nlm.nih.gov/34870899/> 4. <https://www.ncbi.nlm.nih.gov/pmc/articles/PMC5555483/> 5. <https://pubmed.ncbi.nlm.nih.gov/33504991/> 6. <https://www.ncbi.nlm.nih.gov/pmc/articles/PMC11818098/> 7. <https://www.ncbi.nlm.nih.gov/pmc/articles/PMC11286937/> 8. <https://www.semanticscholar.org/paper/87902e7597b9641b450c94e6c020cacab75076e8> |
|  | Evaluation  **Summarization Accuracy & Completeness:**  **Hallucination Rate:**  **Readability & Usability:**  **Literature Coverage & Relevance:**  **Time Efficiency:** | |
| **5** | The genomic variant c.331A>G p.Thr111Ala, also known as rs1426654, is located on the SLC24A5 gene, which plays a crucial role in the regulation of melanin production, a pigment responsible for skin, hair, and eye color. This missense variant results in the substitution of threonine with alanine at the 111th amino acid position of the protein encoded by SLC24A5. The SLC24A5 gene encodes a membrane-associated transporter protein that is involved in melanogenesis, the process by which melanin is produced in melanocytes [2].  The rs1426654 variant has been extensively studied due to its association with pigmentation variation among different populations. It has been observed that the G allele frequency of this SNP decreases gradually from Africa to Europe, suggesting a selection pressure in favor of the A allele in regions with lower ultraviolet (UV) radiation intensity. This pattern is consistent with the hypothesis that lighter skin pigmentation evolved to optimize vitamin D synthesis in areas of lower solar incidence [1].  In addition to its role in pigmentation, the rs1426654 variant has been implicated in the risk of melanoma. Studies have shown that the AA genotype is associated with fair skin and light eyes, and the presence of homozygous genotypes of this SNP (AA) has been associated with an increased risk for the development of melanoma. This association remains significant even after adjusting for other risk factors such as ancestry, gender, age, eye, hair, and skin color, and the number of nevi [1].  The functional impact of the rs1426654 variant has been demonstrated in experimental studies. In melanocyte cultures, the homozygous GG genotype leads to an increase in SLC24A5 gene transcripts, which in turn increases tyrosinase activity and melanin production. This variant was first described in zebrafish as responsible for the golden phenotype due to a delay in melanin production during embryonic development [1].  The rs1426654 variant has also been characterized through resequencing studies, which have shown that the exons of SLC24A5 are highly conserved in humans, with rs1426654 being the only non-synonymous SNP detected in the exonic regions. This highlights the potential functional importance of this particular variant [2].  In summary, the c.331A>G p.Thr111Ala rs1426654 variant on the SLC24A5 gene is a missense variant associated with pigmentation differences and melanoma risk. Its prevalence varies geographically, reflecting historical selection pressures related to UV radiation exposure and vitamin D synthesis. Despite its association with certain phenotypes and disease risks, this variant is classified as benign in ClinVar, indicating that it is not typically associated with severe health consequences. | [[1]](https://pubmed.ncbi.nlm.nih.gov/33167923/) Reis LB et al. (2020). "Skin pigmentation polymorphisms associated with increased risk of melanoma in a case-control sample from southern Brazil." BMC cancer, 20(1)  [[2]](https://pubmed.ncbi.nlm.nih.gov/24244186/) Basu Mallick C et al. (2013). "The light skin allele of SLC24A5 in South Asians and Europeans shares identity by descent." PLoS genetics, 9(11)  [[3]](https://pubmed.ncbi.nlm.nih.gov/29897937/) Wang Y et al. (2018). "Four-dimensional, dynamic mosaicism is a hallmark of normal human skin that permits mapping of the organization and patterning of human epidermis during terminal differentiation." PloS one, 13(6)  [[4]](https://pubmed.ncbi.nlm.nih.gov/19440451/) Giardina E et al. (2008). "Haplotypes in SLC24A5 Gene as Ancestry Informative Markers in Different Populations." Current genomics, 9(2)  [[5]](https://pubmed.ncbi.nlm.nih.gov/33318654/) Jorgenson E et al. (2020). "Genetic ancestry, skin pigmentation, and the risk of cutaneous squamous cell carcinoma in Hispanic/Latino and non-Hispanic white populations." Communications biology, 3(1)  [[6]](https://pubmed.ncbi.nlm.nih.gov/39075179/) Rivera-Paredez B et al. (2024). "Skin pigmentation related variants in Mexican population and interaction effects on serum 25(OH)D concentration and vitamin D deficiency." Scientific reports, 14(1)  [[7]](https://pubmed.ncbi.nlm.nih.gov/23224873/) Wilson S et al. (2013). "NCKX5, a natural regulator of human skin colour variation, regulates the expression of key pigment genes MC1R and alpha-MSH and alters cholesterol homeostasis in normal human melanocytes." Advances in experimental medicine and biology, 961()  [[8]](https://pubmed.ncbi.nlm.nih.gov/23525585/) Wilson RT et al. (2011). "Genetic Ancestry, Skin Reflectance and Pigmentation Genotypes in Association with Serum Vitamin D Metabolite Balance." Hormone molecular biology and clinical investigation, 7(1)  [[9]](https://pubmed.ncbi.nlm.nih.gov/26395555/) Santos HC et al. (2016). "A minimum set of ancestry informative markers for determining admixture proportions in a mixed American population: the Brazilian set." European journal of human genetics : EJHG, 24(5)  [[10]](https://pubmed.ncbi.nlm.nih.gov/26918427/) Jonnalagadda M et al. (2016). "Association of genetic variants with skin pigmentation phenotype among populations of west Maharashtra, India." American journal of human biology : the official journal of the Human Biology Council, 28(5)  [[11]](https://pubmed.ncbi.nlm.nih.gov/27866970/) Mishra A et al. (2017). "Genotype-Phenotype Study of the Middle Gangetic Plain in India Shows Association of rs2470102 with Skin Pigmentation." The Journal of investigative dermatology, 137(3)  [[12]](https://pubmed.ncbi.nlm.nih.gov/39940926/) Mpofana N et al. (2025). "Association of Genetic Polymorphisms in SLC45A2, TYR, HERC2, and SLC24A in African Women with Melasma: A Pilot Study." International journal of molecular sciences, 26(3)  [[13]](https://pubmed.ncbi.nlm.nih.gov/22073278/) Leite TK et al. (2011). "Genomic ancestry, self-reported "color" and quantitative measures of skin pigmentation in Brazilian admixed siblings." PloS one, 6(11)  [[14]](https://pubmed.ncbi.nlm.nih.gov/34068188/) Shan MA et al. (2021). "Analysis of Skin Pigmentation and Genetic Ancestry in Three Subpopulations from Pakistan: Punjabi, Pashtun, and Baloch." Genes, 12(5)  [[15]](https://pubmed.ncbi.nlm.nih.gov/16847698/) Soejima M et al. (2007). "Population differences of two coding SNPs in pigmentation-related genes SLC24A5 and SLC45A2." International journal of legal medicine, 121(1) |
|  | Evaluation  **Summarization Accuracy & Completeness:**  **Hallucination Rate:**  **Readability & Usability:**  **Literature Coverage & Relevance:**  **Time Efficiency:** | |
