## Supplementary material for "Benchmarking generative AI tools for literature retrieval and summarization in genomic variant interpretation": TREX1 c.290G_A.docx

| **Tool** | **Summary** | **References** |
| --- | --- | --- |
| **1** | The TREX1 gene encodes the three prime repair exonuclease 1 enzyme, a DNA exonuclease responsible for trimming nucleotides from the 3' ends of DNA strands. This enzyme plays a crucial role in maintaining genomic stability by removing aberrant DNA fragments that arise during various cellular processes, such as DNA replication, repair, and apoptosis. By degrading these DNA fragments, TREX1 prevents inappropriate activation of the immune system, which could otherwise mistake self-derived DNA for pathogenic nucleic acids, leading to autoimmune responses. The c.290G>A (p.Arg97His) variant in the TREX1 gene results in a non-conservative amino acid substitution at position 97, replacing arginine with histidine. This variant has been identified in individuals with Aicardi-Goutières syndrome (AGS), a rare genetic disorder characterized by severe neurological dysfunction, chronic encephalopathy, and elevated levels of interferon-alpha in the cerebrospinal fluid. Functional studies suggest that such mutations impair the exonuclease activity of TREX1, leading to the accumulation of endogenous DNA fragments that trigger an aberrant immune response, culminating in the clinical manifestations observed in AGS. The c.290G>A variant has been detected at a low frequency in the general population, with an allele frequency of approximately 0.00006 in large-scale genomic databases, indicating its rarity. In silico predictive tools consistently suggest a deleterious effect on protein function. However, the clinical significance of this variant remains uncertain due to limited direct evidence linking it to disease phenotypes. While some studies have reported the presence of this variant in AGS patients, comprehensive genotype-phenotype correlations are lacking, and functional assays specifically assessing the impact of the p.Arg97His substitution are scarce. Given the current evidence, the c.290G>A (p.Arg97His) variant in the TREX1 gene is considered a variant of uncertain significance. Further research, including detailed functional analyses and expanded clinical studies, is essential to elucidate its pathogenicity and potential role in AGS or related autoimmune conditions. | [ncbi.nlm.nih.gov TREX1 three prime repair exonuclease 1 [ (human)] - NCBI](https://www.ncbi.nlm.nih.gov/gene/11277)  [4 marzo 2025 — This gene encodes a nuclear protein with 3' exonuclease activity. ...](https://www.ncbi.nlm.nih.gov/gene/11277)  [genecards.org TREX1 Gene - GeneCards \| TREX1 Protein \| TREX1 Antibody 25 dicembre 2024 — This gene encodes a nuclear protein with 3' exonuclease activity. ...](https://www.genecards.org/cgi-bin/carddisp.pl?gene=TREX1)  [medlineplus.gov TREX1 gene: MedlinePlus Genetics 26 settembre 2024 — The TREX1 gene provides instructions for making the three prime repair exonuclease 1 enzyme. This enzyme is a DNA exonuclease.](https://medlineplus.gov/genetics/gene/trex1/)  [medlineplus.gov [PDF] TREX1 gene - MedlinePlus 26 settembre 2024 — ...](https://medlineplus.gov/download/genetics/gene/trex1.pdf)  [medlineplus.gov [PDF] TREX1 gene - MedlinePlus 26 settembre 2024 — The TREX1 gene provides instructions for making the three prime repair exonuclease 1 enzyme., ...](https://medlineplus.gov/download/genetics/gene/trex1.pdf)  [nature.com Structural basis of human TREX1 DNA degradation and ... - Nature](https://www.nature.com/articles/s41467-022-32055-z)  [25 luglio 2022 — TREX1 is a cytosolic DNA nuclease essential for regulation ...](https://www.nature.com/articles/s41467-022-32055-z)  [frontiersin.org TREX1 as a Novel Immunotherapeutic Target - Frontiers](https://www.frontiersin.org/journals/immunology/articles/10.3389/fimmu.2021.660184/full)  [31 marzo 2021 — The 3' → 5' exonuclease TREX1 acts in vivo to degrade DNA...](https://www.frontiersin.org/journals/immunology/articles/10.3389/fimmu.2021.660184/full)  [uniprot.org TREX1 - Three-prime repair exonuclease 1 - Homo sapiens (Human)](https://www.uniprot.org/uniprotkb/Q9NSU2/entry)  [20 dicembre 2017 — Plays a key role in degradation of DNA fragments at cytosolic micronuclei arising from genome instability: its association with the endoplasmic ...](https://www.uniprot.org/uniprotkb/Q9NSU2/entry)  [wellcomeopenresearch.org](https://wellcomeopenresearch.org/articles/2-106/v1/pdf)  [[PDF] 5' DNA exonuclease TREX1 in early onset small vessel stroke](https://wellcomeopenresearch.org/articles/2-106/v1/pdf)  [2 novembre 2017 — Abstract. Background: Monoallelic and …](https://wellcomeopenresearch.org/articles/2-106/v1/pdf)  [ncbi.nlm.nih.gov NM_033629.6(TREX1):c.290G>A (p.Arg97His) AND Aicardi ... - NCBI](https://www.ncbi.nlm.nih.gov/clinvar/RCV000490435)  [Variant summary: TREX1 c.290G>A (p.Arg97His) results in a non-conservative...](https://www.ncbi.nlm.nih.gov/clinvar/RCV000490435)  [ncbi.nlm.nih.gov NM_033629.6(TREX1):c.290G>A (p.Arg97His) AND Aicardi ... - NCBI](https://www.ncbi.nlm.nih.gov/clinvar/RCV000490435.3/)  [Variant summary: TREX1 c.290G>A (p.Arg97His) results in a non-conservative...](https://www.ncbi.nlm.nih.gov/clinvar/RCV000490435.3/)  [databases.lovd.nl Unique variants in the TREX1 gene - Global Variome shared LOVD](https://databases.lovd.nl/shared/variants/TREX1/unique)  [(?), p.(Arg152His), -, likely pathogenic, pathogenic (recessive), g.48508344G>A, -, 290G>A (Arg97His), TREX1(NM_033629.6):c.290G>A (p.R97H) ... TREX1 c.539A>G, p.](https://databases.lovd.nl/shared/variants/TREX1/unique)  [databases.lovd.nl All variants in the TREX1 gene - Global Variome shared LOVD](https://databases.lovd.nl/shared/variants/TREX1)  [TREX1(NM_033629.6):c.290G>A (p.R97H), -, TREX1_000064, VKGL data sharing initiative Nederland, -, -, -, CLASSIFICATION record, -, -, -, -, -, VKGL-NL_Groningen.](https://databases.lovd.nl/shared/variants/TREX1)  [ClinVar Miner List of variants in gene combination ATRIP, ATRIP-TREX1, TREX1 ...](https://clinvarminer.genetics.utah.edu/variants-by-condition/Aicardi-Goutieres%20syndrome%201/gene/ATRIP%2C%20ATRIP-TREX1%2C%20TREX1/likely%20pathogenic)  [List of variants in gene combination ATRIP, ATRIP-TREX1, TREX1 reported as likely pathogenic for Aicardi-Goutieres syndrome 1 ; NM_033629.6(TREX1):c.290G>A (p.](https://clinvarminer.genetics.utah.edu/variants-by-condition/Aicardi-Goutieres%20syndrome%201/gene/ATRIP%2C%20ATRIP-TREX1%2C%20TREX1/likely%20pathogenic)  [ClinVar Miner List of variants in gene combination ATRIP, ATRIP-TREX1, TREX1 ...](https://clinvarminer.genetics.utah.edu/variants-by-submitter/505765/gene/ATRIP%2C%20ATRIP-TREX1%2C%20TREX1)  [Total variants: 2. Download table as spreadsheet. HGVS, dbSNP, gnomAD frequency. NM_033629.6(TREX1):c.290G>A (p.Arg97His), rs200773268, 0.00003. NM_033629.6( ...](https://clinvarminer.genetics.utah.edu/variants-by-submitter/505765/gene/ATRIP%2C%20ATRIP-TREX1%2C%20TREX1)  [en.wikipedia.org TREX1 - Wikipedia DNA recombination ...](https://en.wikipedia.org/wiki/TREX1)  [neurology.org Aicardi-Goutières Syndrome Presenting as Recurrent Ischemic Stroke](https://www.neurology.org/doi/10.1212/WNL.0000000000200952)  [This patient with TREX1-mutated AGS was a child, carrying the same TREX1 variant of c.290G>A as our patient. However, the stroke was revealed by MRI, and the ...](https://www.neurology.org/doi/10.1212/WNL.0000000000200952)  [v1.marrvel.org TREX1 - MARRVEL: Search Result MIM number: ...](https://v1.marrvel.org/search/gene/TREX1) |
|  | Evaluation  **Summarization Accuracy & Completeness:**  **Hallucination Rate:**  **Readability & Usability:**  **Literature Coverage & Relevance:**  **Time Efficiency:** | |
| **2** | The genomic variant c.290G>A, resulting in the amino acid substitution p.Arg97His (rs200773268), is located in the TREX1 gene, which encodes a 3'→5' exonuclease. TREX1 plays a critical role in degrading cytosolic DNA, thereby acting as a nucleolytic antagonist of the cGAS-STING pathway, which is involved in the innate immune response. The proper functioning of TREX1 is essential for preventing the accumulation of cytosolic DNA that can trigger autoinflammatory responses [2][3].  The p.Arg97His variant has been implicated in the pathogenesis of Aicardi-Goutieres syndrome (AGS), a severe autoinflammatory disorder characterized by chronic systemic and neurological inflammation, as well as elevated type I interferon activity. This variant is located within the dimerization interface of TREX1, which is critical for the protein's folding and enzymatic activity. The substitution of arginine to histidine at position 97 disrupts hydrogen-bond contacts that are essential for the stability of the dimer interface, thereby potentially impairing the protein's function [2].  Structural analyses have demonstrated that the p.Arg97His mutation, along with other mutations such as p.Arg114His, hinders the requisite homodimerization of TREX1, which is necessary for its exonuclease activity. This disruption can lead to a loss of function, contributing to the accumulation of cytosolic DNA and subsequent activation of the cGAS-STING pathway, resulting in the overproduction of type I interferons and the clinical manifestations of AGS [2][3][4].  In addition to its role in AGS, the p.Arg97His variant has been associated with other autoimmune diseases, including systemic lupus erythematosus and familial chilblain lupus. The variant's impact on TREX1's dimerization and function underscores its significance in the pathophysiology of these disorders [2].  ClinVar submissions have classified the c.290G>A p.Arg97His variant as pathogenic, with a total of three submissions supporting this classification. This further corroborates the variant's clinical relevance and its association with disease phenotypes.  In summary, the c.290G>A p.Arg97His variant in the TREX1 gene is a missense mutation that compromises the protein's dimerization and enzymatic activity, leading to the pathogenesis of AGS and other related autoimmune conditions. The variant's detrimental impact on TREX1 function highlights the importance of this exonuclease in maintaining immune homeostasis by regulating cytosolic DNA levels and preventing aberrant immune activation. | [[1]](https://pubmed.ncbi.nlm.nih.gov/31130681/) Garau J et al. (2019). "Molecular Genetics and Interferon Signature in the Italian Aicardi Goutieres Syndrome Cohort: Report of 12 New Cases and Literature Review." Journal of clinical medicine, 8(5)  [[2]](https://pubmed.ncbi.nlm.nih.gov/35879334/) Zhou W et al. (2022). "Structural basis of human TREX1 DNA degradation and autoimmune disease." Nature communications, 13(1)  [[3]](https://pubmed.ncbi.nlm.nih.gov/38260344/) Shim A et al. (2024). "Mutations in the non-catalytic polyproline motif destabilize TREX1 and amplify cGAS-STING signaling." bioRxiv : the preprint server for biology, ()  [[4]](https://pubmed.ncbi.nlm.nih.gov/38796715/) Shim A et al. (2024). "Mutations in the non-catalytic polyproline motif destabilize TREX1 and amplify cGAS-STING signaling." Human molecular genetics, 33(18)  [[5]](https://pubmed.ncbi.nlm.nih.gov/38003924/) Swierczynska M et al. (2023). "Aicardi-Goutieres Syndrome with Congenital Glaucoma Caused by Novel TREX1 Mutation." Journal of personalized medicine, 13(11)  [[6]](https://pubmed.ncbi.nlm.nih.gov/33868310/) Hemphill WO et al. (2021). "TREX1 as a Novel Immunotherapeutic Target." Frontiers in immunology, 12()  [[7]](https://pubmed.ncbi.nlm.nih.gov/25138095/) Ellyard JI et al. (2014). "Identification of a pathogenic variant in TREX1 in early-onset cerebral systemic lupus erythematosus by Whole-exome sequencing." Arthritis & rheumatology (Hoboken, N.J.), 66(12)  [[8]](https://pubmed.ncbi.nlm.nih.gov/34362117/) Wajda A et al. (2021). "Application of NGS Technology in Understanding the Pathology of Autoimmune Diseases." Journal of clinical medicine, 10(15)  [[9]](https://pubmed.ncbi.nlm.nih.gov/35262626/) Giordano AMS et al. (2022). "DNA damage contributes to neurotoxic inflammation in Aicardi-Goutieres syndrome astrocytes." The Journal of experimental medicine, 219(4)  [[10]](https://pubmed.ncbi.nlm.nih.gov/27224741/) Chung SA et al. (2016). "Rare variants, autoimmune disease, and arthritis." Current opinion in rheumatology, 28(4)  [[11]](https://pubmed.ncbi.nlm.nih.gov/35595529/) Qin Z et al. (2022). "Distinct landscapes of deleterious variants in DNA damage repair system in ethnic human populations." Life science alliance, 5(9) *Variant reported in Supplementary Information*  [[12]](https://pubmed.ncbi.nlm.nih.gov/36922933/) White JA et al. (2022). "Whole-exome Sequencing of Nigerian Prostate Tumors from the Prostate Cancer Transatlantic Consortium (CaPTC) Reveals DNA Repair Genes Associated with African Ancestry." Cancer research communications, 2(9) *Variant reported in Supplementary Information*  [[13]](https://pubmed.ncbi.nlm.nih.gov/31589614/) Capalbo A et al. (2019). "Optimizing clinical exome design and parallel gene-testing for recessive genetic conditions in preconception carrier screening: Translational research genomic data from 14,125 exomes." PLoS genetics, 15(10) *Variant reported in Supplementary Information* |
|  | Evaluation  **Summarization Accuracy & Completeness:**  **Hallucination Rate:**  **Readability & Usability:**  **Literature Coverage & Relevance:**  **Time Efficiency:** | |
| **3** | The TREX1 gene encodes the three prime repair exonuclease 1, a DNA exonuclease crucial for the degradation of cytosolic single-stranded DNA, thereby preventing aberrant activation of the innate immune response. Pathogenic variants in TREX1 are associated with a spectrum of autoimmune and neuroinflammatory disorders, including Aicardi-Goutières syndrome (AGS), familial chilblain lupus, and retinal vasculopathy with cerebral leukodystrophy. The c.290G>A variant in TREX1 has been previously reported in at least one case of AGS, suggesting a pathogenic role in the disorder, although broader epidemiological data on its frequency remain limited [1]. Mutations in TREX1 disrupt its exonuclease activity, leading to accumulation of cytosolic DNA and subsequent activation of the interferon response, a mechanism strongly implicated in the pathophysiology of AGS and other interferonopathies [2]. The clinical phenotype associated with TREX1 variants is highly variable, ranging from early-onset severe neurodevelopmental impairment with cerebral calcifications to milder cutaneous or vascular manifestations, depending on the specific genetic alteration and its functional consequences. Computational pathogenicity predictions and functional studies indicate that TREX1 variants affecting conserved exonuclease domains are likely to result in loss-of-function effects, reinforcing their disease-causing potential [3]. However, given the rarity of the c.290G>A variant, further research, including segregation analysis and functional assays, is necessary to confirm its pathogenicity and clinical significance. The classification of this variant according to the American College of Medical Genetics and Genomics (ACMG) guidelines remains uncertain, requiring additional genetic and phenotypic correlation studies. As with other rare TREX1 mutations, clinicians should consider genetic testing in individuals presenting with unexplained neuroinflammatory or autoimmune features and assess variant impact within the broader context of interferonopathy-related conditions [4]. Given the current limitations in published evidence, caution is warranted in interpreting its significance, and functional validation studies are recommended. | [1] Kuang S, et al. (2022) *Child neurology: Aicardi-Goutières syndrome presenting as recurrent ischemic stroke*. Neurology. DOI:10.1212/wnl.0000000000200952  [2] Świerczyńska M, et al. (2023) *Aicardi–Goutières Syndrome with Congenital Glaucoma Caused by Novel TREX1 Mutation*. J Pers Med. DOI:10.3390/jpm13111609  [3] Hofer MJ, et al. (2024) *The prototypical interferonopathy: Aicardi-Goutières syndrome from bedside to bench*. Immunol Rev. [DOI:10.1111/imr.13413](https://onlinelibrary.wiley.com/doi/abs/10.1111/imr.13413)  [4] Garau J, et al. (2019) *Molecular genetics and interferon signature in the Italian Aicardi Goutières syndrome cohort: report of 12 new cases and literature review*. J Clin Med. DOI:10.3390/jcm8050750 |
|  | Evaluation  **Summarization Accuracy & Completeness:**  **Hallucination Rate:**  **Readability & Usability:**  **Literature Coverage & Relevance:**  **Time Efficiency:** | |
| **4** | The TREX1 gene, also known as Three Prime Repair Exonuclease 1, encodes a nuclear protein with 3' exonuclease activity. This protein plays a crucial role in DNA repair and serves as a proofreading function for DNA polymerase. Mutations in TREX1 are associated with several immune-related disorders, including Aicardi-Goutières syndrome, chilblain lupus, and Cree encephalitis. These conditions highlight the importance of TREX1 in maintaining genomic stability and preventing autoimmune responses.  The TREX1 c.290G>A variant is a missense mutation that results in the substitution of glycine with aspartic acid at codon 97 (p.Gly97Asp) in the TREX1 protein. This variant has been identified in individuals with systemic lupus erythematosus (SLE) and other autoimmune disorders. The functional impact of this variant is not fully understood, but it is believed to affect the exonuclease activity of TREX1, leading to the accumulation of DNA fragments in the cytosol and subsequent activation of the innate immune response.  The frequency of the TREX1 c.290G>A variant in the general population is not well documented, but it has been observed in multiple individuals with SLE. The classification of this variant is not definitive, as the available evidence is currently insufficient to determine its role in disease. Algorithms developed to predict the effect of missense changes on protein structure and function are either unavailable or do not agree on the potential impact of this variant.  In vitro studies have shown that the TREX1 c.290G>A variant affects the protein's ability to degrade DNA fragments, which is essential for preventing the activation of the innate immune response. The accumulation of DNA fragments in the cytosol can lead to the activation of the cGAS-STING pathway, resulting in the production of type I interferon and inflammation. This mechanism is believed to contribute to the development of autoimmune disorders in individuals with this variant.  In conclusion, the TREX1 c.290G>A variant is a missense mutation that affects the exonuclease activity of TREX1, leading to the accumulation of DNA fragments in the cytosol and subsequent activation of the innate immune response. The functional impact of this variant is not fully understood, but it has been observed in individuals with SLE and other autoimmune disorders. Further research is needed to determine the role of this variant in disease and to develop targeted therapies for individuals with this mutation. | 1. Lee-Kirsch MA, Gong M, Chowdhury D, et al. Nat Genet. 2007 Sep;39(9):1065-7. Epub 2007 Jul 29.  2. Ellyard JI, Jerjen R, Martin JL, et al. Arthritis Rheumatol. 2014 Dec;66(12):3382-6. doi: 10.1002/art.38824.  3. Kisla Ekinci RM, Balci S, Bisgin A, et al. Eur J Med Genet. 2017 Dec;60(12):690-694. doi: 10.1016/j.ejmg.2017.09.004. Epub 2017 Sep 13.  4. Crow YJ, Rehwinkel H, Odom VT, et al. Cell. 2006 May 19;125(4):661-73. doi: 10.1016/j.cell.2006.03.037.  5. Rice GI, Newman WG, Dean JL, et al. Nat Genet. 2007 Jan;39(1):102-4. Epub 2006 Nov 26. |
|  | Evaluation  **Summarization Accuracy & Completeness:**  **Hallucination Rate:**  **Readability & Usability:**  **Literature Coverage & Relevance:**  **Time Efficiency:** | |
| **5** | The TREX1 gene encodes the major 3'-5' DNA exonuclease, which is crucial for degrading cytosolic DNA to prevent aberrant immune activation[2](https://www.semanticscholar.org/paper/8450025ceca8b0dd82a59c2cf9a1313a2b66eddb)[5](https://pubmed.ncbi.nlm.nih.gov/31475890/). This exonuclease reduces both single- and double-stranded DNA, thereby suppressing self-recognition and protecting the host from inappropriate autoimmune responses[5](https://pubmed.ncbi.nlm.nih.gov/31475890/). When TREX1 is inactivated, DNA accumulates in the cytosol, leading to the induction of the cGAS-STING DNA sensing pathway and subsequent interferon-mediated innate immune signaling and inflammation[2](https://www.semanticscholar.org/paper/8450025ceca8b0dd82a59c2cf9a1313a2b66eddb). Pathogenic mutations in TREX1 can lead to hereditary autoimmune and autoinflammatory disorders[2](https://www.semanticscholar.org/paper/8450025ceca8b0dd82a59c2cf9a1313a2b66eddb). Specifically, mutations in the TREX1 gene have been associated with Aicardi-Goutières syndrome (AGS), familial chilblain lupus (FCL), systemic lupus erythematosus (SLE), and retinal vasculopathy with cerebral leukodystrophy (RVCL)[5](https://pubmed.ncbi.nlm.nih.gov/31475890/). A study involving patients with autoimmune diseases identified several TREX1 variations, reinforcing the gene's involvement in such conditions, and suggesting that the spectrum of autoimmune diseases linked to TREX1 may be broader than currently recognized, though the role of TREX1 variants in SLE was not confirmed in the study[1](https://www.ncbi.nlm.nih.gov/pmc/articles/PMC3810194/). The TREX1 c.290G>A variant is not specifically discussed in the available literature. Given the general role of TREX1 in preventing undue immune activation and its association with several autoimmune disorders, further investigation may be warranted to determine the specific impact and clinical significance of the c.290G>A variant. | **References**  [1](https://www.ncbi.nlm.nih.gov/pmc/articles/PMC3810194/) (PMCID: PMC3810194)  [2](https://www.semanticscholar.org/paper/8450025ceca8b0dd82a59c2cf9a1313a2b66eddb) (Abstract 4345)  [5](https://pubmed.ncbi.nlm.nih.gov/31475890/) (PMID: 31475890)  **Citations:**   1. <https://www.ncbi.nlm.nih.gov/pmc/articles/PMC3810194/> 2. <https://www.semanticscholar.org/paper/8450025ceca8b0dd82a59c2cf9a1313a2b66eddb> 3. <https://pubmed.ncbi.nlm.nih.gov/25604658/> 4. <https://pubmed.ncbi.nlm.nih.gov/23732569/> 5. <https://pubmed.ncbi.nlm.nih.gov/31475890/> 6. <https://www.ncbi.nlm.nih.gov/pmc/articles/PMC5133658/> 7. <https://pubmed.ncbi.nlm.nih.gov/16845398/> 8. <https://pubmed.ncbi.nlm.nih.gov/27566796/> |
|  | Evaluation  **Summarization Accuracy & Completeness:**  **Hallucination Rate:**  **Readability & Usability:**  **Literature Coverage & Relevance:**  **Time Efficiency:** | |
