## Supplementary material for "Benchmarking generative AI tools for literature retrieval and summarization in genomic variant interpretation": UBE3A c.2616__6del.docx

| **Tool** | **Summary** | **References** |
| --- | --- | --- |
| **1** | The UBE3A gene encodes ubiquitin protein ligase E3A, an enzyme integral to the ubiquitin-proteasome system, which tags proteins for degradation, thereby maintaining cellular protein homeostasis. This function is particularly vital in neuronal cells, where UBE3A regulates synaptic plasticity, essential for learning and memory. Notably, UBE3A exhibits genomic imprinting in neurons; only the maternally inherited allele is active, while the paternal allele remains silenced. Disruptions in UBE3A are implicated in Angelman syndrome, a neurodevelopmental disorder characterized by intellectual disability, motor dysfunction, and seizures. The specific variant c.2616_*6del results in the deletion of nucleotides spanning from position 2616 in the coding sequence to 6 nucleotides past the stop codon, leading to the elongation of the protein beyond its normal termination at codon 873. This alteration changes the stop codon to a glutamine residue and extends the protein until a new stop codon is encountered, designated as p.Ter873GlnextTer?. The precise impact of this variant on protein function is currently undetermined due to limited data. Given UBE3A's role in neural development and function, such an extension could potentially disrupt its enzymatic activity or interaction with substrates, possibly contributing to neurological phenotypes. However, without specific functional studies on this variant, its pathogenicity remains uncertain. To date, no peer-reviewed publications have directly examined the c.2616_*6del p.Ter873GlnextTer? variant. Consequently, further research, including functional assays and clinical correlation studies, is essential to elucidate the clinical significance of this genetic alteration. | <https://en.wikipedia.org/wiki/UBE3A>  <https://www.wired.com/2011/12/paternal-gene-targeting/>  <https://medlineplus.gov/genetics/gene/ube3a/>  <https://medlineplus.gov/download/genetics/gene/ube3a.pdf>  <https://neuroscience.wustl.edu/yi-lab-assay-genetic-variants/>  <https://aesnet.org/abstractslisting/seizure-susceptibility-and-neuroinflammation-in-the-ube3a-maternal-deletion-rat-model-of-angelman-syndrome>  <https://atlasgeneticsoncology.org/gene/42756/ube3a-%28ubiquitin-protein-ligase-e3a%29>  <https://www.ncbi.nlm.nih.gov/clinvar/RCV000144308/>  <https://www.ncbi.nlm.nih.gov/clinvar/variation/155985/>  <https://www.ncbi.nlm.nih.gov/gene/7337>  <https://pmc.ncbi.nlm.nih.gov/articles/PMC2843143/>  <https://clinvarminer.genetics.utah.edu/variants-by-gene/SNHG14%2C%20UBE3A/significance/pathogenic>  <https://rgd.mcw.edu/rgdweb/report/gene/main.html?id=1314934>  [https://www.malacards.org/card/angelman_syndrome](https://www.malacards.org/card/angelman_syndrome?search=MTHFR)  <https://www.sciencedirect.com/topics/biochemistry-genetics-and-molecular-biology/ube3a> |
|  | Evaluation  **Summarization Accuracy & Completeness:**  **Hallucination Rate:**  **Readability & Usability:**  **Literature Coverage & Relevance:**  **Time Efficiency:** | |
| **2** | The UBE3A gene encodes ubiquitin protein ligase E3A, an enzyme crucial for targeting proteins for degradation within cells. This process is essential for maintaining normal cellular functions by removing damaged or unnecessary proteins. The UBE3A gene is subject to genomic imprinting, where only the maternal copy is active in certain brain regions, particularly in neurons. This imprinting mechanism is critical because the paternal copy is normally inactive in these areas. The primary function of UBE3A is to attach ubiquitin molecules to proteins, marking them for degradation by the proteasome. This activity is part of the ubiquitin-proteasome pathway, which regulates various cellular processes, including protein quality control and signal transduction. The UBE3A gene is located on chromosome 15q11-13, a region implicated in several neurodevelopmental disorders. Mutations or deletions in the UBE3A gene are associated with Angelman syndrome, a severe neurological disorder characterized by developmental delays, intellectual disability, seizures, and movement disorders. These mutations often result in a nonfunctional or truncated version of the protein, leading to a loss of its enzymatic activity. The variant c.2616_*6del p.Ter873GlnextTer? in the UBE3A gene is a deletion that affects the stop codon and extends the reading frame, likely resulting in a longer, nonfunctional protein. This variant is classified as pathogenic and is associated with Angelman syndrome. The frequency of this specific variant is not well-documented in the general population, but it is known to be rare. The impact of this variant is severe, as it disrupts the normal function of UBE3A, leading to the characteristic features of Angelman syndrome. The classification of this variant as pathogenic is based on its predicted effect on protein function and its association with a well-defined clinical phenotype. However, the exact mechanisms by which this variant contributes to the pathogenesis of Angelman syndrome are not fully understood. Further research is needed to elucidate the molecular consequences of this variant and to develop targeted therapeutic strategies. The current understanding of this variant is limited to its genetic classification and clinical association, highlighting the need for more detailed functional studies. | [1] MedlinePlus Genetics. UBE3A gene. Available at: <https://medlineplus.gov/genetics/gene/ube3a/>  [2] Wikipedia. UBE3A. Available at: <https://en.wikipedia.org/wiki/UBE3A>  [3] OMIM. UBIQUITIN-PROTEIN LIGASE E3A; UBE3A. Available at: <https://omim.org/entry/601623>  [4] ClinVar. NM_130839.5(UBE3A):c.2616_*6del (p.Ter873GlnextTer?) AND Angelman syndrome. Available at: <https://www.ncbi.nlm.nih.gov/clinvar/RCV000144308/> |
|  | Evaluation  **Summarization Accuracy & Completeness:**  **Hallucination Rate:**  **Readability & Usability:**  **Literature Coverage & Relevance:**  **Time Efficiency:** | |
| **3** | There was an issue retrieving peer-reviewed articles directly referencing the UBE3A variant c.2616_*6del (p.Ter873GlnextTer?), as no such studies were identified in academic databases. Given the absence of variant-specific literature, available information about the gene's function and known pathogenic mechanisms must be considered to assess potential clinical implications. The UBE3A gene encodes an E3 ubiquitin-protein ligase involved in targeting proteins for degradation via the ubiquitin-proteasome pathway. It plays a crucial role in synaptic function and neurodevelopment and is particularly notable for its brain-specific imprinting, where the maternal allele is primarily expressed in neurons. Disruptions in the maternal copy of UBE3A are causative for Angelman syndrome, a severe neurodevelopmental disorder characterized by intellectual disability, impaired speech, ataxia, and a unique behavioral phenotype. The variant p.Ter873GlnextTer? is a deletion that removes the canonical stop codon, potentially resulting in translation beyond the normal termination site into the 3′ untranslated region. This class of variant, known as stop-loss, can lead to protein elongation, which may impair normal protein folding, stability, or function, although the precise functional consequences of such elongation are context-dependent and often uncertain. No clinical reports or functional assays assessing the pathogenicity of this specific variant were identified, highlighting the need for further investigation. Given the gene’s established dosage sensitivity and the potential for disruption of tightly regulated protein interactions, variants affecting the C-terminal end of UBE3A warrant caution. However, without functional validation or clinical case correlations, the variant should be classified as a Variant of Uncertain Significance (VUS) per ACMG/AMP guidelines. Additional studies, including segregation analysis, functional assays, or identification in affected individuals with consistent phenotypes, are recommended to clarify pathogenic potential. In the absence of published data directly referencing this variant, no peer-reviewed citations can be provided at this time. | No references available |
|  | Evaluation  **Summarization Accuracy & Completeness:**  **Hallucination Rate:**  **Readability & Usability:**  **Literature Coverage & Relevance:**  **Time Efficiency:** | |
| **4** | The UBE3A gene encodes the E3 ubiquitin ligase E6-associated protein (E6AP), which plays a crucial role in neurodevelopment by regulating protein degradation and synaptic plasticity[4]. The gene is imprinted, with the maternal allele predominantly expressed in neurons, while the paternal allele is silenced[4]. Mutations or deletions in the maternal UBE3A allele lead to Angelman syndrome, characterized by severe neurological symptoms including speech and language impairments, intellectual disability, and seizures[5][6]. Conversely, overexpression of UBE3A due to copy number variations is associated with autism spectrum disorders[4][8].  The variant c.2616_*6del p.Ter873GlnextTer in the UBE3A gene introduces a frameshift mutation that likely disrupts the normal function of the E6AP protein. However, specific research findings directly related to this variant are currently unavailable. Given the critical role of UBE3A in neurodevelopmental disorders, any mutation affecting its function could potentially have significant clinical implications. The classification and frequency of this variant are not well-documented in the literature, and its impact on protein function and disease risk remains uncertain.  Further investigation is recommended to determine the clinical significance of this variant, including its potential to cause or contribute to neurodevelopmental disorders like Angelman syndrome or autism spectrum disorders. This would involve assessing the variant's effect on E6AP expression and function, as well as evaluating its presence in affected individuals and their families. | [1] <https://pubmed.ncbi.nlm.nih.gov/35534672/>  [2] <https://pubmed.ncbi.nlm.nih.gov/36947593/>  [3] <https://pubmed.ncbi.nlm.nih.gov/12210318/>  [4] <https://www.ncbi.nlm.nih.gov/pmc/articles/PMC6502993/>  [5] <https://pubmed.ncbi.nlm.nih.gov/18487518/>  [6] <https://www.ncbi.nlm.nih.gov/pmc/articles/PMC11169893/>  [7] <https://pubmed.ncbi.nlm.nih.gov/38922486/>  [8] <https://www.ncbi.nlm.nih.gov/pmc/articles/PMC10814747/> |
|  | Evaluation  **Summarization Accuracy & Completeness:**  **Hallucination Rate:**  **Readability & Usability:**  **Literature Coverage & Relevance:**  **Time Efficiency:** | |
| **5** | Variant c.2616_*6del p.Ter873GlnextTer? rs587781232 is a deletion that affects the canonical stop codon of the UBE3A gene, extending the open reading frame by an additional six amino acids before the next in-frame stop codon is encountered. This variant results in the loss of the normal termination signal at codon 873, and the introduction of a glutamine residue in place of the stop codon, followed by a short extension of the protein product.  UBE3A encodes an E3 ubiquitin-protein ligase that plays a critical role in targeting proteins for proteasomal degradation. It is involved in the ubiquitin-proteasome pathway, which is essential for the regulated turnover of proteins and for the elimination of misfolded proteins. The gene is known to be imprinted in certain tissues, including the brain, where only the maternal allele is typically expressed.  Given the nature of the variant, which affects the stop codon and results in an elongated protein product, it may impact the normal function of the UBE3A protein. However, without functional studies, the exact consequence of this elongation on the protein's function cannot be explicitly determined from the genomic information alone.  The variant rs587781232 has been submitted to ClinVar with a classification of pathogenic by a single submitter. This classification should be considered in the context of additional evidence, including functional studies, segregation analysis, and the clinical presentation of individuals with the variant, to determine its relevance to disease phenotypes. | [[1]](https://pubmed.ncbi.nlm.nih.gov/28569743/) Yang S et al. (2017). "Sources of discordance among germ-line variant classifications in ClinVar." Genetics in medicine : official journal of the American College of Medical Genetics, 19(10) [SUPP] |
|  | Evaluation  **Summarization Accuracy & Completeness:**  **Hallucination Rate:**  **Readability & Usability:**  **Literature Coverage & Relevance:**  **Time Efficiency:** | |
