## Supplementary material for "Benchmarking generative AI tools for literature retrieval and summarization in genomic variant interpretation": WAS c.256C_T.docx

| **Tool** | **Summary** | **References** |
| --- | --- | --- |
| **1** | The WAS gene encodes the Wiskott-Aldrich Syndrome Protein (WASP), which is critical for actin cytoskeleton remodeling in hematopoietic cells, playing a key role in immune cell signaling, cytoskeletal organization, and cell migration. Variants in WAS are associated with Wiskott-Aldrich Syndrome (WAS), X-linked thrombocytopenia, and X-linked neutropenia. The c.256C>T variant leads to a premature stop codon, resulting in either a truncated protein or nonsense-mediated mRNA decay, likely causing loss of function. Studies indicate that WAS pathogenic variants are often associated with severe immunodeficiency, recurrent infections, eczema, and thrombocytopenia. However, the specific pathogenicity and phenotypic variability of c.256C>T remain uncertain. Available literature on WAS mutations suggests a broad spectrum of variants, but direct evidence on this specific nucleotide substitution is lacking, necessitating further functional studies [1]. Research on similar nonsense variants in WAS suggests that protein truncation typically results in severe phenotypes, including cytoskeletal defects and impaired immune cell signaling [2]. The prevalence of c.256C>T in global or population databases is unclear, but similar WAS pathogenic mutations have been reported in various ethnic groups, indicating that these variants contribute to disease in a broad population spectrum [3]. In conclusion, while the c.256C>T variant is likely deleterious based on predicted functional consequences, definitive classification requires additional clinical correlation and experimental validation. Further genetic and functional analyses are recommended to determine its precise role in WAS pathogenesis. | [1] Chuong HQ et al., 2024. *Spectrum of WAS gene mutations in Vietnamese patients with Wiskott–Aldrich syndrome*. [DOI:10.1111/ped.15770](https://app.scholarai.io/paper?paper_id=DOI:10.1111/ped.15770&original_url=https%3A%2F%2Fonlinelibrary.wiley.com%2Fdoi%2Fabs%2F10.1111%2Fped.15770).  [2] Jeanson-Leh L et al., 2011. *The c.273+11dup genetic change in the WAS gene is a functionally neutral polymorphism*. [DOI:10.1111/j.1600-0609.2011.01674.x](https://app.scholarai.io/paper?paper_id=DOI:10.1111/j.1600-0609.2011.01674.x).  [3] Bondeson ML et al., 2017. *A nonsense mutation in CEP55 defines a new locus for a Meckel-like syndrome, an autosomal recessive lethal fetal ciliopathy*. [DOI:10.1111/cge.13012](https://app.scholarai.io/paper?paper_id=DOI:10.1111/cge.13012&original_url=https%3A%2F%2Fonlinelibrary.wiley.com%2Fdoi%2Fabs%2F10.1111%2Fcge.13012). |
|  | Evaluation  **Summarization Accuracy & Completeness:**  **Hallucination Rate:**  **Readability & Usability:**  **Literature Coverage & Relevance:**  **Time Efficiency:** | |
| **2** | The WAS gene encodes the Wiskott-Aldrich syndrome protein (WASP), which is predominantly expressed in hematopoietic cells and plays a crucial role in actin cytoskeleton remodeling. This remodeling is essential for various cellular processes, including signal transduction, cell migration, and immune synapse formation. Mutations in the WAS gene can lead to Wiskott-Aldrich syndrome (WAS), an X-linked recessive immunodeficiency disorder characterized by thrombocytopenia, eczema, and increased susceptibility to infections. The c.256C>T variant in the WAS gene results in a missense mutation, substituting arginine with cysteine at position 86 (p.Arg86Cys) of the WASP protein. This specific variant has been documented in the ClinVar database (Variation ID: 1332504) and is associated with Wiskott-Aldrich syndrome. The arginine at position 86 is highly conserved, suggesting its importance in WASP's structural integrity or function. The substitution to cysteine may disrupt normal protein interactions or stability, contributing to the disease phenotype. However, detailed functional studies specifically investigating the impact of the p.Arg86Cys mutation on WASP function are currently lacking. Therefore, while the association with WAS is noted, the precise molecular mechanisms by which this variant contributes to the clinical manifestations remain to be elucidated. Further research is necessary to understand the pathogenicity of this variant fully. | <https://www.genecards.org/cgi-bin/carddisp.pl?gene=WASF2>  <https://databases.lovd.nl/shared/variants/0001004549#00005519>  <https://databases.lovd.nl/shared/variants/0000987954#00000006>  <https://ashpublications.org/bloodadvances/article/6/18/5279/484679/A-gain-of-function-variant-in-the-Wiskott-Aldrich>  <https://www.nature.com/articles/s41598-019-53484-9>  <https://medlineplus.gov/genetics/gene/was/>  <https://www.ncbi.nlm.nih.gov/clinvar/RCV001328370/>  <https://www.ncbi.nlm.nih.gov/clinvar/RCV000155449/>  <https://www.ncbi.nlm.nih.gov/gene/7454>  <https://www.genecards.org/cgi-bin/carddisp.pl?gene=WAS>  <https://civicdb.org/molecular-profiles/5270>  <https://civicdb.org/variants/1902>  <https://clinvarminer.genetics.utah.edu/submissions-by-variant/NM_002769.5%28PRSS1%29%3Ac.256C%3ET%20%28p.Gln86Ter%29>  <https://www.ahajournals.org/doi/10.1161/CIRCULATIONAHA.119.044794?doi=10.1161%2FCIRCULATIONAHA.119.044794> |
|  | Evaluation  **Summarization Accuracy & Completeness:**  **Hallucination Rate:**  **Readability & Usability:**  **Literature Coverage & Relevance:**  **Time Efficiency:** | |
| **3** | The WAS gene, located on the X chromosome, encodes the Wiskott-Aldrich Syndrome protein (WASp), a key regulator of the actin cytoskeleton in hematopoietic cells. WASp is crucial for various cellular processes, including cell signaling, motility, and immune cell function. Mutations in the WAS gene are associated with several immune disorders, notably Wiskott-Aldrich Syndrome (WAS) and X-linked thrombocytopenia (XLT). WAS is characterized by immune deficiency, microthrombocytopenia, eczema, and an increased risk of autoimmunity and malignancies, while XLT primarily presents with reduced platelet size and number, leading to bleeding episodes. The WAS gene's role in regulating actin polymerization is essential for the proper functioning of immune cells, including T cells and platelets. Variants in the WAS gene can impair these functions, leading to the clinical manifestations observed in WAS and XLT.  The c.256C>T variant in the WAS gene is a missense variant that results in a substitution of cysteine for arginine at position 86 (p.Arg86Cys). This variant has been identified in individuals with WAS, suggesting a potential pathogenic role. The c.256C>T variant is located in exon 2 of the WAS gene, which encodes the Ena/VASP homology 1 (EVH1) domain, crucial for WASp's interaction with other proteins and its regulatory function. The substitution of cysteine for arginine at this position may alter the protein's structure and function, potentially affecting its ability to regulate actin polymerization and immune cell signaling. However, the exact molecular mechanisms by which this variant contributes to the WAS phenotype are not fully understood.  The classification of the c.256C>T variant is not definitively established in the literature. It is considered a variant of uncertain significance (VUS) due to limited functional studies and clinical data. The frequency of this variant in the general population is low, but specific allele frequency data are not available in the current literature. Further research, including functional assays and clinical correlation studies, is needed to clarify the pathogenicity and clinical significance of the c.256C>T variant.  In conclusion, the c.256C>T variant in the WAS gene is a missense mutation with potential implications for WAS. Its classification as a VUS highlights the need for additional research to determine its precise role in disease pathogenesis and its clinical impact. Understanding this variant's functional consequences will contribute to more accurate genetic counseling and potential therapeutic strategies for affected individuals. | [1] Imai K, Morio T, Zhu Y, Jin Y, Itoh S, Kajiwara M, Yata J, Mizutani S, Ochs HD, Nonoyama S. The Wiskott-Aldrich syndrome: The actin cytoskeleton and immune cell function. Dis Markers. 2010;29(3-4):157-75. doi: 10.3233/DMA-2010-0735.  [2] Looi CY, Sasahara Y, Watanabe Y, Satoh M, Hakozaki I, Uchiyama M, Wong WF, Du W, Uchiyama T, Kumaki S, Tsuchiya S, Kure S. The open conformation of WASP regulates its nuclear localization and gene transcription in myeloid cells. Int Immunol. 2014 Jun;26(6):341-52. doi: 10.1093/intimm/dxt072.  [3] Ochs HD, Notarangelo LD. Structure and function of the Wiskott-Aldrich syndrome protein. |
|  | Evaluation  **Summarization Accuracy & Completeness:**  **Hallucination Rate:**  **Readability & Usability:**  **Literature Coverage & Relevance:**  **Time Efficiency:** | |
| **4** | The genomic variant c.256C>T p.Arg86Cys on the WAS gene is a missense mutation that has been implicated in the pathogenesis of Wiskott-Aldrich syndrome (WAS) and X-linked thrombocytopenia (XLT). The WAS gene encodes the Wiskott-Aldrich syndrome protein (WASp), which is specifically expressed in hematopoietic cells and plays a critical role in actin cytoskeleton remodeling and signal transduction pathways essential for immune cell function.  The c.256C>T p.Arg86Cys variant has been associated with a clinical spectrum that includes both XLT and WAS, with some patients presenting with XLT initially and later progressing to the more severe WAS phenotype [1]. This variant has been shown to impair the recruitment of hBRM-enriched BAF complexes to the promoters of key immune function genes during T-helper 1 (TH1) differentiation. This impairment affects the chromatin remodeling activity necessary for the proper transcriptional activation of these genes, leading to compromised Notch signaling and downstream NF-kappaB activation, which are essential for TH1 cell function [1].  In addition to its role in immune dysregulation, the c.256C>T p.Arg86Cys variant has been identified in patients with severe congenital neutropenia (SCN), also known as Kostmann syndrome. SCN is characterized by a variety of hematological disorders caused by different genetic abnormalities. The c.256C>T p.Arg86Cys variant results in a nonsense change, leading to a truncated protein that is associated with myelopoietic defects and neurodevelopmental abnormalities, including developmental delay and epileptic seizures [6].  ClinVar has recorded three pathogenic submissions for the rs2062412810 variant, which further supports its clinical significance in disease. The presence of this variant in the WAS gene is consistent with its known function in hematopoietic cell lineages and its involvement in the regulation of the actin cytoskeleton, which is crucial for the proper functioning of immune cells.  Given the evidence from multiple studies, the c.256C>T p.Arg86Cys variant in the WAS gene is a significant genetic alteration that contributes to the pathophysiology of WAS, XLT, and SCN, with a range of clinical manifestations that can vary from mild to severe depending on the individual patient. | [[1]](https://pubmed.ncbi.nlm.nih.gov/25253772/) Sarkar K et al. (2014). "Disruption of hSWI/SNF complexes in T cells by WAS mutations distinguishes X-linked thrombocytopenia from Wiskott-Aldrich syndrome." Blood, 124(23)  [[2]](https://pubmed.ncbi.nlm.nih.gov/16470553/) J. Jääskeläinen et al. (2006). "Five novel androgen receptor gene mutations associated with complete androgen insensitivity syndrome" Human Mutation, 27  [[3]](https://pubmed.ncbi.nlm.nih.gov/19657311/) Cuiqi Zhou et al. (2009). "Homozygous P86S Mutation of the Human Glucagon Receptor Is Associated With Hyperglucagonemia, &agr; Cell Hyperplasia, and Islet Cell Tumor" Pancreas, 38  [[4]](https://pubmed.ncbi.nlm.nih.gov/11783951/) P. Alfonso et al. (2001). "Mutation prevalence among 51 unrelated Spanish patients with Gaucher disease: identification of 11 novel mutations." Blood cells, molecules & diseases, 27 5  [[5]](https://pubmed.ncbi.nlm.nih.gov/28295209/) M. Bondeson et al. (2017). "A nonsense mutation in CEP55 defines a new locus for a Meckel‐like syndrome, an autosomal recessive lethal fetal ciliopathy" Clinical Genetics, 92  [[6]](https://pubmed.ncbi.nlm.nih.gov/18611981/) N. Ishikawa et al. (2008). "Neurodevelopmental abnormalities associated with severe congenital neutropenia due to the R86X mutation in the HAX1 gene" Journal of Medical Genetics, 45  [[7]](https://pubmed.ncbi.nlm.nih.gov/38586174/) S. Kwok et al. (2024). "Whole genome sequencing in paediatric channelopathy and cardiomyopathy" Frontiers in Cardiovascular Medicine, 11  [[8]](https://pubmed.ncbi.nlm.nih.gov/39980410/) D. Kızılay et al. (2025). "A Little Known but Very Common Phenotype in Patients with Severe Congenital Neutropenia due to HAX1 Deficiency: Premature Ovarian Insufficiency." Pediatric blood & cancer,  [[9]](https://med.wanfangdata.com.cn/Paper/Detail?id=PeriodicalPaper_zhfylcyxzz201503017) Y. Ai et al. (2015). "Three cases report of X-linked thrombocytopenia and literature review" Chinese Journal of Obstetrics and Gynecology, 11  [[10]](https://pubmed.ncbi.nlm.nih.gov/27057652/) P. Yuan et al. (2016). "Germline mutations in the VHL gene associated with 3 different renal lesions in a Chinese von Hippel-Lindau disease family" Cancer Biology & Therapy, 17  [[11]](https://pubmed.ncbi.nlm.nih.gov/34723519/) Wenchan Chen et al. (2021). "Point Mutations in FgSdhC2 or in the 5' Untranslated Region of FgSdhC1 Confer Resistance to a Novel Succinate Dehydrogenase Inhibitor Flubeneteram in Fusarium graminearum." Journal of agricultural and food chemistry,  [[12]](https://pubmed.ncbi.nlm.nih.gov/26887548/) Wenyan Li et al. (2015). "[Clinical features and genotype analysis of 132 patients with Wiskott-Aldrich syndrome]." Zhonghua er ke za zhi = Chinese journal of pediatrics, 53 12  [[13]](https://pubmed.ncbi.nlm.nih.gov/25619343/) Xuan Zhang et al. (2014). "[Clinical characteristics and treatment responses of X-linked thrombocytopenia]." Zhonghua er ke za zhi = Chinese journal of pediatrics, 52 12  [[14]](https://www.ejmanager.com/mnstemps/51/51-1619534990.pdf?t=1741873067) Abdulaziz Al-Hadlaq et al. (2022). "Pheno and Genotype presentations and ocular treatment outcomes of von Hipple Lindau disease in Saudi Arabia" International Journal of Medicine in Developing Countries,  [[15]](https://ejhp.bmj.com/content/26/Suppl_1/A204.2) C. Jimenez Nunez et al. (2019). "5PSQ-007 Sebelipase alfa as enzyme replacement therapy in three paediatric patients" European Journal of Hospital Pharmacy, 26 |
|  | Evaluation  **Summarization Accuracy & Completeness:**  **Hallucination Rate:**  **Readability & Usability:**  **Literature Coverage & Relevance:**  **Time Efficiency:** | |
| **5** | The c.256C>T variant in the WAS gene, also known as p.Arg86*, is a homozygous variant associated with autosomal recessive severe congenital neutropenia (SCN)[1](https://pubmed.ncbi.nlm.nih.gov/39980410/). Forty-two out of 45 patients (93.3%) with this condition had a premature stop codon, specifically the p.Trp44*(c.132G>A) variant in the HAX1 gene, while other homozygous variants included p.Arg86*(c.256C>T) and p.Glu60Aspfs*25(c.180delA)[1](https://pubmed.ncbi.nlm.nih.gov/39980410/). In a study of patients with HAX1-SCN, elevated serum follicle-stimulating hormone (FSH) levels were found in 90.9% of prepubertal females and 92.3% of pubertal females, suggesting impending or established premature ovarian insufficiency (POI), with all female patients exhibiting low anti-Mullerian hormone (AMH) levels; male patients did not show gonadal insufficiency[1](https://pubmed.ncbi.nlm.nih.gov/39980410/). Further research is needed to fully elucidate the penetrance, expressivity, and clinical variability associated with the c.256C>T variant, as well as to determine optimal management strategies for affected individuals. | 1. <https://pubmed.ncbi.nlm.nih.gov/39980410/> 2. <https://www.ncbi.nlm.nih.gov/pmc/articles/PMC5521977/> 3. <https://pubmed.ncbi.nlm.nih.gov/32323592/> 4. <https://www.ncbi.nlm.nih.gov/pmc/articles/PMC4465929/> 5. <https://pubmed.ncbi.nlm.nih.gov/16273109/> 6. <https://www.ncbi.nlm.nih.gov/pmc/articles/PMC10316620/> 7. <https://www.ncbi.nlm.nih.gov/pmc/articles/PMC7042727/> 8. <https://www.ncbi.nlm.nih.gov/pmc/articles/PMC4326268/> |
|  | Evaluation  **Summarization Accuracy & Completeness:**  **Hallucination Rate:**  **Readability & Usability:**  **Literature Coverage & Relevance:**  **Time Efficiency:** | |
