## Supplementary material for "Benchmarking generative AI tools for literature retrieval and summarization in genomic variant interpretation": guidelines_for_reviewers.docx

**Guide to evaluating reports generated by AI tools**

Each report contains an analysis of a genetic variant generated by five different tools. We ask you to evaluate them individually, but comparing the outputs can help you build a clearer reference scale.

The aim of the study is to evaluate the ability of different tools to provide a summary of the main elements related to a genetic variant supported by scientific literature.

### 1. Summarization Accuracy

### Key question: Does the tool accurately summarize the results of relevant studies? Does it include essential details?

### A low score is given to summaries that are superficial, generic, or lack clear references. A high score corresponds to a clear and well-supported summary that provides all useful information without significant omissions.

### Scale: 1 = Incorrect or very incomplete summary | 5 = Clear, accurate, and complete summary

### 2. Hallucinations Robustness

### Key question: Does the tool introduce errors or unsupported information?

### A serious error is the invention of non-existent studies or data. A less serious error is citing generic or less relevant articles. A high score indicates that all information reported is verifiable and accurate.

### Scale: 1 = High presence of errors | 5 = No errors

### 3. Readability & Usability

### Key question: Is the report clear, well structured, and useful for a physician or researcher?

### A low score is given to texts that are confusing, redundant, or difficult to read. A high score indicates a well-organized report with information that is easy to find and understand.

### Scale: 1 = Difficult to read | 5 = Optimal clarity and structure

### 4. Literature Coverage & Relevance

**Key question:** Does the tool retrieve a complete and relevant set of articles?

A low score indicates few relevant studies or many irrelevant sources. Some variants have abundant literature, while others are less studied. In these cases, the tool should not be penalized for the scarcity of available publications, but for the possible failure to select the most useful ones. For a more accurate assessment, we recommend comparing the references provided with those retrievable via Google Scholar, PubMed, or Bing, checking for any omissions or discrepancies with existing literature. A high score corresponds to an up-to-date and targeted selection.

An important point to note is that the list of variants has been intentionally structured to include both variants with many references and others with few, in order to evaluate the performance of the tools under conditions of varying information availability. Consequently, when evaluating the coverage and relevance of the literature, and more generally the quality of the information provided, remember that the purpose of these tools is not simply to provide data at any cost. The variability you will encounter in the amount and format of information depends on how the tools present the references. For this reason, elements such as the absence of precise citations and the presence of incorrect or unusable references should be considered in the evaluation.

### Scale: 1 = Poor coverage, irrelevant sources | 5 = Complete and targeted coverage

### 5. Time Efficiency

### Key question: How much time does each tool save compared to a manual review?

### Scale: Relative assessment compared to the time needed to do the same job manually.

### 1- No time saved, need to retrieve key information manually, 5- Very significant reduction in the time needed to write an accurate summary

### To assess this indicator, we recommend comparing different summaries, evaluating how readily available key information is, and evaluating one tool against the others. For example, if the summary contains all the information in a clear and referenced manner, making it quick to access and verify its accuracy, the score will be high. If the summary only partially contains important information, poorly referenced (requiring manual research and verification to confirm the accuracy of the information), the score will be lower.

**Considerations on the variability of variants and references**

We intentionally structured the list of variants to include both those with numerous references and others with few publications available. This allows us to evaluate the performance of the tools under conditions of varying information availability.

In each summary, to make the analysis more robust, the order of the tools used is random (so summary 1 is not always generated by the same tool, and so on), with the aim of making the evaluation independent of any “memory effect” linked to the order of the tools.

Consequently, when evaluating the coverage and relevance of the literature, and more generally the quality of the information provided, it is important to consider that the objective of these tools is not simply to return data in every case, but to effectively select and synthesize the most relevant information.

La variabilità riscontrata nella quantità e nel formato delle informazioni dipende dal modo in cui i tool gestiscono le referenze. Per questo motivo, fattori come l’assenza di citazioni precise, la presenza di referenze errate o non utilizzabili devono essere considerati nella valutazione complessiva della qualità del tool.

The variability found in the quantity and format of information depends on how the tools manage references. For this reason, factors such as the absence of precise citations and the presence of incorrect or unusable references must be considered in the overall assessment of the tool's quality.
