## Supplementary material for "Benchmarking generative AI tools for literature retrieval and summarization in genomic variant interpretation": ALK_p.R401L.docx

| **Tool** | **Summary** | **References** |
| --- | --- | --- |
| **1** | The ALK (Anaplastic Lymphoma Kinase) gene encodes a receptor tyrosine kinase that plays a crucial role in the development of the brain and the conduction of nervous system signals. ALK is part of the insulin receptor superfamily and is primarily expressed in neural tissues. The ALK protein is involved in various cellular processes, including proliferation, differentiation, and survival. The p.R401L variant of the ALK gene refers to a missense mutation where arginine is substituted with leucine at amino acid position 401. This variant is classified using the nomenclature p.R401L, following the guidelines of the Human Genome Variation Society (HGVS). The frequency of the p.R401L variant in the general population is not well-documented, and its classification regarding pathogenicity is currently uncertain due to limited available data. The impact of the p.R401L variant on ALK function and its potential role in disease is not fully understood. There is a lack of specific peer-reviewed publications directly addressing the p.R401L variant, which limits the ability to provide detailed insights into its clinical significance. Given the uncertainties surrounding this variant, further investigation through functional studies and clinical correlation is recommended to better understand its implications. | No references available |
|  | Evaluation  **Summarization Accuracy & Completeness:**  **Hallucination Rate:**  **Readability & Usability:**  **Literature Coverage & Relevance:**  **Time Efficiency:** | |
| **2** | The ALK (anaplastic lymphoma kinase) gene encodes a receptor tyrosine kinase involved in neural development and, when aberrantly activated, plays a key role in oncogenesis through constitutive signaling promoting cell proliferation and survival. Mutations, rearrangements, and amplifications of ALK are implicated in various malignancies, most notably anaplastic large cell lymphoma and non-small cell lung cancer. The p.R401L variant in ALK denotes a missense mutation where arginine is substituted by leucine at position 401. Although this specific variant is less well-characterized than other oncogenic ALK alterations, recent studies suggest its presence in head and neck squamous cell carcinoma (HNSCC), a cancer type where ALK genomic aberrations contribute to tumorigenesis. In a study of HNSCC, p.R401L was identified among the spectrum of ALK mutations and appeared to be associated with advanced disease stages, highlighting its potential as a pathogenic variant and biomarker of aggressive tumor behavior [1]. However, this variant is rarely reported in broader population-scale databases, and no specific functional characterization or therapeutic response data currently exists. As such, the clinical significance of p.R401L remains uncertain, requiring further validation through functional assays and larger cohort analyses. In the absence of comprehensive mechanistic data or clinical correlation beyond the HNSCC context, this variant should be classified as a Variant of Uncertain Significance (VUS) per ACMG/AMP guidelines, pending additional evidence. Given its possible involvement in oncogenic signaling, surveillance for future studies and consideration in expanded tumor sequencing panels is recommended for research and diagnostic follow-up. | [[1]](https://aacrjournals.org/cancerres/article/81/13_Supplement/2422/668318) Wang et al., Genetic aberrations of Anaplastic Lymphoma Kinase (ALK) are functional drivers in head and neck squamous cell carcinoma (HNSCC) |
|  | Evaluation  **Summarization Accuracy & Completeness:**  **Hallucination Rate:**  **Readability & Usability:**  **Literature Coverage & Relevance:**  **Time Efficiency:** | |
| **3** | The ALK gene encodes the anaplastic lymphoma kinase, a receptor tyrosine kinase involved in neural development and cellular signaling pathways that regulate proliferation, differentiation, and survival. It is normally expressed in the nervous system but can become oncogenic through chromosomal rearrangements, mutations, or amplifications, leading to the formation of fusion proteins such as EML4-ALK in non-small cell lung cancer (NSCLC) and other malignancies. The p.R401L variant refers to a missense mutation where arginine at position 401 is substituted by leucine, a change that could potentially alter kinase activity or protein interactions; however, specific functional and pathogenic data on this variant are limited. ALK mutations are classified variably, with some mutations being oncogenic drivers, especially in the context of fusion proteins, and others being benign or of uncertain significance. The frequency of this particular variant is not well documented in the literature, and its impact on ALK function remains uncertain, emphasizing the need for further functional studies to clarify its role in oncogenesis or disease susceptibility. In the context of ALK-driven cancers, targeted therapies such as crizotinib and other ALK inhibitors have shown efficacy, particularly in cases involving fusion proteins or activating mutations. The current evidence does not definitively associate p.R401L with increased kinase activity or oncogenic potential, and its classification remains uncertain, warranting additional research to determine its clinical significance. Given the limited data, it is advisable to interpret this variant cautiously, considering the possibility of it being a variant of uncertain significance until more functional and clinical evidence becomes available. Overall, the ALK gene plays a crucial role in neural development and oncogenesis, with its pathogenic alterations being important targets for precision medicine; however, the specific contribution of the p.R401L variant requires further investigation to establish its relevance in disease. | 1. <https://pubmed.ncbi.nlm.nih.gov/37773266/> 2. <https://www.ncbi.nlm.nih.gov/pmc/articles/PMC10526845/> 3. <https://www.ncbi.nlm.nih.gov/pmc/articles/PMC7289932/> 4. <https://www.semanticscholar.org/paper/87b07308901073e73e2b7d4a7b058e9adbe1419a> 5. <https://pubmed.ncbi.nlm.nih.gov/21513493/> 6. <https://www.ncbi.nlm.nih.gov/pmc/articles/PMC2919633/> 7. <https://www.ncbi.nlm.nih.gov/pmc/articles/PMC9884431/> 8. <https://www.ncbi.nlm.nih.gov/pmc/articles/PMC10054655/> |
|  | Evaluation  **Summarization Accuracy & Completeness:**  **Hallucination Rate:**  **Readability & Usability:**  **Literature Coverage & Relevance:**  **Time Efficiency:** | |
| **4** | The genomic variant c.1202G>T p.R401L rs188859061 is located in the ALK gene, which encodes the anaplastic lymphoma kinase, a receptor tyrosine kinase implicated in the development of the brain and exerts its effects on specific neurons in the nervous system. This kinase is also involved in the pathogenesis of several types of cancer, including neuroblastoma, non-small cell lung cancer (NSCLC), and anaplastic large cell lymphoma. The variant results in the substitution of arginine with leucine at position 401 of the ALK protein, which is a missense change.  In the context of cancer, ALK mutations can lead to aberrant activation of the kinase and subsequent oncogenic signaling. The specific variant c.1202G>T p.R401L rs188859061 has been studied for its potential impact on peptide binding affinity to HLA-A0201, which is relevant for immune recognition and response to tumor cells. The mutant peptide containing the leucine substitution demonstrated a significantly lower binding affinity to HLA-A0201 compared to the wild-type peptide, indicating a potential impact on the immune system's ability to recognize and respond to cells harboring this mutation [1].  Furthermore, the variant rs188859061 has been reported in ClinVar with a classification of uncertain significance, suggesting that there is currently insufficient evidence to determine its impact on health definitively. This underscores the need for further research and data accumulation to clarify the clinical relevance of this variant.  In summary, the c.1202G>T p.R401L rs188859061 variant in the ALK gene results in a missense change that may affect the immune system's recognition of cancer cells due to altered peptide-HLA binding affinity. However, its clinical significance remains uncertain, and additional studies are required to elucidate its role in disease pathogenesis and potential as a therapeutic target [1]. | [[1]](https://pubmed.ncbi.nlm.nih.gov/23243591/) Khalili JS et al. (2012). "In silico prediction of tumor antigens derived from functional missense mutations of the cancer gene census." Oncoimmunology, 1(8)  [[2]](https://pubmed.ncbi.nlm.nih.gov/29575851/) Ku BM et al. (2018). "Molecular Screening of Small Biopsy Samples Using Next-Generation Sequencing in Korean Patients with Advanced Non-small Cell Lung Cancer: Korean Lung Cancer Consortium (KLCC-13-01)." Journal of pathology and translational medicine, 52(3)  [[3]](https://pubmed.ncbi.nlm.nih.gov/35865984/) Saifullah et al. (2022). "Integrated analysis of the clinical consequence and associated gene expression of ALK in ALK-positive human cancers." Heliyon, 8(7) [SUPP]  [[4]](https://pubmed.ncbi.nlm.nih.gov/31034466/) Brown AL et al. (2019). "Finding driver mutations in cancer: Elucidating the role of background mutational processes." PLoS computational biology, 15(4) [SUPP]  [[5]](https://pubmed.ncbi.nlm.nih.gov/34389641/) Ganz J et al. (2022). "Rates and Patterns of Clonal Oncogenic Mutations in the Normal Human Brain." Cancer discovery, 12(1) [SUPP]  [[6]](https://pubmed.ncbi.nlm.nih.gov/25054154/) Doss CG et al. (2014). "Integrating in silico prediction methods, molecular docking, and molecular dynamics simulation to predict the impact of ALK missense mutations in structural perspective." BioMed research international, 2014() [SUPP]  [[7]](https://pubmed.ncbi.nlm.nih.gov/21727090/) Reva B et al. (2011). "Predicting the functional impact of protein mutations: application to cancer genomics." Nucleic acids research, 39(17) [SUPP]  [[8]](https://pubmed.ncbi.nlm.nih.gov/29617658/) Way GP et al. (2018). "Machine Learning Detects Pan-cancer Ras Pathway Activation in The Cancer Genome Atlas." Cell reports, 23(1) [SUPP]  [[9]](https://pubmed.ncbi.nlm.nih.gov/32579932/) Triki M et al. (2020). "mTOR Signaling and SREBP Activity Increase FADS2 Expression and Can Activate Sapienate Biosynthesis." Cell reports, 31(12) [SUPP] |
|  | Evaluation  **Summarization Accuracy & Completeness:**  **Hallucination Rate:**  **Readability & Usability:**  **Literature Coverage & Relevance:**  **Time Efficiency:** | |
| **5** | The ALK gene encodes the anaplastic lymphoma kinase, a receptor tyrosine kinase involved in neural development and signal transduction. The p.R401L variant is a missense mutation resulting in the substitution of arginine with leucine at position 401, located within the MAM domain 1 of the ALK protein. Functional studies have demonstrated that this variant enhances cellular proliferation, clonogenic survival, and invasiveness in vitro, suggesting a gain-of-function effect. Specifically, ectopic expression of ALK R401L in head and neck squamous cell carcinoma cell lines led to increased oncogenic behaviors, supporting its potential role in tumorigenesis. However, the clinical significance of the p.R401L variant remains uncertain due to limited data on its prevalence and impact in patient populations. Further research is necessary to elucidate its pathogenicity and potential as a therapeutic target. | <https://www.ncbi.nlm.nih.gov/gene/249>  <https://www.lung.org/lung-health-diseases/lung-disease-lookup/lung-cancer/symptoms-diagnosis/biomarker-testing/alk-lung-cancer>  <https://primarycarenotebook.com/pages/general-information/asymptomatic-raised-alkaline-phosphatase>  <https://aacrjournals.org/cancerres/article/81/13_Supplement/2422/668318/Abstract-2422-Genetic-aberrations-of-Anaplastic>  <https://molecular-cancer.biomedcentral.com/articles/10.1186/s12943-018-0776-2>  <https://medlineplus.gov/genetics/gene/alk/>  <https://medlineplus.gov/lab-tests/alkaline-phosphatase/>  <https://www.oncokb.org/gene/ALK/R401L>  <https://www.oncokb.org/gene/ALK/R401Q>  <https://my.clevelandclinic.org/health/diagnostics/22029-alkaline-phosphatase-alp>  <https://cancer.sanger.ac.uk/cosmic/mutation/overview?id=6387463>  <https://cancer.sanger.ac.uk/cosmic3d/protein/ALK>  <https://www.ncbi.nlm.nih.gov/gene/238>  <https://pmc.ncbi.nlm.nih.gov/articles/PMC6450058/>  <https://pubmed.ncbi.nlm.nih.gov/27573755/>  <https://pmc.ncbi.nlm.nih.gov/articles/PMC5171696/>  <https://pmc.ncbi.nlm.nih.gov/articles/PMC2174237/>  <https://ckb-core.genomenon.com/gene/show?geneId=238>  <https://ckb-core.genomenon.com/molecularProfile/show/41906>  <https://www.mayocliniclabs.com/test-catalog/download-setup.php?format=pdf&unit_code=8340>  <https://www.alkpositive.org/blog/2021/1/15/normal-alk-function>  <https://www.sciencedirect.com/science/article/pii/S0923753419316291>  <https://www.sonicbookings.com.au/our-tests/elft-euc-lft/alkaline-phosphatase/>  <https://en.wikipedia.org/wiki/Anaplastic_lymphoma_kinase>  <https://www.genecards.org/cgi-bin/carddisp.pl?gene=ALK>  <https://eprints.whiterose.ac.uk/id/eprint/139972/8/Thwaites%20et%20al%20Blood.pdf> |
|  | Evaluation  **Summarization Accuracy & Completeness:**  **Hallucination Rate:**  **Readability & Usability:**  **Literature Coverage & Relevance:**  **Time Efficiency:** | |
