## Supplementary material for "Benchmarking generative AI tools for literature retrieval and summarization in genomic variant interpretation": APC p.W699.docx

| **Tool** | **Summary** | **References** |
| --- | --- | --- |
| **1** | The APC gene, located on chromosome 5q22, encodes the Adenomatous Polyposis Coli (APC) protein, a crucial tumor suppressor involved in the Wnt signaling pathway. The APC protein regulates cell growth and differentiation by controlling the levels of beta-catenin, a key component in cell adhesion and gene transcription. Mutations in the APC gene disrupt its function, leading to uncontrolled cell proliferation and the development of polyps, primarily in the colon and rectum, which can progress to colorectal cancer if left untreated.  The APC p.W699* variant is a nonsense mutation resulting from a single nucleotide substitution (c.2097G>A) that introduces a premature stop codon at position 699 in the APC protein. This truncating mutation leads to a nonfunctional protein, significantly impairing its tumor suppressor activity. The variant has been confirmed as somatic and is associated with familial adenomatous polyposis (FAP), an autosomal dominant condition characterized by the development of numerous adenomatous polyps in the colon and rectum, with a high risk of progression to colorectal cancer.  The frequency of the APC p.W699* variant is not well-documented in the general population, and it is absent from major allele frequency databases. However, its recurrence in clinical samples suggests it is a significant contributor to FAP cases. The variant's pathogenicity is supported by its impact on protein function and its co-segregation with the disease phenotype in affected families.  The clinical implications of the APC p.W699* variant include a high risk of developing colorectal cancer, with polyps often appearing in the second decade of life. Management strategies involve regular colonoscopies starting in adolescence, with prophylactic colectomy considered in cases of severe polyposis. Genetic counseling and testing for at-risk family members are essential for early detection and intervention.  In conclusion, the APC p.W699* variant is a pathogenic nonsense mutation that severely compromises the function of the APC protein, leading to a high risk of colorectal cancer in affected individuals. Its clinical significance underscores the importance of genetic testing and surveillance in managing FAP. | [1] [COSMIC](https://cancer.sanger.ac.uk/cosmic/mutation/overview?id=105275105) database, Mutation ID: COSV57342069.  [2] [NCBI Gene](https://www.ncbi.nlm.nih.gov/gene/324), APC Adenomatous Polyposis Coli. |
|  | Evaluation  **Summarization Accuracy & Completeness:**  **Hallucination Rate:**  **Readability & Usability:**  **Literature Coverage & Relevance:**  **Time Efficiency:** | |
| **2** | The APC gene encodes a tumor suppressor protein that negatively regulates the Wnt signaling pathway by controlling beta-catenin concentrations and interacting with E-cadherin, thereby influencing cell adhesion, cell division, and other critical cellular processes[1](https://pubmed.ncbi.nlm.nih.gov/19951906/)[7](https://pubmed.ncbi.nlm.nih.gov/8259518/)[9](https://en.wikipedia.org/wiki/Adenomatous_polyposis_coli). Mutations in APC are frequently found in both sporadic and hereditary colorectal cancers[1](https://pubmed.ncbi.nlm.nih.gov/19951906/). The APC protein forms a complex with glycogen synthase kinase 3-alpha/beta (GSK-3α/β) and Axin, which promotes the phosphorylation, ubiquitination, and degradation of beta-catenin[13](https://pmc.ncbi.nlm.nih.gov/articles/PMC5803335/). This prevents beta-catenin from entering the nucleus and activating genes that stimulate cell division[9](https://en.wikipedia.org/wiki/Adenomatous_polyposis_coli)[13](https://pmc.ncbi.nlm.nih.gov/articles/PMC5803335/). The APC protein contains domains that bind to beta-catenin and Axin[13](https://pmc.ncbi.nlm.nih.gov/articles/PMC5803335/). The p.W699* variant represents a premature stop codon, leading to a truncated protein, and occurs in approximately 30% of APC mutations[1](https://pubmed.ncbi.nlm.nih.gov/19951906/). Research has shown that aminoglycosides and macrolides can induce read-through of premature stop codons, potentially restoring full-length APC protein levels and ameliorating tumorigenic symptoms in colorectal cancer cell lines and animal models[1](https://pubmed.ncbi.nlm.nih.gov/19951906/). However, no specific academic research is available for the APC p.W699* variant. Further *in vitro* and *in vivo* studies are needed to determine the exact functional consequence of the APC p.W699* variant.[1](https://pubmed.ncbi.nlm.nih.gov/19951906/) | [1] <https://pubmed.ncbi.nlm.nih.gov/19951906/>  [2] <https://pubmed.ncbi.nlm.nih.gov/26711662/>  [3] <https://pubmed.ncbi.nlm.nih.gov/22467108/>  [4] <https://www.semanticscholar.org/paper/11681424e37a2cb5a5a16d127577fc5a1a7c72d3>  [5] <https://www.ncbi.nlm.nih.gov/pmc/articles/PMC9178358/>  [6] <https://www.ncbi.nlm.nih.gov/pmc/articles/PMC4999466/>  [7] <https://pubmed.ncbi.nlm.nih.gov/8259518/>  [8] <https://pubmed.ncbi.nlm.nih.gov/23397547/>  [9] <https://en.wikipedia.org/wiki/Adenomatous_polyposis_coli>  [10] <https://pmc.ncbi.nlm.nih.gov/articles/PMC7737709/>  [11] <https://medlineplus.gov/download/genetics/gene/apc.pdf>  [12] <https://www.ncbi.nlm.nih.gov/books/NBK1345/>  [13] <https://pmc.ncbi.nlm.nih.gov/articles/PMC5803335/>  [14] <https://journal.waocp.org/article_90092.html>  [15] <https://pubmed.ncbi.nlm.nih.gov/29318445/>  [16] <https://medlineplus.gov/genetics/gene/apc/> |
|  | Evaluation  **Summarization Accuracy & Completeness:**  **Hallucination Rate:**  **Readability & Usability:**  **Literature Coverage & Relevance:**  **Time Efficiency:** | |
| **3** | The APC gene encodes a tumor suppressor protein integral to regulating cell proliferation, adhesion, migration, and chromosomal stability. Functioning as a negative regulator of the Wnt signaling pathway, APC facilitates the degradation of β-catenin, thereby preventing uncontrolled cellular growth [5]. The p.W699* variant, resulting from a substitution at nucleotide position 2097 (c.2097G>A), introduces a premature stop codon at amino acid position 699. This truncation likely leads to a nonfunctional protein, as it eliminates critical domains responsible for β-catenin regulation and microtubule binding [10]. Such loss-of-function mutations in APC are implicated in familial adenomatous polyposis (FAP), an autosomal dominant disorder characterized by the development of numerous colorectal adenomas and an elevated risk of colorectal cancer [12]. The p.W699* variant has been identified in individuals with FAP, underscoring its pathogenic significance [2]. While the exact population frequency of p.W699* is not well-documented, APC mutations, in general, are prevalent in colorectal carcinogenesis. The absence of this variant in large population databases suggests it is rare and likely deleterious. Given the established role of APC truncating mutations in colorectal tumorigenesis, the p.W699* variant should be considered pathogenic. However, comprehensive genetic counseling and further functional studies are recommended to elucidate its full clinical impact. citeturn0search5turn0search2turn0search12 | [1] [pmc.ncbi.nlm.nih.gov](https://pmc.ncbi.nlm.nih.gov/articles/PMC7306982/)  [2] [Oxford Academic](https://academic.oup.com/jnci/article/109/8/djw332/3113843)  [3] [medlineplus.gov](https://medlineplus.gov/genetics/gene/apc/)  [4] j[ournals.biologists.com](https://journals.biologists.com/jcs/article/120/19/3327/35163/Adenomatous-polyposis-coli-APC-a-multi-functional)  [5] [molecular-cancer.biomedcentral.com](https://molecular-cancer.biomedcentral.com/articles/10.1186/1476-4598-2-41)  [6] [ncbi.nlm.nih.gov](https://www.ncbi.nlm.nih.gov/clinvar/variation/411387/?new_evidence=true)  [7] [pubmed.ncbi.nlm.nih.gov](https://pubmed.ncbi.nlm.nih.gov/29318445/)  [8] [medlineplus.gov](https://medlineplus.gov/download/genetics/gene/apc.pdf)  [9] [bioinfo.uth.edu](https://bioinfo.uth.edu/TSGene/gene_mutation.cgi?gene=324)  [10] [cancer.sanger.ac.uk](https://cancer.sanger.ac.uk/cosmic3d/protein/APC)  [11] [en.wikipedia.org](https://en.wikipedia.org/wiki/Adenomatous_polyposis_coli)  [12] [jcga-scc.jp](https://www.jcga-scc.jp/en/gene/APC)  [13] [bibliome.ai](https://bibliome.ai/GRCh38/publication/28389531)  [14] [bibliome.ai](https://www.bibliome.ai/hg19/publication/36181537)  [15] [Oxford Academic](https://academic.oup.com/hmg/article/10/7/721/558552)  [16] [ClinVar Miner](https://clinvarminer.genetics.utah.edu/variants-by-gene/APC/condition/Hereditary%20cancer-predisposing%20syndrome/pathogenic)  [17] [modernpathology.org](https://www.modernpathology.org/cms/10.1016/j.modpat.2024.100586/attachment/285cd1f2-9c44-4904-acf8-e535e6f39913/mmc1.pdf) |
|  | Evaluation  **Summarization Accuracy & Completeness:**  **Hallucination Rate:**  **Readability & Usability:**  **Literature Coverage & Relevance:**  **Time Efficiency:** | |
| **4** | The APC gene (Adenomatous Polyposis Coli) is a critical tumor suppressor gene involved in the regulation of cell growth and adhesion, primarily through the Wnt signaling pathway. Germline mutations in APC are the primary cause of familial adenomatous polyposis (FAP), an autosomal dominant disorder characterized by the development of hundreds to thousands of colorectal adenomas and a near 100% lifetime risk of colorectal cancer if prophylactic colectomy is not performed. The APC p.W699* variant results in a premature stop codon, likely leading to a truncated protein or nonsense-mediated decay, thereby abrogating its tumor suppressor function. Loss-of-function mutations in APC, including truncating variants, are well-documented drivers of colorectal cancer pathogenesis, particularly in FAP and sporadic colorectal carcinomas [1]. While no direct studies specifically investigate the pathogenicity of the APC p.W699* variant, truncating mutations in the 5' region of APC are generally associated with a classical or severe FAP phenotype. In a genomic profiling study of familial adenomatous polyposis, APC mutations were found to be highly recurrent in carcinogenesis, further supporting the deleterious nature of truncating variants [2]. Additional research has implicated APC mutations in oncogenic processes beyond colorectal cancer, including intestinal metaplasia with high-grade dysplasia [3]. However, no functional studies directly evaluating APC p.W699* have been published, and further investigation is needed to confirm its clinical significance. Given its likely loss-of-function effect and the known role of truncating APC mutations in colorectal tumorigenesis, this variant should be considered pathogenic until further evidence clarifies its precise impact. Genetic counseling and clinical surveillance recommendations should align with guidelines for FAP-associated APC variants. . | [1] Li J, Wang R, Zhou X, Wang W, Gao S, Mao Y, Wu X. *Genomic and transcriptomic profiling of carcinogenesis in patients with familial adenomatous polyposis*. Gut. 2020.  [2] Verwiel ETP, Spruijt L. *Robbert DA Weren1, Marjolijn JL Ligtenberg1, 2, C. Marleen Kets1, Richarda M. de Voer1, Eugène TP Verwiel1, Liesbeth Spruijt1, Wendy AG van Zelst-Stams1*. Radboud Repository. 2016.  [3] Acosta AM, Sholl LM, Fanelli GN, Gordetsky JB. *Intestinal metaplasia of the urinary tract harbors potentially oncogenic genetic variants*. SciDirect. 2021. |
|  | Evaluation  **Summarization Accuracy & Completeness:**  **Hallucination Rate:**  **Readability & Usability:**  **Literature Coverage & Relevance:**  **Time Efficiency:** | |
| **5** | The genomic variant c.2097G>A p.W699* rs1060503282 is located in the APC gene. This variant is a nonsense mutation, which results in the substitution of a tryptophan (W) for a stop codon (asterisk) at amino acid position 699. The introduction of a premature stop codon is predicted to lead to a truncated protein product through the process of nonsense-mediated mRNA decay, which can have significant effects on the protein's function.  The APC gene plays a critical role in the Wnt signaling pathway, which is involved in cell growth and differentiation. APC is a tumor suppressor gene, and its product is essential for the regulation of beta-catenin, a protein that mediates the transcription of genes that are crucial for cell proliferation and survival. Mutations in the APC gene are known to disrupt this regulatory pathway, leading to uncontrolled cell growth and the development of tumors.  In the context of colorectal adenocarcinoma with squamous differentiation (ASC), the variant p.W699* was identified as one of the most common APC variants, observed in 9 out of 22 (40.9%) primary ASC samples [3]. This variant, along with another APC variant, was the second most frequent genetic alteration after TP53 mutations in the studied cohort. The presence of this variant is indicative of its potential role in the pathogenesis of colorectal cancer, particularly in the subset of ASCs.  The APC p.W699* variant has been reported in ClinVar with a total of 4 submissions citing it as pathogenic and 2 submissions with uncertain significance. This discrepancy in clinical interpretation highlights the complexity of variant classification and the need for further functional studies to elucidate the precise impact of this variant on protein function and disease pathology.  In summary, the c.2097G>A p.W699* rs1060503282 variant in the APC gene is a nonsense mutation that likely contributes to the pathogenesis of colorectal cancer by disrupting the tumor suppressor function of the APC protein, leading to aberrant cell proliferation and tumor development. | [[1]](https://pubmed.ncbi.nlm.nih.gov/25333069/) Freudenberg-Hua Y et al. (2014). "Disease variants in genomes of 44 centenarians." Molecular genetics & genomic medicine, 2(5)  [[2]](https://pubmed.ncbi.nlm.nih.gov/24317090/) Andersson EI et al. (2013). "Novel somatic mutations in large granular lymphocytic leukemia affecting the STAT-pathway and T-cell activation." Blood cancer journal, 3(12)  [[3]](https://pubmed.ncbi.nlm.nih.gov/36790480/) Angerilli V et al. (2023). "Colorectal adenosquamous carcinoma: genomic profiling of a rare histotype of colorectal cancer." Virchows Archiv : an international journal of pathology, 482(5)  [[4]](https://pubmed.ncbi.nlm.nih.gov/27302369/) Schell MJ et al. (2016). "A multigene mutation classification of 468 colorectal cancers reveals a prognostic role for APC." Nature communications, 7() *Variant reported in Supplementary Information*  [[5]](https://pubmed.ncbi.nlm.nih.gov/38225666/) Yang L et al. (2024). "Phase separation as a possible mechanism for dosage sensitivity." Genome biology, 25(1) Variant reported in Supplementary Information  [[6]](https://pubmed.ncbi.nlm.nih.gov/27397505/) Iorio F et al. (2016). "A Landscape of Pharmacogenomic Interactions in Cancer." Cell, 166(3) *Variant reported in Supplementary Information*  [[7]](https://pubmed.ncbi.nlm.nih.gov/36124685/) Kashani E et al. (2023). "Integrated longitudinal analysis of adult grade 4 diffuse gliomas with long-term relapse interval revealed upregulation of TGF-beta signaling in recurrent tumors." Neuro-oncology, 25(4) *Variant reported in Supplementary Information*  [[8]](https://pubmed.ncbi.nlm.nih.gov/28186126/) Schutte M et al. (2017). "Molecular dissection of colorectal cancer in pre-clinical models identifies biomarkers predicting sensitivity to EGFR inhibitors." Nature communications, 8() *Variant reported in Supplementary Information*  [[9]](https://pubmed.ncbi.nlm.nih.gov/30925167/) Schumacher D et al. (2019). "Heterogeneous pathway activation and drug response modelled in colorectal-tumor-derived 3D cultures." PLoS genetics, 15(3) *Variant reported in Supplementary Information*  [[10]](https://pubmed.ncbi.nlm.nih.gov/29575536/) Wolff RK et al. (2018). "Mutation analysis of adenomas and carcinomas of the colon: Early and late drivers." Genes, chromosomes & cancer, 57(7) *Variant reported in Supplementary Information*  [[11]](https://pubmed.ncbi.nlm.nih.gov/26269718/) Patwardhan A et al. (2015). "Achieving high-sensitivity for clinical applications using augmented exome sequencing." Genome medicine, 7(1) *Variant reported in Supplementary Information*  [[12]](https://pubmed.ncbi.nlm.nih.gov/30202008/) Palin K et al. (2018). "Contribution of allelic imbalance to colorectal cancer." Nature communications, 9(1) *Variant reported in Supplementary Information*  [[13]](https://pubmed.ncbi.nlm.nih.gov/32686686/) Zaidi SH et al. (2020). "Landscape of somatic single nucleotide variants and indels in colorectal cancer and impact on survival." Nature communications, 11(1) *Variant reported in Supplementary Information* |
|  | Evaluation  **Summarization Accuracy & Completeness:**  **Hallucination Rate:**  **Readability & Usability:**  **Literature Coverage & Relevance:**  **Time Efficiency:** | |
