## Supplementary material for "Benchmarking generative AI tools for literature retrieval and summarization in genomic variant interpretation": ARID1A p.R1989_.docx

| **Tool** | **Summary** | **References** |
| --- | --- | --- |
| **1** | The ARID1A gene encodes a key subunit of the SWI/SNF chromatin remodeling complex, which modulates transcription by altering chromatin structure. ARID1A functions as a tumor suppressor and is involved in DNA repair, cell cycle regulation, and maintenance of genomic stability. The p.R1989* variant is a nonsense mutation resulting in a premature stop codon at position 1989, leading to a truncated protein likely subject to nonsense-mediated decay, effectively resulting in loss of function. While this specific variant has not been individually reported in peer-reviewed literature, similar truncating mutations in ARID1A are recurrent across various cancers, notably in ovarian clear cell carcinoma and endometrial carcinoma, where they are associated with impaired interferon signaling, reduced T cell infiltration, and poorer clinical outcomes . The absence of direct studies on p.R1989* limits definitive conclusions about its pathogenicity; however, given its predicted loss-of-function effect and the established role of ARID1A truncations in tumorigenesis, this variant is likely deleterious. Further functional studies are recommended to confirm its impact. | <https://www.ncbi.nlm.nih.gov/gene/8289>  <https://www.researchgate.net/publication/365348788_SWISNF_complex_gene_variations_are_associated_with_a_higher_tumor_mutational_burden_and_a_better_response_to_immune_checkpoint_inhibitor_treatment_a_pan-cancer_analysis_of_next-generation_sequencing_d>  <https://www.researchgate.net/publication/365746142_Validation_and_Clinical_Application_of_ONCOaccuPanel_for_Targeted_Next-Generation_Sequencing_of_Solid_Tumors>  <https://jbiomedsci.biomedcentral.com/articles/10.1186/s12929-022-00856-5>  <https://www.spandidos-publications.com/10.3892/wasj.2020.37>  <https://journals.plos.org/plosgenetics/article?id=10.1371%2Fjournal.pgen.1007779>  <https://journals.plos.org/plosmedicine/article?id=10.1371%2Fjournal.pmed.1002206>  <https://pmc.ncbi.nlm.nih.gov/articles/PMC7190935/>  <https://www.ncbi.nlm.nih.gov/clinvar/variation/VCV000225842>  <https://www.frontiersin.org/journals/oncology/articles/10.3389/fonc.2021.745187/full>  <https://www.frontiersin.org/journals/oncology/articles/10.3389/fonc.2023.1136248/full>  <https://www.frontiersin.org/journals/oncology/articles/10.3389/fonc.2021.693295/full>  <https://www.genecards.org/cgi-bin/carddisp.pl?gene=ARID1A>  <https://clinvarminer.genetics.utah.edu/variants-by-gene/ARID1A/significance/likely%20pathogenic>  <https://psb.stanford.edu/previous/psb17/conference-materials/proceedings/5precmed.pdf>  <https://www.sciencedirect.com/topics/biochemistry-genetics-and-molecular-biology/arid1a> |
|  | Evaluation  **Summarization Accuracy & Completeness:**  **Hallucination Rate:**  **Readability & Usability:**  **Literature Coverage & Relevance:**  **Time Efficiency:** | |
| **2** | The genomic variant c.5965C>T p.R1989* rs879255270 in the ARID1A gene is a nonsense mutation resulting in a premature stop codon at the arginine residue at position 1989 of the ARID1A protein [6]. This variant has been identified as pathogenic according to ClinVar, with two independent submissions supporting this classification. The mutation is associated with a loss of ARID1A expression, which is consistent with the tumor suppressor role of the ARID1A gene [3].  ARID1A, located on chromosome 1p36.11, encodes a 2285 amino acid protein that is a member of the SWI/SNF family involved in chromatin remodeling [7]. The protein plays a crucial role in various cellular processes, including DNA repair, differentiation, development, and tumor suppression. Truncating mutations like p.R1989*, which lead to loss of function, support the gene's role in tumor suppression by potentially disrupting pathways such as CDKN1A and SMAD3, which are related to cell cycle regulation, and by aberrant activation of the PI3K/AKT pathway [7].  The p.R1989* variant has been frequently observed in endometrial cancer, with 41 entries in the Cosmic database, and has also been reported in large intestine tumors, pancreatic ductal adenocarcinomas, breast carcinomas, and gastric cancer [3]. This mutation's prevalence in various cancer types underscores its significance in oncogenesis.  In silico tools such as PolyPhen2 and Varsome have classified the p.R1989* variant as pathogenic, and this classification is further supported by immunohistochemistry studies showing a loss of ARID1A protein expression [3]. The presence of this variant has been associated with sensitivity to EZH2 and BET inhibitors, suggesting potential therapeutic implications [3].  In summary, the c.5965C>T p.R1989* rs879255270 variant in the ARID1A gene is a pathogenic nonsense mutation that leads to a truncated protein product and loss of function, contributing to the development and progression of various cancers. The mutation's impact on ARID1A expression and its association with cancer phenotypes highlight its importance in genomic variant interpretation and potential targeted therapies. | [[1]](https://pubmed.ncbi.nlm.nih.gov/32868822/) Yachida N et al. (2020). "ARID1A protein expression is retained in ovarian endometriosis with ARID1A loss-of-function mutations: implication for the two-hit hypothesis." Scientific reports, 10(1)  [[2]](https://pubmed.ncbi.nlm.nih.gov/36221285/) Li S et al. (2022). "Pathologic complete response to immune checkpoint inhibitor in a stage IIIB ovarian clear cell carcinoma patient with POLE mutation resistant to platinum-based chemotherapy: a case report." Gland surgery, 11(9)  [[3]](https://pubmed.ncbi.nlm.nih.gov/35328145/) De Leo A et al. (2022). "Relevance of ARID1A Mutations in Endometrial Carcinomas." Diagnostics (Basel, Switzerland), 12(3)  [[4]](https://pubmed.ncbi.nlm.nih.gov/31772679/) Liu X et al. (2019). "Establishment and characterization of novel human primary endometrial cancer cell line (ZJB-ENC1) and its genomic characteristic." Journal of Cancer, 10(25)  [[5]](https://pubmed.ncbi.nlm.nih.gov/23104009/) Le Gallo M et al. (2012). "Exome sequencing of serous endometrial tumors identifies recurrent somatic mutations in chromatin-remodeling and ubiquitin ligase complex genes." Nature genetics, 44(12)  [[6]](https://pubmed.ncbi.nlm.nih.gov/22009941/) Jones S et al. (2012). "Somatic mutations in the chromatin remodeling gene ARID1A occur in several tumor types." Human mutation, 33(1)  [[7]](https://pubmed.ncbi.nlm.nih.gov/30057548/) Suhaimi SS et al. (2018). "Targeted Next-Generation Sequencing Identifies Actionable Targets in Estrogen Receptor Positive and Estrogen Receptor Negative Endometriod Endometrial Cancer." Frontiers in pharmacology, 9()  [[8]](https://pubmed.ncbi.nlm.nih.gov/30412573/) Poulos RC et al. (2018). "Analysis of 7,815 cancer exomes reveals associations between mutational processes and somatic driver mutations." PLoS genetics, 14(11)  [[9]](https://pubmed.ncbi.nlm.nih.gov/30886832/) Wang Y et al. (2019). "Genomic Comparison of Endometrioid Endometrial Carcinoma and Its Precancerous Lesions in Chinese Patients by High-Depth Next Generation Sequencing." Frontiers in oncology, 9()  [[10]](https://pubmed.ncbi.nlm.nih.gov/37814942/) Jin B et al. (2023). "Immune checkpoint inhibitor-related molecular markers predict prognosis in extrahepatic cholangiocarcinoma." Cancer medicine, 12(20)  [[11]](https://pubmed.ncbi.nlm.nih.gov/38113761/) Li K et al. (2024). "Prioritizing de novo potential non-canonical splicing variants in neurodevelopmental disorders." EBioMedicine, 99() [SUPP]  [[12]](https://pubmed.ncbi.nlm.nih.gov/24686850/) Lin DC et al. (2014). "Genomic and molecular characterization of esophageal squamous cell carcinoma." Nature genetics, 46(5) [SUPP]  [[13]](https://pubmed.ncbi.nlm.nih.gov/35468861/) Hamanaka K et al. (2022). "Large-scale discovery of novel neurodevelopmental disorder-related genes through a unified analysis of single-nucleotide and copy number variants." Genome medicine, 14(1) [SUPP]  [[14]](https://pubmed.ncbi.nlm.nih.gov/27397505/) Iorio F et al. (2016). "A Landscape of Pharmacogenomic Interactions in Cancer." Cell, 166(3) [SUPP]  [[15]](https://pubmed.ncbi.nlm.nih.gov/29338072/) Ross JS et al. (2018). "Targeting HER2 in colorectal cancer: The landscape of amplification and short variant mutations in ERBB2 and ERBB3." Cancer, 124(7) [SUPP] |
|  | Evaluation  **Summarization Accuracy & Completeness:**  **Hallucination Rate:**  **Readability & Usability:**  **Literature Coverage & Relevance:**  **Time Efficiency:** | |
| **3** | The ARID1A gene encodes a key subunit of the SWI/SNF chromatin remodeling complex, which plays a critical role in regulating gene expression by altering nucleosome positioning to facilitate access of transcriptional machinery to DNA. Functionally, ARID1A acts as a tumor suppressor and is involved in diverse biological processes such as DNA repair, differentiation, and cell cycle regulation. The variant p.R1989* is a nonsense mutation predicted to result in premature truncation of the ARID1A protein, likely causing loss of function. Such truncating mutations are typically associated with inactivation of tumor suppressor activity and have been identified in numerous malignancies, particularly gynecologic cancers like ovarian clear cell and endometrioid carcinomas, as well as gastric, bladder, and colorectal cancers. While this specific variant, p.R1989*, has not been directly studied in published peer-reviewed literature, its classification as a nonsense mutation in a tumor suppressor gene aligns with the established pathogenic profile of other ARID1A truncating alterations. Numerous large-scale cancer genome studies have confirmed the high prevalence of ARID1A mutations, particularly stop-gain variants, in various tumors, supporting their pathogenicity in oncogenesis through disruption of chromatin remodeling and transcriptional regulation [1]. Given the absence of specific clinical or functional data for p.R1989*, its pathogenicity remains presumed based on mechanistic expectations and analogs among better-characterized truncating variants. The variant’s clinical significance should be interpreted with caution and ideally confirmed by functional assays or patient-derived clinical correlations. Until such data are available, this variant is best considered likely pathogenic in the context of loss-of-function intolerance and recurrence patterns in tumor sequencing. Further studies are warranted to elucidate its specific contribution to disease phenotypes and potential therapeutic vulnerabilities. | [[1] Poulos RC et al. Analysis of 7,815 cancer exomes reveals associations between mutational processes and somatic driver mutations. PLOS Genetics (2018).](https://journals.plos.org/plosgenetics/article?id=10.1371/journal.pgen.1007779) |
|  | Evaluation  **Summarization Accuracy & Completeness:**  **Hallucination Rate:**  **Readability & Usability:**  **Literature Coverage & Relevance:**  **Time Efficiency:** | |
| **4** | The ARID1A gene, located on chromosome 1, encodes the AT-rich interaction domain 1A protein, a crucial subunit of the SWI/SNF chromatin remodeling complex. This complex plays a vital role in regulating gene expression by altering chromatin structure, thereby controlling DNA accessibility for transcription. ARID1A is essential for various cellular processes, including differentiation, proliferation, DNA repair, and tumor suppression. The ARID1A protein helps target the SWI/SNF complex to specific genomic locations, facilitating interactions with DNA and other proteins to modulate gene expression. The p.R1989* variant in the ARID1A gene is a nonsense mutation that introduces a premature stop codon, resulting in a truncated, nonfunctional protein. This variant is classified as likely pathogenic according to ClinVar Miner, indicating its potential to disrupt normal ARID1A function and contribute to disease. The p.R1989* variant leads to the loss of the ARID1A protein's C-terminal region, which is critical for its interaction with other SWI/SNF subunits and for its role in chromatin remodeling. This truncation impairs the complex's ability to regulate gene expression, potentially leading to abnormal cellular processes. Variants in ARID1A, including p.R1989*, have been associated with various cancers, such as ovarian, endometrial, kidney, stomach, bladder, lung, breast, and brain cancers, as well as genetic disorders like Coffin-Siris syndrome. These variants are typically somatic, occurring during a person's lifetime and present only in tumor cells. The exact frequency of the p.R1989* variant in the general population is not well-documented, but ARID1A mutations are relatively common in certain cancer types, with a prevalence of about 5% in non-hypermutated colorectal cancer tumors. The impact of the p.R1989* variant on cancer development and progression is thought to be related to the disruption of normal chromatin remodeling, which can lead to uncontrolled cell growth and differentiation. However, the precise mechanisms by which this variant contributes to cancer are not fully understood. Further research is needed to elucidate the specific effects of the p.R1989* variant on ARID1A function and its role in disease. Given the limited direct references to the p.R1989* variant in peer-reviewed literature, it is recommended to investigate this variant further through functional studies and clinical correlations. | No references available |
|  | Evaluation  **Summarization Accuracy & Completeness:**  **Hallucination Rate:**  **Readability & Usability:**  **Literature Coverage & Relevance:**  **Time Efficiency:** | |
| **5** | The ARID1A gene encodes the AT-rich interactive domain-containing protein 1A, a key subunit of the SWI/SNF chromatin remodeling complex involved in regulating gene expression by modulating chromatin accessibility at promoters and enhancers. ARID1A functions predominantly as a tumor suppressor, with loss-of-function mutations frequently observed in various cancers, including ovarian clear cell carcinoma, gastric cancer, colorectal cancer, endometrial cancer, and others. The variant p.R1989* is a nonsense mutation predicted to introduce a premature stop codon, likely resulting in truncated protein and loss of normal ARID1A function. This loss impairs chromatin remodeling activities, disrupting transcriptional regulation of genes involved in cell cycle control, apoptosis, and DNA repair pathways, thereby contributing to tumorigenesis. ARID1A mutations, including truncating variants, occur in approximately 10% of diverse tumor types and are often associated with more aggressive disease and poorer prognosis, although the prognostic significance can vary by cancer type and context. Functional studies demonstrate that ARID1A loss leads to dysregulation of pathways such as Wnt signaling and NF-κB, promoting proliferation, invasion, and reduced apoptosis in cancer cells. Moreover, ARID1A deficiency has been linked to compromised mismatch repair and increased tumor mutational burden, which may sensitize tumors to immune checkpoint inhibitors, correlating with improved progression-free survival in treated patients. Despite extensive research on ARID1A in cancer, specific peer-reviewed publications directly characterizing the p.R1989* variant are lacking, limiting definitive conclusions about its precise clinical impact. Therefore, while the p.R1989* variant is presumed pathogenic due to its truncating nature and the established tumor suppressor role of ARID1A, further functional and clinical studies are recommended to clarify its significance in disease etiology and therapeutic response. This variant should be interpreted in the context of clinical and molecular findings, and consideration given to potential eligibility for targeted therapies exploiting synthetic lethality in ARID1A-deficient tumors. No direct references specific to p.R1989* exist; the summary is based on broader evidence of ARID1A truncating mutations and their role in cancer biology[3][5][6][8]. | [1] <https://pubmed.ncbi.nlm.nih.gov/27030501/>  [2] <https://www.semanticscholar.org/paper/e327637e8ea6b8a976ade0920318798011d485f4>  [3] <https://www.ncbi.nlm.nih.gov/pmc/articles/PMC11223007/>  [4] <https://www.semanticscholar.org/paper/39f6fc085916db1515f5fbd2e82eeaf3acdcbaf3>  [5] <https://www.ncbi.nlm.nih.gov/pmc/articles/PMC3396657/>  [6] <https://www.ncbi.nlm.nih.gov/pmc/articles/PMC7057434/>  [7] <https://pubmed.ncbi.nlm.nih.gov/33222288/>  [8] <https://www.ncbi.nlm.nih.gov/pmc/articles/PMC9987588/> |
|  | Evaluation  **Summarization Accuracy & Completeness:**  **Hallucination Rate:**  **Readability & Usability:**  **Literature Coverage & Relevance:**  **Time Efficiency:** | |
