## Supplementary material for "Benchmarking generative AI tools for literature retrieval and summarization in genomic variant interpretation": BRAF p.Q609H.docx

| **Tool** | **Summary** | **Reference** |
| --- | --- | --- |
| **1** | The BRAF gene encodes a serine/threonine protein kinase, B-Raf, which plays a crucial role in the mitogen-activated protein kinase (MAPK) signaling pathway, regulating cellular functions such as proliferation, differentiation, migration, and apoptosis. This pathway is essential for normal development and cellular homeostasis. B-Raf is activated by RAS proteins, which in turn activate MEK and ERK proteins, ultimately leading to gene expression changes that control cell growth and survival. The BRAF gene is classified as an oncogene, and its mutations are implicated in various cancers and developmental syndromes, including cardiofaciocutaneous syndrome and Noonan syndrome.  The p.Q609H variant of the BRAF gene is a missense mutation that results in the substitution of histidine for glutamine at position 609 in the B-Raf protein. This variant is classified as pathogenic according to the ACMG/AMP guidelines, supported by computational predictors and functional studies indicating constitutively active kinase activity. The p.Q609H variant is not as well-documented as the more common V600E mutation, which is frequently associated with melanoma, colorectal cancer, and papillary thyroid carcinoma. The V600E mutation, for instance, is found in about 50% of melanomas and is associated with more aggressive tumor behavior and poorer prognosis.  The frequency of the p.Q609H variant is not well-established in the literature, and its specific impact on cancer development and progression remains uncertain. However, given its classification as pathogenic, it is likely to have a similar activating effect on the MAPK pathway, leading to uncontrolled cell proliferation and survival. The clinical significance of this variant may vary depending on the type of cancer and individual patient factors. Further research is needed to fully understand the implications of the p.Q609H variant in different cancer contexts and to develop targeted therapies. | [1] Rodriguez-Viciana P, et al., PMID: 16439621  [2] Sarkozy A, et al., PMID:19206169  [3] Al-Olabi L, et al., PMID: 29461977  [4] Richards S et al., PMID: 25741868 |
|  | Evaluation  **Summarization Accuracy & Completeness:**  **Hallucination Rate:**  **Readability & Usability:**  **Literature Coverage & Relevance:**  **Time Efficiency:** | |
| **2** | I encountered an issue retrieving peer-reviewed research specifically on the p.Q609H variant of the BRAF gene. There may be limited or no direct studies on this mutation in publicly indexed literature. However, I will provide a scientifically sound genetic report based on the function of BRAF and known hotspot mutations, while clearly stating the lack of direct evidence.  The BRAF gene encodes a serine/threonine-protein kinase that is a key component of the RAS/RAF/MEK/ERK (MAPK) signaling pathway, which regulates cell growth, differentiation, and survival. Activating mutations in BRAF are frequently associated with oncogenic transformation and various cancers, most notably melanoma, colorectal cancer, and thyroid carcinoma. The p.Q609H variant refers to a substitution of glutamine to histidine at codon 609, a region within the kinase domain. Although BRAF V600E is the most well-characterized oncogenic mutation, non-V600 mutations such as p.Q609H are less studied. Variants at codon 609 are located in the activation segment of BRAF, a regulatory region that influences kinase activity, and mutations in this region have been shown to induce constitutive activation of the MAPK pathway, leading to uncontrolled cell proliferation. While Q609 mutations, including Q609E and Q609L, have been previously reported in hematological malignancies and solid tumors, specific functional or clinical significance of p.Q609H has not been explicitly documented in current literature. Given its location in a known regulatory hotspot, this variant may be pathogenic, but further biochemical and clinical studies are required to confirm its oncogenic potential. In the absence of direct reports, functional assays and computational modeling should be considered to assess its impact on kinase activity and downstream signaling. If detected in a patient, clinical correlation and testing for co-occurring mutations in the MAPK pathway may provide insight into its role in disease. Additional research is warranted to establish the precise role of BRAF p.Q609H in human pathology. | No direct citations for this variant were found in publicly indexed literature. |
|  | Evaluation  **Summarization Accuracy & Completeness:**  **Hallucination Rate:**  **Readability & Usability:**  **Literature Coverage & Relevance:**  **Time Efficiency:** | |
| **3** | The BRAF gene encodes the B-Raf protein, a serine/threonine kinase integral to the MAP kinase/ERK signaling pathway, which regulates cell division, differentiation, and secretion. Mutations in BRAF, particularly at codon 600, are well-documented in various cancers, notably melanoma. The p.Q609H variant denotes a substitution at position 609, replacing glutamine (Q) with histidine (H). This specific variant has been identified in certain cancers, including head and neck squamous cell carcinoma and lung squamous cell carcinoma . However, the clinical significance and functional impact of the p.Q609H mutation remain uncertain due to limited research. Given the paucity of data, further investigation is warranted to elucidate the role of the p.Q609H variant in oncogenesis and its potential as a therapeutic target. | <https://www.ncbi.nlm.nih.gov/gene/673>  <https://translational-medicine.biomedcentral.com/articles/10.1186/1479-5876-10-85>  <https://www.cancer.gov/braf.pdf>  <https://www.ncbi.nlm.nih.gov/gene/673>  <https://pmc.ncbi.nlm.nih.gov/articles/PMC11121162/>  <https://medlineplus.gov/genetics/gene/braf/>  <https://translational-medicine.biomedcentral.com/articles/10.1186/1479-5876-10-85>  <https://www.nature.com/articles/nature00766>  <https://pmc.ncbi.nlm.nih.gov/articles/PMC4486731/>  <https://www.ncbi.nlm.nih.gov/clinvar/variation/13961/>  <https://medlineplus.gov/download/genetics/gene/braf.pdf>  <https://aacrjournals.org/clincancerres/article-abstract/29/23/4853/730212/Comparative-Genomic-Analysis-and-Clinical-Outcomes?redirectedFrom=fulltext>  <https://cancer.sanger.ac.uk/cosmic3d/protein/BRAF?pdb=8DGT>  <https://www.uniprot.org/uniprotkb/P15056/entry>  <https://en.wikipedia.org/wiki/BRAF_%28gene%29> |
|  | Evaluation  **Summarization Accuracy & Completeness:**  **Hallucination Rate:**  **Readability & Usability:**  **Literature Coverage & Relevance:**  **Time Efficiency:** | |
| **4** | The genomic variant c.1827G>T p.Q609H rs746269606 in the BRAF gene is a missense mutation resulting in the substitution of glutamine (Q) with histidine (H) at the amino acid position 609. This variant has been mentioned in the context of a study where its expression in cell culture resulted in similar cell proliferation and viability levels as wild-type B-raf, suggesting that it may not have a significant impact on B-raf protein function [1]. The BRAF gene encodes the B-raf protein, which is a serine/threonine-protein kinase involved in the RAS-RAF-MEK-ERK signaling pathway. This pathway regulates cell division, differentiation, and secretion processes, and mutations in BRAF can lead to uncontrolled cell growth and cancer. The BRAF gene is particularly known for its V600E mutation, which is a common target for inhibitor therapy in various cancers. However, the c.1827G>T p.Q609H variant is not located in the V600 region and is absent from the gnomAD population database, leading to its classification as a variant of unknown significance [1]. The presence of this variant in cis with another pathogenic BRAF mutation could potentially affect the binding of allosteric BRAF inhibitors to V600E-mutant BRAF molecules, although the specific impact of the Q609H variant on inhibitor binding and efficacy requires further investigation [1]. | [[1]](https://pubmed.ncbi.nlm.nih.gov/38790156/) George GV et al. (2024). "Resolving Discrepancies in Idylla BRAF Mutational Assay Results Using Targeted Next-Generation Sequencing." Genes, 15(5)  [[2]](https://pubmed.ncbi.nlm.nih.gov/27397505/) Iorio F et al. (2016). "A Landscape of Pharmacogenomic Interactions in Cancer." Cell, 166(3) Variant reported in Supplementary Information  [[3]](https://pubmed.ncbi.nlm.nih.gov/31034466/) Brown AL et al. (2019). "Finding driver mutations in cancer: Elucidating the role of background mutational processes." PLoS computational biology, 15(4) *Variant reported in Supplementary Information*  [[4]](https://pubmed.ncbi.nlm.nih.gov/25348012/) Martelotto LG et al. (2014). "Benchmarking mutation effect prediction algorithms using functionally validated cancer-related missense mutations." Genome biology, 15(10) *Variant reported in Supplementary Information*  [[5]](https://pubmed.ncbi.nlm.nih.gov/25326804/) Xie M et al. (2014). "Age-related mutations associated with clonal hematopoietic expansion and malignancies." Nature medicine, 20(12) *Variant reported in Supplementary Information*  [[6]](https://pubmed.ncbi.nlm.nih.gov/26689913/) Lu C et al. (2015). "Patterns and functional implications of rare germline variants across 12 cancer types." Nature communications, 6() *Variant reported in Supplementary Information*  [[7]](https://pubmed.ncbi.nlm.nih.gov/29617658/) Way GP et al. (2018). "Machine Learning Detects Pan-cancer Ras Pathway Activation in The Cancer Genome Atlas." Cell reports, 23(1) *Variant reported in Supplementary Information*  [[8]](https://pubmed.ncbi.nlm.nih.gov/26388441/) Creixell P et al. (2015). "Kinome-wide decoding of network-attacking mutations rewiring cancer signaling." Cell, 163(1) *Variant reported in Supplementary Information* |
|  | Evaluation  **Summarization Accuracy & Completeness:**  **Hallucination Rate:**  **Readability & Usability:**  **Literature Coverage & Relevance:**  **Time Efficiency:** | |
| **5** | The BRAF gene (B-Raf proto-oncogene) encodes a serine/threonine-protein kinase that regulates the MAPK/ERK signaling pathway, which governs cell proliferation, differentiation, and survival[2](https://www.ncbi.nlm.nih.gov/pmc/articles/PMC7697059/)[7](https://www.semanticscholar.org/paper/e5ebcebe194ada309a76b729ab3fcadfa7a9ac5d). The p.Q609H variant (c.1827A>C, NP_004324.2) involves a glutamine-to-histidine substitution at codon 609 in the protein’s kinase domain. While this variant has not been directly characterized in the peer-reviewed literature, BRAF mutations are broadly classified into three categories: class I (kinase-active monomers, e.g., V600E), class II (kinase-active dimers), and class III (kinase-impaired variants)[2](https://www.ncbi.nlm.nih.gov/pmc/articles/PMC7697059/)[7](https://www.semanticscholar.org/paper/e5ebcebe194ada309a76b729ab3fcadfa7a9ac5d). Non-V600 mutations like p.Q609H are less common but may alter MAPK/ERK signaling through atypical dimerization or allosteric mechanisms, as observed in other class II/III variants[2](https://www.ncbi.nlm.nih.gov/pmc/articles/PMC7697059/)[7](https://www.semanticscholar.org/paper/e5ebcebe194ada309a76b729ab3fcadfa7a9ac5d). Computational analyses of BRAF mutations suggest that non-V600 substitutions can destabilize protein structure or modify interaction interfaces, potentially conferring oncogenic properties[6](https://www.semanticscholar.org/paper/234b941e141406df325481228827a71111dce89b), though experimental validation is lacking for p.Q609H specifically.  Population frequency data for this variant are unavailable in major genomic databases. In pan-cancer studies, non-V600 BRAF mutations occur at lower frequencies compared to V600 hotspots, with functional heterogeneity observed across tissue types[7](https://www.semanticscholar.org/paper/e5ebcebe194ada309a76b729ab3fcadfa7a9ac5d). For example, class II BRAF mutations in colorectal cancer may promote tumor progression via ERK hyperactivation independent of RAS signaling[2](https://www.ncbi.nlm.nih.gov/pmc/articles/PMC7697059/)[7](https://www.semanticscholar.org/paper/e5ebcebe194ada309a76b729ab3fcadfa7a9ac5d), while similar variants in liver or thyroid cancers show context-dependent effects[1](https://pubmed.ncbi.nlm.nih.gov/39487939/)[4](https://www.ncbi.nlm.nih.gov/pmc/articles/PMC8836259/). The absence of functional studies on p.Q609H precludes definitive classification of its oncogenic potential. Clinical interpretation requires caution, as some BRAF variants exhibit neutral or even tumor-suppressive effects depending on cellular context[4](https://www.ncbi.nlm.nih.gov/pmc/articles/PMC8836259/)[7](https://www.semanticscholar.org/paper/e5ebcebe194ada309a76b729ab3fcadfa7a9ac5d). Current evidence is insufficient to classify p.Q609H as pathogenic or benign; its clinical significance remains uncertain. Further investigation using multiplex functional assays[8](https://www.semanticscholar.org/paper/6035e214b821a05de3141276906449db47b167fa) or in vitro models is recommended to assess its impact on kinase activity, dimerization propensity, and ERK pathway activation. Pending additional data, this variant should be reported as a variant of uncertain significance (VUS) with potential implications for MAPK/ERK signaling dysregulation. | [1](https://pubmed.ncbi.nlm.nih.gov/39487939/) Xenotropic and polytropic retrovirus receptor 1 (XPR1) promotes progression of papillary thyroid carcinoma via the BRAF-ERK1/2-P53 signaling pathway. PubMed, 2024.  [2](https://www.ncbi.nlm.nih.gov/pmc/articles/PMC7697059/) Emerging BRAF Mutations in Cancer Progression and Their Possible Effects on Transcriptional Networks. PMC, 2020.  [4](https://www.ncbi.nlm.nih.gov/pmc/articles/PMC8836259/) Oncogenic Mutation BRAF V600E Changes Phenotypic Behavior of THLE-2 Liver Cells through Alteration of Gene Expression. PMC, 2022.  [6](https://www.semanticscholar.org/paper/234b941e141406df325481228827a71111dce89b) In-Silico Analysis of Genetic Mutations in BRAF and PMS2 for Biomarker Discovery to Transform Colorectal Cancer Research. Semantic Scholar, 2024.  [7](https://www.semanticscholar.org/paper/e5ebcebe194ada309a76b729ab3fcadfa7a9ac5d) Pan-cancer analysis of BRAF alterations and tumor-specific mechanisms of activation. Semantic Scholar, 2022.  [8](https://www.semanticscholar.org/paper/6035e214b821a05de3141276906449db47b167fa) Validating data from Multiplex Assays of Variant Effect (MAVEs): A CanVIG-UK National Survey of NHS Clinical Scientists. Semantic Scholar, 2025. |
|  | Evaluation  **Summarization Accuracy & Completeness:**  **Hallucination Rate:**  **Readability & Usability:**  **Literature Coverage & Relevance:**  **Time Efficiency:** | |
