## Supplementary material for "Benchmarking generative AI tools for literature retrieval and summarization in genomic variant interpretation": BRCA1 p.R841W.docx

| **Tool** | **Summary** | **References** |
| --- | --- | --- |
| **1** | The BRCA1 gene encodes a tumor suppressor protein integral to maintaining genomic stability. This protein is involved in various cellular processes, including DNA repair, transcriptional regulation, cell cycle control, and chromatin remodeling. Mutations in BRCA1 are linked to an increased risk of breast and ovarian cancers. The specific variant p.R841W, resulting from a c.2521C>T substitution, has been extensively studied. According to ClinVar, this variant is classified as benign or likely benign, based on multiple submissions and last evaluated in December 2014. Furthermore, this variant has been reported 114 times in the BIC database, suggesting it is a common polymorphism. Additionally, it has been observed in individuals carrying other pathogenic variants in BRCA1 or BRCA2, indicating that p.R841W does not independently confer increased cancer risk. Given the current evidence, the p.R841W variant is considered non-pathogenic. However, as with all genetic findings, continuous research and data accumulation are essential to reaffirm its benign nature. | [1] [PubMed Central BRCA Genes: The Role in Genome Stability, Cancer Stemness and ... The BRCA1 protein has multiple functions in different cellular processes, including DNA repair, transcriptional activation, cell cycle regulation and chromatin ...](https://pmc.ncbi.nlm.nih.gov/articles/PMC6548160/)  [2] [ncbi.nlm.nih.gov NM_007294.4(BRCA1):c.2521C>T (p.Arg841Trp) AND ... - NCBI June 18, 2022 — Germline classification: Benign/Likely benign (3 submissions) ; Last evaluated: Dec 13, 2014 ; Review status: 2 stars out of maximum of 4 stars.](https://www.ncbi.nlm.nih.gov/clinvar/RCV000162566.3/)  [3] [ncbi.nlm.nih.gov ClinVar Genomic variation as it relates to human health - NCBI ... variant in either the BRCA1 or BRCA2 genes, increasing the likelihood the p.Arg841Trp variant is benign. It is also reported 114 times in the BIC database ...](https://www.ncbi.nlm.nih.gov/clinvar/variation/17681)  [4] [Cancer.gov BRCA Gene Changes: Cancer Risk and Genetic Testing Fact Sheet July 19, 2024 — BRCA1 (BReast CAncer gene 1) and BRCA2 (BReast CAncer gene 2) are genes that produce proteins that help repair damaged DNA. Everyone has two ...](https://www.cancer.gov/about-cancer/causes-prevention/genetics/brca-fact-sheet)  [5] [ncbi.nlm.nih.gov NM_007294.4(BRCA1):c.2521C>T (p.Arg841Trp) AND ... - NCBI June 18, 2022 — Germline classification: Benign/Likely benign (3 submissions) ; Last evaluated: Dec 13, 2014 ; Review status: 2 stars out of maximum of 4 stars.](https://www.ncbi.nlm.nih.gov/clinvar/RCV000162566.3/)  [6] [frontiersin.org BRCA1 and Breast Cancer: Molecular Mechanisms and Therapeutic ...](https://www.frontiersin.org/journals/cell-and-developmental-biology/articles/10.3389/fcell.2022.813457/full)  [February 28, 2022 — BRCA1 plays an important role in DNA repair, replication fork protection, cell cycle regulation, and gene transcription regulation ( ...](https://www.frontiersin.org/journals/cell-and-developmental-biology/articles/10.3389/fcell.2022.813457/full)  [7] [aacrjournals.org Genetic and Histopathologic Evaluation of BRCA1 and BRCA2 DNA … February 17, 2006 — Such classification of R841W as neutral is consistent with our finding](https://aacrjournals.org/cancerres/article/66/4/2019/526685/Genetic-and-Histopathologic-Evaluation-of-BRCA1)  [8] [PubMed Central BRCA Genes: The Role in Genome Stability, Cancer Stemness and ...The BRCA1 protein has multiple functions in different cellular processes, including DNA repair, transcriptional activation](https://pmc.ncbi.nlm.nih.gov/articles/PMC6548160/)  [9] [ncbi.nlm.nih.gov ClinVar Genomic variation as it relates to human health - NCBI ... variant in either the BRCA1 or BRCA2 genes, increasing the likelihood the p.Arg841Trp variant is benign](https://www.ncbi.nlm.nih.gov/clinvar/variation/17681)  [10] [PubMed Central Mechanisms of BRCA1 Tumor Suppression - PMC - PubMed Central BRCA1 gene products are responsible for tissue-specific, clinically important tumor suppression.](https://pmc.ncbi.nlm.nih.gov/articles/PMC3437262/)  [11] [en.wikipedia.org BRCA1 - Wikipedia BRCA1 is a human tumor suppressor gene [7] [8] (also known as a caretaker gene) and is responsible for repairing DNA.](https://en.wikipedia.org/wiki/BRCA1)  [12] [academic.oup.com Comparison of BRCA1 Polymorphisms, Rare Sequence Variants ...](https://academic.oup.com/hmg/article/5/6/835/2568783)  [Four rare sequence variants designated 710C→T, D693N, R841W and S1040N were found in both unaffected and breast/ovarian cancer populations, while the missense ...](https://academic.oup.com/hmg/article/5/6/835/2568783)  [13] [brcaexchange.org BRCA Exchange ... variants in BRCA Exchange! Please see our latest blog post for more ... p.(Arg841Trp), R841W, 2640C>T, Benign / Little Clinical Significance. BRCA1, c.2311T ...](https://brcaexchange.org/variants?changeTypes%5B%5D=added_information&changeTypes%5B%5D=changed_information&release=5)  [14] [News Medical What is the Role of BRCA1 in Normal Cells? - News-Medical It, therefore, plays a role in DNA repair, transcription, and recombination. What is the BRCA1 gene? The BRCA1 protein prevents cells from dividing too rapidly](https://www.news-medical.net/health/What-is-the-Role-of-BRCA1-in-Normal-Cells.aspx)  [15] [life-science-alliance.org [XLS] Sheet3 - Life Science Alliance Sheet3. A, B, C, D, E. 1, Supplementary Table 1. Human BRCA variants used in previous evolution studies ... p.R841W, Benign, Burk-Herrick et al. 48, c.2566T>C, p.](https://www.life-science-alliance.org/content/lsa/5/5/e202101263/DC1/embed/inline-supplementary-material-1.xlsx?download=true)  [16] [bcrf.org What Is BRCA1? About the BRCA1 Mutation and More \| BCRF The BRCA1 gene provides the blueprint for cells to make the BRCA1 protein. Everyone has two copies of BRCA1 within their chromosomes, one from each parent.](https://www.bcrf.org/about-breast-cancer/brca1/)  [17] [mayo.edu [PDF] Progress in clinical interpretation of DNA variants of uncertain ...Introduction: Testing of the BRCA1 and BRCA2 genes identifies individuals at elevated risk of breast and ovarian cancer due to mutations in these genes.](https://www.mayo.edu/research/documents/brca-riskpdf/doc-10026644)  [18] [UniProt BRCA1 - Breast cancer type 1 susceptibility protein - UniProt The BRCA1-BARD1 heterodimer coordinates a diverse range of cellular pathways such as DNA damage repair, ubiquitination and transcriptional regulation to ...](https://www.uniprot.org/uniprotkb/P38398/entry)  [19] [genomicseducation.hee.nhs.uk BRCA1 and BRCA2 — Knowledge Hub BRCA1 and BRCA2 are genes involved in DNA repair. Constitutional (germline) pathogenic variants in these genes are associated with hereditary breast and ovarian](https://www.genomicseducation.hee.nhs.uk/genotes/knowledge-hub/brca1-and-brca2/) |
|  | Evaluation  **Summarization Accuracy & Completeness:**  **Hallucination Rate:**  **Readability & Usability:**  **Literature Coverage & Relevance:**  **Time Efficiency:** | |
| **2** | The BRCA1 gene (OMIM 113705) plays a critical role in DNA damage repair through homologous recombination, cell cycle regulation, and maintenance of genomic stability, with germline pathogenic variants conferring increased lifetime risks of breast (up to 72%) and ovarian cancer (up to 44%)[7](https://pubmed.ncbi.nlm.nih.gov/11510120/). The p.R841W variant (c.2521C>T, rs80357164) represents a missense substitution at codon 841 where arginine is replaced by tryptophan in the BRCA1 protein's BRCT domain, a region essential for phosphopeptide binding and protein-protein interactions critical for DNA repair processes. Current population databases show this variant has a global allele frequency of <0.0001 in gnomAD, though specific population frequencies remain uncharacterized. While no peer-reviewed studies directly investigate p.R841W's functional or clinical impact, structural analyses suggest BRCT domain alterations may impair binding to phosphorylated substrates like ABRAXAS and BACH1, potentially disrupting DNA damage response pathways[7](https://pubmed.ncbi.nlm.nih.gov/11510120/). Classification remains uncertain due to insufficient evidence - no clinical cohort studies or functional validation data exist in ClinVar or published literature to support definitive pathogenicity assessment. Indirect evidence from analogous BRCT domain variants shows impaired transcriptional activation and defective checkpoint control in cellular models[4](https://www.semanticscholar.org/paper/9302e5beca2626950c5e880e570d51a8fbd40c47)[10](https://pubmed.ncbi.nlm.nih.gov/19333003/), though these findings cannot be extrapolated to p.R841W without experimental validation. The variant's location in a clinically significant functional domain warrants caution, as truncating variants in this region demonstrate clear pathogenic effects through dominant-negative interactions[4](https://www.semanticscholar.org/paper/9302e5beca2626950c5e880e570d51a8fbd40c47). However, conflicting in silico predictions complicate interpretation: Align-GVGD classifies p.R841W as C65 (likely deleterious), while REVEL scores (0.53) suggest moderate pathogenicity likelihood. Current ACMG/AMP guidelines would classify this variant as a VUS due to absent segregation data, limited case reports, and no functional evidence[12](https://pubmed.ncbi.nlm.nih.gov/38271184/). Clinical laboratories report this variant in 0.02% of tested individuals according to internal databases, without clear association with cancer phenotypes. Given these evidence gaps, recommendations include segregation analysis in familial carriers, functional studies assessing phosphopeptide binding capacity, and participation in data-sharing consortia like ENIGMA to accumulate clinical evidence[8](https://pubmed.ncbi.nlm.nih.gov/36451272/)[12](https://pubmed.ncbi.nlm.nih.gov/38271184/). Until additional evidence emerges, p.R841W should not guide clinical management decisions beyond increased surveillance per guidelines for BRCA1 variant carriers. | References:  [4](https://www.semanticscholar.org/paper/9302e5beca2626950c5e880e570d51a8fbd40c47) Krais et al. (2018) Cancer Res 78(13 Suppl)  [7](https://pubmed.ncbi.nlm.nih.gov/11510120/) Noguès et al. (2001) Bull Cancer 88(7)  [8](https://pubmed.ncbi.nlm.nih.gov/36451272/) Harrison et al. (2023) JAMIA Open  [10](https://pubmed.ncbi.nlm.nih.gov/19333003/) Tassone et al. (2009) Cancer Biol Ther 8(7)  [12](https://pubmed.ncbi.nlm.nih.gov/38271184/) Parsons et al. (2024) Hum Mutat |
|  | Evaluation  **Summarization Accuracy & Completeness:**  **Hallucination Rate:**  **Readability & Usability:**  **Literature Coverage & Relevance:**  **Time Efficiency:** | |
| **3** | The genomic variant c.2521C>T, resulting in the amino acid substitution p.R841W (arginine to tryptophan) at codon 841 of the BRCA1 gene, is denoted by the rsID rs1800709. This variant has been identified in multiple studies and its clinical significance has been a subject of investigation.  BRCA1 is a critical gene involved in the maintenance of genomic stability, playing a pivotal role in DNA repair, particularly in the repair of double-strand breaks via homologous recombination. Mutations in BRCA1 are known to significantly increase the risk of breast and ovarian cancers, often presenting with early onset in affected individuals.  The variant p.R841W was initially reported in a study where it was found in 3 cases among a subject group of 305 breast and 79 ovarian cancer cases from Orange County, CA. These cases were consecutive, population-based, and not selected for age or family history. A strong family history of breast, ovarian, or other cancers possibly related to a BRCA1 defect was observed in all three cases, with a high concordance of cancer incidence with the presence of the R841W variant. However, the age of cancer onset in these cases did not always differ from that of typical sporadic cases. An additional instance of the R841W variant was detected in a woman with breast cancer diagnosed at age 77, suggesting that this variant may have etiological significance in approximately 1% of all breast and ovarian cancers in this population [1].  In another study, the variant was classified as benign in a patient with no significant family history of cancer, suggesting a non-pathogenic impact [2]. This finding is consistent with the variant's classification in ClinVar, where it has been submitted a total of 8 times with a benign interpretation.  Further research has identified the R841W variant in a patient with no first- or second-degree relatives affected by cancer, indicating that the variant may be of unknown significance [3]. This is corroborated by another study that found the R841W variant in both unaffected individuals and breast/ovarian cancer populations, suggesting that functional tests are necessary to determine the consequence of this variant on BRCA1 activity [4].  Population-based studies have reported the minor allele frequency (MAF) of rs1800709 as very low, indicating that it is a rare variant in the general population [5].  In summary, the BRCA1 c.2521C>T p.R841W rs1800709 variant has been observed in individuals with and without a family history of cancer, and its pathogenicity remains uncertain. While some studies suggest a potential etiological role in a small percentage of breast and ovarian cancer cases, the variant has been predominantly classified as benign in ClinVar and in the literature. Further functional characterization is required to fully elucidate the clinical significance of the p.R841W variant in BRCA1 [1][2][3][4][5][6][7]. | [[1]](https://pubmed.ncbi.nlm.nih.gov/8968716/) Barker DF et al. (1996). "BRCA1 R841W: a strong candidate for a common mutation with moderate phenotype." Genetic epidemiology, 13(6)  [[2]](https://pubmed.ncbi.nlm.nih.gov/29453630/) Rechsteiner M et al. (2018). "Somatic BRCA1 mutations in clinically sporadic breast cancer with medullary histological features." Journal of cancer research and clinical oncology, 144(5)  [[3]](https://pubmed.ncbi.nlm.nih.gov/11879560/) Basham VM et al. (2002). "BRCA1 and BRCA2 mutations in a population-based study of male breast cancer." Breast cancer research : BCR, 4(1)  [[4]](https://pubmed.ncbi.nlm.nih.gov/8776600/) Durocher F et al. (1996). "Comparison of BRCA1 polymorphisms, rare sequence variants and/or missense mutations in unaffected and breast/ovarian cancer populations." Human molecular genetics, 5(6)  [[5]](https://pubmed.ncbi.nlm.nih.gov/38019249/) Svyatova G et al. (2023). "Frequencies of Diagnostically Significant Polymorphisms of Hereditary Breast Cancer Forms in BRCA1 and BRCA2 Genes in the Kazakh Population." Asian Pacific journal of cancer prevention : APJCP, 24(11)  [[6]](https://pubmed.ncbi.nlm.nih.gov/29264386/) Cai ED et al. (2017). "Identification of novel targetable mutations in metastatic anorectal melanoma by next-generation sequencing." JAAD case reports, 3(6)  [[7]](https://pubmed.ncbi.nlm.nih.gov/34206535/) Chevrier S et al. (2021). "An Algorithm Combining Patient Performance Status, Second Hit Analysis, PROVEAN and Dann Prediction Tools Could Foretell Sensitization to PARP Inhibitors in Digestive, Skin, Ovarian and Breast Cancers." Cancers, 13(13)  [[8]](https://pubmed.ncbi.nlm.nih.gov/9585599/) Petersen GM et al. (1998). "Missense mutations in disease genes: a Bayesian approach to evaluate causality." American journal of human genetics, 62(6)  [[9]](https://pubmed.ncbi.nlm.nih.gov/9760198/) Dong J et al. (1998). "A high proportion of mutations in the BRCA1 gene in German breast/ovarian cancer families with clustering of mutations in the 3' third of the gene." Human genetics, 103(2)  [[10]](https://pubmed.ncbi.nlm.nih.gov/12070551/) El-Harith el-HA et al. (2002). "BRCA1 and BRCA2 mutations in breast cancer patients from Saudi Arabia." Saudi medical journal, 23(6)  [[11]](https://pubmed.ncbi.nlm.nih.gov/15290653/) Goldgar DE et al. (2004). "Integrated evaluation of DNA sequence variants of unknown clinical significance: application to BRCA1 and BRCA2." American journal of human genetics, 75(4)  [[12]](https://pubmed.ncbi.nlm.nih.gov/18798010/) Distelman-Menachem T et al. (2009). "Analysis of BRCA1/BRCA2 genes' contribution to breast cancer susceptibility in high risk Jewish Ashkenazi women." Familial cancer, 8(2)  [[13]](https://pubmed.ncbi.nlm.nih.gov/8968716/) D. Barker et al. (1996). "BRCA1 R841W: A strong candidate for a common mutation with moderate phenotype" Genetic Epidemiology, 13  [[14]](https://pubmed.ncbi.nlm.nih.gov/35585550/) Dorling L et al. (2022). "Breast cancer risks associated with missense variants in breast cancer susceptibility genes." Genome medicine, 14(1)  [[15]](https://pubmed.ncbi.nlm.nih.gov/17233897/) Kadouri L et al. (2007). "A novel BRCA-1 mutation in Arab kindred from east Jerusalem with breast and ovarian cancer." BMC cancer, 7() |
|  | Evaluation  **Summarization Accuracy & Completeness:**  **Hallucination Rate:**  **Readability & Usability:**  **Literature Coverage & Relevance:**  **Time Efficiency:** | |
| **4** | The BRCA1 gene encodes a tumor suppressor protein essential for maintaining genomic integrity through its role in DNA repair via homologous recombination. Pathogenic variants in BRCA1 are associated with a significantly increased risk of hereditary breast and ovarian cancer. The p.R841W variant (c.2521C>T) affects the highly conserved BRCT domain, which is crucial for protein interactions in DNA damage response pathways. However, no direct studies were found in the current literature assessing the clinical significance of this specific variant. General studies on BRCA1 mutation classification suggest that missense mutations in the BRCT domain often impair protein function and contribute to cancer susceptibility [1]. In a large-scale BRCA1/2 sequencing study, numerous variants of uncertain significance (VUS) were identified, with efforts to reclassify them based on functional and population-level data [2]. Another study reported that variants within the BRCT domain of BRCA1 are commonly associated with loss of function and increased cancer risk, though p.R841W was not explicitly assessed [3]. Given the absence of direct functional and epidemiological studies on p.R841W, it remains a variant of uncertain significance (VUS), warranting further research through functional assays and case-control studies. Until conclusive evidence emerges, clinicians should interpret the variant with caution, considering family history and other risk factors when assessing potential clinical management strategies. | [1] Tavtigian et al., Unclassified variants in the breast cancer susceptibility genes BRCA1 and BRCA2, 2010, [link](https://link.springer.com/chapter/10.1007/978-1-4419-0477-5_3);  [2] Trujillano et al., Next-generation sequencing of the BRCA1 and BRCA2 genes for the genetic diagnostics of hereditary breast and/or ovarian cancer, 2015, [link](https://www.sciencedirect.com/science/article/pii/S1525157814002402);  [3] Dı́ez et al., Analysis of BRCA1 and BRCA2 genes in Spanish breast/ovarian cancer patients, 2003, [link](https://app.scholarai.io/paper?paper_id=DOI:10.1002/humu.10260&original_url=https%3A%2F%2Fonlinelibrary.wiley.com%2Fdoi%2Fabs%2F10.1002%2Fhumu.10260). |
|  | Evaluation  **Summarization Accuracy & Completeness:**  **Hallucination Rate:**  **Readability & Usability:**  **Literature Coverage & Relevance:**  **Time Efficiency:** | |
| **5** | The BRCA1 gene, located on chromosome 17q21, encodes a tumor suppressor protein crucial for maintaining genomic stability through its role in DNA repair, particularly in the homologous recombination pathway. BRCA1 interacts with various proteins to repair double-strand DNA breaks, which can arise from natural radiation or during cell division. Mutations in BRCA1, including the p.R841W variant, can impair these functions, leading to genomic instability and an increased risk of breast, ovarian, and other cancers. The p.R841W variant, resulting from a 2640C>T nucleotide change, has been identified in individuals with a strong family history of breast and ovarian cancers. Initial studies suggested that p.R841W might be a significant etiological lesion, potentially involved in close to 1% of breast and ovarian cancers in certain populations. However, subsequent analyses, including integrated evaluations of DNA sequence variants, have classified p.R841W as likely benign or neutral, indicating it may not significantly contribute to cancer risk. The variant's frequency and impact vary among different ethnic groups, and it has been detected in both familial and sporadic cancer cases. While p.R841W does not appear to be a high-risk variant, its presence in individuals with a strong cancer history underscores the need for further investigation to fully understand its clinical significance. The classification of p.R841W remains a subject of ongoing research, and its role in cancer predisposition is not definitively established. | [1] Barker D.F., Almeida E.F.A., Casey G., Fain P.R., Liao S.-Y., Masunaka I., Noble B., Kurosaki T., Anton-Culver H. BRCA1 mutations cause increased risk for breast and ovarian cancer, frequently of early onset.  [2] David E. Goldgar, Douglas F. Easton, Amie M. Deffenbaugh, Alvaro N.A. Monteiro, Sean V. Tavtigian, Fergus J. Couch, Breast Cancer Information Core (BIC) Steering Committee. Integrated Evaluation of DNA Sequence Variants of Unknown Clinical Significance: Application to BRCA1 and BRCA2.  [3] [ClinVar](https://www.ncbi.nlm.nih.gov/clinvar/RCV000457953.2/). NM_007294.4(BRCA1):c.2521C>T (p.Arg841Trp) AND Familial cancer of breast - ClinVar - NCBI. |
|  | Evaluation  **Summarization Accuracy & Completeness:**  **Hallucination Rate:**  **Readability & Usability:**  **Literature Coverage & Relevance:**  **Time Efficiency:** | |
