## Supplementary material for "Benchmarking generative AI tools for literature retrieval and summarization in genomic variant interpretation": BRCA2 p.S1882_.docx

| **Tool** | **Summary** | **References** |
| --- | --- | --- |
| **1** | The BRCA2 gene encodes a tumor suppressor protein essential for maintaining genomic stability through its role in homologous recombination-mediated DNA double-strand break repair. The c.5645C>A (p.Ser1882*) variant, also referred to as 5873C>A, introduces a premature stop codon in exon 11, leading to a truncated or absent BRCA2 protein due to nonsense-mediated mRNA decay. This loss-of-function mechanism is a well-established cause of hereditary breast and ovarian cancer (HBOC) syndrome. The p.Ser1882* variant has been identified in multiple individuals with breast, ovarian, and prostate cancers, including cases of triple-negative breast cancer and metastatic prostate cancer . It has also been reported in two siblings with Wilms tumor who were compound heterozygotes for BRCA2 mutations . This variant is extremely rare in the general population, with an allele frequency of approximately 0.002% in the Genome Aggregation Database (gnomAD) . Based on the predicted impact on protein function, absence in the general population, and presence in multiple affected individuals, the p.Ser1882* variant is classified as pathogenic according to the American College of Medical Genetics and Genomics (ACMG) guidelines . | [https://www.mdpi.com/2073-4425/12/8/1229](https://www.mdpi.com/2073-4425/12/8/1229?utm_source=chatgpt.com)  [https://www.ncbi.nlm.nih.gov/clinvar/variation/37984](https://www.ncbi.nlm.nih.gov/clinvar/variation/37984/?utm_source=chatgpt.com)  [https://www.ncbi.nlm.nih.gov/clinvar/RCV000167830](https://www.ncbi.nlm.nih.gov/clinvar/RCV000167830/?utm_source=chatgpt.com)  <https://www.medrxiv.org/content/10.1101/2024.01.22.24301588v1.full-text>  <https://www.oncotarget.com/article/28561/text/>  <https://www.frontiersin.org/journals/genetics/articles/10.3389/fgene.2021.652718/full>  <https://molecularcasestudies.cshlp.org/content/7/1/a005801.full.pdf>  <https://www.oncotarget.com/article/18815/text/>  <https://www.bmj.com/content/bmj/346/bmj.f2223.full.pdf>  <https://www.ncbi.nlm.nih.gov/clinvar/RCV000167830/>  <https://www.ncbi.nlm.nih.gov/clinvar/variation/37984/>  <https://www.asco.org/abstracts-presentations/ABSTRACT262743>  <https://www.frontiersin.org/journals/genetics/articles/10.3389/fgene.2022.982930/full>  <https://www.researchgate.net/publication/349447646_Treatment-emergent_neuroendocrine_prostate_cancer_with_a_germline_BRCA2_mutation_Identification_of_a_candidate_reversion_mutation_associated_with_platinumPARP-inhibitor_resistance>  <https://www.nature.com/articles/6690892.pdf>  <https://www.mdpi.com/2073-4425/12/8/1229> |
|  | Evaluation  **Summarization Accuracy & Completeness:**  **Hallucination Rate:**  **Readability & Usability:**  **Literature Coverage & Relevance:**  **Time Efficiency:** | |
| **2** | The genomic variant c.5645C>A in the BRCA2 gene results in a premature stop codon at position 1882 (p.S1882*) and is denoted as rs80358785. This variant has been classified as deleterious and pathogenic in multiple studies, indicating a strong association with disease phenotypes, particularly hereditary breast and ovarian cancer syndrome [1][3][5][6]. The presence of this nonsense mutation has been confirmed through various sequencing techniques, including next-generation sequencing (NGS) and Sanger sequencing, which have demonstrated its occurrence in both affected probands and unaffected relatives, suggesting its potential utility in cancer risk assessment and prevention strategies [1].  The BRCA2 gene plays a critical role in maintaining genomic stability by participating in the repair of DNA double-strand breaks through homologous recombination. Mutations in BRCA2 can lead to a loss of protein function, which impairs DNA repair processes and contributes to the accumulation of genetic alterations that drive oncogenesis.  The variant rs80358785 has been reported in ClinVar with a total of 19 pathogenic submissions, reinforcing its clinical significance in the context of hereditary cancer predisposition. The consistent classification of this variant as pathogenic across different studies and databases underscores its importance in genetic counseling and clinical decision-making for individuals carrying this mutation. | [[1]](https://pubmed.ncbi.nlm.nih.gov/25802882/) Hirotsu Y et al. (2015). "Detection of BRCA1 and BRCA2 germline mutations in Japanese population using next-generation sequencing." Molecular genetics & genomic medicine, 3(2)  [[2]](https://pubmed.ncbi.nlm.nih.gov/27533253/) Enyedi MZ et al. (2016). "Simultaneous detection of BRCA mutations and  large genomic rearrangements in germline DNA and FFPE tumor samples." Oncotarget, 7(38)  [[3]](https://pubmed.ncbi.nlm.nih.gov/38298632/) Song Y et al. (2024). "Next-generation sequencing-based analysis of homologous recombination repair gene variant in ovarian cancer." Heliyon, 10(2)  [[4]](https://pubmed.ncbi.nlm.nih.gov/23961350/) Solano AR et al. (2012). "BRCA1 And BRCA2 analysis of Argentinean breast/ovarian cancer patients selected for age and family history highlights a role for novel mutations of putative south-American origin." SpringerPlus, 1()  [[5]](https://pubmed.ncbi.nlm.nih.gov/39783935/) Chevrier S et al. (2025). "Nanopore adaptive sampling accurately detects nucleotide variants and improves the characterization of large-scale rearrangement for the diagnosis of cancer predisposition." Clinical and translational medicine, 15(1)  [[6]](https://pubmed.ncbi.nlm.nih.gov/22666503/) Meyer P et al. (2012). "BRCA2 mutations and triple-negative breast cancer." PloS one, 7(5)  [[7]](https://pubmed.ncbi.nlm.nih.gov/29383094/) Hirotsu Y et al. (2017). "Simultaneous detection of genetic and copy number alterations in BRCA1/2 genes." Oncotarget, 8(70)  [[8]](https://pubmed.ncbi.nlm.nih.gov/28039656/) De Silva S et al. (2017). "Novel and reported pathogenic variants in exon 11 of BRCA2 gene in a cohort of Sri Lankan young breast cancer patients." Familial cancer, 16(3)  [[9]](https://pubmed.ncbi.nlm.nih.gov/26733283/) Petersen AH et al. (2016). "Post-mortem testing; germline BRCA1/2 variant detection using archival FFPE non-tumor tissue. A new paradigm in genetic counseling." European journal of human genetics : EJHG, 24(8)  [[10]](https://pubmed.ncbi.nlm.nih.gov/29215753/) Nakagomi H et al. (2018). "Combined annotation-dependent depletion score for BRCA1/2 variants in patients with breast and/or ovarian cancer." Cancer science, 109(2)  [[11]](https://pubmed.ncbi.nlm.nih.gov/31263500/) Ueda M et al. (2019). "Challenges in Managing Patients with Hereditary Cancer at Gynecological Services." Obstetrics and gynecology international, 2019()  [[12]](https://pubmed.ncbi.nlm.nih.gov/35741847/) Abe A et al. (2022). "Prevalence of Pathogenic Germline BRCA1/2 Variants and Their Association with Clinical Characteristics in Patients with Epithelial Ovarian Cancer in a Rural Area of Japan." Genes, 13(6)  [[13]](https://pubmed.ncbi.nlm.nih.gov/38671360/) Andaluz S et al. (2024). "Using Portuguese BRCA pathogenic variation as a model to study the impact of human admixture on human health." BMC genomics, 25(1)  [[14]](https://pubmed.ncbi.nlm.nih.gov/26221963/) Wong ES et al. (2015). "Predictive Factors for BRCA1 and BRCA2 Genetic Testing in an Asian Clinic-Based Population." PloS one, 10(7)  [[15]](https://pubmed.ncbi.nlm.nih.gov/29649263/) Soukupova J et al. (2018). "Validation of CZECANCA (CZEch CAncer paNel for Clinical Application) for targeted NGS-based analysis of hereditary cancer syndromes." PloS one, 13(4) |
|  | Evaluation  **Summarization Accuracy & Completeness:**  **Hallucination Rate:**  **Readability & Usability:**  **Literature Coverage & Relevance:**  **Time Efficiency:** | |
| **3** | The **BRCA2** gene plays a crucial role in maintaining genomic stability through its involvement in homologous recombination, a process essential for repairing DNA double-strand breaks[5][6]. Mutations in **BRCA2** are associated with an increased risk of breast, ovarian, pancreatic, and other cancers[5][7]. The variant **p.S1882***, a nonsense mutation, is expected to lead to a premature stop codon, potentially resulting in a truncated protein that lacks essential functional domains. This type of mutation is generally considered pathogenic, as it disrupts the gene's ability to perform its critical roles in DNA repair[8]. However, there is evidence that some nonsense mutations in **BRCA2** can lead to in-frame skipping of exons during RNA splicing, potentially retaining partial function of the protein[8]. For **p.S1882*** specifically, there is limited direct research available in the peer-reviewed literature, and its precise impact on protein function and cancer risk remains to be fully elucidated. Therefore, further investigation is recommended to determine its clinical significance accurately. The classification and frequency of this variant in the general population or specific cohorts are not well-documented, highlighting the need for additional studies to understand its implications fully. In the absence of specific data, general principles of nonsense mutations in **BRCA2** suggest a potential for increased cancer risk, but the extent of this risk may vary based on the specific molecular and clinical context[4][5]. | [1] <https://www.ncbi.nlm.nih.gov/pmc/articles/PMC10878975/>  [2] <https://www.ncbi.nlm.nih.gov/pmc/articles/PMC7311593/>  [3] <https://pubmed.ncbi.nlm.nih.gov/33084436/>  [4] <https://pubmed.ncbi.nlm.nih.gov/30638113/>  [5] <https://www.ncbi.nlm.nih.gov/pmc/articles/PMC8307705/>  [6] <https://www.ncbi.nlm.nih.gov/pmc/articles/PMC7188823/>  [7] <https://pubmed.ncbi.nlm.nih.gov/31396961/>  [8] <https://pubmed.ncbi.nlm.nih.gov/32046981/> |
|  | Evaluation  **Summarization Accuracy & Completeness:**  **Hallucination Rate:**  **Readability & Usability:**  **Literature Coverage & Relevance:**  **Time Efficiency:** | |
| **4** | The BRCA2 gene encodes a tumor suppressor protein that plays a critical role in the maintenance of genomic stability, specifically by facilitating homologous recombination (HR) repair of double-strand DNA breaks. Loss-of-function mutations in BRCA2 significantly compromise the HR pathway, predisposing individuals to hereditary breast and ovarian cancers, among other malignancies. The variant p.S1882* (also described as c.5645C>A or S1882X) is a nonsense mutation that introduces a premature stop codon at amino acid position 1882, likely resulting in truncated or absent BRCA2 protein due to nonsense-mediated decay. Such truncating mutations are generally considered pathogenic, particularly when located in regions of the gene essential for DNA repair interactions. While BRCA2 truncations overall are well-documented in association with hereditary cancer syndromes, there is currently no direct peer-reviewed publication specifically addressing the functional or clinical significance of the p.S1882* variant. As such, its classification is inferred based on well-established patterns for similar truncating variants in BRCA2. This variant is absent from or extremely rare in large population sequencing datasets, which supports its likely pathogenic status. BRCA2 truncations, including those near or upstream of S1882, are frequently reported in individuals with breast, ovarian, pancreatic, and prostate cancers, and their presence often correlates with therapeutic vulnerability to PARP inhibitors and platinum-based agents [1,2]. However, it is important to emphasize that no study to date has directly characterized p.S1882* in isolation, and its functional impact remains to be explicitly validated. Further studies are needed to confirm its pathogenicity and to understand its potential implications for therapy resistance or reversion events, which have been observed in other BRCA2 truncating variants leading to acquired resistance to targeted treatments [3]. In summary, p.S1882* is a likely pathogenic BRCA2 nonsense variant predicted to severely disrupt protein function, but direct experimental or clinical data are lacking, and further investigation is warranted to clarify its precise biological and clinical impact. | [1] [Antoci et al., 2025](https://app.scholarai.io/paper?paper_id=DOI:10.1002/cnr2.70200&original_url=https%3A%2F%2Fonlinelibrary.wiley.com%2Fdoi%2Fabs%2F10.1002%2Fcnr2.70200)  [2] [Wei et al., 2020](https://academic.oup.com/oncolo/article-abstract/25/7/e1042/6443420)  [3] [Zhu et al., 2019](https://app.scholarai.io/paper?paper_id=DOI:10.1200/JCO.2019.37.15_suppl.1087&original_url=https%3A%2F%2Fascopubs.org%2Fdoi%2Fabs%2F10.1200%2FJCO.2019.37.15_suppl.1087) |
|  | Evaluation  **Summarization Accuracy & Completeness:**  **Hallucination Rate:**  **Readability & Usability:**  **Literature Coverage & Relevance:**  **Time Efficiency:** | |
| **5** | The BRCA2 gene is a critical tumor suppressor involved in DNA repair, particularly through homologous recombination, which is essential for maintaining genomic stability. The BRCA2 protein interacts with RAD51, a key protein in DNA repair, to facilitate the repair of double-strand breaks in DNA. When BRCA2 is functioning normally, it helps to repair damaged DNA, thereby preventing the accumulation of genetic mutations that can lead to cancer. The BRCA2 gene is located on chromosome 13q13.1 and comprises 27 exons that encode a large protein of 3,418 amino acids. The BRCA2 protein is involved in various cellular processes, including DNA repair, cell cycle regulation, and transcriptional control. The p.S1882* variant in the BRCA2 gene is a nonsense mutation that introduces a premature stop codon at position 1882, leading to a truncated, nonfunctional protein. This variant is classified as pathogenic because it results in a loss of function of the BRCA2 protein, which is crucial for DNA repair and tumor suppression. The nomenclature for this variant is c.5644C>G (p.S1882*), indicating a cytosine to guanine substitution at nucleotide position 5644, resulting in the replacement of serine with a stop codon at amino acid position 1882. The frequency of the p.S1882* variant in the general population is not well-documented, but it is considered rare. The impact of this variant is significant, as it is associated with an increased risk of developing certain cancers, particularly breast and ovarian cancers. Individuals who carry this pathogenic variant have a higher likelihood of developing these cancers compared to the general population. The exact mechanisms by which the p.S1882* variant contributes to cancer development are not fully understood, but it is believed to be related to the impaired DNA repair capacity of the truncated BRCA2 protein. This can lead to the accumulation of genetic mutations and genomic instability, which are hallmarks of cancer. There are no direct references to peer-reviewed publications specifically addressing the p.S1882* variant in the BRCA2 gene. However, the classification and interpretation of this variant are based on established guidelines for genetic variant assessment, such as those provided by the American College of Medical Genetics and Genomics (ACMG). Further research is needed to better understand the clinical implications of this variant and to develop targeted therapies for individuals who carry it. | No references available |
|  | Evaluation  **Summarization Accuracy & Completeness:**  **Hallucination Rate:**  **Readability & Usability:**  **Literature Coverage & Relevance:**  **Time Efficiency:** | |
