## Supplementary material for "Benchmarking generative AI tools for literature retrieval and summarization in genomic variant interpretation": CCND1 p.T286I.docx

| **Tool** | **Summary** | **References** |
| --- | --- | --- |
| **1** | The CCND1 gene encodes cyclin D1, a key regulator of the G1-to-S phase transition in the cell cycle through its interaction with cyclin-dependent kinases (CDK4 and CDK6). Cyclin D1 is critical for the regulation of cellular proliferation, and its dysregulation has been implicated in oncogenesis, particularly in breast, colorectal, and head and neck cancers. The CCND1 p.T286I variant represents a missense mutation at position 286, where threonine is replaced by isoleucine. Threonine 286 is a critical phosphorylation site for glycogen synthase kinase-3 beta (GSK3β), which targets cyclin D1 for ubiquitin-mediated proteasomal degradation, thereby regulating its turnover. The T286I substitution disrupts this regulatory phosphorylation, leading to increased stability and nuclear accumulation of cyclin D1, which promotes sustained CDK activation and cell cycle progression. Experimental models suggest that this mutation can lead to oncogenic transformation by preventing cyclin D1 degradation, resulting in unchecked cell proliferation. While CCND1 amplifications and overexpression are common in multiple cancer types, the prevalence of the p.T286I variant in human tumors is less well defined. Limited research indicates that it may occur in a subset of malignancies, particularly in cancers with aberrant cell cycle regulation, although its precise clinical significance remains under investigation. Due to the functional impact of this variant on cyclin D1 regulation, further studies are required to elucidate its role in tumorigenesis, potential therapeutic targeting, and prognostic implications. No peer-reviewed studies specifically addressing the CCND1 p.T286I variant were identified in the literature, highlighting the need for further investigation into its clinical and molecular consequences. | [1] Simoneschi D. Spatiotemporal Regulation of D-Type Cyclins. 2021. [ProQuest Link](https://search.proquest.com/openview/fc68fcfcc9b198d5ea1e0a3425be48a9/1?pq-origsite=gscholar&cbl=18750&diss=y). |
|  | Evaluation  **Summarization Accuracy & Completeness:**  **Hallucination Rate:**  **Readability & Usability:**  **Literature Coverage & Relevance:**  **Time Efficiency:** | |
| **2** | The genomic variant c.857C>T p.T286I rs2120121034 on the CCND1 gene results in the substitution of threonine with isoleucine at codon 286. CCND1 encodes cyclin D1, a key regulatory protein involved in cell cycle progression, particularly the transition from G1 to S phase. Cyclin D1 forms a complex with cyclin-dependent kinases 4 and 6 (CDK4/6), which phosphorylates the retinoblastoma protein (pRb), leading to the release of E2F transcription factors and the subsequent transcription of genes necessary for S phase entry.  The phosphorylation of cyclin D1 at Thr286 by GSK3beta is crucial for its nuclear export and subsequent cytoplasmic proteolysis via the 26S proteasome. This regulatory mechanism is important for preventing aberrant cell proliferation. Mutations in CCND1 are rare in human cancers, but the T286I mutation has been identified in endometrial cancer. Functional studies of the T286I mutant in HEK-293T cells have shown that it leads to the accumulation of cyclin D1 in the nucleus, decreased expression of pRb, and an increase in cell proliferation as evidenced by an increased number of colonies compared to wild-type cyclin D1[1].  Furthermore, the presence of the T286I mutation in endometrial cancer has been associated with coexisting mutations in other genes involved in the RAS/PI3K pathway, such as K-ras, PIK3CA, and PTEN, which may contribute to the nuclear accumulation of cyclin D1 and the dysregulation of cell cycle control[1]. The T286I mutation has been shown to promote oncogenic activation in endometrial cancer by stabilizing cyclin D1 in the nucleus, thereby overcoming the regulatory effects of pRb and inducing cell proliferation[3].  In the context of endometrioid endometrial carcinoma (EEC), CDK4/6-specific activity (CDK4/6SA) has been proposed as a novel biomarker for prognosis and chemo-sensitivity. High CDK4/6SA has been associated with poor prognosis in pathologically low-risk patients and may indicate a higher likelihood of recurrence. Additionally, tumours with high CDK4/6SA may be more sensitive to platinum-based chemotherapy, suggesting that the cell-cycle augmentation induced by high CDK4/6SA could be associated with sensitivity to such treatments[2].  In summary, the c.857C>T p.T286I rs2120121034 variant in the CCND1 gene is a rare mutation that has functional implications in cell cycle regulation and has been associated with oncogenic activation in endometrial cancer. This variant leads to the stabilization and nuclear accumulation of cyclin D1, which can promote cell proliferation and may serve as a potential biomarker for prognosis and therapeutic response in EEC[1][2][3]. | [[1]](https://pubmed.ncbi.nlm.nih.gov/23733133/) Ikeda Y et al. (2013). "Cyclin D1 harboring the T286I mutation promotes oncogenic activation in endometrial cancer." Oncology reports, 30(2)  [[2]](https://pubmed.ncbi.nlm.nih.gov/26554657/) Ikeda Y et al. (2015). "Prognostic importance of CDK4/6-specific activity as a predictive marker for recurrence in patients with endometrial cancer, with or without adjuvant chemotherapy." British journal of cancer, 113(10)  [[3]](https://aacrjournals.org/cancerres/article/73/8_Supplement/1748/587445/Abstract-1748-Cyclin-D1-harboring-T286I-mutation) Y. Ikeda et al. (2013). "Abstract 1748: Cyclin D1 harboring T286I mutation promotes oncogenic activation in endometrial cancer." Cancer Research, 73 |
|  | Evaluation  **Summarization Accuracy & Completeness:**  **Hallucination Rate:**  **Readability & Usability:**  **Literature Coverage & Relevance:**  **Time Efficiency:** | |
| **3** | The *CCND1* gene encodes cyclin D1, a key regulator of the cell cycle's G1/S transition[11](https://www.genecards.org/cgi-bin/carddisp.pl?gene=CCND1). Cyclin D1 forms a complex with cyclin-dependent kinases CDK4 or CDK6, and this complex is essential for cell cycle progression[11](https://www.genecards.org/cgi-bin/carddisp.pl?gene=CCND1). Overexpression of *CCND1* can lead to unchecked cell proliferation and has been observed in various cancers[9](https://www.nature.com/articles/jhg200475)[11](https://www.genecards.org/cgi-bin/carddisp.pl?gene=CCND1). The *CCND1* gene, located on chromosome 11q13, consists of 13,388 base pairs that translate into a 295 amino acid protein[13](https://en.wikipedia.org/wiki/Cyclin_D1). The T286I variant, a mutation where threonine is substituted for isoleucine at codon 286, has been identified in endometrial cancer[10](https://pubmed.ncbi.nlm.nih.gov/23733133/). Functional analyses indicate that this mutation results in the accumulation of cyclin D1 in the nucleus and promotes cell proliferation[10](https://pubmed.ncbi.nlm.nih.gov/23733133/). Specifically, the mutant cyclin D1 (T286I) inhibits the expression of pRb, a tumor suppressor protein, and increases colony formation compared to wild-type cyclin D1[10](https://pubmed.ncbi.nlm.nih.gov/23733133/). The *CCND1* T286I mutation has been reported in two out of 88 endometrial cancer tissue specimens analyzed[10](https://pubmed.ncbi.nlm.nih.gov/23733133/). Tumors harboring this mutation often present with coexisting mutations in K-ras, PIK3CA, and/or PTEN[10](https://pubmed.ncbi.nlm.nih.gov/23733133/). Further research is needed to fully elucidate the prevalence, specific functional impacts, and clinical significance of the CCND1 p.T286I variant across diverse cancer types.[10](https://pubmed.ncbi.nlm.nih.gov/23733133/) | [1] <https://pubmed.ncbi.nlm.nih.gov/39113852/>  [2] <https://www.ncbi.nlm.nih.gov/pmc/articles/PMC10298313/>  [3] <https://www.ncbi.nlm.nih.gov/pmc/articles/PMC8923789/>  [3] <https://pubmed.ncbi.nlm.nih.gov/26676638/>  [4] <https://www.ncbi.nlm.nih.gov/pmc/articles/PMC7426966/>  [5] <https://www.ncbi.nlm.nih.gov/pmc/articles/PMC9274483/>  [6] <https://www.ncbi.nlm.nih.gov/pmc/articles/PMC8255842/>  [7] <https://pubmed.ncbi.nlm.nih.gov/34564768/>  [8] <https://www.nature.com/articles/jhg200475>  [9] <https://pubmed.ncbi.nlm.nih.gov/23733133/>  [10] <https://www.genecards.org/cgi-bin/carddisp.pl?gene=CCND1>  [11] <https://www.youtube.com/watch?v=H7DqNZ4yPK8>  [12] <https://en.wikipedia.org/wiki/Cyclin_D1>  [13] <https://pmc.ncbi.nlm.nih.gov/articles/PMC4247015/>  [14] <https://www.ncbi.nlm.nih.gov/gene/595>  [15] <https://stackoverflow.com/questions/76792247/unable-to-generate-summary-in-bullet-points-using-langchain> |
|  | Evaluation  **Summarization Accuracy & Completeness:**  **Hallucination Rate:**  **Readability & Usability:**  **Literature Coverage & Relevance:**  **Time Efficiency:** | |
| **4** | The CCND1 gene encodes cyclin D1, a critical regulator of cell cycle progression, particularly in the transition from the G1 to S phase. Cyclin D1 forms a complex with cyclin-dependent kinases CDK4 and CDK6, facilitating the phosphorylation and inactivation of the retinoblastoma protein (pRb), which leads to cell cycle progression. The CCND1 p.T286I variant, located in the C-terminal region of cyclin D1, is a missense mutation that substitutes threonine with isoleucine at codon 286. This variant has been identified in endometrial cancers, where it promotes oncogenic activation by inhibiting the phosphorylation of cyclin D1 at Thr286, a process normally mediated by GSK3β. Phosphorylation at Thr286 typically triggers cyclin D1's nuclear export and subsequent proteolysis via the 26S proteasome. The p.T286I mutation disrupts this regulatory mechanism, leading to the nuclear accumulation of cyclin D1 and enhanced cell proliferation. Research has shown that this variant is functional in inducing cell proliferation and has been detected in approximately 2.3% of endometrial cancer cases, often co-occurring with other mutations such as K-ras, PIK3CA, and PTEN. The frequency of this variant in the general population is not well-documented, and its classification in clinical databases is limited. Further investigation is necessary to fully understand the impact of the CCND1 p.T286I variant on cancer development and progression, as well as its potential as a therapeutic target. | [1] Ikeda, K., et al. (2013). Cyclin D1 harboring the T286I mutation promotes oncogenic activation in endometrial cancer. Oncology Reports, 30(5), 2173-2180.  [2] Xu, J., & Lin, D. I. (2018). Oncogenic c-terminal cyclin D1 (CCND1) mutations are enriched in endometrioid endometrial adenocarcinomas. PLoS ONE, 13(7), e0199688. |
|  | Evaluation  **Summarization Accuracy & Completeness:**  **Hallucination Rate:**  **Readability & Usability:**  **Literature Coverage & Relevance:**  **Time Efficiency:** | |
| **5** | The CCND1 gene encodes cyclin D1, a regulatory protein integral to cell cycle progression, particularly the transition from the G1 to S phase. Cyclin D1 forms complexes with cyclin-dependent kinases CDK4 or CDK6, phosphorylating the retinoblastoma protein (pRb), thereby promoting cell cycle advancement. Alterations in CCND1, such as mutations, amplifications, or overexpression, can disrupt normal cell cycle control and are frequently observed in various cancers. citeturn0search1 The p.T286I variant denotes a threonine-to-isoleucine substitution at amino acid position 286 of cyclin D1. This threonine residue is crucial for phosphorylation-dependent nuclear export and proteasomal degradation of cyclin D1. Phosphorylation at Thr286 facilitates cyclin D1's recognition by the CRM1 export pathway, leading to its translocation from the nucleus to the cytoplasm and subsequent degradation. The T286I mutation impairs this regulatory mechanism, resulting in the aberrant accumulation of cyclin D1 within the nucleus. citeturn0search0 Functionally, the T286I mutation has been identified in endometrial cancers and is associated with increased oncogenic potential. In vitro studies demonstrate that cells expressing the T286I mutant exhibit enhanced proliferation compared to those expressing wild-type cyclin D1. This heightened proliferative capacity is attributed to sustained nuclear presence of cyclin D1, leading to continuous activation of CDK4/6 and persistent phosphorylation of pRb, thereby promoting uncontrolled cell cycle progression. citeturn0search11 The prevalence of the T286I mutation varies among cancer types. Notably, this mutation has been detected in a subset of endometrial carcinomas, suggesting a role in the pathogenesis of these tumors. The exact frequency across different malignancies remains to be fully elucidated, necessitating further large-scale genomic studies to determine its distribution and clinical significance. citeturn0search3 Clinically, the presence of the T286I mutation may have therapeutic implications. The constitutive activation of the cyclin D1-CDK4/6 axis due to impaired degradation of the mutant protein suggests that tumors harboring this mutation could be particularly sensitive to CDK4/6 inhibitors. This hypothesis aligns with the mechanism of action of these inhibitors, which aim to restore control over cell cycle progression by targeting hyperactive CDK4/6 complexes. However, clinical validation is required to confirm the efficacy of CDK4/6 inhibitors in patients with T286I-mutant tumors. citeturn0search6 In summary, the CCND1 p.T286I variant disrupts normal regulatory mechanisms governing cyclin D1 localization and degradation, leading to its nuclear accumulation and enhanced oncogenic activity. While its occurrence has been documented in certain cancers, comprehensive data on its prevalence and impact across diverse tumor types are limited. Further research is warranted to elucidate its role in tumorigenesis and to explore potential targeted therapeutic strategies for affected patients. | [1] [ncbi.nlm.nih.gov595](https://www.ncbi.nlm.nih.gov/gene/595)  [2] [GeneCardsCCND1 Gene - Cyclin D1 - GeneCards](https://www.genecards.org/cgi-bin/carddisp.pl?gene=CCND1)  [3] [NatureAberrant Cyclin D1 splicing in cancer: from molecular mechanism to ...6 aprile 2023](https://www.nature.com/articles/s41419-023-05763-7)  [4] [pmc.ncbi.nlm.nih.govOncogenic c-terminal cyclin D1 (CCND1) mutations are enriched in ...3 luglio 2018](https://pmc.ncbi.nlm.nih.gov/articles/PMC6029777/)  [5] [researchgate.net](https://www.researchgate.net/publication/326163066_Oncogenic_c-terminal_cyclin_D1_CCND1_mutations_are_enriched_in_endometrioid_endometrial_adenocarcinomas)  [Oncogenic c-terminal cyclin D1 (CCND1) mutations are enriched in ...3 luglio 2018](https://www.researchgate.net/publication/326163066_Oncogenic_c-terminal_cyclin_D1_CCND1_mutations_are_enriched_in_endometrioid_endometrial_adenocarcinomas)  [6] [oncotarget.comCCND1 mutations increase protein stability and promote ibrutinib ...8 novembre 2016](https://www.oncotarget.com/article/12434/text/)  [7] [spandidos-publications.comCyclin D1 harboring the T286I mutation promotes oncogenic ...3 giugno 2013](https://www.spandidos-publications.com/10.3892/or.2013.2515)  [8] [aacrjournals.orgAbstract 1748: Cyclin D1 harboring T286I mutation promotes ...15 aprile 2013](https://aacrjournals.org/cancerres/article/73/8_Supplement/1748/587445/Abstract-1748-Cyclin-D1-harboring-T286I-mutation)  [9] [spandidos-publications.comCyclin D1 harboring the T286I mutation promotes oncogenic](https://www.spandidos-publications.com/10.3892/or.2013.2515/download)  [10] [pmc.ncbi.nlm.nih.govt(11;14](https://pmc.ncbi.nlm.nih.gov/articles/PMC6119140/))  [11] [pmc.ncbi.nlm.nih.gov](https://pmc.ncbi.nlm.nih.gov/articles/PMC9722596/)  [12] [en.wikipedia.orgCyclin D1 - Wikipedia Cyclin D1](https://en.wikipedia.org/wiki/Cyclin_D1)  [13] [Nature](https://www.nature.com/articles/s41598-018-25141-0)  [14] [oncokb.org](https://www.oncokb.org/gene/CCND1/T286I)  [15] [UniProtG1/S-specific cyclin-D1 - Homo sapiens (Human)](https://www.uniprot.org/uniprotkb/P24385/entry)  [16] [cancerindex.orgBCL1 Gene (CCND1) \| Cancer Genetics](https://www.cancerindex.org/geneweb/BCL1.htm)  [17] [ccsm.uth.eduPCTfuncDB: functional knowledgebase](https://ccsm.uth.edu/PCTfuncDB/gene_search_result.cgi?page6=2&quick_search=CCND1&type=quick_search) |
|  | Evaluation  **Summarization Accuracy & Completeness:**  **Hallucination Rate:**  **Readability & Usability:**  **Literature Coverage & Relevance:**  **Time Efficiency:** | |
