## Supplementary material for "Benchmarking generative AI tools for literature retrieval and summarization in genomic variant interpretation": EGFR p.A289V.docx

| **Tool** | **Summary** | **References** |
| --- | --- | --- |
| **1** | The EGFR (epidermal growth factor receptor) gene encodes a transmembrane tyrosine kinase receptor involved in regulating cell growth, survival, and differentiation. Dysregulation of EGFR signaling is a key driver in multiple cancers, including non-small cell lung cancer (NSCLC) and glioblastoma. The p.A289V variant is a missense mutation located in exon 7 of the extracellular domain (ECD) of EGFR. While EGFR mutations commonly occur in the kinase domain (exons 18-21), alterations in the ECD, including p.A289V, have been reported in both NSCLC and gliomas. This mutation is rare but has been identified in a subset of tumors, suggesting a potential role in oncogenesis. In NSCLC, p.A289V has been associated with sensitivity to EGFR tyrosine kinase inhibitors (TKIs), including Icotinib, as documented in a case report where a patient harboring this mutation responded favorably to treatment, indicating its potential as a targetable alteration [1]. Another study found that extracellular domain mutations, including A289 variants, might expand the EGFR mutation spectrum relevant for targeted therapies, but further validation is required [2]. The p.A289V mutation has also been identified in gliomas, where EGFR alterations are frequent and associated with tumor progression and therapeutic resistance [3]. In glioblastomas, the mutation may contribute to oncogenesis through ligand-independent receptor activation or dimerization, similar to other extracellular domain mutations. However, its precise functional impact remains unclear, and more studies are necessary to determine its role in glioblastoma pathogenesis. Recent evidence also suggests that EGFR A289V may coexist with other driver mutations or amplifications, complicating its clinical interpretation [4]. Given the rarity of this variant, its exact prevalence in different cancers remains uncertain, but it appears infrequently in large-scale genomic databases. Current data on therapeutic responses are limited, and while some reports suggest potential benefit from EGFR-targeted therapies, further studies are necessary to establish its predictive value for treatment selection. Due to the limited but emerging evidence regarding the oncogenic role of EGFR p.A289V, additional functional and clinical studies are recommended to elucidate its significance and therapeutic implications. | [1] Dai L, Su X, Lu L, Lv D. Nonsmall cell lung cancer with rare exon 7 p.A289V mutation in the EGFR gene responds to Icotinib treatment: a case report. [Medicine (Baltimore)](https://journals.lww.com/md-journal/fulltext/2018/12210/Nonsmall_cell_lung_cancer_with_rare_exon_7_p_A289V.122.aspx), 2018.  [2] Wang W, Xu C, Chen Y, Cai X, Fang Y, Zhang Q, Zhu Y. An EGFR extracellular domain mutation data in the East Asian non-small cell lung cancer populations and response to icotinib: A multicenter study. J Clin Oncol, 2019.  [3] Zhang H, Tao R, Zhu X, Luo N, Qi Y, Li M, Sun T, Qi C. The landscape of EGFR mutation in Chinese patients with glioma. J Clin Oncol, 2022.  [4] He L, Li L. Case report: MSI-H, EGFR mutation, and ground-glass nodules as diffuse pulmonary hematogenous metastases. [Front Immunol](https://www.frontiersin.org/journals/immunology/articles/10.3389/fimmu.2024.1478205/full), 2024. |
|  | Evaluation  **Summarization Accuracy & Completeness:**  **Hallucination Rate:**  **Readability & Usability:**  **Literature Coverage & Relevance:**  **Time Efficiency:** | |
| **2** | The epidermal growth factor receptor (EGFR) gene plays a crucial role in cell proliferation, differentiation, and survival by mediating signals from the extracellular environment to the cell interior. Variants in the EGFR gene, such as p.A289V, can potentially disrupt these processes, although specific research on this variant is limited. Generally, EGFR mutations are well-documented in various cancers, including non-small cell lung cancer and glioblastoma, where they can lead to aberrant activation of downstream signaling pathways, such as the ERK pathway[2](https://pubmed.ncbi.nlm.nih.gov/35850069/)[3](https://www.ncbi.nlm.nih.gov/pmc/articles/PMC9527520/). In cancers, EGFR mutations often result in constitutive activation of the receptor, promoting tumor growth and resistance to therapies like gefitinib[6](https://www.ncbi.nlm.nih.gov/pmc/articles/PMC10169458/). The classification and nomenclature of EGFR variants are typically based on their location and functional impact within the gene. However, without specific studies on the p.A289V variant, its frequency and clinical impact remain uncertain. Further investigation is recommended to determine the functional significance and potential clinical implications of this variant. While there are extensive studies on other EGFR mutations, direct evidence for p.A289V is currently lacking, emphasizing the need for additional research to clarify its role in disease pathology. | [1] <https://www.semanticscholar.org/paper/Key-Experiments-in-Practical-Developmental-Biology%3A-Duffy-Perrimon/ff09264334076786c1306b6e575776df0242031f>  [2] <https://pubmed.ncbi.nlm.nih.gov/35850069/>  [3] <https://pmc.ncbi.nlm.nih.gov/articles/PMC9527520/>  [4] <https://pubmed.ncbi.nlm.nih.gov/33405744/>  [5] <https://pmc.ncbi.nlm.nih.gov/articles/PMC8113371/>  [6] <https://pmc.ncbi.nlm.nih.gov/articles/PMC10169458/>  [7] <https://pubmed.ncbi.nlm.nih.gov/24469720/>  [8] <https://pubmed.ncbi.nlm.nih.gov/30470824/> |
|  | Evaluation  **Summarization Accuracy & Completeness:**  **Hallucination Rate:**  **Readability & Usability:**  **Literature Coverage & Relevance:**  **Time Efficiency:** | |
| **3** | The EGFR gene encodes the epidermal growth factor receptor, a receptor-type tyrosine kinase involved in the regulation of cellular homeostasis. EGFR is a critical component in cell differentiation, proliferation, and various signaling pathways. Dysregulated EGFR signaling is implicated in the pathogenesis of numerous epithelial malignancies, contributing to tumor growth, neovascularization, invasion, and metastasis [1].  The genomic variant c.866C>T p.A289V rs149840192 is a missense mutation located in the extracellular domain of the EGFR protein. This mutation has been associated with a range of neoplasms, including glioblastoma, low-grade glioma, and head and neck neoplasms. It is cataloged in the dbSNP database as a pathogenic allele [1]. In the context of non-small cell lung cancer (NSCLC), the p.A289V mutation is particularly rare, and its presence in NSCLC and the response to tyrosine kinase inhibitors (TKIs) have not been widely reported [1].  A study has shown that a patient with NSCLC harboring the p.A289V mutation responded positively to treatment with the first-generation EGFR-TKI Icotinib, suggesting that this mutation may be a TKI-sensitive site, especially in NSCLCs. The patient exhibited a partial response to the treatment, with a significant decrease in tumor volume after 5 months, and experienced only mild adverse events [1]. This suggests that the p.A289V mutation may activate oncogenic signaling pathways without disrupting the binding between the ATP-binding site of the EGFR protein and TKIs, allowing drugs like Icotinib and Erlotinib to inhibit the signal transduction cascade of the mutant receptor [1].  In glioblastoma, the p.A289V mutation has been associated with a highly invasive phenotype, with studies indicating increased tumor invasion and dependence on ERK-mediated expression of matrix metalloproteinase-1. This mutation has been proposed as a potential molecular marker for responsiveness to therapy with EGFR-targeting antibodies, as tumor growth expressing EGFRA289V was attenuated with an antibody against a cryptic epitope [2].  The detection of oncogenic EGFR extracellular domain mutations, including p.A289V, in lung adenocarcinoma patients who are "pan-negative" for other common driver mutations, underscores the importance of comprehensive genomic profiling in the management of lung cancer. This approach allows for the identification of rare but potentially actionable mutations that may guide targeted therapy [3].  In summary, the c.866C>T p.A289V rs149840192 variant in the EGFR gene represents a rare but clinically significant mutation with implications for targeted therapy in NSCLC and glioblastoma. The mutation's sensitivity to TKIs in NSCLC and its association with an invasive phenotype in glioblastoma highlight the need for further research to understand the molecular mechanisms underlying these observations and to confirm the therapeutic potential of targeting this variant in larger patient cohorts. ClinVar has reported a total of 6 pathogenic submissions for this variant, supporting its clinical relevance. | [[1]](https://pubmed.ncbi.nlm.nih.gov/30572543/) Dai L et al. (2018). "Nonsmall cell lung cancer with rare exon 7 p.A289V mutation in the EGFR gene responds to Icotinib treatment: A case report." Medicine, 97(51)  [[2]](https://pubmed.ncbi.nlm.nih.gov/29990498/) Binder ZA et al. (2018). "Epidermal Growth Factor Receptor Extracellular Domain Mutations in Glioblastoma Present Opportunities for Clinical Imaging and Therapeutic Development." Cancer cell, 34(1)  [[3]](https://pubmed.ncbi.nlm.nih.gov/26486077/) Wang R et al. (2015). "Comprehensive investigation of oncogenic driver mutations in Chinese non-small cell lung cancer patients." Oncotarget, 6(33)  [[4]](https://pubmed.ncbi.nlm.nih.gov/22203214/) Longo SL et al. (2012). "Bay846, a new irreversible small molecule inhibitor of EGFR and Her2, is highly effective against malignant brain tumor models." Investigational new drugs, 30(6)  [[5]](https://pubmed.ncbi.nlm.nih.gov/39697713/) Dempke WCM et al. (2024). "Targeting C797S mutations and beyond in non-small cell lung cancer-a mini-review." Translational cancer research, 13(11)  [[6]](https://www.mdpi.com/2571-6980/5/2/11) G. Mestrallet et al. (2024). "Prediction of Glioma Resistance to Immune Checkpoint Inhibitors Based on Mutation Profile" Neuroglia,  [[7]](https://aacrjournals.org/cancerres/article/76/14_Supplement/4497/611771/Abstract-4497-NYGC-glioblastoma-clinical-outcomes) K. Wrzeszczynski et al. (2016). "Abstract 4497: NYGC glioblastoma clinical outcomes pilot study: Discovering therapeutic potential in glioblastoma through integrative genomics" Cancer Research, 76  [[8]](https://aacrjournals.org/cancerres/article/84/6_Supplement/1229/737806/Abstract-1229-BDTX-1535-A-MasterKey-EGFR-inhibitor) Etienne Dardenne et al. (2024). "Abstract 1229: BDTX-1535: A MasterKey EGFR inhibitor targeting classical and non-classical oncogenic driver mutations and the C797S acquired resistance mutation to address the evolving molecular landscape of EGFR mutant NSCLC" Cancer Research,  [[9]](https://ascopubs.org/doi/10.1200/JCO.2023.41.16_suppl.e16208) Xiaobing Wu et al. (2023). "EGFR/ERBB2 aberrations in Chinese biliary tract carcinomas." Journal of Clinical Oncology,  [[10]](https://aacrjournals.org/cancerres/article/84/6_Supplement/5444/740037/Abstract-5444-EGFR-missense-mutation-s-induce) S. Weaver et al. (2024). "Abstract 5444: EGFR missense mutation(s) induce OLIG2 expression to regulate glioma stem cells maintenance and therapeutic resistance" Cancer Research,  [[11]](https://academic.oup.com/neuro-oncology/article/25/Supplement_5/v299/7406357) Benjamin Lin et al. (2023). "MODL-05. ELUCIDATING THE EPIGENETIC DRIVEN TRANSCRIPTIONAL PROGRAM UNDERLYING THE INVASIVE PHENOTYPE OF THE NOVEL EGFRA289VARIANT OF GBM" Neuro-Oncology,  [[12]](https://pubmed.ncbi.nlm.nih.gov/39959305/) S. Ghisai et al. (2024). "Transcriptomic analysis of EGFR co-expression and activation in glioblastoma reveals associations with its ligands" Neuro-Oncology Advances, 7  [[13]](https://academic.oup.com/neuro-oncology/article/26/Supplement_5/v15/7824865) S. Ghisai et al. (2024). "OS01.4.A MODELING EGFR REGULATION IN GLIOBLASTOMA REVEALS ASSOCIATIONS WITH ITS LIGANDS" Neuro-Oncology,  [[14]](https://academic.oup.com/neuro-oncology/article/21/Supplement_6/vi150/5619905) M. Lobbous et al. (2019). "PATH-35. RETROSPECTIVE ANALYSIS OF 145 PATIENTS WITH GLIOBLASTOMA; CORRELATING MOLECULAR ALTERATION INCIDENCE WITH DEMOGRAPHICS, TUMOR LOCATION, AND PROGNOSIS" Neuro-Oncology,  [[15]](https://aacrjournals.org/cancerres/article/75/23_Supplement/A29/606294/Abstract-A29-Targeting-the-EGFR-in-high-grade) V. Pillay et al. (2015). "Abstract A29: Targeting the EGFR in high-grade glioma" Brain, 75 |
|  | Evaluation  **Summarization Accuracy & Completeness:**  **Hallucination Rate:**  **Readability & Usability:**  **Literature Coverage & Relevance:**  **Time Efficiency:** | |
| **4** | The EGFR gene encodes the epidermal growth factor receptor, a transmembrane protein with tyrosine kinase activity that plays a crucial role in cell proliferation, differentiation, and survival. The p.A289V variant in the EGFR gene is a missense mutation that substitutes valine for alanine at position 289 in the protein sequence. This variant is located in the extracellular domain of the EGFR protein, which is involved in ligand binding and receptor dimerization. The p.A289V variant has been identified in various cancer types, including glioblastoma and non-small cell lung cancer (NSCLC), with a reported frequency of 0.18% in AACR GENIE cases, notably in glioblastoma multiforme, glioblastoma, lung adenocarcinoma, anaplastic astrocytoma, and astrocytoma . Functional studies have shown that the p.A289V mutation results in constitutive phosphorylation of EGFR, leading to persistent activation of downstream signaling pathways, such as ERK phosphorylation, which enhances the expression of MMP1 and promotes a pro-invasive phenotype in glioblastoma cells . Clinical data indicate that patients with glioblastoma harboring the p.A289V mutation have a significantly reduced overall survival compared to those without this mutation . Additionally, the p.A289V variant has been shown to sensitize glioblastoma cells to the antibody-drug conjugate depatux-m in combination with temozolomide, suggesting a potential therapeutic strategy for tumors with this mutation . In NSCLC, the p.A289V mutation has been reported to respond to tyrosine kinase inhibitors (TKIs) such as Icotinib, although further validation in larger patient cohorts is needed . The impact of the p.A289V variant on EGFR signaling and its potential as a therapeutic target highlight the importance of further investigation to understand its role in cancer development and treatment response. While the available data suggest a significant clinical impact, the precise mechanisms by which this variant influences tumor behavior and therapeutic outcomes remain to be fully elucidated. | [[0]](https://pubmed.ncbi.nlm.nih.gov/29990498/) Epidermal Growth Factor Receptor Extracellular Domain Mutations in Glioblastoma Present Opportunities for Clinical Imaging and Therapeutic Development - PMC  [[2]](https://www.mycancergenome.org/content/gene/egfr/) EGFR A289V - My Cancer Genome  [[3]](https://pmc.ncbi.nlm.nih.gov/articles/PMC7212878/) EGFR mutations are associated with response to depatux-m in combination with temozolomide and result in a receptor that is hypersensitive to ligand \| Neuro-Oncology Advances \| Oxford Academic  [[4]](https://pubmed.ncbi.nlm.nih.gov/30572543/) Nonsmall cell lung cancer with rare exon 7 p.A289V mutation in the EGFR gene responds to Icotinib treatment: A case report - PubMed |
|  | Evaluation  **Summarization Accuracy & Completeness:**  **Hallucination Rate:**  **Readability & Usability:**  **Literature Coverage & Relevance:**  **Time Efficiency:** | |
| **5** | The epidermal growth factor receptor (EGFR) gene encodes a transmembrane tyrosine kinase receptor that plays a pivotal role in regulating cellular processes such as proliferation, differentiation, and survival. Mutations in EGFR are implicated in various malignancies, notably glioblastoma (GBM) and non-small cell lung cancer (NSCLC). The p.A289V mutation, a missense alteration where alanine is substituted by valine at codon 289, occurs within the extracellular domain of EGFR. This specific mutation has been identified in approximately 6% of GBM cases, correlating with increased tumor invasiveness and a median overall survival of six months, significantly shorter than the typical 15-month median for GBM patients [1]. Functional studies demonstrate that the A289V mutation leads to EGFR activation, promoting tumor growth and invasiveness. In vivo experiments using murine models have shown that tumors expressing this mutation exhibit enhanced aggressiveness and reduced survival rates compared to those with wild-type EGFR. Importantly, therapeutic interventions targeting EGFR, such as the monoclonal antibody mAb806, have yielded promising results in preclinical settings, effectively reducing tumor burden and extending survival in models harboring the A289V mutation [1]. Beyond GBM, the p.A289V mutation has been sporadically reported in other cancers, including NSCLC. A case study detailed an advanced NSCLC patient with the A289V mutation who achieved a partial response to first-generation EGFR tyrosine kinase inhibitors (TKIs), specifically icotinib, with a progression-free survival of over five months [2]. Another report described a lung adenocarcinoma patient harboring both the A289V and L858R EGFR mutations, who initially responded to icotinib but experienced disease progression after six months, suggesting potential primary resistance associated with the A289V mutation [3]. These findings underscore the need for further research to elucidate the mutation's role in therapeutic resistance and to optimize treatment strategies. In summary, the EGFR p.A289V mutation is associated with aggressive tumor behavior and poorer prognosis, particularly in GBM. While targeted therapies like mAb806 show potential, their efficacy requires validation in clinical trials. The mutation's impact in other cancers, such as NSCLC, remains to be fully characterized, highlighting the necessity for additional studies to inform clinical decision-making. | [1] Binder ZA, Thorne AH, Bakas S, et al. Epidermal growth factor receptor extracellular domain mutations in glioblastoma present opportunities for clinical imaging and therapeutic development. *Cancer Cell*. 2018;34(1):163-177.e7.  [2] Nonsmall cell lung cancer with rare exon 7 p.A289V mutation in the EGFR gene: A case report and literature review. *Medicine (Baltimore)*. 2018;97(52):e13824.  [3] He L, Li L. Case report: MSI-H, EGFR mutation, and ground-glass nodules as diffuse pulmonary hematogenous metastases. *Front Immunol*. 2024;15:1478205.  Others:  <https://europepmc.org/article/PMC/PMC6319919>  <https://www.pharmacytimes.com/view/egfr-mutation-linked-to-deadlier-cases-of-gliobastoma>  <https://www.pennmedicine.org/news/news-releases/2018/july/penn-led-study-finds-mutation-driving-deadlier-brain-tumors-and-potential-therapy-to-stop-it>  <https://www.frontiersin.org/journals/oncology/articles/10.3389/fonc.2022.954026/full>  <https://www.frontiersin.org/journals/immunology/articles/10.3389/fimmu.2024.1478205/full>  <https://www.frontiersin.org/journals/immunology/articles/10.3389/fimmu.2024.1478205/full>  <https://academic.oup.com/neuro-oncology/article/25/Supplement_5/v37/7405755?login=false>  <https://www.frontiersin.org/journals/oncology/articles/10.3389/fonc.2022.954026/full>  <https://academic.oup.com/neuro-oncology/article/24/12/2035/6705404?login=false>  <https://europepmc.org/article/PMC/PMC6319919>  <https://pmkb.weill.cornell.edu/>  <https://cancer.sanger.ac.uk/cosmic/mutation/overview?id=103092003>  <https://cancer.sanger.ac.uk/cosmic/study/overview?pmid=30572543>  <https://www.pennmedicine.org/news/news-releases/2018/july/penn-led-study-finds-mutation-driving-deadlier-brain-tumors-and-potential-therapy-to-stop-it>  <https://www.ncbi.nlm.nih.gov/medgen/1689324> |
|  | Evaluation  **Summarization Accuracy & Completeness:**  **Hallucination Rate:**  **Readability & Usability:**  **Literature Coverage & Relevance:**  **Time Efficiency:** | |
