## Supplementary material for "Benchmarking generative AI tools for literature retrieval and summarization in genomic variant interpretation": HER2 p.G776V.docx

| **Tool** | **Summary** | **References** |
| --- | --- | --- |
| **1** | The HER2 gene (also known as ERBB2) encodes a receptor tyrosine kinase that belongs to the epidermal growth factor receptor (EGFR) family, playing a critical role in the regulation of cell growth, survival, and differentiation through activation of downstream signaling pathways such as PI3K/AKT and MAPK. HER2 is a well-established oncogene frequently amplified or overexpressed in various cancers, most notably breast cancer, where it serves as both a prognostic marker and a therapeutic target. The variant p.G776V in HER2 represents a missense mutation substituting glycine with valine at position 776, located within the tyrosine kinase domain of the protein, a region crucial for its enzymatic activity and signal transduction. Despite the importance of this domain, there is currently no peer-reviewed literature specifically describing the clinical significance, functional impact, or frequency of the p.G776V variant in HER2. This absence of direct evidence precludes definitive classification of this variant’s pathogenicity or its role in disease phenotypes. HER2 mutations, in general, have been implicated in oncogenic activation and resistance to targeted therapies in certain cancer subtypes, but the precise effects are highly variant-specific. Given the lack of published data on p.G776V, it remains a variant of uncertain significance, and its contribution to oncogenesis or therapeutic response cannot be reliably inferred. Functional studies and clinical correlation are recommended to elucidate the biological consequences of this substitution. Until such data are available, caution is warranted in clinical interpretation, and this variant should be considered in the context of comprehensive molecular profiling and patient phenotype. No direct peer-reviewed references on p.G776V were identified in the literature or genomic variant databases at the time of this report, highlighting the need for further research on this specific alteration. | 1. <https://www.ncbi.nlm.nih.gov/pmc/articles/PMC11412846/> 2. <https://www.semanticscholar.org/paper/d511284412ade16cb42b3adc1306fb2c6773bed2> 3. <https://www.ncbi.nlm.nih.gov/pmc/articles/PMC10543094/> 4. <https://www.semanticscholar.org/paper/c57feb1c1a1ff5433e326c9ec092c9bd74acecc8> 5. <https://www.semanticscholar.org/paper/fefd4c0dd393801f5de4ea0976fce131b8dc1b0f> 6. <https://www.semanticscholar.org/paper/6d120bccf683272c9222fe1b64e91abf09b02cfe> 7. <https://www.ncbi.nlm.nih.gov/pmc/articles/PMC10003572/> 8. <https://www.ncbi.nlm.nih.gov/pmc/articles/PMC11454008/> |
|  | Evaluation  **Summarization Accuracy & Completeness:**  **Hallucination Rate:**  **Readability & Usability:**  **Literature Coverage & Relevance:**  **Time Efficiency:** | |
| **2** | The HER2 gene (ERBB2) encodes a receptor tyrosine kinase integral to the EGFR family, playing a pivotal role in regulating cellular proliferation and survival through pathways such as PI3K-AKT-mTOR and RAS-RAF-MEK-ERK. The p.G776V variant, a glycine-to-valine substitution at codon 776 within the kinase domain (exon 20), has been identified in various cancers, including breast and non-small cell lung carcinoma (NSCLC). Although its precise biochemical impact remains uncharacterized, clinical observations suggest oncogenic potential. Notably, a patient with NSCLC harboring the p.G776V mutation exhibited a progression-free survival of 17.5 months when treated with pyrotinib, indicating possible sensitivity to HER2-targeted tyrosine kinase inhibitors (TKIs) . Furthermore, preclinical studies have demonstrated that AP32788, a selective inhibitor of EGFR and HER2, effectively inhibits the growth of cell lines overexpressing ERBB2 G776V . Despite these findings, the functional consequences of the p.G776V mutation are not fully understood, and its role in tumorigenesis requires further investigation. Given the potential therapeutic implications, additional studies are warranted to elucidate the clinical significance of this variant and to explore targeted treatment options. | <https://pmkb.weill.cornell.edu/genes/95>  <https://pmc.ncbi.nlm.nih.gov/articles/PMC11252102/>  <https://www.accessdata.fda.gov/cdrh_docs/pdf16/P160045S035B.pdf>  <https://www.nature.com/articles/s41598-022-13189-y>  <https://www.frontiersin.org/journals/pharmacology/articles/10.3389/fphar.2022.806737/full>  <https://www.nature.com/articles/s41523-022-00390-5>  <https://www.sciencedirect.com/science/article/pii/S1535610821002841>  <https://acsjournals.onlinelibrary.wiley.com/doi/10.1002/cncr.31125>  <https://ascopubs.org/doi/pdf/10.1200/PO.16.00037>  <https://aacrjournals.org/cancerres/article/76/14_Supplement/2644/609601/Abstract-2644-AP32788-a-potent-selective-inhibitor>  <https://aacrjournals.org/cancerres/article/65/5/1642/519413/Somatic-Mutations-of-the-HER2-Kinase-Domain-in>  <https://www.mycancergenome.org/content/alteration/erbb2-exon-20-insertion/>  <https://pmc.ncbi.nlm.nih.gov/articles/PMC11247561/>  <https://pmc.ncbi.nlm.nih.gov/articles/PMC8530969/>  <https://www.sciencedirect.com/science/article/pii/S266700542100017X>  <https://www.sciencedirect.com/science/article/pii/S2667005421000363>  <https://www.oncokb.org/gene/ERBB2/G776V/Breast%20Cancer>  <https://www.oncokb.org/gene/ERBB2/G776V/Hepatobiliary%20Cancer>  <https://pmkb.weill.cornell.edu/genes/95>  <https://www.carislifesciences.com/wp-content/uploads/2022/06/ASCO-Caris-2022-HER2-Alterations-and-Prognostic-Implications-in-All-Subtypes-of-Breast-Cancer.pdf> |
|  | Evaluation  **Summarization Accuracy & Completeness:**  **Hallucination Rate:**  **Readability & Usability:**  **Literature Coverage & Relevance:**  **Time Efficiency:** | |
| **3** | The genomic variant c.2327G>T p.G776V rs144434331 on the HER2 gene is a missense mutation that results in the substitution of glycine with valine at the 776th amino acid position of the HER2 protein. This variant is located within the tyrosine kinase domain (TKD) of HER2, which is critical for the receptor's enzymatic activity and signal transduction. The HER2 gene, also known as ERBB2, encodes a receptor tyrosine kinase that belongs to the epidermal growth factor receptor (EGFR) family. HER2 plays a pivotal role in the regulation of cell growth, survival, and differentiation. Overexpression or amplification of HER2 is associated with various cancers, including breast and gastric cancers, and it is a target for cancer therapeutics.  The p.G776V variant has been identified in the context of cancer and has been evaluated for its transforming potential using the MANO method, which assesses the ability of HER2 variants to induce cellular transformation [3]. Although the specific functional consequences of the p.G776V variant are not detailed in the provided literature, other variants at the same position, such as p.G776S, have been studied, suggesting that alterations at this residue may impact the function of the HER2 protein.  In a study involving patients with lung adenocarcinomas, the p.G776V variant was identified in one patient, indicating its occurrence in cancer [3]. The response of this variant to HER2-targeted tyrosine kinase inhibitors (TKIs) is not explicitly described in the literature provided. However, it is noted that HER2 kinase domain missense mutations, including p.G776V, have been recorded in datasets such as Foundation Medicine or cBioPortal and are recognized as oncogenic [3].  In a case series, the p.G776V variant was detected in a patient's tumor, and while the variant was not found in circulating cell-free DNA (ccfDNA), this does not necessarily reflect the variant's clinical significance or its abundance in the tumor [7]. The variant's presence in the tumor tissue suggests it may contribute to the oncogenic process, although further studies would be required to fully elucidate its role.  Overall, the p.G776V variant is a rare missense mutation within the HER2 gene with potential oncogenic properties. Its clinical significance remains uncertain, as reflected by its classification in ClinVar, and further research is needed to determine its impact on HER2 function and its potential as a therapeutic target. | [[1]](https://pubmed.ncbi.nlm.nih.gov/38715247/) Bon G et al. (2024). "HER2 mutation as an emerging target in advanced breast cancer." Cancer science, 115(7)  [[2]](https://pubmed.ncbi.nlm.nih.gov/24956168/) Sherwood JL et al. (2014). "Panel based MALDI-TOF tumour profiling is a sensitive method for detecting mutations in clinical non small cell lung cancer tumour." PloS one, 9(6)  [[3]](https://pubmed.ncbi.nlm.nih.gov/39012378/) Yang G et al. (2024). "Clinical and structural insights into the rare but oncogenic HER2-activating missense mutations in non-small cell lung cancer: a retrospective ATLAS cohort study." Discover oncology, 15(1)  [[4]](https://pubmed.ncbi.nlm.nih.gov/38287892/) Corne J et al. (2024). "Plasma-based analysis of ERBB2 mutational status by multiplex digital PCR in a large series of patients with metastatic breast cancer." Molecular oncology, 18(11)  [[5]](https://pubmed.ncbi.nlm.nih.gov/30297788/) Pasternack H et al. (2018). "Somatic alterations in circulating cell-free DNA of oesophageal carcinoma patients during primary staging are indicative for post-surgical tumour recurrence." Scientific reports, 8(1)  [[6]](https://pubmed.ncbi.nlm.nih.gov/17088437/) Ikediobi ON et al. (2006). "Mutation analysis of 24 known cancer genes in the NCI-60 cell line set." Molecular cancer therapeutics, 5(11)  [[7]](https://pubmed.ncbi.nlm.nih.gov/37427135/) Xu T et al. (2023). "Sustained complete response to first-line immunochemotherapy for highly aggressive TP53/MDM2-mutated upper tract urothelial carcinoma with ERBB2 mutations, luminal immune-infiltrated contexture, and non-mesenchymal state: a case report and literature review." Frontiers in oncology, 13()  [[8]](https://pubmed.ncbi.nlm.nih.gov/30072744/) Sugimachi K et al. (2018). "Serial mutational tracking in surgically resected locally advanced colorectal cancer with neoadjuvant chemotherapy." British journal of cancer, 119(4)  [[9]](https://pubmed.ncbi.nlm.nih.gov/29975516/) Guo J et al. (2018). "Identification of a Water-Coordinating HER2 Inhibitor by Virtual Screening Using Similarity-Based Scoring." Biochemistry, 57(32)  [[10]](https://pubmed.ncbi.nlm.nih.gov/19459856/) Suzuki T et al. (2009). "Antitumor activity of a dual epidermal growth factor receptor and ErbB2 kinase inhibitor MP-412 (AV-412) in mouse xenograft models." Cancer science, 100(8)  [[11]](https://pubmed.ncbi.nlm.nih.gov/29967253/) Nagano M et al. (2018). "High-Throughput Functional Evaluation of Variants of Unknown Significance in ERBB2." Clinical cancer research : an official journal of the American Association for Cancer Research, 24(20)  [[12]](https://doi.org/10.54112/bcsrj.v2023i1.629) F. Muccee et al. (2023). "IN-SILICO CHARACTERIZATION OF HUMAN HER2 GENE TO PREDICT THE BREAST CANCER ASSOCIATED BIOMARKERS" Biological and Clinical Sciences Research Journal,  [[13]](https://pubmed.ncbi.nlm.nih.gov/28448556/) Inoue Y et al. (2017). "A highly specific and sensitive massive parallel sequencer-based test for somatic mutations in non-small cell lung cancer." PloS one, 12(4)  [[14]](https://pubmed.ncbi.nlm.nih.gov/38509475/) Randall J et al. (2024). "Quantitative proteomic analysis of HER2 protein expression in PDAC tumors." Clinical proteomics, 21(1)  [[15]](https://pubmed.ncbi.nlm.nih.gov/23525077/) Dulak AM et al. (2013). "Exome and whole-genome sequencing of esophageal adenocarcinoma identifies recurrent driver events and mutational complexity." Nature genetics, 45(5) |
|  | Evaluation  **Summarization Accuracy & Completeness:**  **Hallucination Rate:**  **Readability & Usability:**  **Literature Coverage & Relevance:**  **Time Efficiency:** | |
| **4** | The HER2 gene (ERBB2) encodes a member of the epidermal growth factor receptor (EGFR) family of receptor tyrosine kinases, playing a key role in the regulation of cell proliferation and survival through activation of downstream signaling cascades such as the PI3K/AKT and MAPK pathways. Aberrant activation of HER2 through amplification or somatic mutations contributes to oncogenesis in various cancers, particularly breast, gastric, and non-small cell lung cancer (NSCLC). The p.G776V variant is a rare missense mutation occurring within the kinase domain of HER2, involving a glycine-to-valine substitution at position 776. While structurally close to the more well-characterized activating mutation p.G776>VC (a duplication insertion), p.G776V itself has limited direct characterization in the literature. Available data from broader analyses of HER2 mutations in NSCLC suggest that alterations at codon 776 can result in constitutive kinase activity and may be oncogenic; however, most studies have not independently evaluated p.G776V, and the specific biochemical or clinical impact of this variant remains uncertain. A retrospective cohort analysis of HER2 mutations in NSCLC highlighted the structural heterogeneity and oncogenic potential of rare kinase domain variants, including those near G776, but did not provide functional analysis specific to p.G776V [1]. Similarly, studies on circulating tumor DNA profiling in metastatic settings and broader cancer genomic screens list variants at this codon without specific commentary on G776V’s pathogenicity [2][3]. In the absence of variant-specific evidence, it is currently not possible to classify p.G776V with confidence regarding oncogenicity or therapeutic response, particularly to HER2-targeted tyrosine kinase inhibitors (TKIs). Therefore, while the localization of the mutation in a hotspot region of the HER2 kinase domain raises concern for possible functional relevance, its clinical significance remains uncertain. Additional in vitro functional assays and clinical correlation studies are needed to elucidate the precise role of this variant. At present, p.G776V should be classified as a variant of uncertain significance (VUS), and caution should be used when considering it for treatment decisions. | [1] [Yang et al.](https://link.springer.com/article/10.1007/s12672-024-01154-2)  [2] [Corné et al.](https://febs.onlinelibrary.wiley.com/doi/abs/10.1002/1878-0261.13592)  [3] [Ikediobi et al.](https://aacrjournals.org/mct/article-abstract/5/11/2606/234860) |
|  | Evaluation  **Summarization Accuracy & Completeness:**  **Hallucination Rate:**  **Readability & Usability:**  **Literature Coverage & Relevance:**  **Time Efficiency:** | |
| **5** | The HER2 gene, also known as ERBB2, encodes a transmembrane tyrosine kinase receptor belonging to the epidermal growth factor receptor (EGFR) family. This receptor plays a crucial role in regulating cell growth, survival, and differentiation through the activation of downstream signaling pathways, including the PI3K/Akt and MAPK pathways. Amplification or mutation of HER2 can lead to constitutive activation of these pathways, driving oncogenesis and tumor progression. The HER2 gene is notably implicated in various cancers, including breast, gastric, and non-small cell lung cancer (NSCLC). The p.G776V variant of the HER2 gene is a missense mutation that results in the substitution of glycine with valine at amino acid position 776. This variant is present in approximately 0.03% of AACR GENIE cases, with the highest prevalence observed in colon adenocarcinoma, bladder urothelial carcinoma, ampulla of Vater carcinoma, ampulla of Vater pancreatobiliary type adenocarcinoma, and appendix mucinous adenocarcinoma. The ERBB2 G776V mutation is a predictive biomarker for the use of ado-trastuzumab emtansine and fam-trastuzumab deruxtecan in patients, with two therapies having NCCN guidelines in at least one clinical setting. Non-small cell lung carcinoma has the most therapies targeted against ERBB2 G776V or its related pathways. The clinical significance of the p.G776V variant is not fully elucidated, and further investigation is needed to determine its impact on protein function and response to targeted therapies. The absence of direct references to the p.G776V variant in peer-reviewed literature highlights the need for additional research to clarify its role in cancer development and treatment. | No references available |
|  | Evaluation  **Summarization Accuracy & Completeness:**  **Hallucination Rate:**  **Readability & Usability:**  **Literature Coverage & Relevance:**  **Time Efficiency:** | |
