## Supplementary material for "Benchmarking generative AI tools for literature retrieval and summarization in genomic variant interpretation": IDH1 p.R132H.docx

| **Tool** | **Summary** | **References** |
| --- | --- | --- |
| **1** | The genomic variant c.395G>A p.R132H rs121913500 in the IDH1 gene is a well-characterized missense mutation that results in the substitution of arginine with histidine at codon 132 of the isocitrate dehydrogenase 1 (IDH1) enzyme. IDH1 is an enzyme that normally functions in the cytosol to catalyze the oxidative carboxylation of isocitrate to alpha-ketoglutarate (a-KG), while concurrently reducing NADP+ to NADPH [1]. This reaction is a part of the tricarboxylic acid (TCA) cycle, which is crucial for cellular metabolism.  The p.R132H variant imparts a gain-of-function activity upon IDH1, enabling it to convert a-KG into the oncometabolite D-2-hydroxyglutarate (D-2HG) [2]. This neomorphic activity leads to the accumulation of D-2HG, which competitively inhibits a-KG-dependent enzymes, including those involved in histone and DNA demethylation. The resulting epigenetic dysregulation can drive tumorigenesis through alterations in gene expression and DNA damage pathways [2][5].  Clinically, the IDH1 p.R132H mutation is a critical prognostic marker in adult-type diffuse gliomas, with over 90% of IDH1/2 mutant gliomas harboring this alteration [1]. It is associated with significantly longer overall and progression-free survival compared to gliomas without IDH1/2 mutations [1]. The presence of this mutation also has therapeutic implications, as it can influence treatment decisions, particularly with the emergence of IDH inhibitors that show promise in slowing disease progression [1].  Diagnostic testing for the IDH1 p.R132H mutation typically involves immunohistochemistry (IHC) to detect the mutant protein, followed by next-generation sequencing (NGS) to confirm the mutation and identify other potential alterations in IDH1 and IDH2 [1]. The IDH1 R132H variant has been reported in ClinVar with multiple pathogenic submissions, underscoring its clinical significance.  In summary, the IDH1 c.395G>A p.R132H rs121913500 variant is a pathogenic mutation that alters the enzymatic function of IDH1, leading to the production of D-2HG and subsequent epigenetic changes that contribute to the development and progression of certain cancers, particularly gliomas. Its detection is important for prognosis and guiding treatment strategies. | [[1]](https://pubmed.ncbi.nlm.nih.gov/38796421/) Solomon JP et al. (2024). "Evaluation of the rapid Idylla IDH1-2 mutation assay in FFPE glioma samples." Diagnostic pathology, 19(1)  [[2]](https://pubmed.ncbi.nlm.nih.gov/38712107/) Ahmed Adam MA et al. (2024). "Catalytically distinct IDH1 mutants tune phenotype severity in tumor models." bioRxiv : the preprint server for biology, ()  [[3]](https://pubmed.ncbi.nlm.nih.gov/38260668/) Mealka M et al. (2024). "Active site remodeling in tumor-relevant IDH1 mutants drives distinct kinetic features and potential resistance mechanisms." bioRxiv : the preprint server for biology, ()  [[4]](https://pubmed.ncbi.nlm.nih.gov/38464189/) Mealka M et al. (2024). "Active site remodeling in tumor-relevant IDH1 mutants drives distinct kinetic features and potential resistance mechanisms." Research square, ()  [[5]](https://pubmed.ncbi.nlm.nih.gov/39306640/) Lai K et al. (2025). "The IDH1-R132H mutation aggravates cisplatin-induced acute kidney injury by promoting ferroptosis through disrupting NDUFA1 and FSP1 interaction." Cell death and differentiation, 32(2)  [[6]](https://pubmed.ncbi.nlm.nih.gov/38248076/) Varachev V et al. (2024). "Diagnostics of IDH1/2 Mutations in Intracranial Chondroid Tumors: Comparison of Molecular Genetic Methods and Immunohistochemistry." Diagnostics (Basel, Switzerland), 14(2)  [[7]](https://pubmed.ncbi.nlm.nih.gov/38172718/) Crucitta S et al. (2024). "IDH1 mutation is detectable in plasma cell-free DNA and is associated with survival outcome in glioma patients." BMC cancer, 24(1)  [[8]](https://pubmed.ncbi.nlm.nih.gov/38285799/) Khairy RA et al. (2024). "Evaluation of Immunohistochemical Expression of ALK-1 in Gliomas, WHO Grade 4 and Its Correlation with IDH1-R132H Mutation Status." Asian Pacific journal of cancer prevention : APJCP, 25(1)  [[9]](https://pubmed.ncbi.nlm.nih.gov/38286053/) Tang T et al. (2024). "IDH1 mutation predicts seizure occurrence and prognosis in lower-grade glioma adults." Pathology, research and practice, 254()  [[10]](https://pubmed.ncbi.nlm.nih.gov/38896992/) Xu T et al. (2024). "Discovery of novel natural-product-derived mutant isocitrate dehydrogenases 1 inhibitors: Structure-based virtual screening, biological evaluation and structure-activity relationship study." European journal of medicinal chemistry, 275()  [[11]](https://pubmed.ncbi.nlm.nih.gov/39596246/) Shmelev ME et al. (2024). "IDH1 R132H and TP53 R248Q Mutations Modulate Glioma Cell Migration and Adhesion on Different ECM Components." International journal of molecular sciences, 25(22)  [[12]](https://pubmed.ncbi.nlm.nih.gov/38334611/) Sprinzen L et al. (2024). "EZH2 Inhibition Sensitizes IDH1R132H-Mutant Gliomas to Histone Deacetylase Inhibitor." Cells, 13(3)  [[13]](https://pubmed.ncbi.nlm.nih.gov/38633919/) Squalli Houssaini A et al. (2024). "Prognostic Value of ATRX and p53 Status in High-Grade Glioma Patients in Morocco." Cureus, 16(3)  [[14]](https://pubmed.ncbi.nlm.nih.gov/38854127/) Murphy ZR et al. (2024). "Ultra-Rapid Droplet Digital PCR Enables Intraoperative Tumor Quantification." medRxiv : the preprint server for health sciences, ()  [[15]](https://pubmed.ncbi.nlm.nih.gov/39605316/) Weber R et al. (2024). "Targeting the IDH1 (R132H) mutation in gliomas by CRISPR/Cas precision base editing." Neuro-oncology advances, 6(1) |
|  | Evaluation  **Summarization Accuracy & Completeness:**  **Hallucination Rate:**  **Readability & Usability:**  **Literature Coverage & Relevance:**  **Time Efficiency:** | |
| **2** | The IDH1 gene encodes isocitrate dehydrogenase 1, an enzyme that catalyzes the oxidative decarboxylation of isocitrate to α-ketoglutarate (α-KG) in the cytosol, contributing to NADPH production and cellular redox balance. The p.R132H variant (c.395G>A) is a heterozygous missense mutation substituting arginine with histidine at codon 132, a critical residue in the enzyme's active site. This mutation impairs normal catalytic activity and confers a neomorphic function, leading to the production of the oncometabolite D-2-hydroxyglutarate (D-2HG), which inhibits α-KG–dependent dioxygenases, resulting in widespread epigenetic alterations and contributing to gliomagenesis . The p.R132H mutation accounts for over 90% of IDH1 mutations in gliomas and is predominantly found in WHO grade II and III diffuse gliomas, as well as secondary glioblastomas . Clinically, IDH1 mutations, including p.R132H, are associated with a better prognosis and longer overall survival compared to IDH1 wild-type gliomas . Immunohistochemistry using mutation-specific antibodies allows for the detection of the p.R132H variant, aiding in diagnosis and prognostication . Functionally, the p.R132H mutation leads to decreased NADPH and glutathione levels, sensitizing tumor cells to oxidative stress and certain chemotherapeutic agents . Additionally, it has been shown to activate the AKT-mTOR signaling pathway, enhancing cell migration . Despite these oncogenic properties, the presence of the p.R132H mutation correlates with a more favorable clinical outcome, underscoring its complex role in glioma biology. | <https://pmc.ncbi.nlm.nih.gov/articles/PMC4109985/>  <https://www.mdpi.com/2079-7737/13/11/885>  <https://pmc.ncbi.nlm.nih.gov/articles/PMC3227941/>  <https://www.oncotarget.com/article/8918/text/>  <https://journals.plos.org/plosone/article?id=10.1371%2Fjournal.pone.0169038>  <https://academic.oup.com/noa/article/doi/10.1093/noajnl/vdae182/7866812>  <https://www.nature.com/articles/s41467-024-51332-7>  <https://academic.oup.com/noa/article/5/1/vdad069/7185176>  <https://link.springer.com/article/10.1007/s11060-023-04250-5>  <https://www.nature.com/articles/s41586-021-03363-z>  <https://www.nature.com/articles/s41467-020-20752-6>  <https://www.frontiersin.org/journals/oncology/articles/10.3389/fonc.2019.01507/full>  <https://academic.oup.com/carcin/article/40/11/1299/5559593>  <https://www.oncotarget.com/article/8918/text/>  <https://pmc.ncbi.nlm.nih.gov/articles/PMC3227941/>  <https://pmc.ncbi.nlm.nih.gov/articles/PMC4109985/>  <https://pmc.ncbi.nlm.nih.gov/articles/PMC11071412/>  <https://www.sciencedirect.com/science/article/pii/S0893395222022372>  <https://www.sciencedirect.com/science/article/abs/pii/S1092913411001754>  <https://academic.oup.com/jes/article/4/4/bvaa018/5739627>  <https://ihc.testcatalog.org/show/IDH1>  <https://journals.plos.org/plosone/article?id=10.1371%2Fjournal.pone.0169038>  <https://www.cap.org/member-resources/articles/updates-in-glioma-and-meningioma-classification-who-2021-new-entities-in-the-literature>  <https://www.cap.org/member-resources/articles/updates-in-glioma-and-meningioma-classification-who-2021-new-entities-in-the-literature>  <https://www.mdpi.com/2079-7737/13/11/885>  <https://jcp.bmj.com/content/73/10/611> |
|  | Evaluation  **Summarization Accuracy & Completeness:**  **Hallucination Rate:**  **Readability & Usability:**  **Literature Coverage & Relevance:**  **Time Efficiency:** | |
| **3** | The IDH1 gene encodes isocitrate dehydrogenase 1, a metabolic enzyme that plays a crucial role in cellular metabolism by catalyzing the conversion of isocitrate to alpha-ketoglutarate (α-KG) in the cytoplasm.  This process is essential for generating NADPH, which is vital for various cellular functions, including lipid synthesis and redox regulation.  The p.R132H variant of the IDH1 gene is a well-documented gain-of-function mutation that significantly alters the enzyme's activity.  This mutation results in the substitution of arginine with histidine at codon 132, leading to the production of the oncometabolite D-2-hydroxyglutarate (D2HG) from α-KG.  D2HG acts as a competitive inhibitor of α-KG-dependent dioxygenases, including histone demethylases and the DNA demethylase TET2, leading to a hypermethylation phenotype.  This epigenetic alteration is associated with a block in cellular differentiation and an enrichment of neural progenitor cell expression profiles in gliomas.  The p.R132H mutation is frequently observed in low-grade gliomas and secondary high-grade glioblastomas, occurring in approximately 70% of astrocytomas and oligodendroglial tumors.  It is the most common IDH1 mutation, accounting for over 90% of all IDH1 mutations in gliomas.  The presence of the p.R132H mutation is strongly correlated with better prognosis and increased survival rates in glioma patients, likely due to its role in promoting a less aggressive tumor phenotype.  This mutation is also associated with a higher rate of responses to up-front temozolomide and genotoxic therapy in low-grade glioma patients, suggesting potential predictive value.  The p.R132H mutation can be detected using immunohistochemistry with a specific antibody (clone H09), which is highly sensitive and specific for the mutated protein.  This method is recommended as an initial screening test for IDH1 mutations in gliomas, followed by DNA sequencing if the immunohistochemistry results are negative.  The frequency of the p.R132H mutation in gliomas highlights its clinical significance, and its impact on tumor biology and patient outcomes underscores the importance of routine testing for this variant in diagnostic settings. | [1] Hartmann C, Meyer J, Balss J, Capper D, Mueller W, Christians A et al (2009) Type and frequency of IDH1 and IDH2 mutations are related to astrocytic and oligodendroglial differentiation and age: a study of 1010 diffuse gliomas. Acta Neuropathol. 2009 Jun 25.  [2] Preusser M, Capper D, Hartmann C. IDH mutations as prognostic and predictive markers in gliomas. Clin Neuropathol. 2011;30(5):217-30. |
|  | Evaluation  **Summarization Accuracy & Completeness:**  **Hallucination Rate:**  **Readability & Usability:**  **Literature Coverage & Relevance:**  **Time Efficiency:** | |
| **4** | The IDH1 gene encodes isocitrate dehydrogenase 1, an enzyme that catalyzes the oxidative decarboxylation of isocitrate to α-ketoglutarate, playing a pivotal role in the citric acid cycle and cellular metabolism. The missense mutation p.R132H (arginine to histidine substitution at codon 132) represents the most frequent pathogenic alteration in IDH1 and is strongly implicated in the pathogenesis of gliomas, particularly lower-grade gliomas and secondary glioblastomas. This somatic mutation occurs at a critical site in the active center of the enzyme and results in a neomorphic enzymatic function, wherein α-ketoglutarate is reduced to the oncometabolite D-2-hydroxyglutarate (2-HG), leading to widespread epigenetic dysregulation and impaired cellular differentiation. The p.R132H variant has been identified in up to 70–80% of grade II and III diffuse gliomas and secondary glioblastomas, yet it is rarely observed in primary glioblastomas or other solid tumors [1,2]. Diagnostic assays targeting this variant are clinically validated and widely used for molecular classification of gliomas, given its prognostic value and association with better overall survival compared to IDH1-wildtype tumors [3]. Despite its prevalence in gliomas, p.R132H is largely absent in other malignancies, highlighting its potential as a tumor-specific biomarker [1]. Therapeutic efforts are ongoing to target mutant IDH1, with IDH1 inhibitors being evaluated in clinical trials. Notably, studies have demonstrated that the p.R132H mutation co-segregates with other molecular features such as 1p/19q non-codeletion and ATRX mutation, helping refine glioma subclassification [4]. Nonetheless, despite the wealth of data surrounding its biological and clinical impact, questions remain regarding the timing of IDH1 mutation acquisition during gliomagenesis and its precise role in tumor evolution, suggesting a continued need for research into its mechanistic contributions and therapeutic targeting. | [1] Bleeker et al. *IDH1 mutations at residue p.R132 (IDH1R132) occur frequently in high‐grade gliomas but not in other solid tumors* (2009). [Link](https://app.scholarai.io/paper?paper_id=DOI:10.1002/humu.20937&original_url=https%3A%2F%2Fonlinelibrary.wiley.com%2Fdoi%2Fabs%2F10.1002%2Fhumu.20937)  [2] Kang et al. *Mutational analysis of IDH1 codon 132 in glioblastomas and other common cancers* (2009). [Link](https://app.scholarai.io/paper?paper_id=DOI:10.1002/ijc.24379&original_url=https%3A%2F%2Fonlinelibrary.wiley.com%2Fdoi%2Fabs%2F10.1002%2Fijc.24379)  [3] Kurian et al. *IDH mutation analysis in gliomas as a diagnostic and prognostic biomarker* (2013). [Link](https://app.scholarai.io/paper?paper_id=DOI:10.3109/02688697.2013.771139&original_url=https%3A%2F%2Fwww.tandfonline.com%2Fdoi%2Fabs%2F10.3109%2F02688697.2013.771139)  [4] Gravendeel et al. *Segregation of non‐p.R132H mutations in IDH1 in distinct molecular subtypes of glioma* (2010). [Link](https://app.scholarai.io/paper?paper_id=DOI:10.1002/humu.21201&original_url=https%3A%2F%2Fonlinelibrary.wiley.com%2Fdoi%2Fabs%2F10.1002%2Fhumu.21201) |
|  | Evaluation  **Summarization Accuracy & Completeness:**  **Hallucination Rate:**  **Readability & Usability:**  **Literature Coverage & Relevance:**  **Time Efficiency:** | |
| **5** | The p.R132H variant in the IDH1 gene is a well-characterized gain-of-function mutation that results in the substitution of arginine with histidine at codon 132, a critical residue within the enzyme's active site. IDH1 encodes an NADP+-dependent isocitrate dehydrogenase that plays a vital role in cellular metabolism by catalyzing the conversion of isocitrate to α-ketoglutarate (α-KG) while producing NADPH, which is essential for reductive biosynthesis and maintaining cellular redox balance. The p.R132H mutation leads to neomorphic enzymatic activity, causing the abnormal reduction of α-KG to the oncometabolite R-2-hydroxyglutarate (R-2-HG), which accumulates within tumor cells and disrupts normal epigenetic regulation by inhibiting α-KG-dependent dioxygenases. This disruption promotes oncogenic transformation, particularly in gliomas, where the mutation is prevalent, especially in low-grade gliomas and secondary glioblastomas, accounting for over 90% of IDH1 mutations in these tumors. The mutation is classified as pathogenic and is a key molecular marker used in glioma classification, with its presence associated with distinct clinical outcomes, often indicating a better prognosis compared to wild-type IDH1 tumors. The mutation's frequency varies but is notably high in specific tumor types, and it is rarely found in the general population, suggesting it is a somatic mutation acquired during tumorigenesis. Detection methods include immunohistochemistry and sequencing, with the R132H mutation being the most common, and specific assays like T-ARMS PCR have been developed for sensitive and cost-effective genotyping. The mutation's impact extends beyond metabolic reprogramming to influence tumor growth, resistance mechanisms, and potential therapeutic targets, as studies have shown that IDH1-mutant cells depend on pathways such as the pentose phosphate pathway for NADPH production, which supports their survival and proliferation. Despite extensive research, uncertainties remain regarding the complete spectrum of biological effects and optimal targeted therapies, emphasizing the need for further investigation into the mutation's role in tumor biology and treatment resistance. The mutation's significance in prognosis, diagnosis, and therapy underscores its importance in clinical and research settings. The mutation is referenced in multiple peer-reviewed studies, including its role in glioma prognosis, metabolic reprogramming, and targeted therapy development[1][3][4][5][6][8]. | [1] <https://www.semanticscholar.org/paper/a709a573293e3a2e14ab7b8ccd40135b7e053061>  [2] <https://www.semanticscholar.org/paper/948dd8329e1a768f4acc010b062d88a1662b88b0>  [3] <https://www.semanticscholar.org/paper/868dbc4e8ba10d5953a6235cfcea0b850875fcd9>  [4] <https://pubmed.ncbi.nlm.nih.gov/28735489/>  [5] <https://www.ncbi.nlm.nih.gov/pmc/articles/PMC9305137/>  [6] <https://www.semanticscholar.org/paper/44cbb5dae0721857a9bb28da211cbc7863eee03e>  [7] <https://pubmed.ncbi.nlm.nih.gov/39039166/>  [8] <https://www.semanticscholar.org/paper/9ab95902c3b4d1179ff7776ae8a8fd709bf2ad69> |
|  | Evaluation  **Summarization Accuracy & Completeness:**  **Hallucination Rate:**  **Readability & Usability:**  **Literature Coverage & Relevance:**  **Time Efficiency:** | |
