## Supplementary material for "Benchmarking generative AI tools for literature retrieval and summarization in genomic variant interpretation": PIK3CA p.E545K.docx

| **Tool** | **Summary** | **References** |
| --- | --- | --- |
| **1** | The PIK3CA gene encodes the p110α catalytic subunit of phosphoinositide 3-kinase (PI3K), a critical enzyme involved in the PI3K/AKT/mTOR signaling pathway that regulates cell growth, proliferation, survival, and metabolism. Mutations in PIK3CA, particularly gain-of-function variants, lead to constitutive activation of this pathway, contributing to oncogenesis and vascular malformations. The p.E545K variant, classified as a hotspot mutation, results from a glutamic acid to lysine substitution at amino acid position 545 within the helical domain of p110α, which enhances kinase activity and promotes oncogenic processes. This mutation is frequently identified in various cancers, including breast, colorectal, and uterine carcinomas, with studies reporting its presence in approximately 10-20% of breast tumors and other malignancies, often correlating with aggressive disease phenotypes. The p.E545K mutation has been associated with increased cell proliferation, angiogenesis, and resistance to targeted therapies, making it a significant biomarker for prognosis and therapeutic stratification. Recent research indicates that this mutation can generate neoantigens that are targetable by T-cell receptor (TCR) gene therapy, highlighting its potential in immunotherapeutic approaches. Despite its established role as a driver mutation, the precise impact of p.E545K on protein structure and function has been elucidated through molecular dynamics simulations, confirming its contribution to increased kinase activity and oncogenic signaling. The mutation's prevalence and functional consequences underscore its importance in both cancer biology and vascular malformations such as capillary lymphatic venous malformations, where somatic PIK3CA mutations are implicated in pathogenesis. Currently, targeted inhibitors of the PI3K/AKT/mTOR pathway are under investigation for treating PIK3CA-mutant tumors, although resistance mechanisms necessitate ongoing research. The classification of p.E545K as a pathogenic, gain-of-function mutation is supported by its recurrent detection in tumor samples and functional studies demonstrating enhanced kinase activity and oncogenic potential. Further research is warranted to clarify its full spectrum of biological effects and to optimize targeted therapeutic strategies for patients harboring this mutation. Overall, p.E545K in PIK3CA represents a critical molecular alteration with significant implications for cancer progression, treatment resistance, and emerging immunotherapies. | [1] <https://www.semanticscholar.org/paper/f99479b56055975675e7f0f6d9f4bff18e5ec57f>  [2] <https://pubmed.ncbi.nlm.nih.gov/27511117/>  [3] <https://pubmed.ncbi.nlm.nih.gov/33428122/>  [4] <https://pubmed.ncbi.nlm.nih.gov/25679319/>  [5] <https://pubmed.ncbi.nlm.nih.gov/35484265/>  [6] <https://pubmed.ncbi.nlm.nih.gov/24908061/>  [7] <https://www.semanticscholar.org/paper/12fc89872151ad751c255e4c66ae8158b002af3f>  [8] <https://www.semanticscholar.org/paper/016918919c8e63804eb013266fa6bbc0a7087345> |
|  | Evaluation  **Summarization Accuracy & Completeness:**  **Hallucination Rate:**  **Readability & Usability:**  **Literature Coverage & Relevance:**  **Time Efficiency:** | |
| **2** | The genomic variant c.1633G>A, resulting in the amino acid substitution p.E545K (rs104886003), is located in the PIK3CA gene, which encodes the p110α catalytic subunit of phosphoinositide 3-kinase (PI3K). PI3K plays a pivotal role in cellular processes such as growth, proliferation, and survival by synthesizing phosphatidylinositol-3,4,5-triphosphate (PIP3), which activates several downstream signaling pathways, including AKT/mTOR, VEGFR, and EGFR [2].  The p.E545K variant in PIK3CA is a hotspot mutation that has been identified as oncogenic, with a high frequency of occurrence in various cancers, including colorectal cancer (CRC). This missense mutation leads to a charge reversal at the amino acid position 545, which disrupts the interaction between the protein's catalytic and regulatory subunits, resulting in constitutive activation of the PIK3CA alpha subunit and oncogenic signaling [2].  In CRC patients, the p.E545K variant has been associated with poor/moderate tumor differentiation and advanced TNM stage, indicating a correlation with more aggressive disease characteristics. Kaplan-Meier analysis has demonstrated a worse prognosis in overall survival for patients harboring this mutation, and multivariate analysis has confirmed the p.E545K variant as a detrimental factor for overall survival, with a hazard ratio of 6.497 [2].  Furthermore, the presence of the p.E545K variant has been shown to confer higher sensitivity to PI3K pathway inhibitors, such as p110α or AKT inhibitors, suggesting potential therapeutic implications for targeted treatment strategies in cancers with this mutation [3].  Given the significant clinical impact of the p.E545K variant on prognosis and treatment response, it is considered a potential molecular biomarker for predicting prognosis in CRC patients. The variant is well-documented in ClinVar with numerous pathogenic submissions, highlighting its clinical relevance. | [[1]](https://pubmed.ncbi.nlm.nih.gov/39401325/) Yasin F et al. (2024). "Molecular characteristics of advanced colorectal cancer and multi-hit PIK3CA mutations." The oncologist, 29(12)  [[2]](https://pubmed.ncbi.nlm.nih.gov/32099598/) Ranjbar R et al. (2019). "Prevalence and prognostic role of PIK3CA E545K mutation in Iranian colorectal cancer patients." Gastroenterology and hepatology from bed to bench, 12(Suppl1)  [[3]](https://pubmed.ncbi.nlm.nih.gov/39543937/) Zhang S et al. (2025). "Vertical inhibition of p110alpha/AKT and N-cadherin enhances treatment efficacy in PIK3CA-aberrated ovarian cancer cells." Molecular oncology, 19(4)  [[4]](https://pubmed.ncbi.nlm.nih.gov/38215117/) Chen J et al. (2024). "Whether specific genetic feature predicted immunotherapy efficacy: A case report." Medicine, 103(2)  [[5]](https://pubmed.ncbi.nlm.nih.gov/38260414/) Miranda AX et al. (2024). "Genomic dissection and mutation-specific target discovery for breast cancer PIK3CA hotspot mutations." bioRxiv : the preprint server for biology, ()  [[6]](https://pubmed.ncbi.nlm.nih.gov/38848145/) Liu YL et al. (2024). "APOA5 alleviates reactive oxygen species to promote oxaliplatin resistance in PIK3CA-mutated colorectal cancer." Aging, 16(11)  [[7]](https://pubmed.ncbi.nlm.nih.gov/38943378/) Debouki-Joudi S et al. (2024). "A novel PIK3CA hot-spot mutation in breast cancer patients detected by HRM-COLD-PCR analysis." Breast disease, 43(1)  [[8]](https://pubmed.ncbi.nlm.nih.gov/39444377/) Nieto-Coronel T et al. (2024). "PI3K Mutation Profiles on Exons 9 (E545K and E542K) and 20 (H1047R) in Mexican Patients With HER-2 Overexpressed Breast Cancer and Its Relevance on Clinical-Pathological and Survival Biological Effects." International journal of breast cancer, 2024()  [[9]](https://pubmed.ncbi.nlm.nih.gov/38802751/) Miranda AX et al. (2024). "Genomic dissection and mutation-specific target discovery for breast cancer PIK3CA hotspot mutations." BMC genomics, 25(1)  [[10]](https://pubmed.ncbi.nlm.nih.gov/39404875/) Wei Y et al. (2024). "Ultrasensitive detection of circulating tumor DNA using a CRISPR/Cas9 nickase-driven 3D DNA walker based on a COF-AuNPs sensing platform." Mikrochimica acta, 191(11)  [[11]](https://pubmed.ncbi.nlm.nih.gov/39711963/) Smilkou S et al. (2024). "Direct comparison of an ultrasensitive real-time PCR assay with droplet digital PCR for the detection of PIK3CA hotspot mutations in primary tumors, plasma cell-free DNA and paired CTC-derived gDNAs." Frontiers in oncology, 14()  [[12]](https://pubmed.ncbi.nlm.nih.gov/39890189/) Kim HG et al. (2025). "Clinicopathological and Molecular Insights into Primary Retroperitoneal Squamous Cell Carcinoma: HPV Association and Discovery of Recurrent PIK3CA E545K Mutation." Anticancer research, 45(2)  [[13]](https://pubmed.ncbi.nlm.nih.gov/38320429/) Martens GA et al. (2024). "Rational thresholding of circulating tumor DNA concentration for improved surveillance of metastatic breast cancer." ESMO open, 9(2)  [[14]](https://pubmed.ncbi.nlm.nih.gov/38616230/) Zheng S et al. (2024). "Understanding PI3K/Akt/mTOR signaling in squamous cell carcinoma: mutated PIK3CA as an example." Molecular biomedicine, 5(1)  [[15]](https://pubmed.ncbi.nlm.nih.gov/31666350/) Jiang W et al. (2020). "Targeting of beta-Catenin Reverses Radioresistance of Cervical Cancer with the PIK3CA-E545K Mutation." Molecular cancer therapeutics, 19(2) |
|  | Evaluation  **Summarization Accuracy & Completeness:**  **Hallucination Rate:**  **Readability & Usability:**  **Literature Coverage & Relevance:**  **Time Efficiency:** | |
| **3** | The PIK3CA gene encodes the p110α catalytic subunit of phosphatidylinositol 3-kinase (PI3K), a critical enzyme in the PI3K/AKT/mTOR signaling pathway that regulates cellular growth, metabolism, and survival. Somatic mutations in PIK3CA are among the most common oncogenic alterations across a range of human cancers, with the p.E545K variant—resulting from a substitution of glutamic acid to lysine at codon 545—classified as a gain-of-function, hotspot mutation. This variant occurs within the helical domain of the protein and disrupts inhibitory interactions with the regulatory subunit p85, thereby leading to constitutive PI3K activation and downstream oncogenic signaling. The E545K variant is recurrently observed in breast, colorectal, endometrial, cervical, and gallbladder carcinomas, and has been associated with tumor progression, increased glycolysis, and therapeutic resistance in several contexts. Mechanistic studies, including molecular dynamics simulations, have demonstrated that E545K stabilizes an active conformation of PI3Kα that promotes persistent membrane binding and enzymatic activity, mimicking growth factor stimulation in the absence of upstream signals [1]. In gallbladder carcinoma, this variant enhances binding to EGFR, augmenting tumorigenicity [2], and in cervical cancer, it contributes to proliferation via the β-catenin/SIRT3 pathway [3]. Functional assays confirm its oncogenic potential both in vitro and in murine tumor models [4], and its clinical relevance is underscored by the development and FDA approval of PI3Kα-selective inhibitors such as alpelisib for patients with PIK3CA-mutated, HR+/HER2– breast cancer [5]. Prevalence of the E545K variant varies by tumor type but has been frequently detected among the top three PIK3CA hotspot mutations alongside E542K and H1047R [6]. Despite significant research, the prognostic value of E545K remains context-dependent and variable between cancer types. The variant’s presence may influence therapeutic response to targeted agents or endocrine therapy, particularly in breast cancer, though prospective validation remains limited. Therefore, while p.E545K is a well-established oncogenic driver with functional and therapeutic implications, further investigation is warranted to delineate its role in resistance mechanisms and to refine its clinical utility in precision oncology. | [1] [Leontiadou et al., 2018](https://www.nature.com/articles/s41598-018-27044-6)  [2] [Zhao et al., 2016](https://link.springer.com/article/10.1186/s13046-016-0370-7)  [3] [Jiang et al., 2018](https://link.springer.com/article/10.1186/s13045-018-0674-5)  [4] Bader et al., 2006  [5] [Martínez-Sáez et al., 2020](https://link.springer.com/article/10.1186/S13058-020-01284-9)  [6] [Karakas et al., 2006](https://www.nature.com/articles/6602970) |
|  | Evaluation  **Summarization Accuracy & Completeness:**  **Hallucination Rate:**  **Readability & Usability:**  **Literature Coverage & Relevance:**  **Time Efficiency:** | |
| **4** | The PIK3CA gene encodes the p110α catalytic subunit of phosphatidylinositol 3-kinase (PI3K), a key component of the PI3K/AKT/mTOR signaling pathway that regulates cell proliferation, survival, and metabolism. The c.1633G>A (p.E545K) variant is a well-characterized somatic hotspot mutation located in exon 9 within the helical domain of PIK3CA. This substitution disrupts the inhibitory interaction between p110α and its regulatory subunit p85, resulting in constitutive activation of PI3K signaling. The E545K mutation is among the most prevalent PIK3CA alterations across various cancers. In breast cancer, it accounts for approximately 17% of all PIK3CA mutations , while in colorectal cancer, it represents about 9.2% of PIK3CA mutations . In non-small cell lung cancer (NSCLC), E545K constitutes 57.1% of PIK3CA mutations . Functionally, E545K enhances oncogenic signaling, promoting tumorigenesis and progression. In vivo studies have demonstrated that cells expressing E545K exhibit increased tumorigenic potential . Clinically, the presence of E545K has been associated with poor prognosis in certain cancers, including colorectal and gallbladder carcinoma . Moreover, E545K may confer resistance to standard therapies; for instance, cervical cancer cells harboring this mutation show reduced sensitivity to cisplatin and radiation . However, preclinical studies suggest that PI3K inhibitors, such as alpelisib, can effectively target tumors with E545K mutations . Given its pathogenic nature and therapeutic implications, the E545K variant is classified as pathogenic and warrants consideration in clinical decision-making. | [https://www.nature.com/articles/s41598-020-74086-w](https://www.nature.com/articles/s41598-020-74086-w?utm_source=chatgpt.com)  <https://aacrjournals.org/clincancerres/article/30/17/3868/747287/An-Analysis-of-PIK3CA-Hotspot-Mutations-and>  [https://breast-cancer-research.biomedcentral.com/articles/10.1186/s13058-020-01284-9](https://breast-cancer-research.biomedcentral.com/articles/10.1186/s13058-020-01284-9?utm_source=chatgpt.com)  <https://www.frontiersin.org/journals/pharmacology/articles/10.3389/fphar.2024.1494802/full>  [https://www.sciencedirect.com/science/article/pii/S1043661824000677](https://www.sciencedirect.com/science/article/pii/S1043661824000677?utm_source=chatgpt.com)  https://www.pnas.org/doi/10.1073/pnas.0510857103  <https://pmc.ncbi.nlm.nih.gov/articles/PMC7011074/>  <https://jeccr.biomedcentral.com/articles/10.1186/s13046-016-0370-7>  <https://pmc.ncbi.nlm.nih.gov/articles/PMC5347702/>  <https://ckb.jax.org/geneVariant/show?geneVariantId=2569>  <https://pmc.ncbi.nlm.nih.gov/articles/PMC11402363/>  <https://pmc.ncbi.nlm.nih.gov/articles/PMC11641866/>  <https://www.nature.com/articles/s41598-023-31593-w>  <https://www.nature.com/articles/s41467-023-35789-6>  <https://www.e-crt.org/journal/view.php?number=3382>  <https://www.frontiersin.org/journals/oncology/articles/10.3389/fonc.2021.644737/full>  <https://www.nature.com/articles/s41598-020-74086-w>  <https://breast-cancer-research.biomedcentral.com/articles/10.1186/s13058-020-01284-9>  <https://www.dovepress.com/the-distinct-clinicopathological-and-prognostic-implications-of-pik3ca-peer-reviewed-fulltext-article-CMAR>  <https://www.nature.com/articles/s41598-018-27044-6>  <https://jeccr.biomedcentral.com/articles/10.1186/s13046-016-0370-7>  <https://www.oncotarget.com/article/2834/text/>  <https://pmc.ncbi.nlm.nih.gov/articles/PMC4681596/>  <https://pmc.ncbi.nlm.nih.gov/articles/PMC5347702/>  <https://www.frontiersin.org/journals/pharmacology/articles/10.3389/fphar.2024.1494802/full>  <https://www.elsevier.es/en-revista-clinics-22-articulo-pik3ca-exon-20-mutations-are-S1807593222021986>  <https://ckb.jax.org/geneVariant/show?geneVariantId=2569>  <https://www.sciencedirect.com/science/article/pii/S1043661824000677>  <https://www.sciencedirect.com/science/article/pii/S2405844022026846>  <https://www.pnas.org/doi/10.1073/pnas.0510857103>  <https://www.researchgate.net/figure/PIK3CA-mutations-in-breast-cancer-frequency-of-mutations-including-information-on-the_fig1_284208979>  <https://aacrjournals.org/clincancerres/article/30/17/3868/747287/An-Analysis-of-PIK3CA-Hotspot-Mutations-and>  https://samev-dz.com/upload/articles/Non-hotspot%20PIK3CA%20mutations%20are%20more%20frequent%20in%20CLOVES%20than%20in%20common%20or%20combined%20lymphatic%20malformations.pdf |
|  | Evaluation  **Summarization Accuracy & Completeness:**  **Hallucination Rate:**  **Readability & Usability:**  **Literature Coverage & Relevance:**  **Time Efficiency:** | |
| **5** | The PIK3CA gene encodes the p110α catalytic subunit of phosphatidylinositol-4,5-bisphosphate 3-kinase (PI3K), a crucial enzyme in the PI3K/AKT/mTOR signaling pathway, which regulates cell growth, proliferation, and survival. PIK3CA is one of the most frequently mutated oncogenes in human cancers, with a significant role in tumorigenesis. The p.E545K (c.1633G>A) variant is a missense mutation in exon 9 of PIK3CA, resulting in a glutamic acid to lysine substitution at codon 545. This variant is commonly referred to as E545K and is one of the hotspot mutations in the helical domain of PIK3CA, alongside E542K and E545G. The E545K mutation is prevalent in various cancers, notably in breast cancer, where it occurs in approximately 7.0% of cases, and is also found in endometrial carcinoma, colorectal cancer, and other solid tumors. This mutation leads to increased PI3K activity by altering the protein's structure, which disrupts its interaction with the regulatory subunit and results in constitutive activation of the PI3K pathway. The E545K mutation has been shown to enhance glucose metabolism and cell proliferation, contributing to tumor growth and progression. In cervical cancer, patients with the E545K mutation exhibited higher SUVmax values, indicating increased glucose uptake and metabolic activity. The mutation has also been associated with resistance to certain chemotherapy drugs, highlighting its potential as a prognostic and predictive biomarker. Clinical trials have included E545K as an eligibility criterion, exploring targeted therapies such as alpelisib in combination with fulvestrant for treating PIK3CA-mutated cancers. The E545K variant is classified as pathogenic/likely pathogenic according to the American College of Medical Genetics and Genomics (ACMG) guidelines, underscoring its clinical significance. However, the exact frequency and impact of E545K may vary across different cancer types and populations, and further research is needed to fully elucidate its role in cancer development and treatment response. | [1] Scientific Reports - PIK3CA mutation enrichment and quantitation from blood and tissue.  [2] My Cancer Genome - PIK3CA E545K.  [3] Scientific Reports - Insights into the mechanism of the PIK3CA E545K activating mutation using MD simulations.  [4] Journal of Hematology & Oncology - The PIK3CA E542K and E545K mutations promote glycolysis and proliferation via induction of the β-catenin/SIRT3 signaling pathway in cervical cancer.  [5] PMC - Prevalence and prognostic role of PIK3CA E545K mutation in Iranian colorectal cancer patients. |
|  | Evaluation  **Summarization Accuracy & Completeness:**  **Hallucination Rate:**  **Readability & Usability:**  **Literature Coverage & Relevance:**  **Time Efficiency:** | |
