## Supplementary material for "Benchmarking generative AI tools for literature retrieval and summarization in genomic variant interpretation": POLE p.T457R.docx

| **Tool** | **Summary** | **References** |
| --- | --- | --- |
| **1** | The POLE gene encodes the catalytic subunit of DNA polymerase epsilon, a key enzyme involved in chromosomal DNA replication and repair. It plays a critical role in leading strand synthesis during DNA replication and possesses intrinsic 3′–5′ exonuclease activity for proofreading, thereby maintaining replication fidelity and genomic stability. Mutations in POLE, particularly within the exonuclease domain, have been associated with a predisposition to various cancers, including colorectal and endometrial cancers, often characterized by ultramutated tumor phenotypes. The specific variant p.T457R (threonine to arginine substitution at codon 457) has not been reported in peer-reviewed literature or major variant databases such as ClinVar or gnomAD as of this writing. Consequently, there is no established clinical significance, frequency data, or functional characterization available for this variant. Given the location of this residue within the exonuclease domain, which is critical for the proofreading function of the enzyme, alterations at this site could potentially impact the enzyme's ability to correct replication errors. However, without empirical evidence, the pathogenicity of the p.T457R variant remains uncertain. Further studies, including functional assays and clinical correlation, are necessary to determine the impact of this variant on protein function and its potential role in disease. In the absence of such data, the p.T457R variant should be classified as a Variant of Uncertain Significance (VUS), and caution should be exercised in its interpretation in a clinical context. | <https://www.ncbi.nlm.nih.gov/gene/5426>  <https://jeccr.biomedcentral.com/articles/10.1186/s13046-022-02422-1>  <https://www.ncbi.nlm.nih.gov/clinvar/63316314>  <https://www.nature.com/articles/s41436-020-0828-z>  <https://en.wikipedia.org/wiki/POLE_%28gene%29>  <https://ntrs.nasa.gov/api/citations/19670017968/downloads/19670017968.pdf>  <https://www.genecards.org/cgi-bin/carddisp.pl?gene=POLE>  <https://ckb.jax.org/gene/show?geneId=5426>  <https://www.mskcc.org/cancer-care/patient-education/about-mutations-pole-gene>  <https://www.mycancergenome.org/content/gene/POLE>  <https://stanfordlab.com/articles/POLE.html>  <https://omim.org/entry/174762> |
|  | Evaluation  **Summarization Accuracy & Completeness:**  **Hallucination Rate:**  **Readability & Usability:**  **Literature Coverage & Relevance:**  **Time Efficiency:** | |
| **2** | The POLE gene encodes the catalytic subunit of DNA polymerase epsilon, which plays a crucial role in DNA replication and repair. Specifically, POLE is involved in the proofreading and correction of errors that occur during DNA synthesis, ensuring the fidelity of DNA replication. The gene is located on chromosome 12q24.33 and its protein product contains an exonuclease domain that facilitates the removal of misincorporated nucleotides. Germline variants in POLE, particularly those affecting the exonuclease domain, are associated with an increased risk of various cancers, including colorectal, endometrial, ovarian, brain, and breast cancers, as well as polyposis. These variants can lead to a condition known as polymerase proofreading-associated polyposis (PPAP), characterized by a high mutational burden and a specific mutational spectrum in tumors. The p.T457R variant in POLE is a missense variant that results in the substitution of threonine with arginine at amino acid position 457. This variant is located within the exonuclease domain of POLE, which is critical for its proofreading function. The classification of the p.T457R variant is not explicitly detailed in the available literature, and its frequency and impact are not well-documented. However, given its location within the exonuclease domain, it is plausible that this variant could affect the proofreading function of POLE, potentially leading to an increased mutational burden and a higher risk of cancer. Further research, including functional studies and genetic testing in affected individuals and families, is necessary to determine the pathogenicity and clinical significance of the p.T457R variant. Until more data is available, the classification of this variant remains uncertain, and it should be interpreted with caution in genetic reports. | [1] Mur, P., Viana-Errasti, J., García-Mulero, S., Magraner-Pardo, L., Muñoz, I. G., Pons, T., ... & Valle, L. (2023). Recommendations for the classification of germline variants in the exonuclease domain of POLE and POLD1. Genome Medicine, 15(1), 1-24.  [2] Castellsagué, E., Li, R., Aligue, R., ... (2019). Novel POLE pathogenic germline variant in a family with multiple primary tumors results in distinct mutational signatures.  [3] Elsayed, F. A., Kets, C. M., Ruano, D., ... (2014). Germline variants in POLE are associated with early onset mismatch repair deficient colorectal cancer.  [4] Albertson, D. G. (2020). Role of POLE and POLD1 in familial cancer. Genetics in Medicine, 22(9), 1477-1486 |
|  | Evaluation  **Summarization Accuracy & Completeness:**  **Hallucination Rate:**  **Readability & Usability:**  **Literature Coverage & Relevance:**  **Time Efficiency:** | |
| **3** | There was an internal issue retrieving peer-reviewed articles about the POLE p.T457R variant from the academic database. As a result, no direct publications referencing this specific variant (p.T457R) were found. However, POLE encodes the catalytic subunit of DNA polymerase epsilon, a key enzyme involved in leading-strand DNA synthesis and DNA repair via proofreading exonuclease activity. Germline and somatic mutations in POLE have been associated with polymerase proofreading-associated polyposis and a predisposition to ultramutated tumors, particularly colorectal and endometrial cancers. Pathogenic mutations typically cluster within the exonuclease domain (residues ~268–471), where disruptions impair fidelity of DNA replication, resulting in hypermutator phenotypes. The p.T457R variant lies near the C-terminal boundary of the exonuclease domain and involves a threonine-to-arginine substitution, potentially altering local structure or function; however, no functional data or case reports specifically assessing p.T457R have been published to date. As such, this variant remains of uncertain significance, and its pathogenicity cannot be confirmed without further computational modeling, segregation analysis, or experimental validation. Clinical interpretation should be cautious, and further studies are warranted to determine its role in tumorigenesis or hereditary cancer predisposition. | No references available. |
|  | Evaluation  **Summarization Accuracy & Completeness:**  **Hallucination Rate:**  **Readability & Usability:**  **Literature Coverage & Relevance:**  **Time Efficiency:** | |
| **4** | The POLE gene encodes the catalytic subunit of DNA polymerase epsilon, a critical enzyme involved in DNA replication and repair, particularly responsible for leading strand synthesis and possessing 3' to 5' exonuclease proofreading activity to ensure replication fidelity. Variants in POLE, especially those affecting the exonuclease domain, have been implicated in increased mutational burden and cancer predisposition, notably polymerase proofreading-associated polyposis (PPAP), characterized by colorectal polyps and colorectal cancer risk. The p.T457R variant is a missense substitution located within the POLE gene; however, there is currently no direct peer-reviewed literature specifically describing the clinical or functional impact of the p.T457R variant. As such, its classification remains uncertain, and its frequency in the general population or disease cohorts is not well established. Given the known role of POLE exonuclease domain mutations in impairing proofreading function and leading to ultramutated tumor phenotypes, variants in this region are generally considered candidates for pathogenicity pending functional validation. The absence of direct evidence for p.T457R necessitates cautious interpretation, recommending further functional studies and segregation analyses to clarify its effect on polymerase activity and cancer risk. In the broader context, POLE mutations that disrupt exonuclease function result in a hypermutated state with distinctive mutational signatures and an increased tumor mutation burden, which can influence tumor behavior and response to immunotherapy. Until specific data on p.T457R emerge, clinical correlation and comprehensive genetic counseling are advised, with consideration of this variant as a variant of uncertain significance (VUS). No direct references to p.T457R exist in the current literature, highlighting a gap in knowledge and the need for further research to determine its pathogenicity and clinical relevance[1][4]. | [1] <https://www.ncbi.nlm.nih.gov/pmc/articles/PMC7521106/>  [2] <https://www.ncbi.nlm.nih.gov/pmc/articles/PMC8227676/>  [3] <https://pubmed.ncbi.nlm.nih.gov/36314330/>  [4] <https://pubmed.ncbi.nlm.nih.gov/35860951/>  [5] <https://pubmed.ncbi.nlm.nih.gov/21498422/>  [6] <https://www.semanticscholar.org/paper/aa0c0ba7b12799cef87eba880946e148cd5604c6>  [7] <https://pubmed.ncbi.nlm.nih.gov/1768621/>  [8] <https://pubmed.ncbi.nlm.nih.gov/9301023/> |
|  | Evaluation  **Summarization Accuracy & Completeness:**  **Hallucination Rate:**  **Readability & Usability:**  **Literature Coverage & Relevance:**  **Time Efficiency:** | |
| **5** | The genomic variant c.1370C>G p.T457R rs878854842 in the POLE gene has been identified and characterized in several studies, with implications for its role in cancer predisposition. The POLE gene encodes the catalytic subunit of DNA polymerase epsilon, which is involved in leading strand DNA replication and DNA repair, playing a critical role in maintaining genomic stability.  The variant c.1370C>G results in the substitution of threonine with arginine at codon 457 (p.T457R) in the POLE protein. This variant has been classified as of uncertain significance according to the ACMG guidelines, with in silico predictive tools yielding mixed results regarding its potential pathogenicity. PolyPhen-2 predicts the variant as probably damaging, while SIFT categorizes it as tolerated. MutationTaster suggests the variant is disease-causing, and it has a CADD score of 25.1, indicating a potential deleterious effect [5].  The c.1370C>G variant has been reported as absent in the ExAC population database, suggesting it is not a common polymorphism in the general population [1]. This rarity in the general population may contribute to the uncertainty regarding its clinical significance. Conservation analysis shows that the threonine at position 457 is conserved in four out of six species, which may indicate functional importance [1].  Clinically, the variant has been identified in a patient diagnosed with papillary thyroid carcinoma (PTC) at the early age of 18, suggesting a possible association with early-onset cancer [1]. However, the evidence is not sufficient to establish a definitive causal relationship between the variant and PTC, and further functional studies are required to elucidate the impact of the p.T457R substitution on POLE function and its role in tumorigenesis.  In summary, the c.1370C>G p.T457R rs878854842 variant in the POLE gene is a rare missense change with uncertain clinical significance. Its potential pathogenicity is supported by some in silico predictions and its absence in population databases, but further research is necessary to clarify its role in cancer predisposition and to provide more definitive guidance for clinical management. | [[1]](https://pubmed.ncbi.nlm.nih.gov/32992294/) Siraj AK et al. (2020). "POLE and POLD1 pathogenic variants in the proofreading domain in papillary thyroid cancer." Endocrine connections, 9(9)  [[2]](https://pubmed.ncbi.nlm.nih.gov/37990341/) Weber CAM et al. (2023). "Rare germline variants in POLE and POLD1 encoding the catalytic subunits of DNA polymerases epsilon and delta in glioma families." Acta neuropathologica communications, 11(1)  [[3]](https://pubmed.ncbi.nlm.nih.gov/25860647/) Hansen MF et al. (2015). "A novel POLE mutation associated with cancers of colon, pancreas, ovaries and small intestine." Familial cancer, 14(3)  [[4]](https://pubmed.ncbi.nlm.nih.gov/25124163/) Stenzinger A et al. (2014). "Mutations in POLE and survival of colorectal cancer patients--link to disease stage and treatment." Cancer medicine, 3(6)  [[5]](https://pubmed.ncbi.nlm.nih.gov/32567205/) Siraj AK et al. (2020). "POLE and POLD1 germline exonuclease domain pathogenic variants, a rare event in colorectal cancer from the Middle East." Molecular genetics & genomic medicine, 8(8)  [[6]](https://pubmed.ncbi.nlm.nih.gov/32801757/) Li Y et al. (2020). "POLE Mutation Characteristics in a Chinese Cohort with Endometrial Carcinoma." OncoTargets and therapy, 13()  [[7]](https://pubmed.ncbi.nlm.nih.gov/36629684/) Kanno DT et al. (2023). "MULTIPLE NEUROENDOCRINE NEOPLASIA IN A PATIENT WITH TYPE I NEUROFIBROMATOSIS (NF1): REPORT OF A NEW MUTATION (NF1, EXONS 2-30 DELETION) AND LITERATURE REVIEW." Arquivos brasileiros de cirurgia digestiva : ABCD = Brazilian archives of digestive surgery, 35() |
|  | Evaluation  **Summarization Accuracy & Completeness:**  **Hallucination Rate:**  **Readability & Usability:**  **Literature Coverage & Relevance:**  **Time Efficiency:** | |
