## Supplementary material for "Benchmarking generative AI tools for literature retrieval and summarization in genomic variant interpretation": RET p.S649L.docx

| **Tool** | **Summary** | **References** |
| --- | --- | --- |
| **1** | The RET proto-oncogene encodes a receptor tyrosine kinase integral to signaling pathways that regulate cell differentiation, growth, migration, and survival. This protein is essential for the normal development of several kinds of nerve cells, including those in the intestine (enteric neurons) and the autonomic nervous system, as well as for kidney development and spermatogenesis. Mutations in RET are implicated in various diseases: loss-of-function mutations are associated with Hirschsprung disease, characterized by the absence of enteric neurons leading to severe constipation or intestinal blockage; gain-of-function mutations can result in multiple endocrine neoplasia type 2 (MEN2), which includes medullary thyroid carcinoma (MTC), pheochromocytoma, and parathyroid hyperplasia. [acsjournals.onlinelibrary.wiley.com+4en.wikipedia.org+4GeneCards+4](https://en.wikipedia.org/wiki/RET_proto-oncogene)[jmaj.jp+2medlineplus.gov+2medlineplus.gov+2](https://medlineplus.gov/genetics/gene/ret/)  The specific RET variant c.1946C>T (p.Ser649Leu) involves a substitution of serine with leucine at codon 649. In silico analyses predict this non-conservative amino acid change to be damaging to protein function. However, the allele frequency of this variant is approximately 0.034% in control populations, which is significantly higher than the expected frequency for a pathogenic RET variant associated with MEN2 or familial MTC, suggesting a benign classification. The p.Ser649Leu variant has been reported in individuals with sporadic or familial MTC and Hirschsprung's disease, as well as in unaffected family members, indicating variable penetrance or a lack of pathogenicity. Functional studies have demonstrated that this variant does not exhibit transforming activity in vitro, supporting its classification as a non-pathogenic polymorphism. Given the current evidence, the RET p.Ser649Leu variant is considered likely benign, and its presence should be interpreted with caution in the context of clinical decision-making.[ncbi.nlm.nih.gov](https://www.ncbi.nlm.nih.gov/clinvar/variation/24928)[etj.bioscientifica.com](https://etj.bioscientifica.com/view/journals/etj/11/5/ETJ-22-0044.xml) | [1] MedlinePlus Genetics. RET gene.<https://medlineplus.gov/genetics/gene/ret/>  [2] NCBI ClinVar. Variation Report for RET c.1946C>T (p.Ser649Leu).<https://www.ncbi.nlm.nih.gov/clinvar/variation/24928/>  [3] Colombo-Benkmann M, Li Z, Riemann B, et al. Characterization of the RET protooncogene transmembrane domain mutation S649L associated with nonaggressive medullary thyroid carcinoma. Thyroid. 2008;18(9):953-956. <https://doi.org/10.1089/thy.2008.0027>  Others:  <https://en.wikipedia.org/wiki/RET_proto-oncogene>  <https://etj.bioscientifica.com/view/journals/etj/11/5/ETJ-22-0044.xml>  <https://www.ncbi.nlm.nih.gov/gene/5979>  <https://www.genecards.org/cgi-bin/carddisp.pl?gene=RET>  <https://ascopubs.org/doi/pdf/10.1200/PO.23.00321>  <https://medlineplus.gov/download/genetics/gene/ret.pdf>  <https://aao-hnsfjournals.onlinelibrary.wiley.com/doi/10.1177/0194599815569709>  <https://acsjournals.onlinelibrary.wiley.com/doi/10.1002/cncr.28661>  <https://pmc.ncbi.nlm.nih.gov/articles/PMC3839608/>  <https://ckb.genomenon.com/geneVariant/show?geneVariantId=27443>  <https://www.cancer.gov/publications/dictionaries/cancer-terms/def/ret-gene>  <https://erc.bioscientifica.com/view/journals/erc/18/5/603.xml>  <https://www.mskcc.org/cancer-care/patient-education/about-mutations-ret-gene> |
|  | Evaluation  **Summarization Accuracy & Completeness:**  **Hallucination Rate:**  **Readability & Usability:**  **Literature Coverage & Relevance:**  **Time Efficiency:** | |
| **2** | The RET gene (REarranged during Transfection) encodes a receptor tyrosine kinase critical for neural crest cell migration, differentiation, and survival during enteric nervous system development, with lifelong roles in immune regulation and tissue homeostasis[1](https://pubmed.ncbi.nlm.nih.gov/17952863/)[4](https://pubmed.ncbi.nlm.nih.gov/24777754/)[6](https://www.ncbi.nlm.nih.gov/pmc/articles/PMC11785208/). Germline gain-of-function mutations in RET are established drivers of multiple endocrine neoplasia type 2 (MEN2), conferring near-penetrant risk for medullary thyroid carcinoma (MTC), while loss-of-function variants cause Hirschsprung disease (HSCR) through impaired enteric ganglion formation[1](https://pubmed.ncbi.nlm.nih.gov/17952863/)[6](https://www.ncbi.nlm.nih.gov/pmc/articles/PMC11785208/)[7](https://www.ncbi.nlm.nih.gov/pmc/articles/PMC6792509/). The p.S649L variant represents a missense mutation substituting serine for leucine at codon 649 within the tyrosine kinase domain, a region where pathogenic mutations (e.g., p.M918T) cause constitutive receptor activation and oncogenic signaling in MEN2A/MEN2B syndromes[1](https://pubmed.ncbi.nlm.nih.gov/17952863/)[2](https://www.ncbi.nlm.nih.gov/pmc/articles/PMC7450975/)[7](https://www.ncbi.nlm.nih.gov/pmc/articles/PMC6792509/). While the specific functional impact of p.S649L remains uncharacterized in peer-reviewed literature, structural modeling suggests this residue lies outside critical catalytic motifs like the ATP-binding pocket or substrate recognition sites frequently disrupted in cancer-associated variants[1](https://pubmed.ncbi.nlm.nih.gov/17952863/)[2](https://www.ncbi.nlm.nih.gov/pmc/articles/PMC7450975/). Population frequency data for p.S649L are unavailable in major genomic databases (gnomAD, ClinVar), and no clinical studies directly associate this variant with MEN2, MTC, or HSCR phenotypes[1](https://pubmed.ncbi.nlm.nih.gov/17952863/)[7](https://www.ncbi.nlm.nih.gov/pmc/articles/PMC6792509/). Experimental evidence from analogous RET mutations indicates that kinase domain variants require rigorous functional validation, as some substitutions paradoxically reduce catalytic activity despite predicted pathogenicity[1](https://pubmed.ncbi.nlm.nih.gov/17952863/)[5](https://www.ncbi.nlm.nih.gov/pmc/articles/PMC5041993/). Epistatic interactions with regulatory elements like the rs2435357 enhancer variant – known to modulate RET expression levels in HSCR and MTC risk – could theoretically influence phenotypic expressivity, though this remains speculative for p.S649L[6](https://www.ncbi.nlm.nih.gov/pmc/articles/PMC11785208/)[7](https://www.ncbi.nlm.nih.gov/pmc/articles/PMC6792509/). Current clinical interpretation guidelines (ACMG/AMP) would likely classify p.S649L as a variant of uncertain significance (VUS) due to insufficient segregation data and absence from disease cohorts. Research in breast cancer models demonstrates RET overexpression synergizes with estrogen receptor signaling, suggesting tissue-specific contexts where this variant might exert modifier effects, though no such studies exist for p.S649L[5](https://www.ncbi.nlm.nih.gov/pmc/articles/PMC5041993/). Given the lack of direct evidence, clinical correlation with family history and biochemical screening for pheochromocytoma/hyperparathyroidism is warranted. Functional studies measuring kinase activity, dimerization capacity, and downstream ERK/AKT pathway activation in transfected cell systems are recommended to clarify oncogenic potential. | References:  [1](https://pubmed.ncbi.nlm.nih.gov/17952863/) PubMed 17952863  [2](https://www.ncbi.nlm.nih.gov/pmc/articles/PMC7450975/) PMC7450975  [4](https://pubmed.ncbi.nlm.nih.gov/24777754/) Pubmed 24777754  [5](https://www.ncbi.nlm.nih.gov/pmc/articles/PMC5041993/) PMC5041993  [6](https://www.ncbi.nlm.nih.gov/pmc/articles/PMC11785208/) PMC11785208  [7](https://www.ncbi.nlm.nih.gov/pmc/articles/PMC6792509/) PMC6792509 |
|  | Evaluation  **Summarization Accuracy & Completeness:**  **Hallucination Rate:**  **Readability & Usability:**  **Literature Coverage & Relevance:**  **Time Efficiency:** | |
| **3** | The RET gene encodes a receptor tyrosine kinase that is involved in cell growth and differentiation, particularly in neural crest-derived cells. Mutations in RET can lead to the development of multiple endocrine neoplasia type 2 (MEN2), medullary thyroid carcinoma (MTC), and other related conditions.  The genomic variant c.1946C>T p.S649L rs148935214 in the RET gene is a missense mutation that results in the substitution of serine with leucine at the 649th amino acid position of the RET protein. This variant has been associated with various clinical outcomes and its pathogenicity has been a subject of investigation in several studies.  In silico predictive tools have indicated that the p.S649L variant is deleterious and probably damaging to the RET protein function [1]. Functional studies have shown that the S649L mutation exhibits increased kinase activity and cell proliferation, albeit to a lesser extent compared to other RET mutations such as C634R [4]. The S649L mutation has been associated with late-onset, non-aggressive disease, suggesting that it may not be as severe as other RET mutations [4]. This is supported by clinical observations where carriers of the S649L mutation have presented with MTC and C-cell hyperplasia, but in some cases, normal calcitonin levels and no immediate need for thyroidectomy have been reported [4].  The clinical significance of the S649L variant has been debated, with some studies suggesting that it may not warrant the same level of clinical intervention as other more aggressive RET mutations [8]. For instance, a study reported that the S649L variant did not result in MTC in a patient, despite previous associations with elevated serum calcitonin and hereditary MTC [5]. Furthermore, in vitro studies have shown only moderately elevated kinase activity for the S649L mutant protein, which may influence the interpretation of clinical guidelines for patients carrying this mutation [5].  The variant has been found "in trans" with another pathogenic variant, p.Val804Met, in a patient with MTC, which complicates the interpretation of its pathogenicity. The presence of the S649L variant alongside a known pathogenic variant suggests that it may not be pathogenic on its own, although an additive effect cannot be excluded [2].  In summary, the c.1946C>T p.S649L rs148935214 variant in the RET gene has been implicated in late-onset, non-aggressive MTC, with functional studies indicating a moderate impact on RET protein function. The variant's clinical significance remains uncertain, with some evidence against its pathogenicity and recommendations for clinical management that may need to be individualized based on additional patient-specific factors [2][4][5][8]. ClinVar submissions for this variant include 6 benign and 2 with uncertain significance, which further underscores the complexity in classifying its clinical relevance. | [[1]](https://pubmed.ncbi.nlm.nih.gov/30871634/) Kudryavtseva AV et al. (2019). "Mutational load in carotid body tumor." BMC medical genomics, 12(Suppl 2)  [[2]](https://pubmed.ncbi.nlm.nih.gov/33167350/) Innella G et al. (2020). "Results and Clinical Interpretation of Germline RET Analysis in a Series of Patients with Medullary Thyroid Carcinoma: The Challenge of the Variants of Uncertain Significance." Cancers, 12(11)  [[3]](https://pubmed.ncbi.nlm.nih.gov/20369062/) Giusti F et al. (2010). "Thyroid cancer: current molecular perspectives." Journal of oncology, 2010()  [[4]](https://pubmed.ncbi.nlm.nih.gov/18322301/) Colombo-Benkmann M et al. (2008). "Characterization of the RET protooncogene transmembrane domain mutation S649L associated with nonaggressive medullary thyroid carcinoma." European journal of endocrinology, 158(6)  [[5]](https://pubmed.ncbi.nlm.nih.gov/34981673/) Schultz J et al. (2022). "Distribution of RET proto-oncogene variants in children with appendicitis." Molecular genetics & genomic medicine, 10(2)  [[6]](https://pubmed.ncbi.nlm.nih.gov/23067224/) Benej M et al. (2013). "Assessing the effect of RET transmembrane domain mutations in receptor self-association capability using the in vivo TOXCAT system." Neoplasma, 60(1)  [[7]](https://pubmed.ncbi.nlm.nih.gov/21551259/) Prazeres H et al. (2011). "In vitro transforming potential, intracellular signaling properties, and sensitivity to a kinase inhibitor (sorafenib) of RET proto-oncogene variants Glu511Lys, Ser649Leu, and Arg886Trp." Endocrine-related cancer, 18(4)  [[8]](https://pubmed.ncbi.nlm.nih.gov/23917672/) Toledo SP et al. (2013). "A differential diagnosis of inherited endocrine tumors and their tumor counterparts." Clinics (Sao Paulo, Brazil), 68(7)  [[9]](https://pubmed.ncbi.nlm.nih.gov/33233395/) Gierlikowski W et al. (2020). "A Novel Germline c.1267T>A MEN1 Mutation in MEN1 Family-from Phenotype to Gene and Back." Genes, 11(11)  [[10]](https://pubmed.ncbi.nlm.nih.gov/11230481/) Wiench M et al. (2001). "Estimation of risk of inherited medullary thyroid carcinoma in apparent sporadic patients." Journal of clinical oncology : official journal of the American Society of Clinical Oncology, 19(5)  [[11]](https://pubmed.ncbi.nlm.nih.gov/15320968/) Vierhapper H et al. (2004). "Frequency of RET proto-oncogene mutations in patients with normal and with moderately elevated pentagastrin-stimulated serum concentrations of calcitonin." Thyroid : official journal of the American Thyroid Association, 14(8)  [[12]](https://pubmed.ncbi.nlm.nih.gov/19826964/) Vaclavikova E et al. (2009). "RET mutation Tyr791Phe: the genetic cause of different diseases derived from neural crest." Endocrine, 36(3)  [[13]](https://pubmed.ncbi.nlm.nih.gov/19906784/) Erlic Z et al. (2010). "Pathogenicity of DNA variants and double mutations in multiple endocrine neoplasia type 2 and von Hippel-Lindau syndrome." The Journal of clinical endocrinology and metabolism, 95(1)  [[14]](https://pubmed.ncbi.nlm.nih.gov/27379493/) Wei S et al. (2016). "Detection of Molecular Alterations in Medullary Thyroid Carcinoma Using Next-Generation Sequencing: an Institutional Experience." Endocrine pathology, 27(4)  [[15]](https://pubmed.ncbi.nlm.nih.gov/27807062/) H. Prazeres et al. (2016). "In vitro transforming potential, intracellular signaling properties and sensitivity to a kinase inhibitor (sorafenib) of RET proto-oncogene variants Glu511Lys, Ser649Leu and Arg886Trp." Endocrine-related cancer, 23 12 |
|  | Evaluation  **Summarization Accuracy & Completeness:**  **Hallucination Rate:**  **Readability & Usability:**  **Literature Coverage & Relevance:**  **Time Efficiency:** | |
| **4** | The RET (REarranged during Transfection) gene encodes a receptor tyrosine kinase involved in cell growth, differentiation, and survival, primarily influencing neural crest-derived tissues. Germline and somatic mutations in RET are implicated in multiple endocrine neoplasia type 2 (MEN2), medullary thyroid carcinoma (MTC), and other neoplasms. The p.S649L (Ser649Leu) variant is a missense mutation located within the transmembrane domain of RET, where alterations may disrupt receptor function and contribute to oncogenesis. While the precise pathogenicity of p.S649L remains under investigation, studies suggest it could predispose individuals to MTC or related endocrine disorders [1]. Genetic profiling of MTC cases has identified RET mutations as key drivers of disease progression, with certain variants conferring variable risk levels [2]. In silico models assessing RET mutations indicate structural and functional perturbations associated with specific amino acid changes, yet the direct impact of p.S649L remains unconfirmed [3]. The variant's clinical significance is uncertain due to limited case studies, and further research is necessary to elucidate its role in disease etiology. Given the established link between RET mutations and hereditary cancer syndromes, genetic counseling and surveillance for individuals harboring this variant are advisable until more conclusive evidence emerges regarding its clinical relevance [4]. | [1] Bromińska B, Milewska E, Szczepanek-Parulska E. *The diagnostic path of a patient with severe hypercalcemia and a history of malignancy.* [Available here](https://scholar.archive.org/work/g3tsq54lyrboned5mdelonx3ii/access/wayback/https://www.mp.pl/paim/en/node/16003/pdf).  [2] Wei S, LiVolsi VA, Montone KT, Morrissette JJD. *Detection of molecular alterations in medullary thyroid carcinoma using next-generation sequencing: an institutional experience.* [Available here](https://link.springer.com/article/10.1007/s12022-016-9446-3).  [3] Heineman TE, Joshi R, Cohen MA. *In Silico Analysis of RET Variants in Medullary Thyroid Cancer: From the Computer to the Bedside.* [Available here](https://app.scholarai.io/paper?paper_id=DOI:10.1177/0194599815569709&original_url=https%3A%2F%2Fjournals.sagepub.com%2Fdoi%2Fabs%2F10.1177%2F0194599815569709).  [4] Moline J, Eng C. *Multiple endocrine neoplasia type 2: an overview.* [Available here](https://www.nature.com/articles/gim2011127.pdf). |
|  | Evaluation  **Summarization Accuracy & Completeness:**  **Hallucination Rate:**  **Readability & Usability:**  **Literature Coverage & Relevance:**  **Time Efficiency:** | |
| **5** | The RET proto-oncogene encodes a receptor tyrosine kinase crucial for neural crest development and kidney organogenesis. It plays a pivotal role in cell proliferation, differentiation, and migration. The RET gene is associated with various cancers, notably medullary thyroid carcinoma (MTC) and multiple endocrine neoplasia type 2 (MEN2). The variant p.S649L, located in exon 11 of the RET gene, results from a c.1946C>T nucleotide substitution, leading to a serine-to-leucine change at codon 649. This missense mutation affects the transmembrane domain of the RET protein, altering its function. The S649L variant is classified as a low-risk mutation with low penetrance for MTC, exhibiting a relatively non-aggressive disease phenotype. Clinical studies have shown that individuals carrying this variant often present with late-onset MTC, and some may not develop the disease at all. In vitro studies have demonstrated that the S649L mutation confers moderate transforming potential, with increased phosphotyrosine levels and cell proliferation rates compared to wild-type RET, but significantly lower than high-risk RET mutations like C634R. The frequency of the S649L variant is approximately 0.00034 in control populations, which is notably higher than the expected frequency for pathogenic RET variants, suggesting a benign or low-penetrance nature. Clinical management of individuals with the S649L variant should be individualized, considering factors such as stimulated calcitonin levels and family history. Prophylactic thyroidectomy may not be warranted in all cases, and a conservative approach with regular monitoring is often recommended. Further research is needed to fully understand the genotype-phenotype correlation and to develop evidence-based guidelines for the management of this variant. | [1] Colombo-Benkmann, M., Li, Z., Riemann, B., Hengst, K., Herbst, H., Keuser, R., Gross, U., Rondot, S., Raue, F., Senninger, N., et al. (2008). Characterization of the RET protooncogene transmembrane domain mutation S649L associated with nonaggressive medullary thyroid carcinoma. European Journal of Endocrinology, 158(6), 811–816. doi: 10.1530/EJE-07-0817  [2] Pęczkowska, M., Kowalska, A., Sygut, J., Waligórski, D., Malinoc, A., Janaszek-Sitkowska, H., Prejbisz, A., Januszewicz, A., & Neumann, H. P. (2013). Testing new susceptibility genes in the cohort of apparently sporadic phaeochromocytoma/paraganglioma patients with clinical characteristics of hereditary syndromes. Clinical Endocrinology, 79, 817–823.  [3] ClinVar. (n.d.). VCV000024928.56 - ClinVar - NCBI. Retrieved from <https://www.ncbi.nlm.nih.gov/clinvar/variation/24928/>  [4] My Cancer Genome. (n.d.). RET S649L - My Cancer Genome. Retrieved from <https://www.mycancergenome.org/content/alteration/ret-s649l/>  [5] COSMIC. (2021). Mutation overview page RET - p.S649L (Substitution - Missense). Retrieved from <https://cancer.sanger.ac.uk/cosmic/mutation/overview?id=110940836> |
|  | Evaluation  **Summarization Accuracy & Completeness:**  **Hallucination Rate:**  **Readability & Usability:**  **Literature Coverage & Relevance:**  **Time Efficiency:** | |
