## Supplementary material for "Benchmarking generative AI tools for literature retrieval and summarization in genomic variant interpretation": SMAD4 p.R361C.docx

| **Tool** | **Summary** | **References** |
| --- | --- | --- |
| **1** | The SMAD4 gene, located on chromosome 18q21.2, encodes the Smad4 protein, a critical mediator in the transforming growth factor-beta (TGF-β) signaling pathway. This pathway regulates various cellular processes, including growth, differentiation, and apoptosis. Smad4 functions as a common mediator (co-SMAD) that forms heterotrimeric complexes with receptor-regulated SMADs (R-SMADs) to translocate into the nucleus and regulate gene expression. The p.R361C variant in SMAD4, also known as c.1081C>T, results from a C to T substitution at nucleotide position 1081 in exon 8, leading to an arginine to cysteine substitution at codon 361. This variant is classified as pathogenic and is associated with several clinical conditions, notably hereditary hemorrhagic telangiectasia (HHT) and juvenile polyposis syndrome (JPS). The p.R361C variant has been reported in multiple individuals affected with these syndromes and is absent from general population databases, indicating it is not a common polymorphism. Functional studies have shown that the p.R361C variant confers a loss of function to the Smad4 protein, demonstrated by reduced transactivating activity in cell assays. This loss of function is likely oncogenic, as studies of somatic mutations and loss of heterozygosity (LOH) at the SMAD4 locus have shown an association with poor prognosis in various cancers, including pancreatic and colorectal cancers. The variant is present in 0.26% of AACR GENIE cases, with the highest prevalence in colon adenocarcinoma, colorectal adenocarcinoma, pancreatic adenocarcinoma, lung adenocarcinoma, and rectal adenocarcinoma. The p.R361C variant abolishes the interaction between Smad4 and R-SMADs, affecting the formation of functional Smad complexes and subsequently impairing the regulation of gene expression. This disruption in the TGF-β signaling pathway contributes to the pathogenesis of associated syndromes and cancers. The exact molecular mechanisms by which the p.R361C variant exerts its effects are still under investigation, and further studies are needed to fully elucidate its role in disease progression. | Gallione CJ, et al. J Med Genet. 2006 Oct;43(10):793-7.  Gallione CJ, et al. Am J Med Genet A. 2010 Feb;152A(2):333-9.  Houlston RS, et al. Hum Mol Genet. 1998 Nov;7(12):1907-12.  Aretz S, et al. J Med Genet. 2007 Nov;44(11):702-9.  Liu X, et al. Human Mutation. 2015;37:235-241.  The AACR Project GENIE Consortium. Cancer Discovery. 2017;7(8):818-831.  Howe JR, et al. J Med Genet. 2004 Jul;41(7):484-91.  Shi Y, et al. J Med Genet. 1998 Nov;7(12):1907-12. |
|  | Evaluation  **Summarization Accuracy & Completeness:**  **Hallucination Rate:**  **Readability & Usability:**  **Literature Coverage & Relevance:**  **Time Efficiency:** | |
| **2** | The genomic variant c.1081C>T p.R361C rs80338963 in the SMAD4 gene has been identified in multiple studies and is associated with various clinical implications. The SMAD4 gene encodes a protein that is a central component of the TGF-beta signaling pathways, which are involved in a range of cellular processes including cell growth, differentiation, and apoptosis. Mutations in SMAD4 can lead to dysregulation of these pathways and contribute to the development of diseases such as hereditary hemorrhagic telangiectasia (HHT), juvenile polyposis syndrome (JPS), and various cancers.  The p.R361C variant has been found in three patients from three different families, with one case being a proven de novo occurrence [1]. This missense mutation is located in the MH1 domain of the SMAD4 protein and has been previously described in ovarian and colonic cancers. It disrupts the hetero-oligomerisation of SMAD4 with SMAD2, blocking signal transduction and leading to a loss of function phenotype [1]. This loss of function is consistent with the observation that other frameshift variants in SMAD4 also lead to disease phenotypes, suggesting that the loss of SMAD4 function contributes to the associated clinical manifestations [1].  In the context of cancer, the p.R361C variant has been identified as a significant predictor of intrinsic resistance (IR) to therapy. Specifically, patients with tumors harboring the SMAD4 p.R361C mutation experienced a more significant increase in tumor size following treatment and had materially worse survival outcomes compared to those with wild-type SMAD4 [3]. This mutation, along with other hotspot mutations in the MH2 domain, was found to be associated with resistance to BVZ-C therapy, a treatment for colorectal cancer [3].  In pancreatic ductal adenocarcinomas (PDACs), the p.R361C variant is one of several types of mutations found in the SMAD4 gene, which is mutated in approximately 50% of PDAC cases [4]. The variant has also been implicated in gallbladder cancer (GBC), where in silico studies have predicted that it results in the formation of a deformed or unstable protein structure, potentially affecting the protein's ability to interact with other molecules and leading to functional impairment [5].  Furthermore, the p.R361C variant has been associated with more aggressive features in colorectal cancers, such as higher stage, tumor deposits, nodal metastasis, and mucinous features [6]. It has been observed in colorectal carcinoma alongside other mutations commonly seen in this cancer type, such as BRAF p.V600E and mutations in TP53 [6].  Despite its association with cancer, the p.R361C variant has also been reported in a patient with juvenile polyposis syndrome, confirming SMAD4 as a cause of JPS but also indicating genetic heterogeneity in this disease [8]. ClinVar has recorded a total of 25 pathogenic submissions for this variant, underscoring its clinical significance.  In summary, the c.1081C>T p.R361C rs80338963 variant in the SMAD4 gene is a pathogenic mutation with high penetrance, associated with a range of clinical phenotypes including HHT, JPS, and various cancers. Its presence can influence treatment response and prognosis, particularly in the context of cancer therapy resistance. | [[1]](https://pubmed.ncbi.nlm.nih.gov/38575304/) Caillot C et al. (2024). "Phenotypic characterisation of SMAD4 variant carriers." Journal of medical genetics, 61(8)  [[2]](https://pubmed.ncbi.nlm.nih.gov/39660097/) Yamamoto S et al. (2025). "Comprehensive genomic profiling of primary bladder mucinous adenocarcinoma, a rare genitourinary cancer: A case report." Urology case reports, 58()  [[3]](https://pubmed.ncbi.nlm.nih.gov/39631402/) Shi M et al. (2024). "Genetic and microenvironmental evolution of colorectal liver metastases under chemotherapy." Cell reports. Medicine, 5(12)  [[4]](https://pubmed.ncbi.nlm.nih.gov/38791887/) Olaoba OT et al. (2024). "Driver Mutations in Pancreatic Cancer and Opportunities for Targeted Therapy." Cancers, 16(10)  [[5]](https://pubmed.ncbi.nlm.nih.gov/32151199/) Kumar R et al. (2021). "Deciphering the impact of missense mutations on structure and dynamics of SMAD4 protein involved in pathogenesis of gall bladder cancer." Journal of biomolecular structure & dynamics, 39(6)  [[6]](https://pubmed.ncbi.nlm.nih.gov/35178262/) Marji N et al. (2022). "Colonic Adenocarcinoma with Plasmacytoid Feature: Histopathology and Molecular Characteristics of a Rare Neoplasm with an Unusual Presentation." Case reports in pathology, 2022()  [[7]](https://pubmed.ncbi.nlm.nih.gov/25523272/) Tone AA et al. (2014). "Intratumoral heterogeneity in a minority of ovarian low-grade serous carcinomas." BMC cancer, 14()  [[8]](https://pubmed.ncbi.nlm.nih.gov/9811934/) Houlston R et al. (1998). "Mutations in DPC4 (SMAD4) cause juvenile polyposis syndrome, but only account for a minority of cases." Human molecular genetics, 7(12)  [[9]](https://pubmed.ncbi.nlm.nih.gov/32904697/) Tian X et al. (2020). "A Novel Mutation in MYH Gene Associated with Aggressive Colorectal Cancer in a Child: A Case Report and Review of Literature." OncoTargets and therapy, 13()  [[10]](https://pubmed.ncbi.nlm.nih.gov/32440349/) Ye J et al. (2020). "Tissue gene mutation profiles in patients with colorectal cancer and their clinical implications." Biomedical reports, 13(1)  [[11]](https://pubmed.ncbi.nlm.nih.gov/32944796/) Hashimoto Y et al. (2020). "Juvenile polyposis syndrome-hereditary hemorrhagic telangiectasia associated with a SMAD4 mutation in a girl." Clinical journal of gastroenterology, 13(6)  [[12]](https://ashpublications.org/blood/article/138/Supplement%201/1048/480990/Resveratrol-As-a-Novel-Treatment-for-Recurrent) Shahbegh Gill et al. (2021). "Resveratrol As a Novel Treatment for Recurrent Epistaxis in SMAD4+ Hereditary Hemorrhagic Telangiectasia" Blood,  [[13]](https://aacrjournals.org/cancerimmunolres/article/10/12_Supplement/B55/711271/Abstract-B55-Patient-Derived-Organoids-model-time) M. Mastropaolo et al. (2022). "Abstract B55: Patient-Derived Organoids model time-dependent sensitivities to PARP inhibitors in patients with metastatic colorectal cancer" Cancer Immunology Research,  [[14]](https://pubmed.ncbi.nlm.nih.gov/38406801/) Barzi A et al. (2024). "ERBB2 (HER2) amplifications and co-occurring KRAS alterations in the circulating cell-free DNA of pancreatic ductal adenocarcinoma patients and response to HER2 inhibition." Frontiers in oncology, 14()  [[15]](https://pubmed.ncbi.nlm.nih.gov/39377143/) Ito S et al. (2024). "Usefulness of multigene liquid biopsy of bile for identifying driver genes of biliary duct cancers." Cancer science, 115(12) |
|  | Evaluation  **Summarization Accuracy & Completeness:**  **Hallucination Rate:**  **Readability & Usability:**  **Literature Coverage & Relevance:**  **Time Efficiency:** | |
| **3** | The SMAD4 gene encodes a critical transcription factor within the TGF-β signaling pathway, which regulates cellular proliferation, differentiation, and apoptosis. Pathogenic variants in SMAD4 are associated with hereditary hemorrhagic telangiectasia (HHT) and juvenile polyposis syndrome (JPS), both of which predispose individuals to cancer. The missense variant p.R361C (c.1081C>T) in SMAD4 has been reported in neuroendocrine tumors, particularly in rectal neuroendocrine tumors (NETs), and may be implicated in tumorigenesis through disruption of SMAD4-mediated signaling [1]. This variant is not commonly reported in SNP databases or control populations, suggesting it is not a benign polymorphism [2]. The functional consequence of p.R361C remains uncertain, though amino acid substitutions at this position, such as the p.R361G variant, have been linked to altered SMAD4 protein function, potentially affecting its specificity and selectivity in cellular signaling [3]. Given the established role of SMAD4 mutations in tumorigenesis, p.R361C warrants further investigation to clarify its pathogenicity, potential impact on protein stability, and clinical relevance. At present, no definitive classification exists for this variant in clinical guidelines, and additional studies using functional assays and patient-derived data are necessary to determine its effect. Genetic testing and family history evaluation remain important for individuals carrying this variant, particularly in the context of hereditary cancer syndromes. | [1] Park et al., 2019 (<https://www.sciencedirect.com/science/article/pii/S0046817719300243>);  [2] Gallione et al., 2010 (<https://app.scholarai.io/paper?paper_id=DOI:10.1002/ajmg.a.33206&original_url=https%3A%2F%2Fonlinelibrary.wiley.com%2Fdoi%2Fabs%2F10.1002%2Fajmg.a.33206>);  [3] Pan et al., 2020 (<https://app.scholarai.io/paper?paper_id=DOI:10.3892/ol.2020.11851&original_url=https%3A%2F%2Fwww.spandidos-publications.com%2F10.3892%2Fol.2020.11851>). |
|  | Evaluation  **Summarization Accuracy & Completeness:**  **Hallucination Rate:**  **Readability & Usability:**  **Literature Coverage & Relevance:**  **Time Efficiency:** | |
| **4** | The SMAD4 gene plays a crucial role as a mediator in the transforming growth factor-beta (TGF-β) signaling pathway, acting primarily as a tumor suppressor. It is frequently mutated in various cancers, including colorectal and pancreatic cancers, where its loss is associated with poorer patient outcomes[3](https://www.ncbi.nlm.nih.gov/pmc/articles/PMC10158796/). The variant p.R361C in the SMAD4 gene has not been specifically detailed in the available literature, suggesting a need for further investigation to understand its clinical significance. Generally, SMAD4 mutations can lead to the inactivation of its tumor suppressive functions, facilitating tumor progression and metastasis[1](https://www.ncbi.nlm.nih.gov/pmc/articles/PMC10983983/). In some cases, SMAD4 mutations are associated with specific syndromes, such as juvenile polyposis syndrome due to loss-of-function variants, while gain-of-function variants are linked to Myhre syndrome, which may increase cancer susceptibility[8](https://pubmed.ncbi.nlm.nih.gov/31837202/). The impact of SMAD4 mutations on cancer progression highlights the importance of understanding the specific effects of different variants, including p.R361C, to develop targeted therapeutic strategies. Given the lack of direct evidence regarding the p.R361C variant, further research is necessary to classify its pathogenicity and clinical implications. | [1] <https://www.ncbi.nlm.nih.gov/pmc/articles/PMC10983983/>  [2] <https://pubmed.ncbi.nlm.nih.gov/36740808/>  [3] <https://www.ncbi.nlm.nih.gov/pmc/articles/PMC10158796/>  [4] <https://pmc.ncbi.nlm.nih.gov/articles/PMC10795321/>  [5] <https://pmc.ncbi.nlm.nih.gov/articles/PMC11310865/>  [6] <https://pmc.ncbi.nlm.nih.gov/articles/PMC10832098/>  [7] <https://pmc.ncbi.nlm.nih.gov/articles/PMC11496865/>  [8] <https://pubmed.ncbi.nlm.nih.gov/31837202/> |
|  | Evaluation  **Summarization Accuracy & Completeness:**  **Hallucination Rate:**  **Readability & Usability:**  **Literature Coverage & Relevance:**  **Time Efficiency:** | |
| **5** | The SMAD4 gene, located on chromosome 18q21.2, encodes a protein integral to the transforming growth factor-beta (TGF-β) signaling pathway, which regulates cell proliferation, differentiation, and apoptosis. SMAD4 functions as a common mediator (co-SMAD), partnering with receptor-regulated SMADs (R-SMADs) to form complexes that translocate to the nucleus and modulate gene expression. The p.R361C variant denotes a missense mutation where arginine at position 361 is substituted by cysteine. This specific residue lies within the MH2 domain, crucial for SMAD4's interaction with other SMAD proteins and its transcriptional activity. Mutations at this position, including p.R361C, have been implicated in several conditions. Notably, the p.R361C mutation has been associated with juvenile polyposis syndrome (JPS), characterized by multiple benign polyps in the gastrointestinal tract and an elevated risk of colorectal cancer. Functional studies indicate that this mutation disrupts SMAD4's ability to form heteromeric complexes with R-SMADs, thereby impairing canonical TGF-β signaling. Additionally, research suggests that the p.R361C mutation may enhance Wnt/β-catenin signaling, potentially contributing to tumorigenesis in colorectal cancer. Given the pathogenic nature of the p.R361C variant and its association with hereditary cancer syndromes, individuals harboring this mutation should undergo regular surveillance for early detection and management of potential malignancies. Further research is warranted to elucidate the full spectrum of clinical manifestations associated with this variant and to develop targeted therapeutic strategies. | <https://learn.mapmygenome.in/smad4>  <https://en.wikipedia.org/wiki/SMAD_%28protein%29>  <https://www.genecards.org/cgi-bin/carddisp.pl?gene=SMAD4>  <https://aacrjournals.org/mcr/article-abstract/19/5/823/672924/Colorectal-Cancer-Associated-Smad4-R361-Hotspot?redirectedFrom=fulltext>  <https://www.ncbi.nlm.nih.gov/clinvar/variation/24832/>  <https://www.ncbi.nlm.nih.gov/clinvar/variation/8543/>  <https://www.ncbi.nlm.nih.gov/clinvar/RCV001376609.14/>  <https://www.mskcc.org/cancer-care/patient-education/about-mutations-smad4-gene>  <https://www.ncbi.nlm.nih.gov/clinvar/RCV002311508.1/>  <https://medlineplus.gov/genetics/gene/smad4/>  <https://medlineplus.gov/download/genetics/gene/smad4.pdf>  <https://en.wikipedia.org/wiki/SMAD_%28protein%29>  <https://cancer.sanger.ac.uk/cosmic/mutation/overview?id=187736951>  <https://www.proteinatlas.org/ENSG00000141646-SMAD4>  <https://learn.mapmygenome.in/smad4> |
|  | Evaluation  **Summarization Accuracy & Completeness:**  **Hallucination Rate:**  **Readability & Usability:**  **Literature Coverage & Relevance:**  **Time Efficiency:** | |
