## Supplementary material for "Benchmarking generative AI tools for literature retrieval and summarization in genomic variant interpretation": SMARCA4 p.R1192C.docx

| **Tool** | **Summary** | **References** |
| --- | --- | --- |
| **1** | The SMARCA4 gene encodes BRG1, a critical ATPase subunit of the SWI/SNF chromatin remodeling complex, which modulates transcription by altering nucleosome positioning and accessibility of DNA to transcription factors. Germline or somatic mutations in SMARCA4 are linked to various malignancies, including rhabdoid tumor predisposition syndrome and non-small cell lung cancer, with emerging implications across a range of aggressive cancers. The variant p.R1192C results in an amino acid substitution at codon 1192 from arginine to cysteine, potentially impacting the protein's ATPase domain function or altering protein stability, though its pathogenic significance remains insufficiently characterized. At present, there are no peer-reviewed publications directly assessing the p.R1192C variant in SMARCA4, and its clinical significance has not been definitively established in population databases or functional assays. While SMARCA4-deficient tumors, particularly in the central nervous system and esophageal carcinoma, have shown distinct pathological and clinical features [1,2], no data currently links the p.R1192C variant specifically to these phenotypes. Due to the absence of variant-specific evidence, this change should be interpreted cautiously, and further functional studies and case reports are needed to clarify its clinical impact. | [1] [Morris et al., 2024 – SMARCA4-deficient central nervous system metastases](https://academic.oup.com/jnen/article-abstract/83/8/638/7660236) [2] [Cui et al., 2023 – Esophageal carcinoma with SMARCA4 mutation](https://www.sciencedirect.com/science/article/pii/S0344033823003928) |
|  | Evaluation  **Summarization Accuracy & Completeness:**  **Hallucination Rate:**  **Readability & Usability:**  **Literature Coverage & Relevance:**  **Time Efficiency:** | |
| **2** | The SMARCA4 gene encodes BRG1, a catalytic subunit of the SWI/SNF chromatin remodeling complex, which modulates chromatin structure to regulate gene expression, DNA repair, replication, and cell differentiation. BRG1 functions as a tumor suppressor, and its inactivation has been implicated in various cancers. The p.R1192C variant (c.3574C>T) results in an arginine-to-cysteine substitution at position 1192 within the ATPase helicase domain of BRG1, a region critical for ATP binding and chromatin remodeling activity. Functional studies have demonstrated that the R1192C mutation impairs chromatin remodeling and fails to compensate for SMARCA2 loss, indicating a loss-of-function effect . This variant has been identified in various cancers, including non-small cell lung carcinoma and Burkitt lymphoma . ClinVar classifies the germline occurrence of p.R1192C as a variant of uncertain significance for rhabdoid tumor predisposition syndrome 2 . Given its location in a conserved functional domain and demonstrated loss-of-function impact, the p.R1192C variant may contribute to tumorigenesis, but further studies are necessary to elucidate its clinical significance fully. | <https://medlineplus.gov/download/genetics/gene/smarca4.pdf>  [https://www.nature.com/articles/s41467-020-19402-8](https://www.nature.com/articles/s41467-020-19402-8?utm_source=chatgpt.com)  [https://www.oncokb.org/gene/SMARCA4/R1192C](https://www.oncokb.org/gene/SMARCA4/R1192C?utm_source=chatgpt.com)  <https://www.ncbi.nlm.nih.gov/clinvar/97738136/>  <https://academic.oup.com/jnen/article/83/8/638/7660236?rss=1>  <https://www.nature.com/articles/s41388-023-02773-9>  <https://www.ncbi.nlm.nih.gov/clinvar/97738136/>  <https://medlineplus.gov/genetics/gene/smarca4/>  <https://medlineplus.gov/download/genetics/gene/smarca4.pdf>  <https://www.nature.com/articles/s41467-020-19402-8>  <https://www.ncbi.nlm.nih.gov/gene/6597>  <https://pmc.ncbi.nlm.nih.gov/articles/PMC8643328/>  <https://pmc.ncbi.nlm.nih.gov/articles/PMC8524462/>  <https://pmc.ncbi.nlm.nih.gov/articles/PMC8831463/>  <https://pmc.ncbi.nlm.nih.gov/articles/PMC11296419/>  <https://www.uniprot.org/uniprotkb/P51532/entry>  <https://www.sciencedirect.com/science/article/abs/pii/S0169500220305596>  <https://www.genecards.org/cgi-bin/carddisp.pl?gene=SMARCA4>  <https://www.cell.com/cancer-cell/fulltext/S1535-6108%2824%2900055-2> |
|  | Evaluation  **Summarization Accuracy & Completeness:**  **Hallucination Rate:**  **Readability & Usability:**  **Literature Coverage & Relevance:**  **Time Efficiency:** | |
| **3** | The SMARCA4 gene, also known as BRG1, encodes a protein that is a central component of the SWI/SNF chromatin remodeling complex, which plays a crucial role in regulating gene expression by altering chromatin structure. This complex uses the energy from ATP hydrolysis to mobilize nucleosomes, thereby controlling access to DNA for processes such as transcription, replication, and repair. The SMARCA4 protein is essential for various cellular functions, including differentiation, proliferation, and DNA repair. Mutations in SMARCA4 have been implicated in multiple cancers and developmental disorders, highlighting its significance in maintaining cellular homeostasis. The p.R1192C variant in the SMARCA4 gene is a missense mutation that results in the substitution of arginine with cysteine at position 1192 in the protein sequence. This variant is located within the helicase_C domain of the protein, a region critical for its ATPase activity and chromatin remodeling function. The p.R1192C variant has been detected in various cancer types, including lung, ovarian, and kidney cancers, suggesting its potential role in tumorigenesis. Functional studies have shown that this variant can impair the ATPase activity of the SMARCA4 protein, leading to reduced chromatin remodeling and altered gene expression. The frequency of the p.R1192C variant is relatively low in the general population, but it has been identified in multiple cancer cohorts, indicating its relevance in cancer development. The classification of this variant according to the American College of Medical Genetics and Genomics (ACMG) guidelines is likely pathogenic, based on its impact on protein function and association with cancer. However, the exact mechanisms by which the p.R1192C variant contributes to cancer development and progression remain unclear. Further research is needed to elucidate the molecular consequences of this variant and its potential as a therapeutic target. | [[1]](https://www.nature.com/articles/s41467-020-19402-8) Nature Communications  [[2]](https://www.frontiersin.org/journals/immunology/articles/10.3389/fimmu.2021.762598/full) Frontiers  [[3]](https://www.ncbi.nlm.nih.gov/pmc/articles/PMC10932324/) PMC  [[4]](https://www.cell.com/cancer-cell/fulltext/S1535-6108(24)00055-2) Cancer Cell  [[5]](https://pubmed.ncbi.nlm.nih.gov/32777674/) PMC |
|  | Evaluation  **Summarization Accuracy & Completeness:**  **Hallucination Rate:**  **Readability & Usability:**  **Literature Coverage & Relevance:**  **Time Efficiency:** | |
| **4** | The genomic variant c.3574C>T p.R1192C rs1568509370 in the SMARCA4 gene results in the substitution of arginine with cysteine at the protein position 1192. This variant has been identified in the context of various cancer types, with a notable presence in lung cancer and ovarian cancer cases. The SMARCA4 gene encodes a protein that is a part of the SWI/SNF chromatin remodeling complex, which is involved in altering chromatin structure to regulate gene expression. The protein plays a critical role in various cellular processes, including differentiation, proliferation, and DNA repair.  The p.R1192C variant is located within the C-terminal helicase domain of the SMARCA4 protein, a region that is highly conserved and functionally significant. This domain is involved in ATP binding and DNA binding, which are essential for the chromatin remodeling activity of the protein [1]. The arginine residue at position 1192, which is mutated in the p.R1192C variant, is part of the so-called "Arg finger," a critical component of the ATP-binding pocket. Mutations in this region, including the p.R1192C variant, are predicted to alter the physiochemical properties of the protein, potentially affecting its charge and, consequently, its function [1].  Functional studies have shown that the p.R1192C variant is associated with a loss-of-function phenotype. In cellular models, the presence of this variant was unable to rescue the growth defect observed after SMARCA2 knockdown, suggesting that the p.R1192C variant impairs the ability of SMARCA4 to compensate for the loss of SMARCA2, another member of the SWI/SNF complex [1]. This finding is significant because SMARCA2 has been considered a therapeutic target for tumors with SMARCA4 mutations, and the ability of SMARCA4 mutants to compensate for SMARCA2 loss is relevant for the development of SMARCA2-targeting agents.  In clinical studies, the p.R1192C variant has been detected in multiple cancer cases. In lung cancer, it has been associated with reduced overall survival, although the statistical significance was marginal [1]. In ovarian cancer, the variant was identified in 16 cases, and its presence was associated with clinical outcomes in certain cancer types, such as worse prognosis in lung adenocarcinoma [2]. Additionally, the variant has been observed in the context of intratumor heterogeneity and resistance to EGFR-TKI treatment in lung cancer, where it was found alongside other mutations in a primary lung lesion that metastasized and acquired additional resistance mutations [3].  In a case report, the p.R1192C variant was identified in a patient with gallbladder cancer that metastasized to the ovary. The variant was part of a genomic signature that included other alterations, but no targetable genes were identified for therapeutic intervention [4].  In summary, the c.3574C>T p.R1192C rs1568509370 variant in the SMARCA4 gene is a missense mutation with functional evidence supporting a loss-of-function effect. It has been implicated in the pathogenesis of various cancers and may have prognostic implications. However, its clinical significance remains uncertain, as reflected by its current classification in ClinVar. Further research is needed to fully elucidate the clinical impact of this variant and its potential role in targeted therapy. | [[1]](https://pubmed.ncbi.nlm.nih.gov/33144586/) Fernando TM et al. (2020). "Functional characterization of SMARCA4 variants identified by targeted exome-sequencing of 131,668 cancer patients." Nature communications, 11(1)  [[2]](https://pubmed.ncbi.nlm.nih.gov/34675941/) Peng L et al. (2021). "A Pan-Cancer Analysis of SMARCA4 Alterations in Human Cancers." Frontiers in immunology, 12()  [[3]](https://pubmed.ncbi.nlm.nih.gov/32777674/) Kunimasa K et al. (2020). "Multiregional sequence revealed SMARCA4 R1192C mutant clones acquired EGFR C797S mutation in the metastatic site of an EGFR-mutated NSCLC patient." Lung cancer (Amsterdam, Netherlands), 148()  [[4]](https://pubmed.ncbi.nlm.nih.gov/37373946/) Grizzi G et al. (2023). "Krukenberg Tumor Related to Gallbladder Cancer in a Young Woman: A Case Report and Review of the Literature." Journal of personalized medicine, 13(6)  [[5]](https://pubmed.ncbi.nlm.nih.gov/26469284/) J. Masliah-Planchon et al. (2016). "SMARCA4‐Mutated Atypical Teratoid/Rhabdoid Tumor with Retained BRG1 Expression" Pediatric Blood & Cancer, 63 |
|  | Evaluation  **Summarization Accuracy & Completeness:**  **Hallucination Rate:**  **Readability & Usability:**  **Literature Coverage & Relevance:**  **Time Efficiency:** | |
| **5** | The SMARCA4 gene encodes the ATPase subunit of the SWI/SNF chromatin remodeling complex, which plays a crucial role in regulating gene expression by altering chromatin structure[2]. SMARCA4, also known as BRG1, is essential for various cellular processes, including transcriptional regulation and DNA repair[2]. Variants in SMARCA4 have been associated with several conditions, including rhabdoid tumor predisposition syndrome-2 (RTPS2) and Coffin-Siris syndrome[2]. In cancer, SMARCA4 acts primarily as a tumor suppressor, with alterations found in approximately 5-7% of human malignancies, particularly in small cell carcinoma of the ovary, hypercalcemic type (SCCOHT), and certain sarcomas[2]. The specific variant p.R1192C has not been directly referenced in the available literature, suggesting that its clinical significance and impact on SMARCA4 function remain unclear. Given the importance of SMARCA4 in chromatin remodeling and its role in cancer development, further investigation is recommended to determine the potential effects of this variant on gene function and disease risk. While SMARCA4 alterations can lead to loss of function or dominant-negative effects, the specific consequences of p.R1192C would require additional research for accurate classification and clinical interpretation[2][3]. | [1] <https://www.ncbi.nlm.nih.gov/pmc/articles/PMC9602240/>  [2] <https://pubmed.ncbi.nlm.nih.gov/34642211/>  [3] <https://pubmed.ncbi.nlm.nih.gov/38742308/>  [4] <https://pubmed.ncbi.nlm.nih.gov/30339381/>  [5] <https://www.ncbi.nlm.nih.gov/pmc/articles/PMC10003091/>  [6] <https://pubmed.ncbi.nlm.nih.gov/36813544/>  [7] <https://www.semanticscholar.org/paper/7e41fd9124dbdb73ccc3763abff4cbaaaa611c14>  [8] <https://www.ncbi.nlm.nih.gov/pmc/articles/PMC10106838/> |
|  | Evaluation  **Summarization Accuracy & Completeness:**  **Hallucination Rate:**  **Readability & Usability:**  **Literature Coverage & Relevance:**  **Time Efficiency:** | |
