## Supplementary material for "Benchmarking generative AI tools for literature retrieval and summarization in genomic variant interpretation": STK11 rs121913321.docx

| **Tool** | **Summary** | **References** |
| --- | --- | --- |
| **1** | The genomic variant c.841_842del p.Pro281AlafsTer3 rs121913321 in the STK11 gene is a frameshift deletion that results in a truncated protein with a premature termination codon. This variant leads to the loss of a significant portion of the kinase domain and the complete loss of the C-terminal regulatory domain of the STK11 protein [1]. The presence of this variant has been associated with Peutz-Jeghers syndrome (PJS), a condition characterized by mucocutaneous pigmentations and gastrointestinal hamartomatous polyps, with an increased risk of cancer development in various organs [2].  The STK11 gene, also known as LKB1, encodes a serine/threonine kinase that plays a crucial role in cell metabolism, cell cycle regulation, and apoptosis. It is ubiquitously expressed in both adult and fetal tissues and is involved in chromatin remodeling and cell polarization. The protein consists of an N-terminal nuclear localization signal domain, a highly conserved serine/threonine kinase domain, and a C-terminal regulatory domain. Mutations in the kinase domain can impair kinase activity, affecting cell growth, while mutations in the C-terminal region may disrupt cell polarization and are associated with cancer risk [2].  The variant rs121913321 has been reported in ClinVar with a total of 7 pathogenic submissions, indicating a consensus on its pathogenicity in relation to PJS. The variant's association with PJS phenotypes and its potential role in the disease's clinical manifestations have been documented, with a genotype-phenotype correlation suggested in some studies. This correlation may have implications for risk assessment and genetic counseling for individuals with PJS [2]. | [[1]](https://pubmed.ncbi.nlm.nih.gov/30092773/) Jiang YL et al. (2018). "The altered activity of P53 signaling pathway by STK11 gene mutations and its cancer phenotype in Peutz-Jeghers syndrome." BMC medical genetics, 19(1)  [[2]](https://pubmed.ncbi.nlm.nih.gov/28352323/) Zhang Y et al. (2017). "Correlation between genotype and phenotype in three families with Peutz-Jeghers Syndrome." Experimental and therapeutic medicine, 13(2)  [[3]](https://pubmed.ncbi.nlm.nih.gov/30458888/) Hu ZI et al. (2018). "Acquired resistance to immunotherapy in MMR-D pancreatic cancer." Journal for immunotherapy of cancer, 6(1)  [[4]](https://pubmed.ncbi.nlm.nih.gov/37377590/) Jiang LX et al. (2023). "Peutz-Jeghers syndrome without STK11 mutation may correlate with less severe clinical manifestations in Chinese patients." World journal of gastroenterology, 29(21)  [[5]](https://pubmed.ncbi.nlm.nih.gov/39062695/) Uliana V et al. (2024). "Phenotypic Expansion of Autosomal Dominant LZTR1-Related Disorders with Special Emphasis on Adult-Onset Features." Genes, 15(7)  [[6]](https://pubmed.ncbi.nlm.nih.gov/12865922/) Lim W et al. (2003). "Further observations on LKB1/STK11 status and cancer risk in Peutz-Jeghers syndrome." British journal of cancer, 89(2)  [[7]](https://pubmed.ncbi.nlm.nih.gov/28521409/) Lee SJ et al. (2017). "Evaluation of a novel approach to circulating tumor cell isolation for cancer gene panel analysis in patients with breast cancer." Oncology letters, 13(5)  [[8]](https://pubmed.ncbi.nlm.nih.gov/17319781/) Bartosova Z et al. (2007). "STK11/LKB1 germline mutations in the first Peutz-Jeghers syndrome patients identified in Slovakia." Neoplasma, 54(2)  [[9]](https://pubmed.ncbi.nlm.nih.gov/28938614/) Zhang J et al. (2017). "Phenformin enhances the therapeutic effect of selumetinib in KRAS-mutant non-small cell lung cancer irrespective of LKB1 status." Oncotarget, 8(35)  [[10]](https://pubmed.ncbi.nlm.nih.gov/34422335/) Devarakonda S et al. (2021). "A phase II study of everolimus in patients with advanced solid malignancies with TSC1, TSC2, NF1, NF2 or STK11 mutations." Journal of thoracic disease, 13(7)  [[11]](https://pubmed.ncbi.nlm.nih.gov/26917230/) Chen L et al. (2016). "A Sensitive NanoString-Based Assay to Score STK11 (LKB1) Pathway Disruption in Lung Adenocarcinoma." Journal of thoracic oncology : official publication of the International Association for the Study of Lung Cancer, 11(6)  [[12]](https://pubmed.ncbi.nlm.nih.gov/33534223/) Bennett JA et al. (2021). "A Distinctive Adnexal (Usually Paratubal) Neoplasm Often Associated With Peutz-Jeghers Syndrome and Characterized by STK11 Alterations (STK11 Adnexal Tumor): A Report of 22 Cases." The American journal of surgical pathology, 45(8)  [[13]](https://pubmed.ncbi.nlm.nih.gov/25375092/) Xiao Z et al. (2014). "Metformin and salinomycin as the best combination for the eradication of NSCLC monolayer cells and their alveospheres (cancer stem cells) irrespective of EGFR, KRAS, EML4/ALK and LKB1 status." Oncotarget, 5(24)  [[14]](https://pubmed.ncbi.nlm.nih.gov/28391030/) Xiao Z et al. (2017). "Metformin Triggers Autophagy to Attenuate Drug-Induced Apoptosis in NSCLC Cells, with Minor Effects on Tumors of Diabetic Patients." Neoplasia (New York, N.Y.), 19(5)  [[15]](https://pubmed.ncbi.nlm.nih.gov/36505399/) Lin J et al. (2022). "An EGFR L858R mutation identified in 1862 Chinese NSCLC patients can be a promising neoantigen vaccine therapeutic strategy." Frontiers in immunology, 13() |
|  | Evaluation  **Summarization Accuracy & Completeness:**  **Hallucination Rate:**  **Readability & Usability:**  **Literature Coverage & Relevance:**  **Time Efficiency:** | |
| **2** | The STK11 gene, also known as LKB1, encodes the serine/threonine kinase 11 enzyme, a crucial tumor suppressor involved in regulating cell polarity, metabolism, and apoptosis. Germline mutations in STK11 are the primary cause of Peutz-Jeghers syndrome (PJS), an autosomal dominant disorder characterized by mucocutaneous pigmentation, gastrointestinal hamartomatous polyps, and an elevated risk of various cancers. The specific variant rs121913321, corresponding to NM_000455.5(STK11):c.842dup (p.Leu282fs), is a duplication mutation leading to a frameshift and premature stop codon, resulting in a truncated, likely nonfunctional protein. This variant has been classified as pathogenic and is associated with hereditary cancer-predisposing syndromes, particularly PJS. It is absent in population databases such as gnomAD, indicating it is not a common polymorphism. The pathogenicity of this variant is supported by its predicted loss-of-function effect, consistent with the tumor suppressor role of STK11. Individuals with this mutation may present with typical PJS manifestations, including pigmented mucocutaneous lesions and gastrointestinal polyps, and face an increased risk of developing malignancies in organs such as the breast, pancreas, and lungs. Given the serious implications of this variant, genetic counseling and proactive surveillance for associated cancers are recommended for carriers. Further research is necessary to fully elucidate the clinical spectrum associated with this specific mutation and to optimize management strategies for affected individuals. | [1] [medlineplus.gov STK11 gene: MedlinePlus Genetics 1 maggio 2015 — The STK11 gene (also called LKB1) provides instructions for making an enzyme called serine/threonine kinase 11.](https://medlineplus.gov/genetics/gene/stk11/)  [2] [wjgnet.com Peutz-Jeghers syndrome without STK11 mutation may correlate with … 7 giugno 2023 Peutz-Jeghers syndrome without STK11 mutation may correlate with…](https://www.wjgnet.com/1007-9327/full/v29/i21/3302.htm)  [3] [ncbi.nlm.nih.gov NM_000455.5(STK11):c.842dup (p.Leu282fs) AND Hereditary … dbSNP: rs121913321; NCBI 1000 Genomes Browser: rs121913321;](https://www.ncbi.nlm.nih.gov/clinvar/RCV000492505)  [4] [ncbi.nlm.nih.gov ClinVar Genomic variation as it relates to human health - NCBI Loss-of-function variants in STK11 are known to be pathogenic (PMID: 15188174, 16287113).](https://www.ncbi.nlm.nih.gov/clinvar/variation/192227/)  [5] [wjgnet.com Peutz-Jeghers syndrome without STK11 mutation may correlate with ...](https://www.wjgnet.com/1007-9327/tables/v29/i21/3302.htm)  [6] [F6 Publishing [PDF] World Journal of - Gastroenterology - NET 7 giugno 2023 — STK11 is a tumor suppressor gene comprised of 433 amino acids with nine coding exons …](https://f6publishing.blob.core.windows.net/e1b1763a-3786-4b12-8aeb-4df2c91ff0f1/WJG-29-3302.pdf)  [7] [pmc.ncbi.nlm.nih.gov Peutz-Jeghers syndrome without STK11 mutation may correlate with …](https://pmc.ncbi.nlm.nih.gov/articles/PMC10292148/)  [8] [ClinVar Miner List of variants in gene STK11 reported as pathogenic by Women's …](https://clinvarminer.genetics.utah.edu/variants-by-gene/STK11/submitter/500026/pathogenic)  [9] [ClinVar Miner Submissions for variant NM_000455.5(STK11) - ClinVar Miner This variant has not been reported in the literature in individuals with STK11-related conditions.](https://clinvarminer.genetics.utah.edu/submissions-by-variant/NM_000455.5%28STK11%29%3Ac.841_842dup%20%28p.Leu282fs%29)  [10] [ClinVar Miner List of variants in gene STK11 reported as pathogenic for Peutz ...List of variants in gene STK11 reported as pathogenic for Peutz-Jeghers syndrome ...](https://clinvarminer.genetics.utah.edu/variants-by-gene/STK11/condition/Peutz-Jeghers%20syndrome/pathogenic)  [11] [Genopedia Genetic Variant rs121913321 \| Genopedia rs121913321 is a genetic variant on gene STK11. This variant is located on chromosome 19.](https://ng-genopedia-dev-web.azurewebsites.net/variant/121913321)  [12] [wjgnet.com Peutz-Jeghers syndrome without STK11 mutation may correlate with ...Peutz-Jeghers syndrome without STK11 mutation may correlate with less severe clinical manifestations in Chinese patients.](https://www.wjgnet.com/1007-9327/full/v29/i21/3302-T3.htm)  [13] [mgend.ncgm.go.jp chr19:1221319:C> Detail (hg19) (STK11) - Variant Detail : MGeND](https://mgend.ncgm.go.jp/variant/info/DELETION%7Chg19%7Cchr19%3A1221319-1221319%7CgC%3E?entrez_gene_id=6794&transcript_id=ENST00000326873.12)  [rs121913321 dbSNP; Genome: hg19; Position: chr19:1,221,319-1,221,319; Variant Type: snv](https://mgend.ncgm.go.jp/variant/info/DELETION%7Chg19%7Cchr19%3A1221319-1221319%7CgC%3E?entrez_gene_id=6794&transcript_id=ENST00000326873.12)  [14] [ScholarWorks Characterizing STK11 Variants of Unknown Significance Identified ...](https://scholarworks.uvm.edu/src/2020/healthscienceslinical/10/)  [Pathogenic, loss of function mutations in the STK11 gene corresponds with poorer survival and response to immunotherapy, the first line treatment for advanced- ...](https://scholarworks.uvm.edu/src/2020/healthscienceslinical/10/)  [15] [Ambrygen [PDF] Mutations in STK11 identified exclusively in individuals with clinical … Pathogenic mutations in STK11 are detected in an estimated 80% of individuals clinically diagnosed with Peutz Jeghers syndrome (PJS).](https://www.ambrygen.com/file/view/2674/Mutations%20in%20STK11%20identified%20exclusively%20in%20individuals%20with%20clinical%20histories%20suggestive%20of%20Peutz-Jeghers%20syndrome.pdf)  [16] [Malacards Peutz-Jeghers Syndrome (PJS) - MalaCards This syndrome is associated with growths in the gastrointestinal tract and a high risk of cancer.](https://www.malacards.org/card/peutz_jeghers_syndrome)  [20] [github.com falco/ref/filters/clinvar_00-latest.f.vcf at master - GitHub This is used to match alleles with other corresponding clinical (CLN) INFO tags.](https://github.com/tgac-vumc/falco/blob/master/ref/filters/clinvar_00-latest.f.vcf) |
|  | Evaluation  **Summarization Accuracy & Completeness:**  **Hallucination Rate:**  **Readability & Usability:**  **Literature Coverage & Relevance:**  **Time Efficiency:** | |
| **3** | The STK11 gene, also known as liver kinase B1 (LKB1), encodes serine/threonine kinase 11, a critical tumor suppressor protein involved in various cellular processes. STK11 plays a pivotal role in regulating cell growth, apoptosis, cell polarity, and energy metabolism by activating the AMP-activated protein kinase (AMPK) pathway and influencing the mTOR and HIF-1α signaling networks. Germline mutations in STK11 are associated with Peutz-Jeghers syndrome (PJS), an autosomal dominant condition characterized by the development of hamartomatous polyps in the gastrointestinal tract and an increased risk of various cancers, including breast, gastrointestinal, and lung cancers. Somatic mutations in STK11 are also frequently observed in lung adenocarcinoma and other cancer types, where they contribute to tumorigenesis and resistance to therapies, particularly immunotherapies targeting the PD-1 axis.  The STK11 rs121913321 variant is a specific genetic alteration within the STK11 gene. This variant is classified as pathogenic, as it results in a premature termination codon, leading to a truncated or absent protein product due to nonsense-mediated decay. The frequency of this variant is extremely low, with an allele frequency of 4.1e-06 in control populations, indicating its rarity. The clinical significance of this variant is substantial, as it is associated with the development of PJS and an increased risk of cancer. Individuals carrying this variant may exhibit the characteristic features of PJS, including the formation of hamartomatous polyps and mucocutaneous pigmentation. The loss of functional STK11 protein due to this variant can also contribute to the development of various cancers, particularly in the gastrointestinal tract and lungs.  The impact of the STK11 rs121913321 variant on cancer risk and therapeutic response highlights the importance of genetic testing and counseling for affected individuals and their families. Early identification of this variant can facilitate surveillance and management strategies to mitigate cancer risk and improve outcomes. However, the precise mechanisms by which this variant contributes to tumorigenesis and therapeutic resistance remain areas of active research. Ongoing studies and clinical trials are essential to elucidate these mechanisms and develop targeted therapies for individuals with STK11 mutations.  In conclusion, the STK11 rs121913321 variant is a rare and pathogenic alteration in the STK11 gene, associated with Peutz-Jeghers syndrome and an increased risk of cancer. Its clinical significance underscores the need for genetic testing and personalized management strategies for affected individuals. Further research is warranted to fully understand the molecular consequences of this variant and optimize therapeutic approaches for related cancers. | [1] ClinVar. NM_000455.5(STK11):c.842dup (p.Leu282fs) AND Peutz-Jeghers syndrome - ClinVar - NCBI. Available at: <https://www.ncbi.nlm.nih.gov/clinvar/RCV000474843/>  [2] MedlinePlus. STK11 gene: MedlinePlus Genetics. Available at: <https://medlineplus.gov/genetics/gene/stk11/>  [3] ScienceDirect. STK11 - an overview \| ScienceDirect Topics. Available at: <https://www.sciencedirect.com/topics/biochemistry-genetics-and-molecular-biology/stk11> |
|  | Evaluation  **Summarization Accuracy & Completeness:**  **Hallucination Rate:**  **Readability & Usability:**  **Literature Coverage & Relevance:**  **Time Efficiency:** | |
| **4** | The STK11 gene, also known as LKB1, provides instructions for making serine/threonine kinase 11, an enzyme that functions as a tumor suppressor by controlling cell growth and division[2](https://www.semanticscholar.org/paper/922654005d232329ae930f1e0436f9eb84b15156). This enzyme is involved in cell polarization, energy regulation, and apoptosis[2](https://www.semanticscholar.org/paper/922654005d232329ae930f1e0436f9eb84b15156). Germline loss-of-function mutations in STK11 are associated with Peutz-Jeghers syndrome (PJS), an autosomal dominant inherited disease characterized by mucocutaneous pigmentation and gastrointestinal hamartomatous polyps[5](https://www.ncbi.nlm.nih.gov/pmc/articles/PMC6107527/)[7](https://pubmed.ncbi.nlm.nih.gov/34479873/). However, some patients with PJS do not have STK11 mutations, suggesting other genetic factors may be involved[5](https://www.ncbi.nlm.nih.gov/pmc/articles/PMC6107527/). Somatic STK11 mutations are observed in a variety of sporadic cancers, particularly lung adenocarcinomas, where they occur in approximately 15% of cases[3](https://www.semanticscholar.org/paper/f0b7d618073d1eee9204981cb5d082e4ca2dd6de)[7](https://pubmed.ncbi.nlm.nih.gov/34479873/). The rs121913321 variant represents a specific alteration within the STK11 gene. Though the precise classification, nomenclature, and frequency of rs121913321 were not found in the literature, mutations in STK11 have been associated with worse prognoses in various malignancies and may predict poor response to immunotherapy in lung cancer[7](https://pubmed.ncbi.nlm.nih.gov/34479873/). At the cellular level, STK11 loss of function is linked to global DNA hypomethylation and S-adenosyl-methionine depletion in lung adenocarcinoma[8](https://pubmed.ncbi.nlm.nih.gov/34045189/). STK11 mutations can lead to resistance to immune checkpoint blockade and may impact treatment outcomes[3](https://www.semanticscholar.org/paper/f0b7d618073d1eee9204981cb5d082e4ca2dd6de). Some studies have focused on functionally characterizing STK11 mutations to understand their impact, and efforts are underway to classify non-annotated variants using functional screening approaches[3](https://www.semanticscholar.org/paper/f0b7d618073d1eee9204981cb5d082e4ca2dd6de). Further research is needed to fully elucidate the clinical implications of the rs121913321 variant and its potential role in cancer development and treatment response. | [1] <https://pmc.ncbi.nlm.nih.gov/articles/PMC6848484/>  [2] <https://www.semanticscholar.org/paper/STK11-Gene/922654005d232329ae930f1e0436f9eb84b15156>  [3] <https://www.semanticscholar.org/paper/Abstract-5584%3A-Experimental-%E2%80%98loss-of-function%E2%80%99-of-Ahronian-Shahagadkar/f0b7d618073d1eee9204981cb5d082e4ca2dd6de>  [4] <https://pmc.ncbi.nlm.nih.gov/articles/PMC6085611/>  [5] <https://pmc.ncbi.nlm.nih.gov/articles/PMC6107527/>  [6] <https://www.semanticscholar.org/paper/Association-of-cachexia-with-loss-of-function-in-Infante-Ahn/ab92e937ccd7e523e6842eb6537c9de4d01e5509>  [7] <https://pubmed.ncbi.nlm.nih.gov/34479873/>  [8] <https://pubmed.ncbi.nlm.nih.gov/34045189/> |
|  | Evaluation  **Summarization Accuracy & Completeness:**  **Hallucination Rate:**  **Readability & Usability:**  **Literature Coverage & Relevance:**  **Time Efficiency:** | |
| **5** | The STK11 gene (serine/threonine kinase 11), also known as LKB1, encodes a tumor suppressor kinase that plays a crucial role in cellular energy regulation, apoptosis, and growth inhibition via activation of the AMPK pathway. Mutations in STK11 are strongly associated with Peutz-Jeghers syndrome (PJS), an autosomal dominant disorder characterized by gastrointestinal polyposis and an elevated risk of malignancies. The variant STK11 rs121913321 is a pathogenic mutation that has been identified in individuals with PJS, contributing to disrupted kinase function and loss of tumor suppressive activity. The clinical implications of this variant include an increased predisposition to multiple cancers, including gastrointestinal, pancreatic, breast, and gynecological malignancies. While STK11 mutations are well-documented in PJS, some patients exhibit clinical manifestations without detectable mutations, suggesting additional genetic or epigenetic mechanisms that influence disease severity [1]. The classification of rs121913321 as pathogenic is based on its consistent association with PJS and tumorigenesis, reinforcing its clinical significance in hereditary cancer predisposition. However, the precise frequency of this specific variant remains unclear due to limited large-scale population studies. Further research is warranted to elucidate genotype-phenotype correlations and potential therapeutic interventions targeting the STK11 pathway [1][2]. | [1] [Peutz-Jeghers syndrome without STK11 mutation may correlate with less severe clinical manifestations in Chinese patients - PMC](https://pmc.ncbi.nlm.nih.gov/articles/PMC10292148/)  [2] [Microsoft Word - Thesis Proposal 3rd draft.docx](https://khazna.ku.ac.ae/ws/portalfiles/portal/6824392/file) |
|  | Evaluation  **Summarization Accuracy & Completeness:**  **Hallucination Rate:**  **Readability & Usability:**  **Literature Coverage & Relevance:**  **Time Efficiency:** | |
