## Supplementary material for "Benchmarking generative AI tools for literature retrieval and summarization in genomic variant interpretation": TET2 p.H1778R.docx

| **Tool** | **Summary** | **References** |
| --- | --- | --- |
| **1** | The genomic variant c.5333A>G p.H1778R rs62621450 in the TET2 gene is a missense mutation that results in the substitution of histidine with arginine at position 1778 of the TET2 protein. This variant has been studied in the context of myeloid malignancies and has been observed in patient samples with acute myeloid leukemia (AML) and myelodysplastic syndromes (MDS) [1]. The TET2 gene encodes a protein that is involved in the conversion of 5-methylcytosine (5-mC) to 5-hydroxymethylcytosine (5-hmC), a process that plays a critical role in DNA demethylation and, consequently, in the regulation of gene expression.  Molecular dynamics simulations have suggested that the H1778R variant can alter the global and local structure of the TET2 catalytic domain, potentially affecting its interaction with DNA and its enzymatic activity [1]. The variant was found to increase the compactness of the low complexity insert (LCI) region within the TET2 protein, which is thought to have regulatory roles in the TET gene family. This alteration in the LCI region could influence the catalytic activity of TET2 by enhancing its interactions with DNA and the cysteine-rich domain of the protein [1].  In a study of glioma and control brain tissue, the H1778R variant was identified in control samples, suggesting that it may occur in the general population [2]. However, the frequency of this variant in the healthy population and its potential role in disease pathogenesis remain unclear. In a benchmark sample for next-generation sequencing (NGS), the H1778R variant was detected with a median allelic frequency of 51.00% in two out of fifteen participants, indicating its presence in a heterozygous state [3].  The impact of the H1778R variant on TET2 expression and function has been evaluated in the context of myeloid neoplasms. In one study, a patient with this variant exhibited low TET2 protein expression and 5-hmC levels, which could be indicative of altered catalytic activity of the TET2 protein [1]. However, the variant has also been listed as benign in ClinVar, suggesting that its pathogenicity may not be fully established or that it may have variable penetrance or expressivity in different individuals.  In summary, the c.5333A>G p.H1778R rs62621450 variant in the TET2 gene is a missense mutation that has been associated with changes in the structure and function of the TET2 protein. While it has been detected in both patient and control populations, its clinical significance and contribution to disease phenotypes require further investigation. The TET2 protein's role in DNA demethylation underscores the importance of understanding how variants such as H1778R can impact gene regulation and the pathogenesis of myeloid malignancies [1][2][3][4][5]. | [[1]](https://pubmed.ncbi.nlm.nih.gov/37884893/) Seethy AA et al. (2023). "Alterations of the expression of TET2 and DNA 5-hmC predict poor prognosis in Myelodysplastic Neoplasms." BMC cancer, 23(1)  [[2]](https://pubmed.ncbi.nlm.nih.gov/26284134/) Kraus TF et al. (2015). "Genetic Characterization of Ten-Eleven-Translocation Methylcytosine Dioxygenase Alterations in Human Glioma." Journal of Cancer, 6(9)  [[3]](https://pubmed.ncbi.nlm.nih.gov/33138022/) Delcourt T et al. (2020). "NGS for (Hemato-) Oncology in Belgium: Evaluation of Laboratory Performance and Feasibility of a National External Quality Assessment Program." Cancers, 12(11)  [[4]](https://pubmed.ncbi.nlm.nih.gov/26984174/) Scopim-Ribeiro R et al. (2016). "Low Ten-eleven-translocation 2 (TET2) transcript level is independent of TET2 mutation in patients with myeloid neoplasms." Diagnostic pathology, 11()  [[5]](https://pubmed.ncbi.nlm.nih.gov/33643703/) Shaikh ARK et al. (2021). "TET2 mutations in acute myeloid leukemia: a comprehensive study in patients of Sindh, Pakistan." PeerJ, 9()  [[6]](https://pubmed.ncbi.nlm.nih.gov/28400619/) Lyu Y et al. (2017). "Dysfunction of the WT1-MEG3 signaling promotes AML leukemogenesis via p53-dependent and -independent pathways." Leukemia, 31(12)  [[7]](https://pubmed.ncbi.nlm.nih.gov/31453016/) Lobry C et al. (2019). "Analysis of TET2 mutations in paroxysmal nocturnal hemoglobinuria (PNH)." Experimental hematology & oncology, 8()  [[8]](https://pubmed.ncbi.nlm.nih.gov/27555360/) Oudejans CB et al. (2016). "Noncoding RNA-regulated gain-of-function of STOX2 in Finnish pre-eclamptic families." Scientific reports, 6() [SUPP]  [[9]](https://pubmed.ncbi.nlm.nih.gov/28882180/) Su D et al. (2017). "High performance of targeted next generation sequencing on variance detection in clinical tumor specimens in comparison with current conventional methods." Journal of experimental & clinical cancer research : CR, 36(1) [SUPP]  [[10]](https://pubmed.ncbi.nlm.nih.gov/30894629/) Stringer BW et al. (2019). "A reference collection of patient-derived cell line and xenograft models of proneural, classical and mesenchymal glioblastoma." Scientific reports, 9(1) [SUPP]  [[11]](https://pubmed.ncbi.nlm.nih.gov/27275012/) Guerreiro RJ et al. (2016). "Mutation of TBCK causes a rare recessive developmental disorder." Neurology. Genetics, 2(3) [SUPP]  [[12]](https://pubmed.ncbi.nlm.nih.gov/24348198/) Gemovic B et al. (2013). "Feature-based classification of amino acid substitutions outside conserved functional protein domains." TheScientificWorldJournal, 2013() [SUPP]  [[13]](https://pubmed.ncbi.nlm.nih.gov/28324520/) Rawal RM et al. (2015). "Tobacco habituated and non-habituated subjects exhibit different mutational spectrums in head and neck squamous cell carcinoma." 3 Biotech, 5(5) [SUPP]  [[14]](https://pubmed.ncbi.nlm.nih.gov/30256453/) Gaare JJ et al. (2018). "Rare genetic variation in mitochondrial pathways influences the risk for Parkinson's disease." Movement disorders : official journal of the Movement Disorder Society, 33(10) [SUPP]  [[15]](https://pubmed.ncbi.nlm.nih.gov/25352556/) Tang X et al. (2014). "The eSNV-detect: a computational system to identify expressed single nucleotide variants from transcriptome sequencing data." Nucleic acids research, 42(22) [SUPP] |
|  | Evaluation  **Summarization Accuracy & Completeness:**  **Hallucination Rate:**  **Readability & Usability:**  **Literature Coverage & Relevance:**  **Time Efficiency:** | |
| **2** | The TET2 gene encodes an enzyme belonging to the ten-eleven translocation (TET) family of dioxygenases that catalyze the conversion of 5-methylcytosine to 5-hydroxymethylcytosine, playing a critical role in DNA demethylation and epigenetic regulation of gene expression. TET2 is widely expressed, with a prominent function in hematopoietic stem and progenitor cells, where it acts as a tumor suppressor by regulating cellular proliferation and differentiation processes. Loss-of-function mutations in TET2 are among the most common somatic alterations in myeloid malignancies, including myelodysplastic syndromes, acute myeloid leukemia (AML), and other clonal hematopoietic disorders, and are considered driver mutations contributing to leukemogenesis. These mutations often result in decreased TET2 enzymatic activity, leading to aberrant hypermethylation of DNA, particularly at gene promoter regions, which disrupts normal hematopoietic differentiation and promotes malignant transformation. The specific variant p.H1778R in TET2 has been documented in the literature, although direct functional studies on this mutation are limited; however, given the mutation’s location within the catalytic domain, it is likely to impair enzymatic activity, contributing to the pathogenic mechanisms observed in TET2-associated hematologic diseases. Studies have demonstrated that TET2 mutations are associated with decreased gene expression and increased promoter methylation, which can influence disease prognosis and response to therapy. The variant’s classification remains uncertain without functional validation, but it is potentially pathogenic based on the gene’s mutation spectrum and its established role in epigenetic dysregulation in myeloid malignancies. Further research is warranted to elucidate the precise functional impact of p.H1778R and its contribution to disease phenotype, emphasizing the importance of integrating genomic, epigenetic, and clinical data for comprehensive interpretation. Overall, TET2 mutations, including the p.H1778R variant, are significant in the context of hematologic malignancies, with implications for diagnosis, prognosis, and targeted therapeutic strategies. The current evidence underscores the critical role of TET2 in maintaining epigenetic homeostasis and preventing malignant transformation in hematopoietic cells, with loss-of-function mutations serving as key drivers in myeloid neoplasms. | [1] <https://www.semanticscholar.org/paper/fe87ef3abd28cc3f430abc835b95ffd4cbd7557b>  [2] <https://pubmed.ncbi.nlm.nih.gov/38312186/>  [3] <https://www.semanticscholar.org/paper/6bde0f63f8b3882a670ad312719d385de0e87bdc>  [4] <https://pubmed.ncbi.nlm.nih.gov/39196062/>  [5] <https://www.ncbi.nlm.nih.gov/pmc/articles/PMC9898569/>  [6] <https://www.ncbi.nlm.nih.gov/pmc/articles/PMC10629367/>  [7] <https://www.semanticscholar.org/paper/b101631e5adc3d0270f9db6645c51b4597923658>  [8] <https://www.semanticscholar.org/paper/3f113a781590674e5ce052d1b597013b988c0886> |
|  | Evaluation  **Summarization Accuracy & Completeness:**  **Hallucination Rate:**  **Readability & Usability:**  **Literature Coverage & Relevance:**  **Time Efficiency:** | |
| **3** | The TET2 gene encodes a methylcytosine dioxygenase, an enzyme that catalyzes the conversion of methylcytosine to 5-hydroxymethylcytosine, a critical process in DNA demethylation. This enzyme plays a significant role in regulating gene expression by modifying the methylation status of DNA, particularly in hematopoietic cells. TET2 is involved in myelopoiesis, the process of producing myeloid cells, and its dysfunction is linked to various myeloproliferative disorders. The TET2 protein acts as a tumor suppressor, preventing uncontrolled cell growth and division, and its inactivation through mutations can lead to the development of hematological malignancies, including myelodysplastic syndromes (MDS) and acute myeloid leukemia (AML). Somatic mutations in TET2 are not inherited but arise spontaneously in cells, contributing to diseases characterized by abnormal blood cell production. The p.H1778R variant of the TET2 gene is a single nucleotide variant that results in an amino acid substitution from histidine to arginine at position 1778 in the TET2 protein. This variant has been identified in the context of hematological malignancies, although its specific impact on protein function and clinical outcomes is not well-documented in the available literature. The frequency of the p.H1778R variant in the general population is not widely reported, and its classification as benign or pathogenic is uncertain due to limited research. The variant's effect on TET2 function and its potential role in disease development remain areas of ongoing investigation. Further studies are needed to elucidate the clinical significance of the p.H1778R variant and its implications for patient prognosis and treatment. | [1] MedlinePlus Genetics. TET2 gene. Available at: <https://medlineplus.gov/genetics/gene/tet2/>  [2] ClinVar. VCV000135289.3 - TET2 gene variant H1778R. Available at: <https://www.ncbi.nlm.nih.gov/clinvar/variation/135289/>  [3] Rat Genome Database. TET2 gene. Available at: <https://rgd.mcw.edu/rgdweb/report/gene/main.html?id=1606790> |
|  | Evaluation  **Summarization Accuracy & Completeness:**  **Hallucination Rate:**  **Readability & Usability:**  **Literature Coverage & Relevance:**  **Time Efficiency:** | |
| **4** | No peer-reviewed literature currently exists specifically evaluating the TET2 variant p.H1778R. The TET2 gene encodes an enzyme involved in DNA demethylation through the oxidation of 5-methylcytosine to 5-hydroxymethylcytosine, contributing to epigenetic regulation and cellular differentiation, particularly in hematopoietic cells. Pathogenic variants in TET2 have been implicated in a variety of hematologic malignancies including myelodysplastic syndromes, chronic myelomonocytic leukemia, and acute myeloid leukemia, where loss-of-function alterations are frequently observed. The p.H1778R variant represents a missense change in the highly conserved catalytic domain of TET2, substituting histidine with arginine at position 1778; however, its clinical significance remains uncertain due to a lack of published functional or population data. No studies have yet assessed the biochemical impact of this specific substitution on TET2 enzymatic activity, substrate binding, or its downstream epigenetic effects. Additionally, this variant is not reported with significant frequency in population databases such as gnomAD, suggesting it may be rare or private, though the absence of data precludes definitive classification. In silico predictive tools might suggest a deleterious effect due to the conservation and position within a critical domain, but such predictions lack validation in this context. Given these limitations, the p.H1778R variant in TET2 should be categorized as a Variant of Uncertain Significance (VUS) according to ACMG/AMP guidelines, pending further investigation through functional studies or association with clinical phenotypes. Clinicians and geneticists are advised to consider this variant in the broader context of other somatic or germline mutations and clinical presentation. Absence of direct peer-reviewed references specific to this variant necessitates caution in interpretation and underscores the need for future research. | No references available |
|  | Evaluation  **Summarization Accuracy & Completeness:**  **Hallucination Rate:**  **Readability & Usability:**  **Literature Coverage & Relevance:**  **Time Efficiency:** | |
| **5** | The TET2 gene encodes the ten-eleven translocation 2 protein, a methylcytosine dioxygenase that catalyzes the conversion of 5-methylcytosine to 5-hydroxymethylcytosine, playing a crucial role in active DNA demethylation and epigenetic regulation. TET2 is essential for normal hematopoiesis, acting as a tumor suppressor by regulating gene expression involved in cell proliferation and differentiation. Mutations in TET2 are frequently observed in myeloid malignancies, including myelodysplastic syndromes, myeloproliferative neoplasms, and acute myeloid leukemia. The p.H1778R variant (c.5333A>G) results in a histidine-to-arginine substitution at position 1778. This missense variant has been identified in both individuals with myeloid disorders and healthy controls, suggesting it may represent a benign polymorphism. Notably, p.H1778R is located outside the conserved catalytic domain of TET2, and its functional impact remains uncharacterized. Current evidence does not support a pathogenic role for this variant; however, due to limited functional studies, its clinical significance is uncertain. Further investigation is warranted to elucidate the potential effects of p.H1778R on TET2 function and its contribution to disease pathogenesis. | <https://www.ncbi.nlm.nih.gov/gene/54790>  <https://pmc.ncbi.nlm.nih.gov/articles/PMC10601240/>  <https://www.nature.com/articles/s42003-020-01391-5>  <https://www.ahajournals.org/doi/10.1161/CIRCULATIONAHA.120.046155>  <https://medlineplus.gov/genetics/gene/tet2/>  <https://www.ncbi.nlm.nih.gov/clinvar/RCV000122104.1/>  <https://medlineplus.gov/download/genetics/gene/tet2.pdf>  <https://ckb-core.genomenon.com/geneVariant/show?geneVariantId=18338>  <https://en.wikipedia.org/wiki/Tet_methylcytosine_dioxygenase_2>  <https://ashpublications.org/blood/article/118/14/3803/28728/TET2-mutation-is-an-unfavorable-prognostic-factor>  <https://www.researchgate.net/figure/Structure-of-TET2-and-the-effect-of-H1778R-variant-a-Gene-structure-of-full-length_fig4_375083156>  <https://www.researchgate.net/figure/Sequencing-of-TET2-gene-a-TET2-variants-c5060-5061delpQ1687fsX3-cA5333GpH1778R_fig3_375083156>  <https://www.jcancer.org/v06p0832.htm>  <https://www.frontiersin.org/journals/cell-and-developmental-biology/articles/10.3389/fcell.2021.665897/full>  <https://omim.org/entry/612839>  <https://clinvarminer.genetics.utah.edu/variants-by-gene/TET2/significance/any>  <https://www.genecards.org/cgi-bin/carddisp.pl?gene=TET2>  <https://scholar.ufs.ac.za/bitstreams/886ff136-2d28-4a7a-9afa-e1e965919c73/download> |
|  | Evaluation  **Summarization Accuracy & Completeness:**  **Hallucination Rate:**  **Readability & Usability:**  **Literature Coverage & Relevance:**  **Time Efficiency:** | |
