## Supplementary material for "Benchmarking generative AI tools for literature retrieval and summarization in genomic variant interpretation": TOP1 p.G359E.docx

| **Tool** | **Summary** | **References** |
| --- | --- | --- |
| **1** | The genomic variant c.1076G>A, resulting in the amino acid substitution p.G359E in the TOP1 gene, has been identified as a mutation associated with resistance to antibody-drug conjugates (ADCs) in patients with metastatic breast cancer (MBC). TOP1, or DNA topoisomerase I, is an enzyme that plays a critical role in DNA replication and transcription by inducing reversible single-strand breaks in the DNA to alleviate torsional strain during these processes.  The p.G359E variant was found in 12.9% of MBC patients at the time of disease progression on ADC treatment, which is significantly higher than the incidence observed in MBC patients not treated with ADCs (0.7%) and in the general cancer population as reported in The Cancer Genome Atlas (0.5%) [1]. This suggests that the p.G359E variant is selected for under the pressure of ADC treatment, indicating a potential mechanism of acquired resistance.  Functional characterization of the p.G359E variant revealed that the mutant protein exhibits reduced enzymatic activity and attenuated covalent DNA binding. Moreover, the variant confers resistance to the TOP1 inhibitor ADC payloads SN38 and Deruxtecan, which are commonly used in the treatment of MBC [1]. The presence of this mutation was also associated with clinical cross-resistance, as evidenced by a significantly shorter median duration of response to a second ADC following the emergence of the mutation.  Given these findings, the c.1076G>A p.G359E variant in the TOP1 gene may serve as a biomarker for resistance to ADC therapy in MBC, and its identification in patients could have implications for the management of their treatment, including the potential sequential use of ADCs [1]. Further research is needed to optimize biomarkers for ADC resistance and to improve ADC payload design to overcome resistance mechanisms mediated by TOP1 mutations. | [[1]](https://pubmed.ncbi.nlm.nih.gov/39745368/) Abelman RO et al. (2025). "TOP1 Mutations and Cross-Resistance to Antibody-Drug Conjugates in Patients with Metastatic Breast Cancer." Clinical cancer research : an official journal of the American Association for Cancer Research, ()  [[2]](https://aacrjournals.org/cancerres/article/84/6_Supplement/3888/740394/Abstract-3888-TOP1-mutations-mediate-cross) R. Abelman et al. (2024). "Abstract 3888: TOP1 mutations mediate cross resistance to ADCs in metastatic breast cancer" Cancer Research, |
|  | Evaluation  **Summarization Accuracy & Completeness:**  **Hallucination Rate:**  **Readability & Usability:**  **Literature Coverage & Relevance:**  **Time Efficiency:** | |
| **2** | The p.G359E variant in the *TOP1* gene does not currently appear in the peer-reviewed literature, and there are no published studies specifically evaluating its pathogenicity, clinical significance, or frequency in the population. *TOP1* (DNA topoisomerase I) encodes an essential nuclear enzyme that relaxes supercoiled DNA during replication and transcription by inducing transient single-strand breaks, a function critical for maintaining genomic stability. It is a well-established target of chemotherapeutic agents such as camptothecin and its derivatives, which exploit the enzyme’s catalytic mechanism to induce cytotoxicity in proliferating cells. Pathogenic alterations in *TOP1* have been implicated in cancer biology, primarily through mechanisms of drug resistance or genomic instability; however, missense variants like p.G359E remain poorly characterized. This specific substitution replaces a conserved glycine at position 359 with glutamic acid, which may potentially affect protein structure or interaction with DNA or drugs, but no functional assays or clinical correlations are available in the literature to confirm this hypothesis. In the absence of variant-specific data, its clinical interpretation remains uncertain, and further investigation through in vitro studies or cancer cohort sequencing is warranted to assess its potential role in tumor biology or therapy response. No direct peer-reviewed publications currently exist on TOP1 p.G359E [no references available]. | No references available |
|  | Evaluation  **Summarization Accuracy & Completeness:**  **Hallucination Rate:**  **Readability & Usability:**  **Literature Coverage & Relevance:**  **Time Efficiency:** | |
| **3** | The TOP1 gene encodes DNA topoisomerase I, an essential nuclear enzyme that modulates DNA topology by introducing transient single-strand breaks to relieve torsional strain during replication, transcription, and chromatin remodeling. The p.G359E variant denotes a missense substitution at codon 359, replacing glycine with glutamic acid. This variant has been identified in patients with metastatic breast cancer who developed resistance to antibody–drug conjugates (ADCs), specifically those targeting topoisomerase I, such as sacituzumab govitecan. In a study involving 31 patients, the p.G359E mutation was detected in 12.9% of cases (4 patients) at the time of disease progression on ADC therapy, suggesting a potential role in acquired resistance mechanisms. However, the precise functional impact of the G359E substitution on TOP1 enzymatic activity, DNA binding, or drug interaction has not been elucidated. No experimental studies to date have characterized the biochemical or structural consequences of this mutation, and its clinical significance remains uncertain. Given the limited data, the p.G359E variant should be considered of uncertain significance, and further functional analyses are warranted to determine its role in therapy resistance and potential implications for treatment strategies citeturn0search0. | [https://en.wikipedia.org/wiki/Topoisomerase_inhibitor](https://en.wikipedia.org/wiki/Topoisomerase_inhibitor?utm_source=chatgpt.com)  [https://www.researchgate.net/publication/387672413_TOP1_Mutations_and_Cross-Resistance_to_Antibody-Drug_Conjugates_in_Patients_with_Metastatic_Breast_Cancer](https://www.researchgate.net/publication/387672413_TOP1_Mutations_and_Cross-Resistance_to_Antibody-Drug_Conjugates_in_Patients_with_Metastatic_Breast_Cancer?utm_source=chatgpt.com)  [https://discovery.researcher.life/topic/resistance-in-metastatic-breast-cancer/2747714?page=1&topic_name=Resistance+In+Metastatic+Breast+Cancer](https://discovery.researcher.life/topic/resistance-in-metastatic-breast-cancer/2747714?page=1&topic_name=Resistance+In+Metastatic+Breast+Cancer&utm_source=chatgpt.com)  <https://www.ncbi.nlm.nih.gov/gene/7150>  <https://www.ncbi.nlm.nih.gov/gene/854156>  <https://aacrjournals.org/clincancerres/article-pdf/doi/10.1158/1078-0432.CCR-24-2771/3529791/ccr-24-2771.pdf>  <https://www.nature.com/articles/s41580-022-00452-3>  <https://aacrjournals.org/cancerres/article/67/18/8752/533560/Nonclassic-Functions-of-Human-Topoisomerase-I>  <https://www.genecards.org/cgi-bin/carddisp.pl?gene=TOP1>  <https://www.researchgate.net/publication/387672413_TOP1_Mutations_and_Cross-Resistance_to_Antibody-Drug_Conjugates_in_Patients_with_Metastatic_Breast_Cancer>  <https://www.researchgate.net/scientific-contributions/Ryan-B-Corcoran-16368067>  <https://discovery.researcher.life/topic/mutant-metastatic-breast-cancer/19152082?page=1&topic_name=Mutant+Metastatic+Breast+Cancer>  <https://www.sciencedirect.com/topics/biochemistry-genetics-and-molecular-biology/top1>  <https://www.sciencedirect.com/science/article/pii/S1672022916300389>  <https://www.cell.com/cms/10.1016/j.ccell.2018.06.005/attachment/75e388db-d6cb-4579-a49d-706d0069c863/mmc6.xlsx>  <https://www.yeastgenome.org/locus/s000005366>  <https://omim.org/entry/126420>  <https://www.pubmedplus.cn/?q=%22clin+cancer+res%22%5Bjournal%5D> |
|  | Evaluation  **Summarization Accuracy & Completeness:**  **Hallucination Rate:**  **Readability & Usability:**  **Literature Coverage & Relevance:**  **Time Efficiency:** | |
| **4** | The TOP1 gene, located on chromosome 20, encodes DNA topoisomerase I, an enzyme crucial for DNA replication, transcription, and repair. TOP1 catalyzes the transient breaking and rejoining of a single DNA strand, allowing strands to pass through one another and altering DNA topology. This process is essential for relaxing supercoiled DNA during replication and transcription, ensuring genomic stability. TOP1 is involved in various cellular pathways, including RNA binding and core promoter sequence-specific DNA binding, and its dysregulation has been linked to diseases such as hematologic cancer and spinocerebellar ataxia with axonal neuropathy. The enzyme's activity is regulated by post-translational modifications like sumoylation, which partitions TOP1 between the nucleoli and nucleoplasm. TOP1's role in DNA repair and its interaction with other DNA metabolism proteins suggest a potential involvement in human aging, though further studies are needed to confirm this hypothesis. The p.G359E variant of TOP1 has been identified in metastatic breast cancer patients, where it is associated with resistance to antibody-drug conjugates (ADCs) containing topoisomerase-I inhibitor payloads. This variant was found in 6.0% of patients at the time of disease progression on ADC treatment, compared to a frequency of 0.5% in primary breast cancer. The clinical impact of the p.G359E variant includes cross-resistance to ADCs, defined as progressive disease or short treatment duration on a second ADC. The exact molecular mechanisms by which p.G359E confers resistance are not fully understood, and further research is needed to elucidate its functional consequences and potential therapeutic implications. The frequency and impact of the p.G359E variant highlight the need for personalized treatment strategies in metastatic breast cancer, taking into account specific genetic mutations that may influence drug response. | no references available |
|  | Evaluation  **Summarization Accuracy & Completeness:**  **Hallucination Rate:**  **Readability & Usability:**  **Literature Coverage & Relevance:**  **Time Efficiency:** | |
| **5** | The TOP1 gene encodes DNA topoisomerase I, a nuclear enzyme essential for maintaining DNA topology by inducing reversible single-strand breaks to relieve torsional strain during DNA replication, transcription, and chromatin remodeling. This enzyme is classified as a type IB topoisomerase and is critical for normal cellular function and genome stability. Variants in TOP1 can influence cellular responses to DNA damage and sensitivity to topoisomerase-targeting chemotherapeutics. The p.G359E variant involves a missense substitution at amino acid position 359, replacing glycine with glutamic acid. This residue is located within the core domain of the enzyme, which is involved in DNA binding and catalysis; however, specific functional studies on p.G359E are lacking in the literature, and no peer-reviewed publications directly characterize its biochemical or clinical impact. The variant is not reported as a common polymorphism in large population databases, suggesting it may be rare or novel. Given the critical role of TOP1 in DNA relaxation and its targeting by anticancer drugs such as camptothecins, alterations in its sequence could potentially affect enzyme activity, drug binding, or cellular sensitivity to DNA damage, but this remains speculative without direct evidence. Existing studies on TOP1 polymorphisms have primarily focused on other variants and their associations with drug response or cancer susceptibility, with no conclusive data linking p.G359E to disease phenotypes or altered protein function. Therefore, the clinical significance of the p.G359E variant is currently uncertain, and further functional assays and clinical correlation studies are recommended to elucidate its potential pathogenicity or pharmacogenomic relevance. In summary, TOP1 is a vital gene encoding DNA topoisomerase I, with p.G359E representing a rare missense variant of unknown significance requiring additional research for definitive interpretation. No direct peer-reviewed references describing the p.G359E variant were identified in the literature. | 1. <https://www.ncbi.nlm.nih.gov/pmc/articles/PMC5641335/> 2. <https://www.ncbi.nlm.nih.gov/pmc/articles/PMC2360536/> 3. <https://www.semanticscholar.org/paper/4e6bce676f0c2a89ebb79633d082ba834eefc43a> 4. <https://www.ncbi.nlm.nih.gov/pmc/articles/PMC4694935/> 5. <https://www.ncbi.nlm.nih.gov/pmc/articles/PMC7456420/> |
|  | Evaluation  **Summarization Accuracy & Completeness:**  **Hallucination Rate:**  **Readability & Usability:**  **Literature Coverage & Relevance:**  **Time Efficiency:** | |
