## Supplementary material for "Benchmarking generative AI tools for literature retrieval and summarization in genomic variant interpretation": TP53 c.11C_T.docx

| **Tool** | **Summary** | **References** |
| --- | --- | --- |
| **1** | The genomic variant c.11C>T p.Pro4Leu rs878854064 is located on the TP53 gene, which is a well-known tumor suppressor gene encoding the p53 protein. The p53 protein plays a critical role in maintaining genomic stability by regulating the cell cycle, DNA repair, apoptosis, and cellular senescence in response to cellular stress. Mutations in TP53 are associated with a variety of human cancers and can lead to the loss of normal p53 function, contributing to tumorigenesis.  The c.11C>T p.Pro4Leu variant results in the substitution of a proline (Pro) with a leucine (Leu) at the fourth amino acid position of the p53 protein. This missense variant has been identified in multiple studies, with varying interpretations of its clinical significance. In one study, the variant was detected in a case of thin melanoma (TM) but was not associated with metastatic disease or other pathological features such as Breslow thickness or tumor-infiltrating lymphocytes (TIL) grade [4]. This suggests that the variant may not have a significant impact on the aggressiveness of the melanoma in this context.  In another study, the variant was interpreted as a variant of uncertain significance (VUS) and was not reported according to the institutional policy, which may reflect a conservative approach to variant reporting and the need for further evidence to clarify its clinical relevance [3]. The variant was also mentioned in a study that did not provide direct evidence of its functional impact but discussed the importance of TP53 mutations and polymorphisms in cancer susceptibility, highlighting the complexity of interpreting TP53 variants [5].  Given the conflicting interpretations and the presence of both benign and uncertain significance submissions in ClinVar, it is clear that the clinical significance of the c.11C>T p.Pro4Leu rs878854064 variant remains uncertain. Further functional studies and clinical correlation are needed to determine the impact of this variant on p53 function and its role in disease pathogenesis. The presence of this variant in the context of cancer should be interpreted with caution, and it may be considered as part of a broader genetic assessment rather than a definitive marker of disease. | [[1]](https://pubmed.ncbi.nlm.nih.gov/27640185/) Faden DL et al. (2016). "Targeted next-generation sequencing of TP53 in oral tongue carcinoma from non-smokers." Journal of otolaryngology - head & neck surgery = Le Journal d'oto-rhino-laryngologie et de chirurgie cervico-faciale, 45(1)  [[2]](https://pubmed.ncbi.nlm.nih.gov/31341169/) Zhang Y et al. (2019). "Deep single-cell RNA sequencing data of individual T cells from treatment-naive colorectal cancer patients." Scientific data, 6(1)  [[3]](https://pubmed.ncbi.nlm.nih.gov/35935605/) Pandzic T et al. (2022). "Five Percent Variant Allele Frequency Is a Reliable Reporting Threshold for TP53 Variants Detected by Next Generation Sequencing in Chronic Lymphocytic Leukemia in the Clinical Setting." HemaSphere, 6(8)  [[4]](https://pubmed.ncbi.nlm.nih.gov/30181807/) Richetta AG et al. (2018). "Metastases risk in thin cutaneous melanoma: prognostic value of clinical-pathologic characteristics and mutation profile." Oncotarget, 9(63)  [[5]](https://pubmed.ncbi.nlm.nih.gov/29348863/) Zhang A et al. (2017). "No association between TP53 Arg72Pro polymorphism and ovarian cancer risk: evidence from 10113 subjects." Oncotarget, 8(68)  [[6]](https://pubmed.ncbi.nlm.nih.gov/19424414/) Noureddine MA et al. (2009). "Probing the functional impact of sequence variation on p53-DNA interactions using a novel microsphere assay for protein-DNA binding with human cell extracts." PLoS genetics, 5(5)  [[7]](https://pubmed.ncbi.nlm.nih.gov/23785431/) Yuan RH et al. (2013). "S100P expression is a novel prognostic factor in hepatocellular carcinoma and predicts survival in patients with high tumor stage or early recurrent tumors." PloS one, 8(6)  [[8]](https://pubmed.ncbi.nlm.nih.gov/22427977/) Cai Y et al. (2012). "CIBZ, a novel BTB domain-containing protein, is involved in mouse spinal cord injury via mitochondrial pathway independent of p53 gene." PloS one, 7(3)  [[9]](https://pubmed.ncbi.nlm.nih.gov/19536795/) Glick J et al. (2009). "The influence of cytosine methylation on the chemoselectivity of benzo[a]pyrene diol epoxide-oligonucleotide adducts determined using nanoLC/MS/MS." Journal of mass spectrometry : JMS, 44(8)  [[10](https://ashpublications.org/blood/article/124/21/5570/101013/Targeted-Exome-Sequencing-Identifies-Novel)] Nader I. Al-Dewik et al. (2014). "Targeted Exome Sequencing Identifies Novel Mutations in Familial Myeloproliferative Neoplasms Patients in the State of Qatar" Blood, 124  [[11]](https://pubmed.ncbi.nlm.nih.gov/23844043/) Huang HC et al. (2013). "UVB irradiation regulates ERK1/2- and p53-dependent thrombomodulin expression in human keratinocytes." PloS one, 8(7)  [[12]](https://pubmed.ncbi.nlm.nih.gov/24121268/) Sparks A et al. (2014). "The degradation of p53 and its major E3 ligase Mdm2 is differentially dependent on the proteasomal ubiquitin receptor S5a." Oncogene, 33(38)  [[13]](https://pubmed.ncbi.nlm.nih.gov/30886117/) Quinn EA et al. (2019). "From uncertainty to pathogenicity: clinical and functional interrogation of a rare TP53 in-frame deletion." Cold Spring Harbor molecular case studies, 5(4) [SUPP]  [[14]](https://pubmed.ncbi.nlm.nih.gov/23258702/) Guha M et al. (2013). "Differential strand separation at critical temperature: a minimally disruptive enrichment method for low-abundance unknown DNA mutations." Nucleic acids research, 41(3) [SUPP]  [[15]](https://pubmed.ncbi.nlm.nih.gov/33585199/) Bannon SA et al. (2020). "Next-Generation Sequencing of DDX41 in Myeloid Neoplasms Leads to Increased Detection of Germline Alterations." Frontiers in oncology, 10() [SUPP] |
|  | Evaluation  **Summarization Accuracy & Completeness:**  **Hallucination Rate:**  **Readability & Usability:**  **Literature Coverage & Relevance:**  **Time Efficiency:** | |
| **2** | The TP53 gene, located on chromosome 17p13.1, encodes the tumor suppressor protein p53, which plays a critical role in maintaining genomic stability and preventing tumor formation. Known as the "guardian of the genome," p53 regulates cellular responses to diverse stresses, including DNA damage, oncogene activation, and hypoxia. It functions primarily as a transcription factor, activating the expression of genes involved in cell cycle arrest, apoptosis, DNA repair, and senescence. The importance of p53 is underscored by its frequent mutation in human cancers, with over 50% of all tumors harboring TP53 mutations. These mutations often impair p53's tumor suppressor function and can confer oncogenic properties, contributing to tumor initiation and progression.  The variant c.11C>T in the TP53 gene results in a nonsense mutation, leading to a premature stop codon and a truncated, non-functional p53 protein. This variant is classified as pathogenic according to the American College of Medical Genetics and Genomics (ACMG) guidelines. The loss of functional p53 due to this variant compromises its ability to regulate cell cycle control, DNA repair, and apoptosis, thereby increasing the risk of cancer development. Individuals carrying this variant may exhibit a higher predisposition to various cancers, including early-onset breast cancer, sarcomas, brain tumors, and adrenocortical carcinomas, which are characteristic of Li-Fraumeni syndrome (LFS).  The frequency of the c.11C>T variant in the general population is not well-documented, likely due to its rarity and the severe phenotypic consequences associated with it. However, its presence in families with LFS highlights its significance in cancer predisposition syndromes. The impact of this variant is profound, as it abolishes the protective functions of p53, leading to unchecked cell proliferation and genomic instability.  In summary, the c.11C>T variant in TP53 is a pathogenic mutation that results in a non-functional p53 protein, significantly increasing cancer risk. Its classification and impact underscore the critical role of p53 in tumor suppression and the importance of genetic testing for individuals with a family history of LFS or related cancers. | [1] Tumor-Suppressor Functions of the TP53 Pathway - PMC. [1](https://pmc.ncbi.nlm.nih.gov/articles/PMC4852799/)  [2] Targeting p53 pathways: mechanisms, structures and advances in therapy \| Signal Transduction and Targeted Therapy. [2](https://www.nature.com/articles/s41392-023-01347-1)  [3] p53 signaling in cancer progression and therapy \| Cancer Cell International \| Full Text. [3](https://cancerci.biomedcentral.com/articles/10.1186/s12935-021-02396-8)  [4] TP53 germline pathogenic variants in modern humans were likely originated during recent human history \| NAR Cancer \| Oxford Academic. [4](https://academic.oup.com/narcancer/article/5/3/zcad025/7192725)  Specifications of the ACMG/AMP variant interpretation guidelines for germline TP53 variants - PubMed. [5](https://pubmed.ncbi.nlm.nih.gov/33300245/) |
|  | Evaluation  **Summarization Accuracy & Completeness:**  **Hallucination Rate:**  **Readability & Usability:**  **Literature Coverage & Relevance:**  **Time Efficiency:** | |
| **3** | The TP53 gene encodes the tumor protein p53, a critical transcription factor involved in cellular response to DNA damage, cell cycle regulation, apoptosis, and genomic stability. TP53 mutations are frequently implicated in various malignancies, with certain variants conferring a high risk for cancer development. The c.11C>T variant results in a missense or nonsense change depending on transcript context, potentially disrupting the p53 function. This variant's pathogenicity remains uncertain, as no direct peer-reviewed studies explicitly characterize its clinical significance. Although TP53 mutations are prevalent in myeloid neoplasms and solid tumors, including melanoma and chronic lymphocytic leukemia, current research does not establish a direct oncogenic role for c.11C>T. Population frequency data for this variant is not well documented, and its classification as pathogenic, benign, or a variant of uncertain significance (VUS) remains unresolved. Given the absence of literature directly associating c.11C>T with disease, further functional studies and clinical correlation are necessary to determine its impact. Genetic counseling and additional molecular analyses, including loss-of-function assessments and co-segregation studies, are recommended for individuals harboring this variant to clarify potential clinical relevance. | [Quesada et al., 2019](https://onlinelibrary.wiley.com/doi/abs/10.1002/ajh.25486),  [Pandzic et al., 2022](https://journals.lww.com/hemasphere/fulltext/2022/08000/Five_Percent_Variant_Allele_Frequency_Is_a.9.aspx). |
|  | Evaluation  **Summarization Accuracy & Completeness:**  **Hallucination Rate:**  **Readability & Usability:**  **Literature Coverage & Relevance:**  **Time Efficiency:** | |
| **4** | The TP53 gene encodes the p53 tumor suppressor protein, which plays a critical role in genome stability by regulating DNA repair, cell cycle arrest, and apoptosis in response to cellular stress[4](https://www.ncbi.nlm.nih.gov/pmc/articles/PMC10742156/)[6](https://www.semanticscholar.org/paper/63eb7a38bc1d08d3860941c7ebcb93cb470eb17e). The c.11C>T variant (GRCh38) represents a missense mutation in exon 1 of TP53, resulting in a p.(Pro4Leu) amino acid substitution within the transactivation domain, which is essential for transcriptional regulation. This variant is classified as pathogenic by ClinVar (RCV003324917.1) due to its location in a functional domain and in silico predictions of deleterious effects, though direct functional evidence remains limited. TP53 mutations occur in >50% of human cancers, with missense variants constituting ~60% of all mutations, though c.11C>T itself appears rare based on current databases[6](https://www.semanticscholar.org/paper/63eb7a38bc1d08d3860941c7ebcb93cb470eb17e)[7](https://www.ncbi.nlm.nih.gov/pmc/articles/PMC11788706/). While no peer-reviewed studies specifically address c.11C>T, analogous N-terminal mutations impair p53's ability to transactivate canonical targets like CDKN1A and BAX, potentially compromising tumor suppression[1](https://pubmed.ncbi.nlm.nih.gov/39520074/)[4](https://www.ncbi.nlm.nih.gov/pmc/articles/PMC10742156/). In myeloid neoplasms, TP53 variants with variant allele frequency (VAF) ≥10% correlate with adverse outcomes, though c.11C>T's clinical impact remains uncharacterized[2](https://www.semanticscholar.org/paper/21f319e06e955dd4b45f4ae31d1a077751bdea72)[5](https://www.semanticscholar.org/paper/3e049da7544c87a5ead0e0eebde4878e3c80bddd)[9](https://pubmed.ncbi.nlm.nih.gov/36655582/). The proximal location of this mutation suggests possible disruption of SP1 interaction sites implicated in non-canonical tumor suppression pathways[1](https://pubmed.ncbi.nlm.nih.gov/39520074/). Current classification systems consider TP53 mutation status critical for prognostication in hematologic malignancies, with multi-hit TP53 mutations (VAF ≥10% plus cytogenetic abnormalities) portending particularly poor survival[5](https://www.semanticscholar.org/paper/3e049da7544c87a5ead0e0eebde4878e3c80bddd)[7](https://www.ncbi.nlm.nih.gov/pmc/articles/PMC11788706/)[9](https://pubmed.ncbi.nlm.nih.gov/36655582/). While this variant meets ACMG criteria for pathogenicity (PS1, PM1, PM2, PP3), its functional consequences require validation through transcriptional activity assays and clinical correlation studies. Given the absence of specific therapeutic data, carriers should be monitored per guidelines for TP53 mutation-associated cancer risk. | [1](https://pubmed.ncbi.nlm.nih.gov/39520074/) PubMed 39520074  [2](https://www.semanticscholar.org/paper/21f319e06e955dd4b45f4ae31d1a077751bdea72) Semanticscholar 21f319e0  [4](https://www.ncbi.nlm.nih.gov/pmc/articles/PMC10742156/) PMC10742156  [5](https://www.semanticscholar.org/paper/3e049da7544c87a5ead0e0eebde4878e3c80bddd) Semanticscholar 3e049da7  [6](https://www.semanticscholar.org/paper/63eb7a38bc1d08d3860941c7ebcb93cb470eb17e) Semanticscholar e0656d06  [7](https://www.ncbi.nlm.nih.gov/pmc/articles/PMC11788706/) PMC11788706  [9](https://pubmed.ncbi.nlm.nih.gov/36655582/) PubMed 36655582 |
|  | Evaluation  **Summarization Accuracy & Completeness:**  **Hallucination Rate:**  **Readability & Usability:**  **Literature Coverage & Relevance:**  **Time Efficiency:** | |
| **5** | The TP53 gene encodes the tumor protein p53, a crucial transcription factor that maintains genomic stability by regulating cell cycle arrest, apoptosis, and DNA repair in response to cellular stress. The c.11C>T variant in TP53 results in a missense mutation at codon 4, substituting proline with leucine (p.Pro4Leu). This variant is absent from population databases, including gnomAD, indicating it is not a common polymorphism. Functional assays have demonstrated that p.Pro4Leu does not significantly impair p53's transactivation or growth suppression capabilities. In silico predictive models also suggest a benign effect on protein function. Based on this evidence, the ClinGen TP53 Variant Curation Expert Panel has classified this variant as likely benign for Li-Fraumeni syndrome as of January 16, 2025. However, due to limited clinical observations, continued surveillance and functional studies are recommended to conclusively determine its clinical significance. | <https://genomemedicine.biomedcentral.com/articles/10.1186/s13073-025-01429-5>  <https://www.nature.com/articles/s41598-024-67810-3>  <https://link.springer.com/article/10.1007/s10689-024-00391-2>  <https://exrna-atlas.org/evrepo/ui/classification/08a0ff2e-ec84-4336-94ff-fb468ba9dc88>  <https://jmg.bmj.com/content/jmedgenet/early/2020/10/13/jmedgenet-2020-107059.full.pdf>  <https://www.nature.com/articles/s10038-019-0681-6>  <https://www.ncbi.nlm.nih.gov/clinvar/RCV000229754/>  <https://www.ncbi.nlm.nih.gov/clinvar/RCV000759369.5/>  <https://www.ncbi.nlm.nih.gov/clinvar/variation/481148/>  <https://clinvarminer.genetics.utah.edu/variants-by-condition/Li-Fraumeni%20syndrome%201/gene/TP53/pathogenic>  <https://clinvarminer.genetics.utah.edu/variants-in-conflict-by-gene/TP53>  <https://clinvarminer.genetics.utah.edu/variants-by-gene/TP53/condition/Li-Fraumeni%20syndrome/likely%20benign>  <https://jmg.bmj.com/content/60/11/1057>  <https://aacrjournals.org/cancerres/article/77/6/1250/624858/Recommended-Guidelines-for-Validation-Quality>  <https://aacrjournals.org/cancerres/article-abstract/82/19/3420/709411/Benign-SNPs-in-the-Coding-Region-of-TP53-Finding?redirectedFrom=fulltext>  <https://www.explorationpub.com/Journals/em/Article/1001246>  <https://journals.lww.com/hemasphere/fulltext/2022/08000/five_percent_variant_allele_frequency_is_a.9.aspx> |
|  | Evaluation  **Summarization Accuracy & Completeness:**  **Hallucination Rate:**  **Readability & Usability:**  **Literature Coverage & Relevance:**  **Time Efficiency:** | |
