## Supplementary material for "Benchmarking generative AI tools for literature retrieval and summarization in genomic variant interpretation": TSC1 rs118203599.docx

| **Tool** | **Summary** | **References** |
| --- | --- | --- |
| **1** | The TSC1 gene provides instructions for producing hamartin, a protein that interacts with tuberin (produced by the TSC2 gene) to control cell growth, division, and size; hamartin and tuberin function as tumor suppressors[1](https://medlineplus.gov/genetics/gene/tsc1/). Variants in TSC1 are associated with tuberous sclerosis complex (TSC), lymphangioleiomyomatosis (LAM), and focal cortical dysplasia[1](https://medlineplus.gov/genetics/gene/tsc1/). TSC1-related TSC is characterized by developmental problems and benign tumors in multiple organs, resulting from a loss of functional hamartin[1](https://medlineplus.gov/genetics/gene/tsc1/). LAM, a destructive lung disease predominantly affecting women, can be caused by TSC1 variants, either sporadically or in combination with TSC; sporadic LAM may arise from mosaicism, where some cells have a normal TSC1 gene while others have an altered version[1](https://medlineplus.gov/genetics/gene/tsc1/). Focal cortical dysplasia of Taylor balloon cell type, another condition linked to TSC1 variants, involves brain malformations and severe epilepsy[1](https://medlineplus.gov/genetics/gene/tsc1/). The TSC1 gene is located on chromosome 9 (9q34)[3](https://www.lovd.nl/tsc1). As of February 14, 2025, 4681 variants have been reported for TSC1[3](https://www.lovd.nl/tsc1). Molecular diagnosis of TSC can be established with the identification of a heterozygous pathogenic variant in TSC1[5](https://www.ncbi.nlm.nih.gov/books/NBK1220/).  Absence of direct references for the clinical impact of the rs118203599 variant necessitates further investigation to ascertain its specific functional consequences and clinical relevance. | [1] <https://medlineplus.gov/genetics/gene/tsc1/>  [2] <https://www.lovd.nl/tsc1>  [3] <https://www.malattierare.eu/pages/rivista/La-sclerosi-tuberosa-idA63>  [4] <https://www.ncbi.nlm.nih.gov/books/NBK1220/> |
|  | Evaluation  **Summarization Accuracy & Completeness:**  **Hallucination Rate:**  **Readability & Usability:**  **Literature Coverage & Relevance:**  **Time Efficiency:** | |
| **2** | The TSC1 gene encodes hamartin, a protein involved in the regulation of the mechanistic target of rapamycin (mTOR) signaling pathway, which is crucial for cell growth, proliferation, and autophagy. Pathogenic variants in TSC1 are associated with tuberous sclerosis complex (TSC), a multisystem disorder characterized by benign tumor formation in multiple organs, including the brain, kidneys, heart, and skin. The rs118203599 variant of TSC1 is a single nucleotide polymorphism (SNP) whose clinical significance remains poorly characterized in the literature. Although mutations in TSC1 contribute to the pathogenesis of various renal and neurological disorders, specific evidence regarding the functional impact of rs118203599 is scarce. A study by Wu et al. (2019) [1] investigated germline mutations in TSC1 among patients with renal tumors and reported a second somatically acquired mutation in some individuals, suggesting a potential role in tumor predisposition. However, this study did not explicitly discuss rs118203599, leaving its pathogenicity unresolved. Further research is needed to determine its frequency in different populations, potential genotype-phenotype correlations, and functional consequences in mTOR signaling. Given the current uncertainty surrounding this variant, clinical interpretation should be approached with caution, and additional genetic and functional studies are recommended to clarify its role in disease. | [1] Wu J, Wang H, Ricketts CJ, Yang Y, Merino MJ. Germline mutations of renal cancer predisposition genes and clinical relevance in Chinese patients with sporadic, early-onset disease. *Cancer*. 2019. [DOI:10.1002/cncr.31908](https://app.scholarai.io/paper?paper_id=DOI:10.1002/cncr.31908&original_url=https%3A%2F%2Facsjournals.onlinelibrary.wiley.com%2Fdoi%2Fabs%2F10.1002%2Fcncr.31908). |
|  | Evaluation  **Summarization Accuracy & Completeness:**  **Hallucination Rate:**  **Readability & Usability:**  **Literature Coverage & Relevance:**  **Time Efficiency:** | |
| **3** | The genomic variant c.1905_1906del p.Glu636GlyfsTer51 rs118203599 is located on the TSC1 gene, which encodes hamartin, a protein that acts as a negative regulator of the mTOR pathway. The mTOR pathway is crucial for cell growth and proliferation, and its dysregulation can lead to tumorigenesis. The TSC1 protein, along with TSC2, forms a complex that inhibits mTORC1 signaling, thereby controlling cell size and preventing abnormal cellular proliferation.  The c.1905_1906del p.Glu636GlyfsTer51 variant results in a frameshift mutation, leading to a premature termination codon at the 51st amino acid downstream of the mutation site. This is expected to result in a truncated protein product that lacks normal function, which can disrupt the inhibitory action on the mTOR pathway. The variant has been identified in patients with focal cortical dysplasia (FCD), a malformation of cortical development associated with epilepsy. In one reported case, the variant was found with a variant allele frequency (VAF) of 3.7% in the brain tissue, indicating a somatic origin of the mutation [2].  In the context of bladder cancer, the TSC1 c.1905_1906del mutation has been reported with a frequency of 11.7%, suggesting a role in tumorigenesis in this tissue as well [3]. Moreover, patients with TSC1 mutations, including the c.1905_1906del variant, have been shown to have a longer duration of response to mTOR inhibitor treatment, such as Everolimus, compared to those with wild-type tumors [4].  The presence of the c.1905_1906del p.Glu636GlyfsTer51 rs118203599 variant in the TSC1 gene is consistent with the loss-of-function mechanism that is characteristic of pathogenic variants in this gene. Given the role of TSC1 in regulating the mTOR pathway, the loss of function due to this frameshift mutation is likely to contribute to the development of diseases associated with mTOR dysregulation, such as FCD and cancer. ClinVar has recorded this variant with two pathogenic submissions, supporting its significance in disease. | [[1]](https://pubmed.ncbi.nlm.nih.gov/31444548/) Baldassari S et al. (2019). "Dissecting the genetic basis of focal cortical dysplasia: a large cohort study." Acta neuropathologica, 138(6)  [[2]](https://pubmed.ncbi.nlm.nih.gov/38710875/) Ribierre T et al. (2024). "Targeting pathological cells with senolytic drugs reduces seizures in neurodevelopmental mTOR-related epilepsy." Nature neuroscience, 27(6)  [[3]](https://pubmed.ncbi.nlm.nih.gov/27352265/) Zhang X et al. (2015). "Bladder Cancer and Genetic Mutations." Cell biochemistry and biophysics, 73(1)  [[4]](https://pubmed.ncbi.nlm.nih.gov/28789352/) Zhang Z et al. (2017). "PI3K/AKT/mTOR pathway in pulmonary carcinoid tumours." Oncology letters, 14(2)  [[5]](https://pubmed.ncbi.nlm.nih.gov/22923433/) Iyer G et al. (2012). "Genome sequencing identifies a basis for everolimus sensitivity." Science (New York, N.Y.), 338(6104)  [[6]](https://pubmed.ncbi.nlm.nih.gov/30548481/) Wu J et al. (2019). "Germline mutations of renal cancer predisposition genes and clinical relevance in Chinese patients with sporadic, early-onset disease." Cancer, 125(7)  [[7]](https://pubmed.ncbi.nlm.nih.gov/29338072/) Ross JS et al. (2018). "Targeting HER2 in colorectal cancer: The landscape of amplification and short variant mutations in ERBB2 and ERBB3." Cancer, 124(7) Variant reported in Supplementary Information  [[8]](https://pubmed.ncbi.nlm.nih.gov/38225666/) Yang L et al. (2024). "Phase separation as a possible mechanism for dosage sensitivity." Genome biology, 25(1) Variant reported in Supplementary Information  [[9]](https://pubmed.ncbi.nlm.nih.gov/30140723/) Cravero K et al. (2018). "Biotinylated amplicon sequencing: A method for preserving DNA samples of limited quantity." Practical laboratory medicine, 12() Variant reported in Supplementary Information  [[10]](https://pubmed.ncbi.nlm.nih.gov/26503331/) Li C et al. (2015). "Whole Exome Sequencing Identifies Frequent Somatic Mutations in Cell-Cell Adhesion Genes in Chinese Patients with Lung Squamous Cell Carcinoma." Scientific reports, 5() Variant reported in Supplementary Information |
|  | Evaluation  **Summarization Accuracy & Completeness:**  **Hallucination Rate:**  **Readability & Usability:**  **Literature Coverage & Relevance:**  **Time Efficiency:** | |
| **4** | The TSC1 gene encodes hamartin, a tumor suppressor protein that interacts with tuberin (encoded by TSC2) to form a complex regulating cell growth and proliferation by inhibiting the mammalian target of rapamycin complex 1 (mTORC1) signaling pathway. This inhibition is crucial for controlling anabolic cell growth. Additionally, hamartin functions as a co-chaperone for Hsp90, modulating its ATPase activity and facilitating the folding of various client proteins, including TSC2, thereby preventing their degradation. Variants in TSC1 are associated with tuberous sclerosis complex (TSC), an autosomal dominant disorder characterized by the development of benign tumors in multiple organs, including the brain, skin, kidneys, heart, lungs, and retina. citeturn0search1  The specific variant rs118203599 corresponds to a two-nucleotide deletion at positions c.1907_1908 in the TSC1 gene, leading to a frameshift and premature stop codon (p.Glu636Glyfs*51). This frameshift is predicted to result in a truncated, non-functional hamartin protein, impairing its tumor suppressor function. citeturn0search0  This variant has been classified as pathogenic in ClinVar (Variation ID: 93429), with submissions from clinical testing laboratories supporting its association with TSC. citeturn0search0  The rs118203599 variant is located on chromosome 9 at position 132,905,670-132,905,671 (GRCh38 assembly). citeturn0search0  Given the pathogenic classification of the rs118203599 variant and its predicted impact on hamartin function, individuals harboring this variant are at an increased risk for developing TSC. Clinical management should include regular monitoring for TSC manifestations across various organ systems to facilitate early detection and intervention. Genetic counseling is recommended to discuss the implications of this variant, inheritance patterns, and potential risks to family members. | [1][GeneCardsTSC1 Gene - GeneCards \| TSC1 Protein \| TSC1 Antibody25](https://www.genecards.org/cgi-bin/carddisp.pl?gene=TSC1)  [2] [medlineplus.govTSC1 gene: MedlinePlus Genetics1 aprile 2022](https://medlineplus.gov/genetics/gene/tsc1/)  [3] [snpedia.comrs118203712(-;G) - SNPedia5 febbraio 2020](https://www.snpedia.com/index.php/Rs118203712%28-%3BG%29)  [4] [snpedia.com rs118203657 - SNPedia7 dicembre 2019](https://www.snpedia.com/index.php/Rs118203657)  [5] [acsjournals.onlinelibrary.wiley.comGermline mutations of renal cancer predisposition genes and ...12 dicembre 2018](https://acsjournals.onlinelibrary.wiley.com/doi/full/10.1002/cncr.31908)  [6] [ncbi.nlm.nih.govNM_000368.5(TSC1):c.1907_1908de](https://www.ncbi.nlm.nih.gov/clinvar/RCV000824628.8/)  [7] [ncbi.nlm.nih.govc.1579C>T (p.Gln527Ter)](https://www.ncbi.nlm.nih.gov/clinvar/RCV000660336/)  [8] [ncbi.nlm.nih.gov7248 -](https://www.ncbi.nlm.nih.gov/gene/7248)  [9] [pubmed.ncbi.nlm.nih.govRole of TSC1 in physiology and diseases](https://pubmed.ncbi.nlm.nih.gov/33575875/)  [10] [ClinVar MinerList of variants in gene TSC1](https://clinvarminer.genetics.utah.edu/variants-by-gene/TSC1/condition/Tuberous%20sclerosis%20syndrome)  [11] [ClinVar MinerList of variants in gene TSC1](https://clinvarminer.genetics.utah.edu/variants-by-mondo-condition/6026/gene/TSC1/not%20provided)  [12] [en.wikipedia.orgTSC1](https://en.wikipedia.org/wiki/TSC1)  [13] [thermofisher.comProduct](https://www.thermofisher.com/order/genome-database/details/pcr-sequencing-primers/Hs00626885_CE)  [14] [gedipnet.bicnirrh.res.inComprehensive Gene Data - OMIM, HGNC, Ensembl](https://gedipnet.bicnirrh.res.in/GgeneT.php?GI=7248&gens=TSC1)  [15] [UniProtTSC1 - Hamartin - Homo sapiens (Human) \| UniProtKB](https://www.uniprot.org/uniprotkb/Q92574/entry) |
|  | Evaluation  **Summarization Accuracy & Completeness:**  **Hallucination Rate:**  **Readability & Usability:**  **Literature Coverage & Relevance:**  **Time Efficiency:** | |
| **5** | The TSC1 gene, located on chromosome 9q34.13, encodes the protein hamartin, which functions as a tumor suppressor. Hamartin interacts with tuberin, the product of the TSC2 gene, forming a complex that regulates cell growth and proliferation by inhibiting the mTORC1 signaling pathway. This complex acts as a GTPase-activating protein (GAP) for the small GTPase Rheb, which is essential for mTORC1 activation. The TSC1-TSC2 complex negatively regulates mTORC1, preventing excessive cell growth and division. Mutations in TSC1 can lead to the autosomal dominant disorder tuberous sclerosis complex (TSC), characterized by the development of benign tumors in multiple organs, particularly the brain, skin, kidneys, heart, and lungs. TSC is associated with neurological manifestations, including seizures, intellectual disability, and autism spectrum disorder. The severity of TSC can vary widely, even within families, due to the variable expression of the mutated gene. The TSC1 protein, hamartin, also functions as a co-chaperone for Hsp90, facilitating the folding of kinase and non-kinase clients and preventing their ubiquitination and proteasomal degradation. This role is crucial for maintaining the stability and function of various proteins involved in cellular processes. The TSC1 gene is highly conserved across species, and its function is essential for normal cellular physiology. Loss of function mutations in TSC1 can lead to uncontrolled cell growth and the formation of hamartomas, benign tumor-like growths composed of a mixture of normal and abnormal tissues. These hamartomas can occur in various organs and contribute to the clinical manifestations of TSC. The frequency of TSC1 mutations in the population is not well-documented, but it is estimated that TSC affects approximately 1 in 6,000 individuals. The majority of TSC cases are sporadic, resulting from de novo mutations, while familial cases account for about one-third of all cases. The specific mechanisms by which TSC1 mutations cause disease are not fully understood, but it is suggested that at least two hits are necessary for tumor formation, as TSC1 and TSC2 are tumor suppressor genes.  In summary, the TSC1 gene plays a critical role in regulating cell growth and proliferation through its interaction with TSC2 and the mTORC1 signaling pathway. Mutations in TSC1 can lead to tuberous sclerosis complex, a disorder characterized by the development of benign tumors in multiple organs and various neurological manifestations. The function of TSC1 as a tumor suppressor and co-chaperone is essential for maintaining normal cellular physiology and preventing uncontrolled cell growth. | [1] MedlinePlus Genetics. TSC1 gene. Available at: <https://medlineplus.gov/genetics/gene/tsc1/> |
|  | Evaluation  **Summarization Accuracy & Completeness:**  **Hallucination Rate:**  **Readability & Usability:**  **Literature Coverage & Relevance:**  **Time Efficiency:** | |
